# Environmentally relevant bisphenol mixtures modify heart–brain physiology, cerebrovascular structures, and transcriptomic profiles in zebrafish larvae

**DOI:** 10.64898/2026.10.06.752046

**Authors:** Shana Lallemand, Sandrine Ellero-Simatos, Laurence Gamet-Payrastre, Gaëlle Payros, Yannick Lippi, Océane Lassié, Marine Blaquière, Alicia Janvier, Adèle Faucherre, Aurélien Drouard, Angelo G. Torrente, Chris Jopling, Nicola Marchi, Ieremias Chousidis

## Abstract

Environmental exposure involves pollutant mixtures. Building on evidence that individual bisphenols alter zebrafish larval physiology and transcriptomic profiles at ng/L concentrations, we assessed the effects of specific mixtures of BPS, BPF, and BPAF without (M1) or with (M2) BPA. Experimentally, M1 and M2 induced 4.16% and 6.54% mortality in zebrafish larvae, despite the concentration addition model predicting no toxicity. Both mixtures reduced swimming-locomotor activities. These behaviors were accompanied by organ-level functional defects, as *in vivo* optical mapping revealed bradycardia and increased high-frequency (30-80 Hz) brain oscillations for both mixtures, while M2 additionally altered low-frequency (0.2-4 Hz) brain activity. The neurophysiological changes were accompanied by neurovascular remodeling, with *in vivo* two-photon microscopy revealing decreased vascular length and branching. Finally, transcriptomic analysis revealed common M1- and M2-induced alterations linked to pathways of vascular function, neuronal communication, and metabolism. M2 altered additional pathways of cardiac and neuronal signaling, consistent with the reported organ-physiology effects. We discuss the environmental relevance of mixture exposures and assess whether BPA intensifies mixture-induced responses.

**Graphical abstract:** 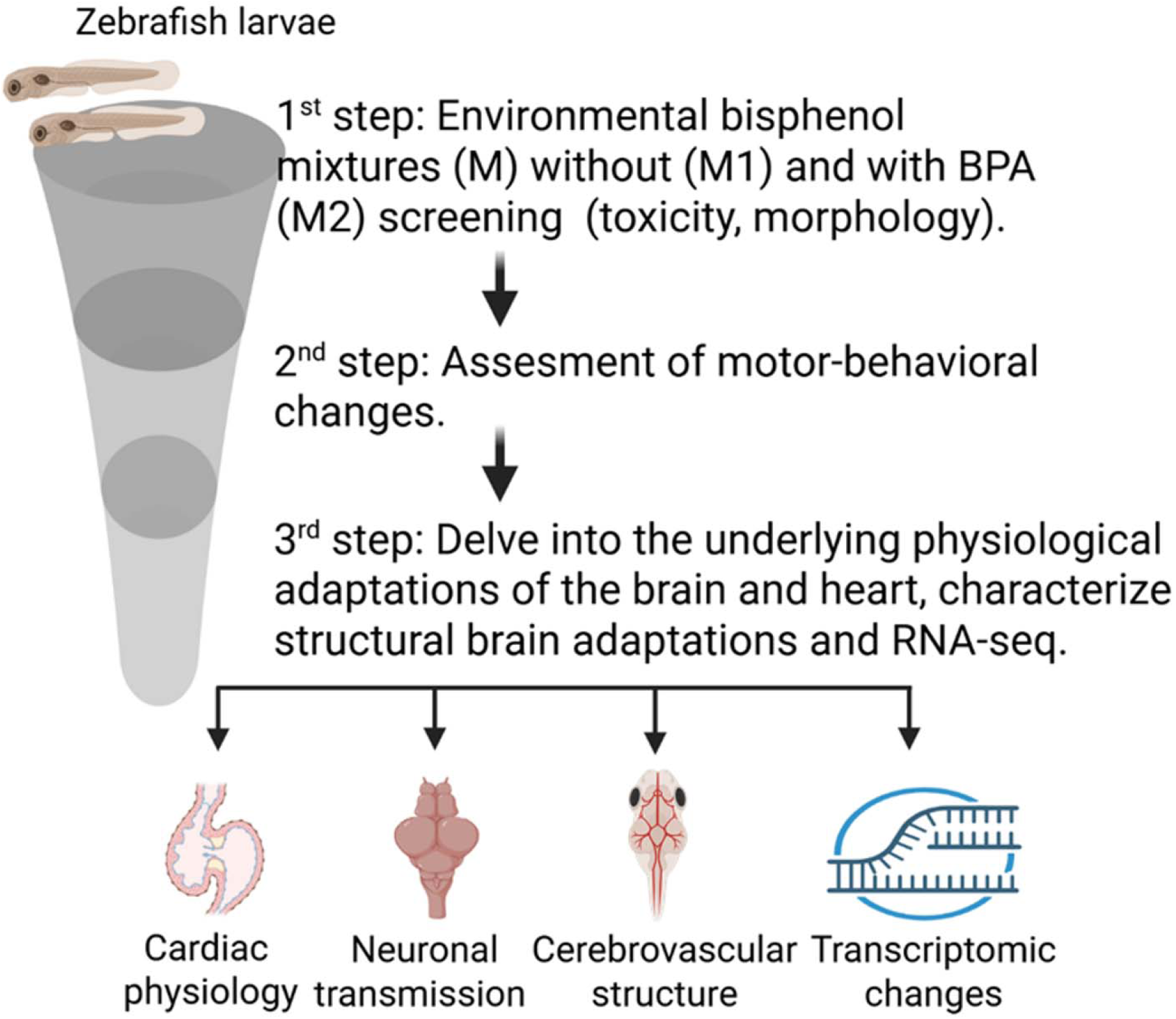

## 1. Introduction

Bisphenols are synthetic industrial chemicals that are widely used in the production of polycarbonate plastics and epoxy resins [1]. Their presence has raised significant concern for human health and environmental safety. Regulatory restrictions on bisphenol A (BPA) have been accompanied by the introduction of structural analogues (e.g., BPS, BPF, BPAF), intended to curb health and environmental risks [2–4]. However, bisphenol analogues are not entirely safer alternatives to BPA. Epidemiological studies have linked exposure to BPS, BPF, BPAF, and other analogues with adverse metabolic, reproductive, cardiovascular, and neurodevelopmental outcomes [5–7]. Studies in zebrafish have unveiled endocrine-disrupting, neurobehavioral, and cardiotoxic effects [8, 9]. In this context, we previously demonstrated that, in zebrafish larvae, individual BPS, BPF, BPAF, or BPA at environmentally relevant levels (ng/L) induced heart-brain extraphysiological adaptations along with transcriptomic changes [10].

Environmental exposure involves bisphenol mixtures, yet most studies focus on single compounds [11]. Some evidence indicates that bisphenol mixtures can induce additive cardiac or developmental toxicity in zebrafish embryos [12]. The reported effects were observed under acute exposure conditions and at concentrations exceeding those found in aquatic environments [12]. The latter creates a knowledge gap in ecotoxicology and risk assessment. Here, we examined physiological and molecular responses to environmentally relevant concentrations (ng/L) of a BPS, BPF, and BPAF mixture, with or without BPA. Specific concentrations (100 ng/L BPS, 100 ng/L BPAF, 10 ng/L BPF, BPA 100 ng/L) were selected based on our previous work [10]. When tested individually, these concentrations were below the LD_50_, non-lethal, and represented the lowest and environmental exposure levels at which functional and transcriptomic adaptations occur [10, 11, 13–15]. We employed the OECD Fish Embryo Acute Toxicity Test guidelines (Directive 2010/63/EU) [16]. We determined that mixture exposure elicits effects across multiple levels of biological organization, from acute toxicity and developmental morphology to locomotor behavior, cardiac and brain electrophysiology, neurovascular structure assessed by in vivo two-photon microscopy, and transcriptomic signatures that may underlie these physiological effects. We discuss these findings in the context of mixture composition and the presence of BPA and, where possible, compare the observed responses with those elicited by individual bisphenols under the same experimental framework [10].

## 2. Materials and Methods

### 2.1 Chemicals

Analytical-grade bisphenol A (BPA, 99%, CAS 80-05-7), bisphenol S (BPS, 98%, CAS 80-09-1), bisphenol F (BPF, 99%, CAS 620-92-8), and bisphenol AF (BPAF, 99%, CAS 1478-61-1) were purchased from Sigma-Aldrich (France). Each compound was separately dissolved in ethanol (70%) and further diluted in 50 mL E3 culture medium to prepare 0.1 mg/mL stock solutions (EtOH concentration in each stock solution ≤ 0.1% v/v). Stocks were stored at -20°C and renewed weekly to ensure stability. For each bisphenol mixture, we prepared a corresponding ethanol vehicle control to match the ethanol content. We assessed vehicle toxicity (n=84/condition, triplicate) and found no significant differences between the E3 culture medium control and the vehicle control (Supporting Information, Table S1).

### 2.2 Zebrafish husbandry

Zebrafish were maintained under standardized conditions, and experiments were conducted in accordance with local approval (APAFIS#4054-2016021116464098 v5) and European Communities directive 2010/63/EU. Sexually mature (3-month-old) zebrafish were used for spawning, and collected embryos were pooled in E3 culture medium. For testing, larvae (up to 5 days or 120 h post-fertilization) were used. Tricaine methanesulfonate (MS222, Sigma-Aldrich, France) was used for anesthesia or euthanasia.

### 2.3 Toxicological, morphological, and functional screening of bisphenol analogue exposure

At 1.5 hours post-fertilization (hpf), embryos were placed in 24-well plates (2 embryos per well, 2 mL solution per well). Exposures lasted 96 hours, ending at 5 days post-fertilization (dpf). We renewed solutions daily and rinsed embryos twice with E3 culture medium after exposure to remove residual chemicals. The procedure followed OECD guidelines [16]. We exposed zebrafish larvae to two environmentally relevant bisphenol mixtures. Mixture 1 (M1) consisted of the lowest concentrations of each analogue that provoked significant biological alterations in our previous study (100 ng/L BPS, 100 ng/L BPAF, and 10 ng/L BPF). Mixture 2 (M2) consisted of M1 supplemented with 100 ng/L BPA. Comparing the two mixtures, one containing BPA and one without, enabled us to systematically investigate whether the inclusion of BPA enhances the biological effects of bisphenol mixtures. To determine the nature of the interactions between chemicals within the mixtures, we calculated the expected acute toxicity of each bisphenol mixture using the Concentration Addition model (CA). This model assumes that chemicals share a common toxic mode of action and contribute additively to the overall mixture toxicity [17–19]. This enables us to compare the predicted and observed toxicity, providing insight into whether the combined effects deviated from concentration addition. Under this model, the contribution of each compound is expressed as a toxic unit (TU), calculated as the ratio of its exposure concentration to its median lethal concentration (LD_50_):

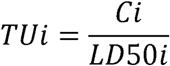

where (Ci) is the concentration of compound i in the mixture and (LD_50_i) is the experimentally determined LD_50_ of the same compound, as determined in our previous study [10].

The overall toxic pressure of each mixture was then calculated as the sum of the individual toxic units:

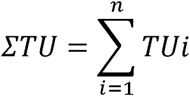

The LD_50_ values used for the calculations were 16.024 mg/L for BPS, 1.280 mg/L for BPAF, 2.113 mg/L for BPF and 8.710 mg/L for BPA (Informed by our previous study [10]).

For Mixture 1 (100 ng/L BPS, 100 ng/L BPAF, and 10 ng/L BPF), the calculated toxic units were 6.24×10^−6^, 7.81×10^−5^, and 4.73×10^−6^, respectively, resulting in a total ΣTU of 8.91×10^−5^. For Mixture 2 (100 ng/L BPS, 100 ng/L BPAF, 10 ng/L BPF, and 100 ng/L BPA), the additional toxic unit contributed by BPA was 1.15×10^−5^, yielding a total ΣTU of 1.01×10^−4^. Accordingly, the calculated ΣTU values indicate that Mixtures 1 and 2 corresponded to 0.0089% and 0.0101%, respectively, of the concentration expected to produce 50% mortality under an additive model. Therefore, both mixtures were predicted to exhibit negligible acute toxicity (∼0%).

After toxicity determination, we examined the possible presence of embryonic malformations in M1- and M2-exposed larvae. Morphological parameters, including total body length (TL, mm), swim bladder area (SWB, mm²), and eye diameter (ED, mm), were analyzed at 5 dpf using a stereoscope equipped with a camera (Olympus SZX16, ProgRes CapturePro 2.8.8 software) (n=10/group, duplicate). Hatching rates (%) were also recorded (n=224/group, triplicate). Muscle formation defects were further assessed via phalloidin staining, as described [10] (n=10/group). The effects of bisphenol mixtures were evaluated by functional screening of zebrafish larvae using two tests at the end of the exposure: (1) Basal Locomotor Activity (BLA) and (2) Vibrational Startle Response (VSR). Before behavioral testing, larvae were screened to confirm the absence of morphological malformations after 5 days of exposure. This ensured that observed differences in swimming activity were not due to physical impairments. For the BLA test, larvae were subjected to alternating light-dark phases for 30 minutes after a 10-minute acclimation period (10 min dark-10 min light-10 min dark, see Fig. 2A). This protocol assessed the larvae’s ability to adapt to baseline environmental conditions [20]. Each larva was individually placed into a 24-well plate and transferred to the DanioVision Observation Chamber (Noldus Information Technology, VA, USA) equipped with a temperature control unit. Locomotor activity was tracked using an infrared camera (25 frames/sec, Basler acA1300-60gm, Basler Inc., PA, USA) and movement parameters (distance traveled [cm]) were analyzed (n= 24/group, duplicate; see Fig. 2A-C). The VSR test evaluated locomotor activity, sensory organ function, and stress–like behaviors [21]. A mechanical tapping stimulus was applied using a piston in the behavior recording system (DanioVision Tapping Device DVOC-004x/T, Noldus, The Netherlands). After a 10-minute acclimation period, a 6-minute video recording (60 frames/sec) was conducted in complete darkness. The tapping stimulus was delivered at the 3-minute mark (180 sec). The response was analyzed by measuring (1) the distance traveled (cm) within 3 minutes post-stimulus and (2) the latency to respond to the stimulus (time to first movement, sec) (n= 24/group, duplicate; see Fig. 2E-F). Both behavioral tests (BLA and VSR) were conducted between 10:00 AM and 12:00 PM to account for circadian rhythms. Background noise interference was minimized using a 10% body-length filter (∼0.4 mm).

### 2.4 In vivo neuro and cardiac electrophysiological screening by optical mapping

At the end of each exposure, zebrafish larvae were incubated in blebbistatin (8.5 μM prepared in E3 culture medium) (Selleckchem, Cologne, Germany) for 30 minutes to prevent muscle contractions. Larvae were then incubated in the dark for 40 minutes with the voltage-sensitive probe Di-4-Anepps (9 μM prepared in E3 culture medium) (AAT Bioquest, Pleasanton, CA, USA). Larvae were immobilized in 35 mm glass-bottom petri dishes using low-melting agarose (LMA, Sigma-Aldrich, France) and positioned laterally for cardiac recordings and dorsally for cerebral recordings, as shown in our previous work [22]. To maintain viability and prevent muscle contractions during recordings, we added 2 ml of E3 culture medium with blebbistatin (8.5 μM) to each Petri dish. Each 35 mm Petri dish was positioned under the microscope setup within a warming chamber (QE-1, Warner Instruments, Holliston, MA, USA), connected to a dual automatic temperature controller (TC-344B, Warner Instruments, Holliston, MA, USA), and maintained at a constant temperature of 28°C. Cardiac and cerebral recording acquisitions (30 sec) were performed with an acquisition frequency of 500 frames/sec using a 150W halogen lighting system (SciMedia, Costa Mesa, CA, USA) as an excitation source coupled with a 531/50 nm excitation filter, a 580 nm dichroic mirror, and a 580 long-pass emission filter. The optical mapping system comprised an SDF PLAN FLUOR 0.3X stereo microscope objective mounted as an eyepiece lens (Olympus, France), a Plan Apo 5.0X/0.50 LWD objective mounted as a magnification lens (Leica, France), and a MiCAM03 optical camera with a 17.6 × 17.6 mm sensor (SciMedia, Costa Mesa, CA, USA). This configuration provided a field of view of 1.05 × 1.05 mm corresponding to 256 × 256 pixels, as described [22]. We used BV Workbench software (SciMedia 4.6, Costa Mesa, CA, USA) to perform the acquisitions. Cardiac depolarization was recorded as a fluorescence change in the atrial region, corresponding to cyclical heart rate traces (See Fig. 3A). Brain depolarization was recorded by capturing the cumulative fluorescent signal (See Fig. 4A). Fluorescent signals deriving from the heart were extracted and analyzed using ClampFit 11.3 (Molecular Devices, LLC, San Jose, CA, USA). We assessed the effects of bisphenol analogs by analyzing heart rate (beats per minute, BPM) and the coefficient of variability of the interval between two heartbeats (coefficient of variability, % CV), recorded as optical action potentials in the atrial region (n = 8/condition, executed in duplicate). For brain field voltage activity, signals were recorded from the brain region shown in Fig. 4A. Signals from this region were exported to OriginPro software for Fast Fourier Transform (FFT) analysis (OriginPro 2018, Northampton, MA, USA; n = 10/condition, executed in duplicate) to examine low-amplitude voltage frequency patterns. We assessed FT frequency ranges of 0.2–4 Hz, 4.1 - 7 Hz, and 30 - 80 Hz to allow calculations (Nyquist sampling law [23]) with the sampling rate used (500 frames/s). To minimize cardiac signal interference, frequencies between 2–3 Hz were removed from the analyses. The recordings were visually inspected for high-amplitude spike activities, while artifacts were excluded. In Fig. 4B, the Y-axis units in brain activity plots are derived from the mathematical FT elaboration, which indicates the relative abundance of low-amplitude waves in the targeted frequency ranges and over the recorded time. This solution analyzes the frequency range of activities emerging from excitable cells [24].

### 2.5. In vivo two-photon neuro-glio-vascular analysis

*Tg(fli1a:GFP)y1Tg:Tg(mpeg1:mCherry)* zebrafish larvae, labeling endothelial cells and microglia/macrophages, were used for the analysis. Larvae from the CTRL, M1, and M2 groups (n = 10/group) were anesthetized with tricaine (0.01%, 100mg/L) and immobilized in a drop of low-melting-point agarose. N-phenylthiourea (PTU) was added daily to the E3 medium from 24 to 96 hpf to prevent pigmentation, to a final concentration of 0.002 mL PTU/mL E3. In vivo imaging of zebrafish was performed using a FVMPE-RS two-photon microscope (Evident) equipped with an InSight X3 femtosecond-pulsed infrared laser (Spectra-Physics) and a Ti:Sapphire laser (Chameleon Vision2, Coherent) in a collimated configuration. Imaging was performed using a 25×, numerical aperture 1.05, water-immersion objective (XLPLN25XWMP2, Evident). The laser wavelength was tuned to 1040 nm to simultaneously excite GFP and DsRed, enabling faster acquisition, reduced channel misalignment, and simpler image processing. Fluorescence emission was collected using bandpass filters at 495–540 nm for GFP and 580–638 nm for DsRed. Whole brains were imaged by Z-stack acquisition (z-step = 1 µm). 3D cerebrovascular maps were generated using IMARIS 10.0 (Oxford Instruments) from z-stack images (512 × 512 pixels). We selected the midbrain as the region of interest (ROI), specifically the area between the metencephalic artery (MtA) and anterior cerebral vein (ACeV), following the annotations of the Interactive atlas of zebrafish anatomy (Fig. 5A)[25]. The following parameters were quantified in three dimensions: (i) cerebrovascular length and volume; (ii) the number of vascular branches and terminal segments; and (iii) microglial number. Microglia were quantified within the predefined midbrain ROI, corresponding to the region located between the metencephalic artery (MtA) and the anterior cerebral vein (ACeV). The ROI was subdivided into three equally sized adjacent subregions to ensure systematic and representative sampling across the entire midbrain ROI while minimizing potential bias arising from local heterogeneity in microglial distribution (Fig. 5C). Microglia were manually counted (by a blinded operator) within each subregion, and the counts were summed to obtain a single value per larva (Fig. 5I).

### 2.6 RNA sequencing and analysis

A pool of n = 50 zebrafish larvae (5 dpf) constituted one sample. The experimental conditions included one control group, M1, and M2. Each condition was analyzed in six replicates, for a total of 18 samples and 900 larvae. Total RNA was extracted using Trizol (Invitrogen) and sequenced at the GeT-TRiX platform at the TOXALIM Institute in Toulouse, France. For each sample, RNAseq libraries were constructed from 500 of total RNA at the GeT-TRiX facility (GénoToul, Génopole Toulouse Midi-Pyrénées) using QuantSeq 3’ mRNA-Seq V2 Library Prep Kit with UDI (Lexogen) following the manufacturer’s instructions. Libraries were amplified with fourteen cycles of PCR. The yield and the quality of the libraries were assessed using D1000 ScreenTapes on TapeStation 4200 (Agilent Technologies, Santa Clara). The libraries were then pooled to equimolar concentrations and transferred to INRAE PGTB sequencing facility for sequencing. The library pool was loaded into one P2 Flow Cell on Illumina NextSeq 2000 using 1x76 bp single read sequencing mode with NextSeq 1000/2000 P2 XLEAP-SBS Reagent kit (100 Cycles). The GeT-TRiX facility (GénoToul, Génopole Toulouse Midi-Pyrénées) performed bioinformatics processing of the sequencing data using the computing resources of the Genotoul bioinformatics platform Toulouse Midi-Pyrénées (Bioinfo Genotoul). The analysis pipeline was executed with Nextflow v25.10.0 [26] and and processed using nf-core/rnaseq v3.19.0 [27], from the nf-core collection of workflows [28] utilizing reproducible software environments from the Bioconda [29] and Biocontainers [30] projects. Reads were aligned to the Zebrafish(*Danio rerio*) genome reference GRCz11 (build GCA_000002035.4) and the quantification of reads was resumed to genes defined in Ensembl release 116 annotation.

Biostatistical analyses were performed in R 4.3.0 (R Core Team, 2023) using edgeR [31], limma [32], and Bioconductor packages [33]. Briefly, the raw count table was filtered to eliminate undetected genes using filterByExpr function with min.count = 10. Normalization factors were calculated using TMM method. Then, the common, trended, and tagwise negative binomial dispersions were estimated using the robust estimateDisp function. Experiment date was included in the design matrix as a fixed blocking factor. A negative binomial generalized log-linear model was fitted to count data with the quasi-likelihood (QL) methods using glmQLFit function[34]. Pairwise comparisons between biological conditions were extracted using specific contrasts with glmQLFtest function to identify differential expression. A correction for multiple testing was applied using the Benjamini-Hochberg procedure [35] to control the False Discovery Rate (FDR). Genes with FDR ≤ 0.05 were differentially expressed between conditions.

Hierarchical clustering was applied to samples and genes using the normalized logCPM expression values of the selected genes (Pvalue < 0.001 / VIP > 1). Pearson correlation (1 - cor) was used as the distance, and Ward’s criterion was used for agglomeration. Gene set enrichment analysis (GSEA) was applied to the full list of log fold changes for each comparison using the gseGO and gseKEGG functions. RNA-Seq data are available in the ArrayExpress database (http://www.ebi.ac.uk/arrayexpress) under accession number E-MTAB-17635.

### 2.7 Statistical analyses

We performed statistical analyses using GraphPad Prism 10.4.1. When the data met the assumption of normality, we used parametric tests such as one-way ANOVA followed by Dunnett’s multiple comparisons test. Otherwise, we used Kruskal-Wallis (non-parametric) followed by Dunn’s multiple comparisons test (p < 0.05). Results are presented using violin plots. The cumulative hatching rate was analyzed using two-way ANOVA followed by Dunnett’s multiple comparisons test (p < 0.05). Asterisks indicate statistical differences compared to the control group (CTRL): * (p < 0.05), ** (p < 0.01), *** (p< 0.001), **** (p < 0.0001).

## 3. Results

### 3.1. Expected and experimental toxicity and anatomical screening following low-level bisphenol mixture exposure

First, we assessed whether exposure to the selected bisphenol concentrations, shown to be non-toxic individually [10], produces an additive or synergistic effect as a mixture. Because the tested bisphenols belong to the same chemical class and are predicted to share similar modes of action, we employed the concentration addition (CA; see Methods) model. The calculated summed toxic units (ΣTU) were 8.91 × 10⁻ for Mixture 1 and 1.01 × 10⁻ for Mixture 2, corresponding to values approximately four orders of magnitude below one toxic unit; expected mortality was 0% for both mixtures. However, experimentally, we observed mortality of 4.16% in M1 and 6.54% in M2 at 5 dpf (Fig. 1A). Although these values were low, the mixture produced greater toxicity than the model predicted. Vehicle controls did not induce toxicity (Supplemental Table S1). This indicates that both mixtures produce effects that differ from those expected under an additive model.

**Figure 1.**
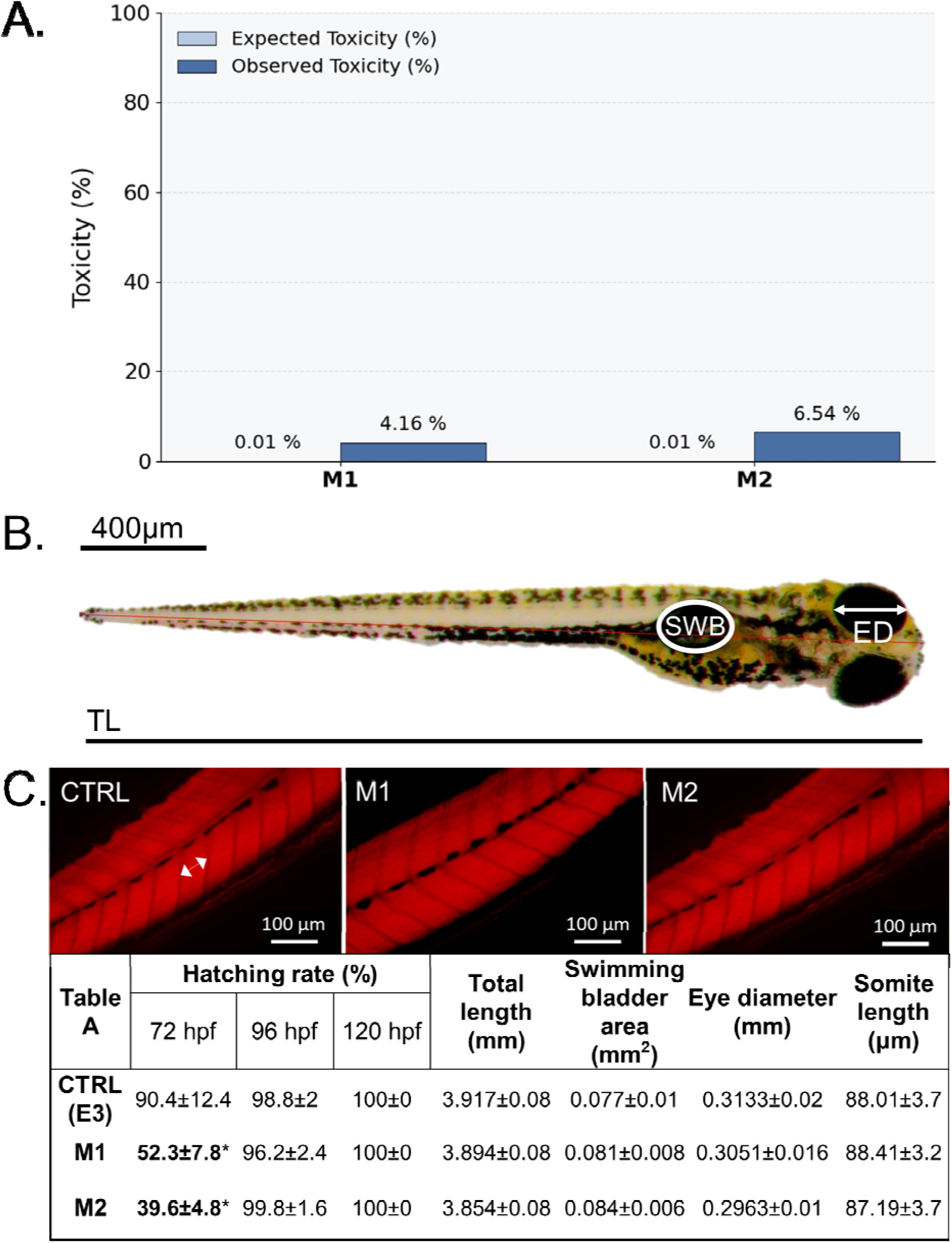
Bisphenol mixtures induce toxicity exceeding concentration addition model predictions and in the absence of morphological and muscular modifications. (A) Bar-plot showing the predicted versus observed toxicity upon exposure to bisphenol mixtures; (B) Examples of measurements for morphological assessment: Total body length (TL, μm), swimming bladder (SWB, μm2), eye diameter (ED, μm) of zebrafish larvae at 5 dpf. (C) Lateral views of trunk somites in zebrafish larvae stained using phalloidin. Dashed lines with arrows indicate the length of a typical muscle fiber measured. Table (A): statistically significant changes only in hatching rates at 72hpf. All other parameters remained statistically insignificant. As shown in Supplemental Table 1, controls with matching ethanol content and E3 exhibit no statistically significant differences in the tested parameters. For morphology measurements, data are reported as means ± SD (n = 15/group, duplicate). Cumulative hatching rates consecutively assessed at 72, 96, and 120 hpf are expressed as hatched eggs / total x 100 (experiments conducted in triplicate, n = 224/group; 2-way ANOVA, Dunnett’s multiple comparisons test, p < 0.05). Muscle measures were reported as means of somite length (µm, n = 10/group, Two-way ANOVA, Dunnett’s multiple comparisons test, p < 0.05).

We next examined whether exposure to bisphenol mixtures was associated with developmental and morphological alterations. At 72 hpf, hatching rates were significantly delayed after exposure to M1 and M2 compared with controls. Total length (TL), swim bladder area (SWB), and eye diameter (ED) showed no statistically significant changes after mixture exposure compared with controls. (See Fig. 1, Table A). Furthermore, phalloidin immunostaining ruled out muscle deformities at 5 dpf (Fig. 1B, Table A). Vehicle controls did not significantly alter the parameters analyzed (Supporting Information, Table S1). Overall, M1 and M2 induced low but above-predicted toxicity and no overall morphological abnormalities.

### 3.2 Swimming-motor alterations following low-level bisphenol mixture exposure

Next, we investigated whether exposure to bisphenol mixtures modifies swimming-motor behaviors, in the absence of morphological abnormalities. Fig. 2A shows the paradigm used, and Fig. 2B shows the capacity of 5 dpf zebrafish larvae to adapt to alternating light and dark conditions. Exposure to M1 and M2 induced a statistically significant reduction in distance traveled (cm) under all conditions (dark/light) compared to control. No statistically significant changes were observed between M1 and M2 (Fig. 2B). Exposure to vehicle controls (EtOH) did not result in statistically significant alterations compared to E3 medium (Fig. S1). We also examined the startle response using a mechanical tapping stimulus. Fig. 2D illustrates the experimental paradigm used. We report no statistically significant differences in the latency to the first movement following the mechanical stimulus in larvae exposed to bisphenol mixtures compared to controls (Fig. 2F). However, larvae exposed to M1 and M2 exhibited a significant reduction in the total distance traveled after the tapping stimulus compared to controls (Fig.2E), with no significant differences observed between M1 and M2 groups. Overall, the swimming-motor alterations were observed regardless of the presence of BPA in the mixture.

**Figure 2.**
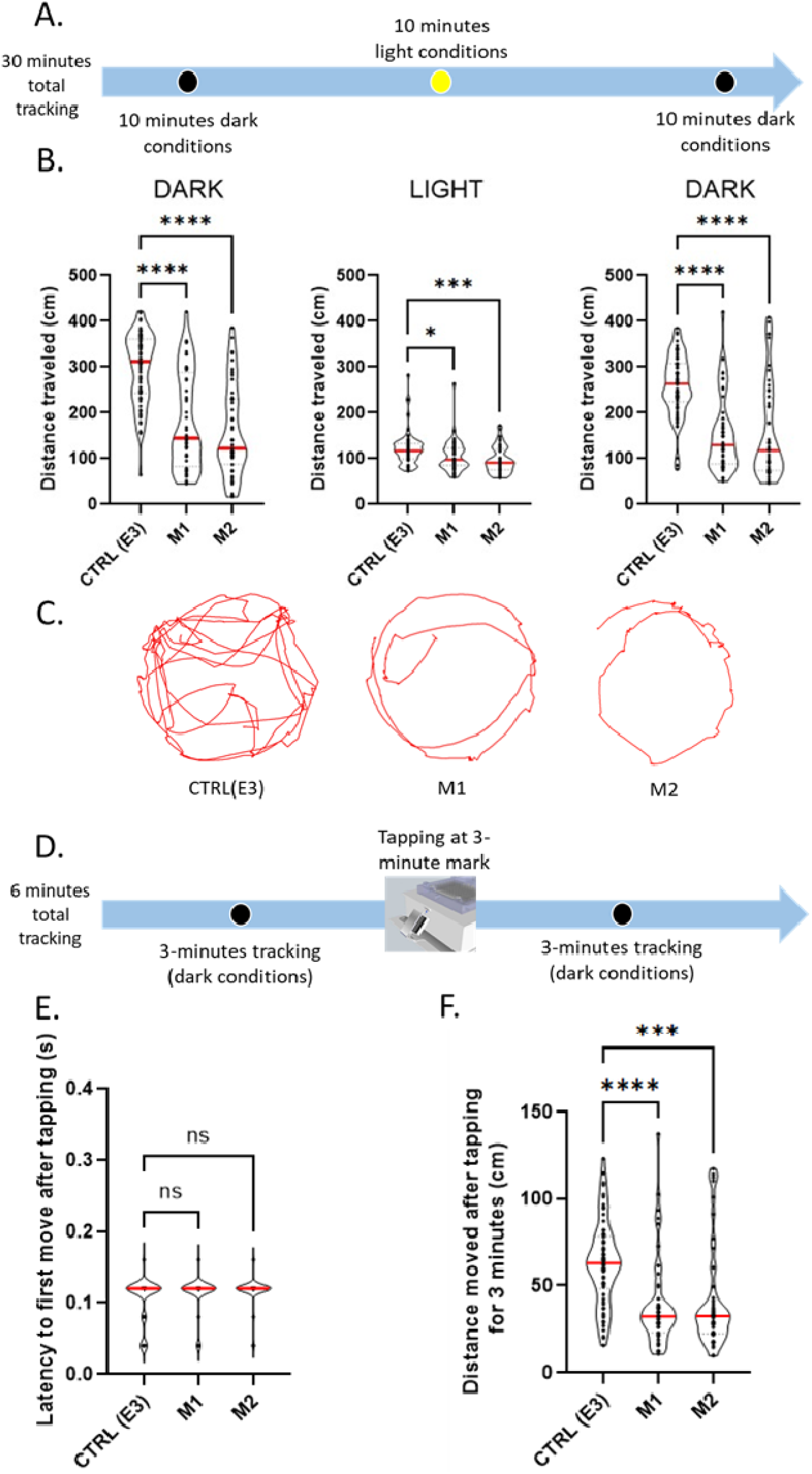
Environmentally relevant exposure to bisphenol mixtures alters locomotor activity patterns in both dark and light conditions and after mechanical stimulation in zebrafish larvae at 5 dpf. (A) Protocol utilized to identify locomotor activities and changes. CTRL refers to E3. (B) Locomotion outputs obtained during the 30 min tracking and according to light/dark conditions for mixtures tested (n = 24 zebrafish each condition, executed in duplicate; non-parametric test, Kruskal-Wallis, Dunn’s multiple comparisons test, p < 0.05). (C) Examples of motor-activity for each condition during 30 seconds in dark conditions. (D) Protocol of vibrational startle response test to examine reflex modifications potentially triggered by M1 and M2 exposure. CTRL refers to E3. (E) Latency to first movement after tapping for M1 and M2. (F) Total distance moved for 3 min after tapping with exposure to M1 and M2. For the latency dataset, we report means ± SD (n = 24/group), executed in duplicate; non-parametric tests (Kruskal-Wallis) with Dunn’s multiple-comparisons test; p < 0.05. As shown in Supplementary Figure S1, controls with matching ethanol content and E3 exhibit no statistically significant changes in motor behavior.

### 3.3 Cerebral and cardiac physiological adaptations to low-level bisphenol mixture exposure in zebrafish larvae

Guided by the observed changes in locomotor behavior in the absence of gross morphological alterations, we investigated whether these functional deficits were associated with changes in organ physiology, focusing on the heart and brain. We used optical mapping [22] to non-invasively image the spatiotemporal and anatomical propagation of field voltage activity *in vivo* (Fig. 3A-B). Compared to controls (CTRL E3 medium), exposure to M1 and M2 resulted in a statistically significant reduction in heart rate (bradycardia) (Fig. 3C), whereas the coefficient of variability of heart rate remained unchanged, suggesting the absence of cardiac arrhythmias (Fig. 3D). We next examined spatio-temporal patterns of neuronal activity [45] in the forebrain (Fig. 4). Using spectrogram analyses, we report changes in the abundance of low- and high-frequency oscillations (delta 0.2-4Hz and gamma 30-80 Hz, respectively, Fig. 4C, Fig. 4E), indicating adaptations in neuronal activity to both bisphenol mixtures. Specifically, M1 provoked a trend toward an increase in gamma-wave abundance (p = 0.067). M2 significantly increased the abundance of both delta and gamma waves, with the stronger effect likely attributable to the presence of BPA. No statistically significant changes were observed in the other frequency ranges analyzed (4.1-7 Hz, 5-15 Hz, and 15-30 Hz, Fig. S4). Vehicle controls showed no alterations in either cardiac or brain function (Fig. S2, S3). Our results showed that both mixtures induced a shared pattern of cardiac physiological alterations, while the presence of BPA in M2 was associated with additional effects on neuronal activity.

**Figure 3.**
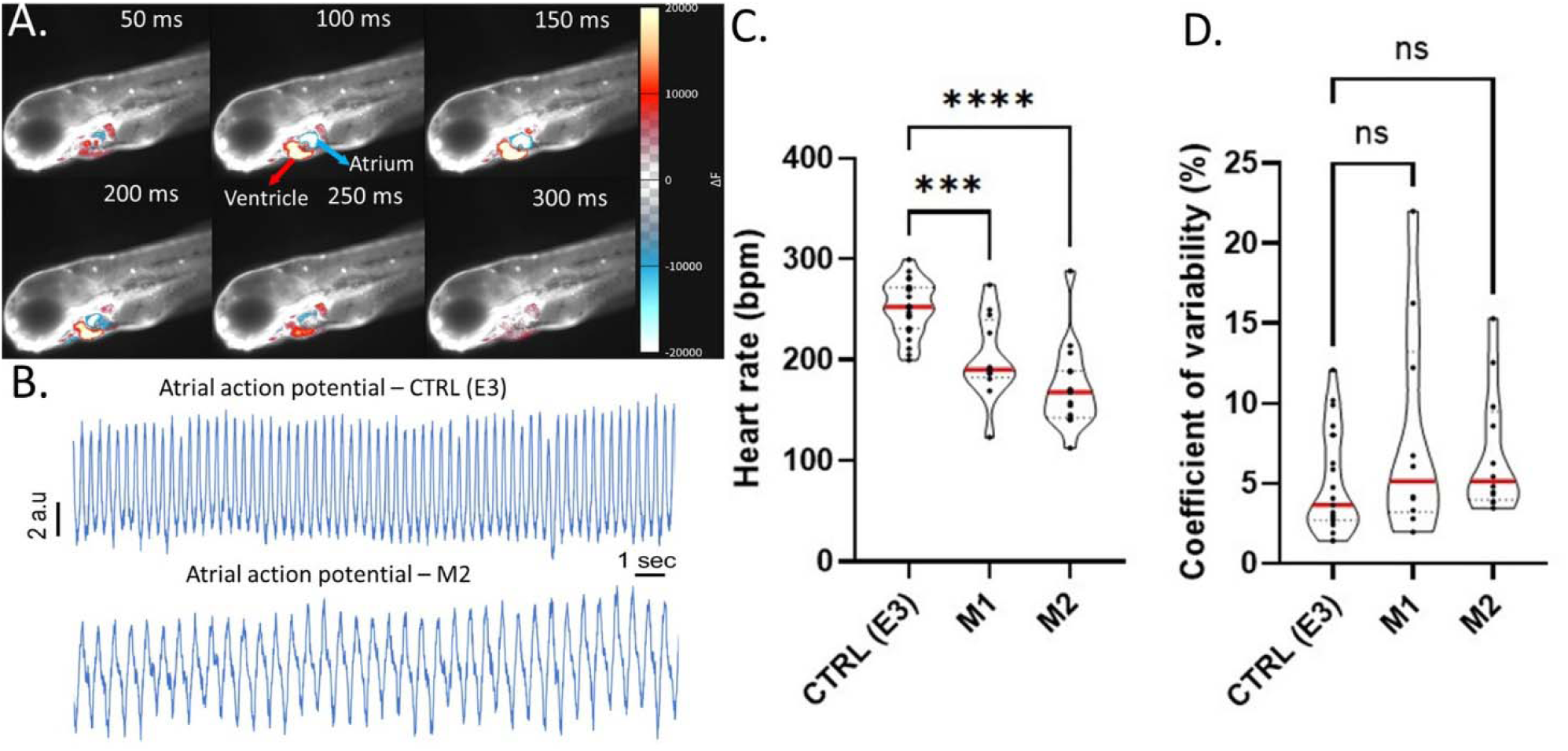
Environmentally relevant exposure to bisphenol mixtures induces bradycardia. (A) Zebrafish larvae in a lateral position, exposing the heart charged with a voltage-sensitive dye (red indicates depolarization) and a time series of the voltage depolarization signal across heart regions. The image at time 50ms corresponds to the beginning of atrial depolarization and the end of ventricular depolarization. (B) Action potentials (APs) recorded in the atrial regions of the zebrafish heart. (C) Quantification of cardiac APs rate (n = 8 for each condition, executed in duplicate, non-parametric test, Kruskal-Wallis, Dunn’s multiple comparisons test, p < 0.05) and (D) coefficient of variability (interval between two APs) in zebrafish exposed to bisphenol mixtures (n = 8 for each condition, executed in duplicate, non-parametric test, Kruskal-Wallis, Dunn’s multiple comparisons test, p < 0.05). As shown in Supplementary Figure S2, controls with matching ethanol content and E3 exhibit no statistically significant changes cardiac function

**Figure 4.**
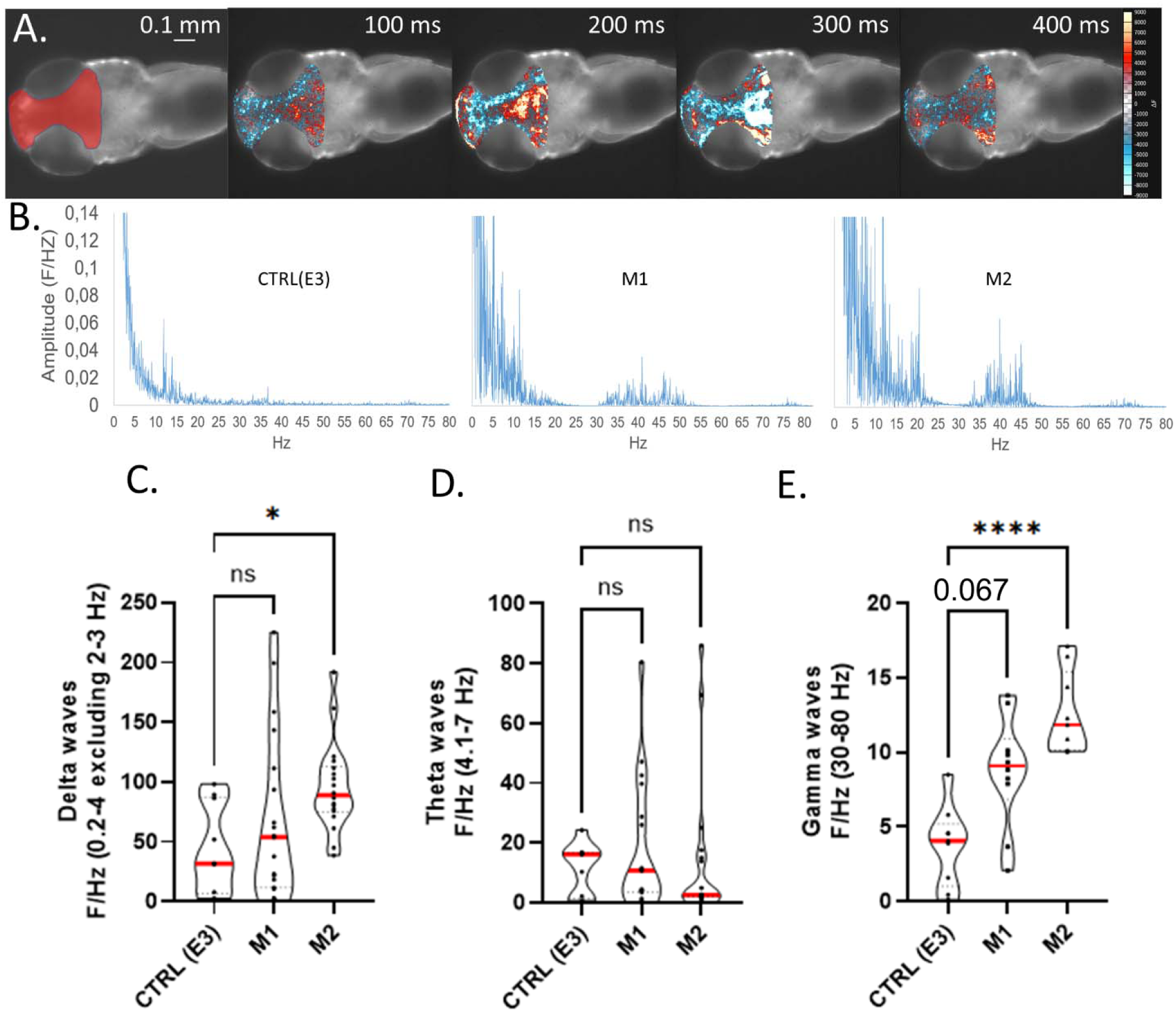
Environmentally relevant exposure to bisphenol mixtures induces neurophysiological alterations, with broader effects if BPA is present. (A) Example of zebrafish larvae in the dorsal position to expose the targeted brain region (indicated in transparent red) and a time (100 ms) series of voltage signal dynamics (red/yellow indicates depolarization) as captured by *in vivo* microscopy. (B) Example of Fourier Transform (FT; F/Hz) of brain voltage activity. Quantifications of brain voltage activities for the frequency intervals (C) 0.2–4 Hz, (D) 4.1–7 Hz and (E) 30–80 Hz in zebrafish exposed to the bisphenol mixtures (n > 7 zebrafish larvae for each condition, executed in duplicates; non-parametric test, Kruskal-Wallis, Dunn’s multiple comparisons test, p < 0.05). As shown in Supplementary Figure S3, controls with matching ethanol content and E3 exhibited no statistically significant changes in brain wave oscillations. The remaining frequency spectrum was also statistically unaffected following exposure to the bisphenol mixtures (Supplementary Figure S4).

### 3.4 Impact of bisphenol mixtures on neurovascular structures imaged in living zebrafish larvae

The brain physiological and motor behavioral alterations induced by bisphenol mixture exposure prompted us to assess cellular-level changes in the developing brain using in vivo two-photon microscopy. We specifically reconstructed the three-dimensional cerebrovascular architecture of *Tg(fli1a:GFP)y1Tg:Tg(mpeg1:mCherry)* zebrafish (Fig. 5), providing a readout of neurovascular unit integrity [36]. Representative Z-stack reconstructions of the entire midbrain vasculature are shown in Fig. 5A1–A3. Three-dimensional (3D) skeleton analysis of the cerebrovascular network (Fig. 5B1–B3) showed that developmental exposure to bisphenol mixtures reduced total vascular length (Fig. 5E), vascular volume (Fig. 5F), the number of branches (Fig. 5G), and the number of terminal segments for both mixtures (Fig. 5H). As a corollary, we examined whether bisphenol mixture exposure could affect glial cells, focusing on microglial cells using Tg(mpeg1:mCherry) zebrafish larvae (Fig. 5C, 5D1-D3). We observed no significant differences in microglial abundance (Fig. 5I). These findings indicate that bisphenol mixture exposure interferes with neurovascular development, with comparable effects observed between M1 and M2.

**Figure 5.**
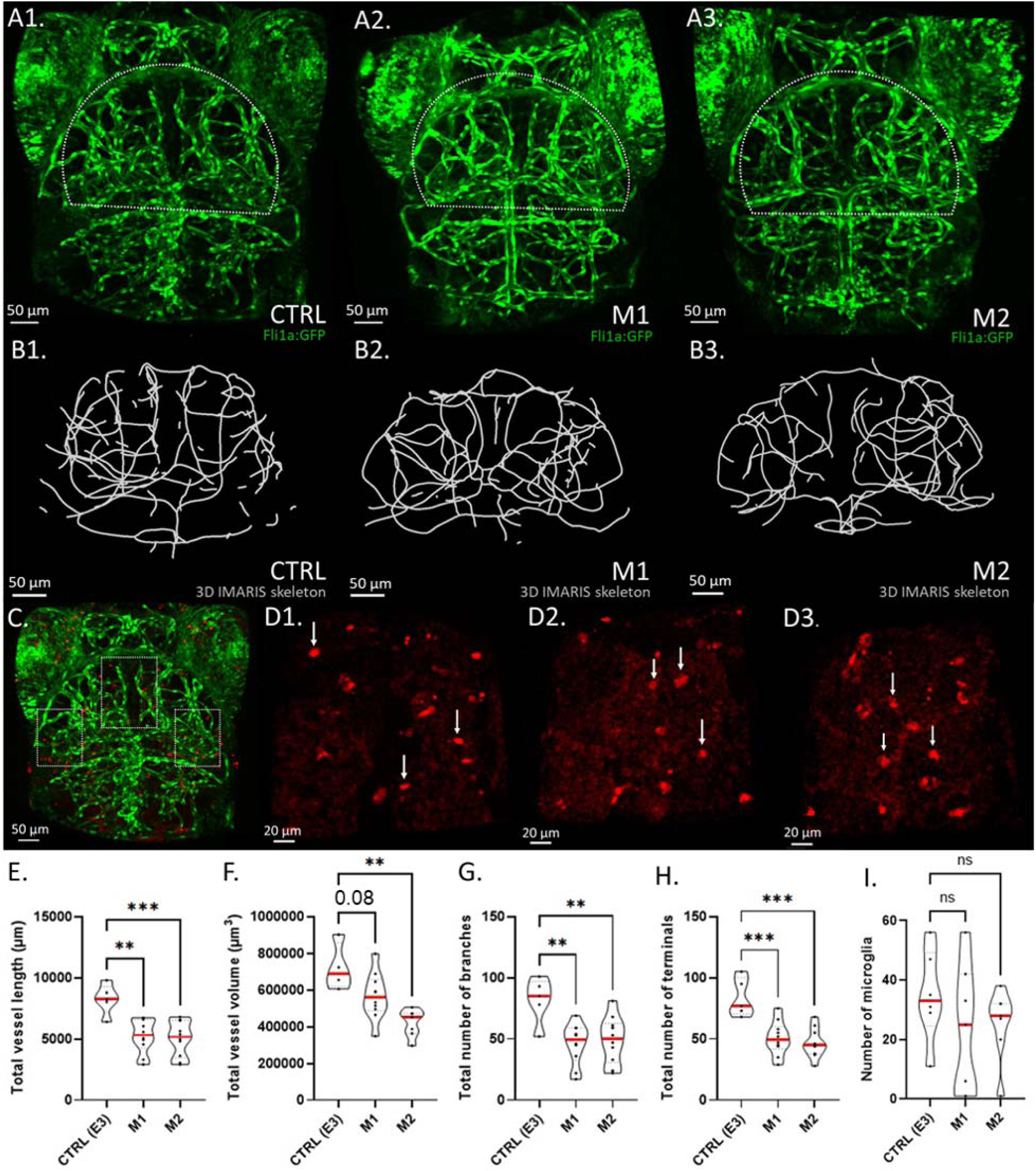
Environmentally relevant exposure to bisphenol mixtures impairs cerebrovascular architecture but does not alter microglia in zebrafish larvae. (A1-A3) Representative two-photon microscopy images of the brain cerebrovasculature in 5 dpf *Tg(fli1a:GFP)y1Tg* zebrafish larvae exposed to control (E3), mixture 1 (M1), or mixture 2 (M2). The dashed outline indicates the region of interest (ROI) used for quantitative analysis. Scale bar = 50 μm. (B1–B3) Representative 3D skeleton reconstructions generated with IMARIS software for quantitative analysis of the cerebrovascular network. (C) two-photon Z-stack reconstructions of a *Tg(fli1a:GFP)y1Tg:Tg(mpeg1:mCherry)* zebrafish larvae showing the brain vasculature in green and microglia in red. Dorsal view with the caudal tail oriented downward. Three equally sized adjacent subregions (white dashed squares) within the predefined region of interest (ROI) are shown. We manually counted microglia in each subregion and summed the counts to obtain the total number of microglia per larva. Scale bar = 50 µm. Arrows indicate representative microglial cells. (D) Examples of microglia in (D1) CTRL(E3), (D2) M1, and D3) M2. Scale bar = 20 µm. (E–I) Quantification of total vessel length (E), total vessel volume (F), total number of branches (G), total number of terminal segments (H), and total number of microglia within the selected ROIs (I). Data are presented as violin plots with median (red line) and individual biological replicates (black dots). Statistical significance was determined using a non-parametric Kruskal-Wallis test with Dunn’s multiple comparisons test (p < 0.05).

### 3.5 Transcriptomic signatures associated with exposure to bisphenol mixtures

Lastly, we performed bulk RNA-seq to assess shared or specific transcriptomic alterations induced by M1 and M2 in zebrafish larvae, which may underpin the modifications reported so far. No individual genes reached the significance threshold for differential expression (FDR < 0.05). We then performed GSEA on ranked gene lists to test whether transcriptional changes occur at the pathway level. We used KEGG and GO Biological Process analyses to characterize enriched molecular pathways and biological processes underlying the transcriptional response. We provide full lists of enriched KEGG and GO pathways identified by GSEA, along with the implicated genes and corresponding NES and FDR values, in the Supplementary Material (Tables S2-S5). Overall, M1 and M2 exhibited highly overlapping transcriptomic signatures at the pathway level, with most enriched KEGG pathways shared between the two mixtures. We report a shared set of neuronal and vascular transcriptomic responses, with 11 positively enriched and 8 negatively enriched KEGG pathways due to M1 or M2 exposure (Fig. 6B1–B2). We found shared negative enrichment of neuronal signaling pathways, including neuroactive ligand–receptor and calcium signaling. Consistent with vascular changes, we found negative enrichment for cell adhesion molecule (CAM), IgSF CAM (Immunoglobulin Superfamily Cell Adhesion Molecules), and cadherin signaling. GO Biological process analysis further revealed shared negative enrichment of neuronal processes involved in axon development, synaptic signaling, ion transport, and neurotransmission (see Tables S2-S5). Next, both M1 and M2 showed positive enrichment of pathways related to protein synthesis and cellular metabolism, including ribosome and aminoacyl-tRNA biosynthesis, primary bile acid and steroid biosynthesis, and oxidative phosphorylation, among others (Fig. 6A1–A2). Overall, the convergence of KEGG and GO analyses indicates common alterations in neuronal and vascular functions following M1 and M2 exposure, consistent with the *in vivo* functional findings.

**Figure 6.**
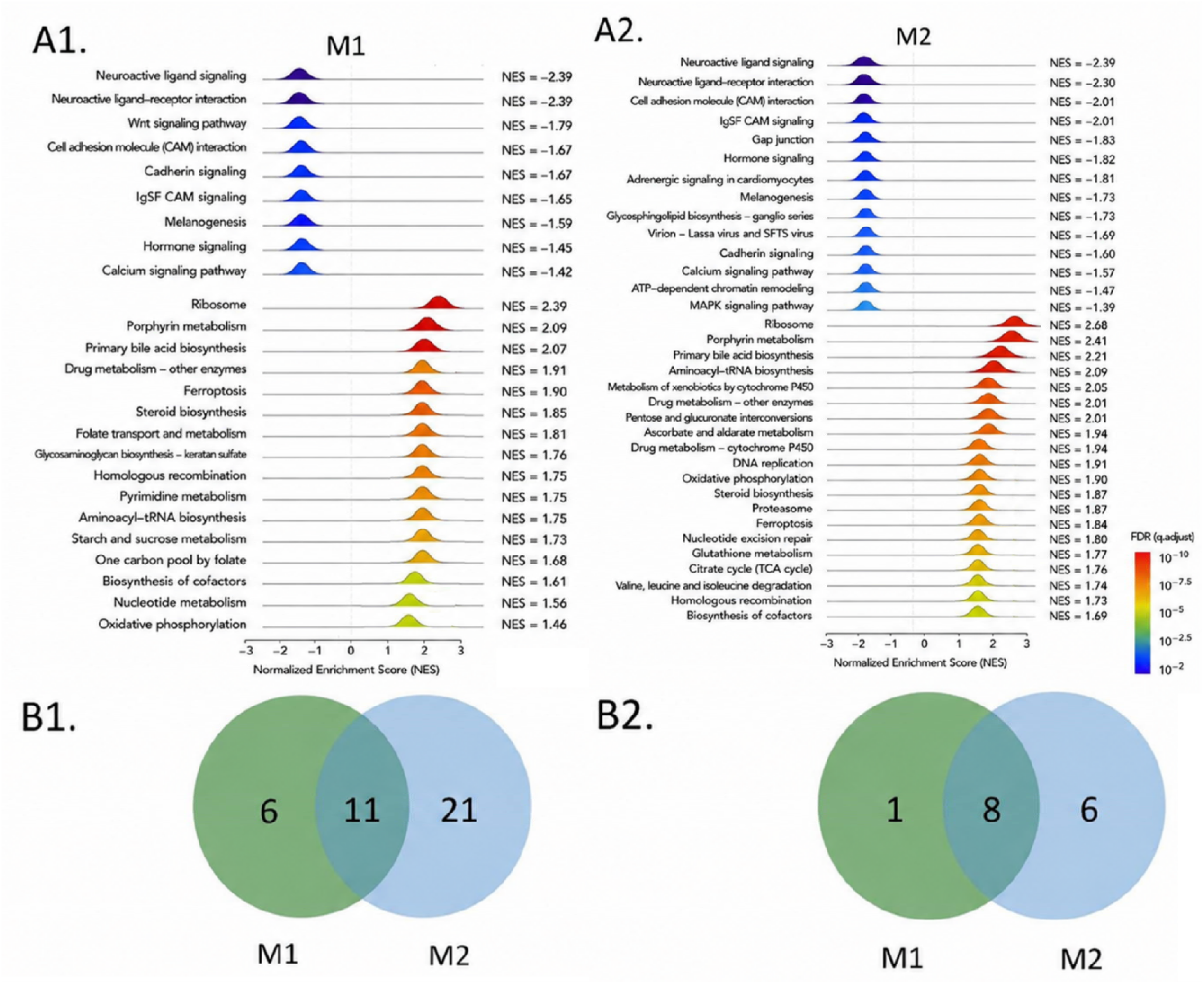
Environmentally relevant exposure to bisphenol mixtures induces shared and mixture-specific transcriptomic signatures. A) Positive or negative enriched biological pathways ranked by normalized enrichment score (NES), with color indicating significance (−log₁₀FDR) for A1) M1 exposure and A2) M2 exposure. Differentially expressed pathways were defined by shared genes (see Tables S2-S5 for complete pathway lists and implicated genes). B) Venn diagrams illustrating the overlap of enriched pathways between bisphenol mixture exposure conditions. B1) Venn diagram illustrating commonly positively enriched pathways shared between M1 and M2. B2) Venn diagram illustrating commonly negatively enriched pathways shared between M1 and M2.

We also identified pathways enriched in M1 or M2 alone, reflecting mixture-specific transcriptomic alterations. Notably, M1 exposure induced only 6 pathways and repressed only 1 pathway specifically (Fig 6B), whereas M2 exposure modulated many more specific pathways (21 induced, 6 repressed) (Fig. 6B). Among these mixture-specific pathways, M1 showed specific negative enrichment of the WNT signaling pathway, which is important for vascular integrity (Fig. 6A1). M2 showed specific negative enrichment of adrenergic signaling in cardiomyocytes and MAPK signaling among other pathway-level alterations (Fig. 6A2, see Tables S2-S5). These findings suggest that including BPA in the mixture is associated with additional pathway-level alterations.

## 4. Discussion

Examining the effects of environmentally relevant pollutant mixtures, such as bisphenols, is fundamental for generating datasets that can inform regulatory organisations. Here, we performed a multi-level investigation using zebrafish larvae, exposing them to environmentally relevant mixtures of BPF, BPAF, and BPS, with or without BPA, to characterize physiological and molecular adaptations. This builds on our examination of individual bisphenols at environmentally relevant concentrations [10]. We report that mixture exposure induced greater-than-predicted toxicity and reduced locomotor activity. These alterations were accompanied by changes in cardiac and neuronal physiological outputs and neurovascular structure. Transcriptomic analysis identified both shared and mixture-specific alterations in pathways related to neuronal communication, vascular development, and metabolism.

### 4.1 Toxicity and behavioral changes induced by bisphenols: mixture effects and BPA presence

Experimentally, we reported toxicity following M1 and M2 exposure (4.16% and 6.54%), contrary to model prediction and to the lack of effects previously shown following exposure to the individual bisphenols under an identical experimental framework [10]. Concentration addition modeling assumes similar modes of action and additive effects for these bisphenols [37], whereas our experimental results indicate that mixture exposure can activate unique biological processes (e.g., transcriptomic changes, see discussion below). Along these lines, synergistic toxicity of bisphenol mixtures have been reported in *in vitro* models, algal models, and zebrafish [37–40]. We did not detect gross morphological malformations or muscle deformities following M1 or M2 exposure, consistent with previous studies in zebrafish [41–43], as developmental defects have been reported at higher concentrations [41, 44].

At the motor-behavioral level, the M1 and M2 bisphenol mixtures provoked significant hypoactivity under light/dark conditions regardless of the presence of BPA (Fig. 2). In comparison, exposure to individual bisphenols at low concentrations induced hypoactivity only during the dark phases [10]. This general behavioral response has been reported following exposure to individual bisphenols [13, 14, 45] and reflects alterations in motor neurons, or broader neurotransmitter systems rather than visual dysfunction [46, 47]. Taken together, these findings indicate that mixture exposure broadens the toxic and behavioral effects of individual bisphenols without inducing morphological defects.

### 4.2 Cardiac and neurovascular adaptations accompany motor-swimming alterations in zebrafish larvae

We found that exposure to bisphenol mixtures caused statistically significant bradycardia, regardless of the presence of BPA in the mixture [9, 12, 48]. In our previous study, we observed bradycardia only under specific exposure conditions of individual bisphenols [10]. Notably, exposure to single BPA and BPS at the concentrations used in the present mixtures (100 ng/L) did not alter heart rate, whereas BPF (10ng/L) and BPAF (100ng/L) induced bradycardia. Together, these results suggest that M1 and M2 exposure retains the cardiac effects generated by single bisphenols in the mixture. At the neuronal level, exposure to M1 was associated with a trend toward increased high-frequency gamma activity, whereas M2 exposure resulted in a statistically significant increase in gamma activity and, in addition, increased delta oscillations. This readout is associated with sleep perturbation and memory consolidation in mammals [49–53], and may serve as a biomarker of neuropathological conditions [54–60]. Furthermore, the reported increased gamma activity could be linked to neuronal network excitability [61–63]. In our previous study, individual bisphenols affected only gamma activity [10]. We also identified reduced cerebrovascular complexity (Fig. 5) [36, 64, 65]. Previous studies have reported that impaired cerebrovascular development is associated with neurological or synaptic alterations which are consistent with the hypoactivity observed herein (Fig. 2) [64–66]. Because cerebrovascular development depends on cardiac blood flow [67, 68], mixture-induced bradycardia (Fig. 3) may also have contributed to the observed outcome. These findings were also consistent with negative enrichment of vascular pathways, as discussed below (Fig. 6 and Tables S2-S5). As a corollary, we observed no changes in microglial number in either M1 or M2 (Fig. 5I), suggesting that neuroinflammation is, at least, not a primary mechanism involved.

### 4.3 Bisphenol mixtures induced shared and specific transcriptomic signatures

Both mixtures provoked negative enrichment of pathways involved in neuronal signaling, including axon development (Tables S2–S5), calcium signaling, and cell adhesion pathways [69, 70], that may contribute to the observed neurophysiological adaptations. In addition, M2 exposure was associated with a negative enrichment of MAPK signaling, implicated in synaptic plasticity [25, 71–74]. This may have contributed to the broader neurophysiological response observed following M2 exposure (Fig. 4). We identified shared negative enrichment of signaling pathways relevant to vascular development and function, including CAM and cadherin signaling, following both M1 and M2 exposure. WNT signaling was negatively enriched after M1 exposure (Fig. 6). These signaling pathways are important regulators of angiogenesis and neurovascular network formation [75–80], and may be linked to the observed neurological and neurovascular changes.

We also observed negative enrichment in calcium signaling in both M1- and M2-exposed larvae, which may contribute to bradycardia. Previous studies have shown that bisphenols can affect key regulators of cardiac Ca²⁺ handling, including an L-type Ca²⁺ channel and SERCA pump, thereby potentially impairing excitation–contraction coupling and cardiac function [12, 81]. In M2-exposed larvae, we also found negative enrichment of the adrenergic signaling pathway in cardiomyocytes, suggesting impaired β-adrenergic cardiac signaling. This pathway regulates cAMP production, Ca²⁺ handling, and excitation–contraction coupling, which are important determinants of heart rate [37, 38, 82–84]. Our data also show a positive enrichment of metabolic pathways in both M1- and M2-exposed larvae, including ribosome biogenesis, oxidative phosphorylation, aminoacyl-tRNA biosynthesis, and xenobiotic metabolism [69, 70, 85–87].

Our data indicate that mixture exposure induces distinct qualitative molecular responses compared to individual bisphenols. The presence of BPA also modified the molecular response, suggesting that combined bisphenol exposure engages biological processes not observed with individual compounds. The mixture exposure did not replicate the gene expression changes we previously observed following exposure to the individual compounds, which could reflect non-additive mixture effects [88]. Previous work in zebrafish showed that mixture-induced transcriptional responses were not simply the sum of those induced by individual chemicals, with mixtures instead producing distinct biological responses [89]. Importantly, bisphenols act through overlapping but distinct receptor targets, and mixtures can diversify pathway activation rather than amplifying a shared signature [90, 91]. Furthermore, mixtures display non-monotonic, dose-dependent effects, as shown for adipogenic gene markers, where a BPA, BPS, and BPF combination produced a U-shaped response absent in individual bisphenols [92]. Comparable work in ovarian granulosa cells reported that bisphenol cocktails did not potentiate single-chemical effects, but rather triggered qualitatively distinct outcomes [93]. Our data indicate that metabolic remodeling represented the main point of convergence between mixture and single bisphenol exposures [69, 70, 85–87]. In summary, these datasets support the notion that bisphenol mixtures elicit qualitatively distinct transcriptomic responses rather than simply amplifying those induced by single compounds.

## 5. Limitations and Conclusions

Our study provides a snapshot of the impact of bisphenol mixtures on zebrafish physiology and transcriptomics. Longer-term studies are necessary to determine whether these alterations are persistent or reversible. We formulated M1 and M2 using the lowest environmentally relevant concentrations of each bisphenol that induced effects in our previous study. Our goal was to combine these bisphenol concentrations to generate environmentally relevant mixtures, with or without BPA, and capture the resulting biological response. However, this design does not allow us to fully characterize the interactions between bisphenols. This would require testing multiple mixture concentrations and generating concentration–response curves. We acknowledge that direct comparison with our previous study [10] should be interpreted with caution, as we identified pathway-level alterations based on significantly regulated individual genes, whereas the present study relied on GSEA. Nevertheless, this comparison provides a biological framework for identifying unique responses associated with mixture exposure. The links between the identified transcriptomic alterations and the observed physiological phenotypes remain correlative and require functional validation.

In conclusion, environmentally relevant bisphenol mixtures induced extra-physiological alterations in zebrafish larvae. We report the overlap between M1 and M2 and show that BPA contributed to specific, additional changes. Comparison with our previous study indicates that mixture exposure produces broader functional and molecular alterations than individual bisphenol exposure. Overall, these findings highlight the importance of assessing mixtures to better capture real-world exposure scenarios and to inform risk assessment and regulation.

## Acknowledgments

This work was fully supported by ANSES Optofish to NM and CJ. We also thank the IGF zebrafish platform ZEFIX-ZebKO. RNA-Seq library was performed at GeT Genopole Toulouse Midi-Pyrénées facility. We thank Zoé Compagnie and Prescillia Alves-Gomes for the sequencing of libraries at INRAE PGTB sequencing facility (doi:10.15454/1.5572396583599417E12). We are grateful to the genotoul bioinformatics platform Toulouse Midi-Pyrenees for providing computing resources for RNAseq bioinformatics and biostatistics analyses (GeT-TRiX). We thank IPAM-BCM Platform, a member of the national infrastructure France-BioImaging supported by the French National Research Agency (ANR-10-INBS-04 and ANR-24-INBS-0005 "Biogen").

