## supplementary material for "Environmentally relevant bisphenol mixtures modify heart–brain physiology, cerebrovascular structures, and transcriptomic profiles in zebrafish larvae"

**Supplementary Tables and Figures**

**Table S1.** **Toxicological and Morphological Screening of zebrafish larvae exposed to E3 or vehicle (EtOH) controls (as compared to Figure 1).** Toxicity, screening hatching rates, and gross morphological parameters of zebrafish larvae exposed to vehicle controls. No statistically significant differences were observed between E3 medium control and the vehicle controls for all tested parameters (One-way ANOVA, Dunnett’s multiple comparisons test, p<0.05). For morphology measurements, data are reported as means ± SD (experiments conducted in duplicate, n=15/group). Cumulative hatching rate consecutively assessed at 72, 96, and 120 hpf are expressed as hatched eggs / total x 100 (experiments conducted in triplicate, n=84/condition; Two-way ANOVA, Dunnett’s multiple comparisons test, p < 0.05). Muscle formation defects were reported as means of somite length (µm, n=15/group, duplicate, Two-way ANOVA, Dunnett’s multiple comparisons test, p < 0.05).

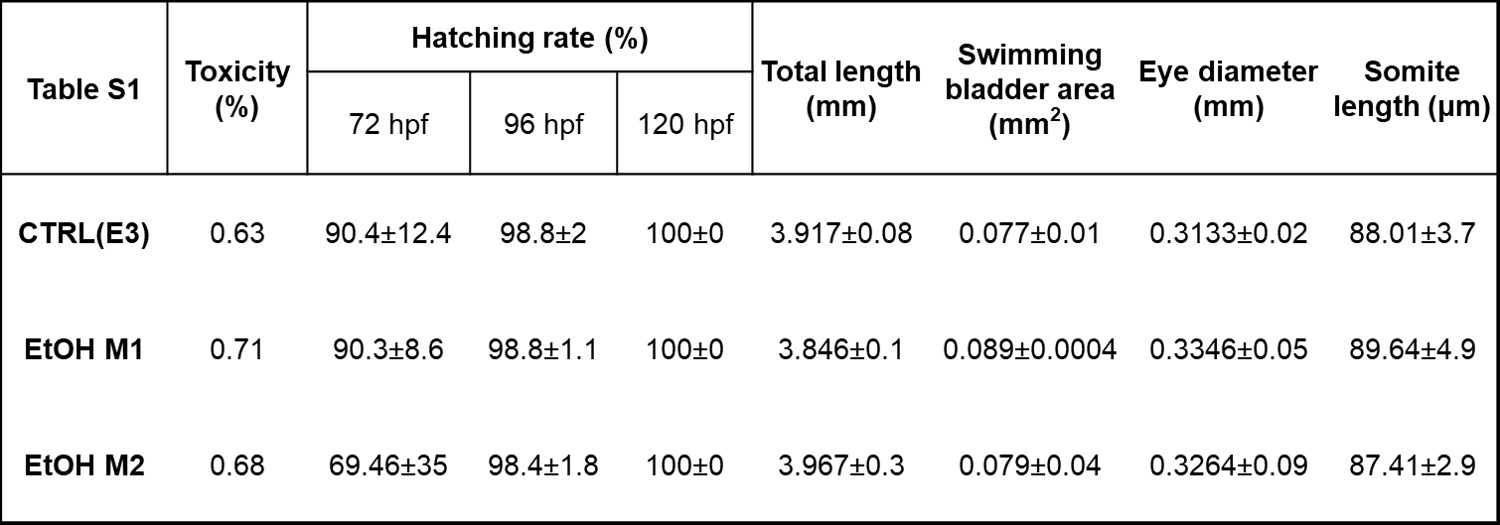

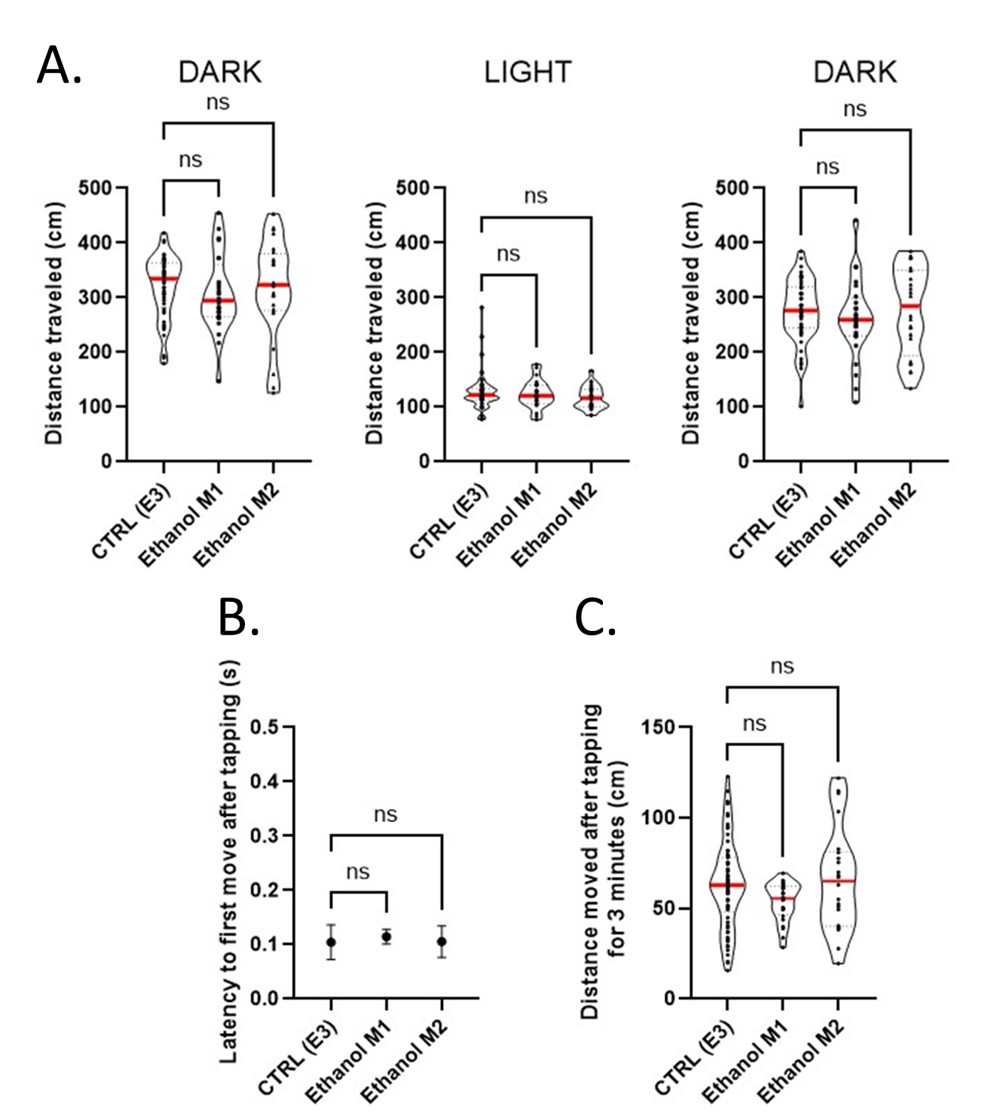

**Figure S1. Motor-behavior patterns of zebrafish larvae exposed to E3 or vehicle (EtOH) controls (as compared to Figure 2).** No significant differences were observed in zebrafish larvae exposed to E3 culture medium or vehicle controls in both behavioral tests. A) Comparison of basal locomotion in light and dark conditions in zebrafish larvae exposed to E3 culture medium vs ethanol vehicle control (N=24 for E3 control and N=24 for ethanol controls, executed in duplicate, Non-parametric test, Kruskal-Wallis test, Dunn’s multiple comparisons test, p<0.05). B-C) Patterns of vibrational startle response of zebrafish larvae exposed to E3 or ethanol vehicle controls (N=24 for E3 control and N=24 for EtOH controls, executed in duplicates, parametric test, One-way Anova test, Dunnett’s multiple comparisons tests): B) Latency to first move after tapping. Data were represented as mean ± SD. C) Distance moved after tapping for 3 minutes.

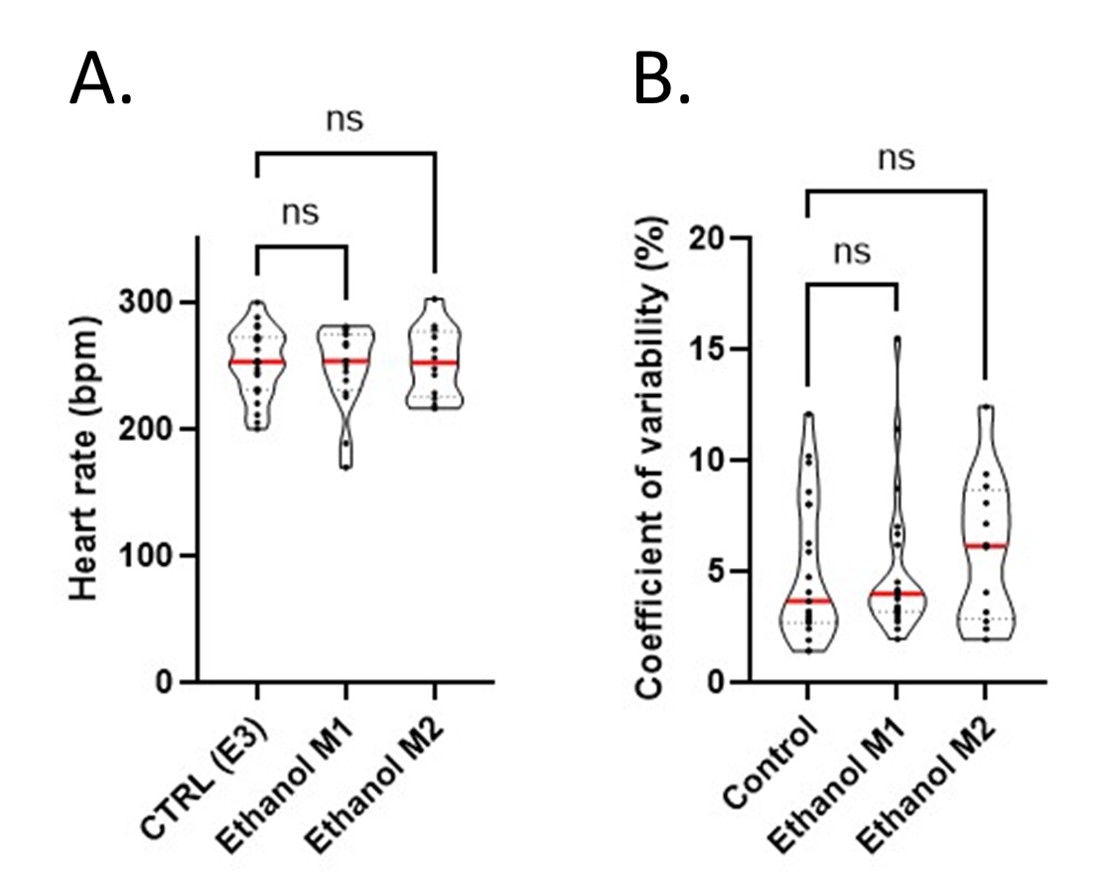

**Figure S2.** **Cardiac physiological modifications** **of zebrafish larvae exposed to E3 or vehicle controls (as compared to Figure 3).** A) Quantification of cardiac APs rate and B) coefficient of variability (interval between two APs) in zebrafish larvae exposed to E3 culture medium or ethanol vehicle (n=10 for each condition, executed in duplicates, non-parametric test, Kruskal-Wallis test, Dunn’s multiple comparisons test, p< 0.05). No statistically significant differences were observed between E3 and ethanol vehicle controls.

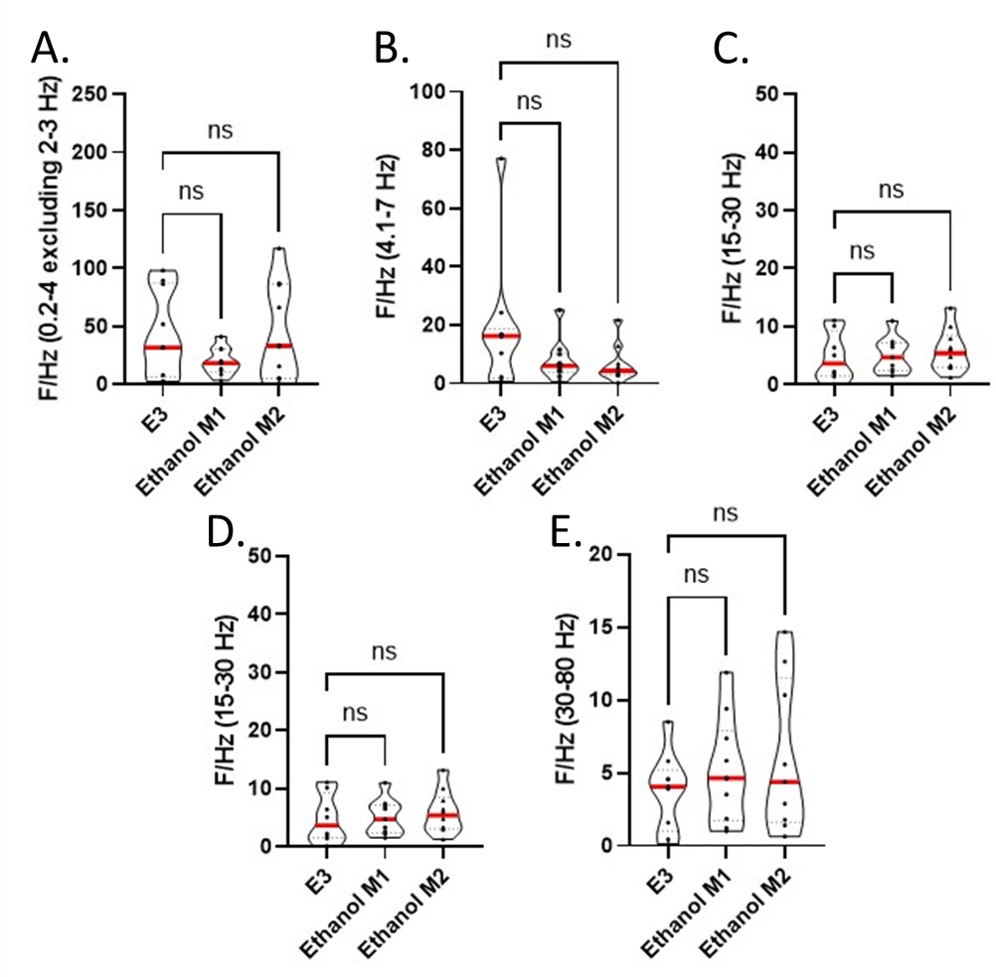

**Figure S3. Neurophysiological modifications of of zebrafish larvae exposed to E3 or vehicle controls (as compared to Fig. 4).** Quantifications of brain voltages activities for the following frequencies intervals: A) 0.2-4 Hz, B) 4.1-7 Hz, C) 8-15 Hz, D) 15-30 Hz and E) 30-80 Hz in zebrafish exposed to E3 culture medium or vehicle ethanol control (n= 10 zebrafish larvae for each condition, executed in duplicates, parametric test, One-way Anova, Dunnett’s multiple comparisons tests, p< 0.05). E3 and ethanol vehicle controls showed no stastically significant differences.

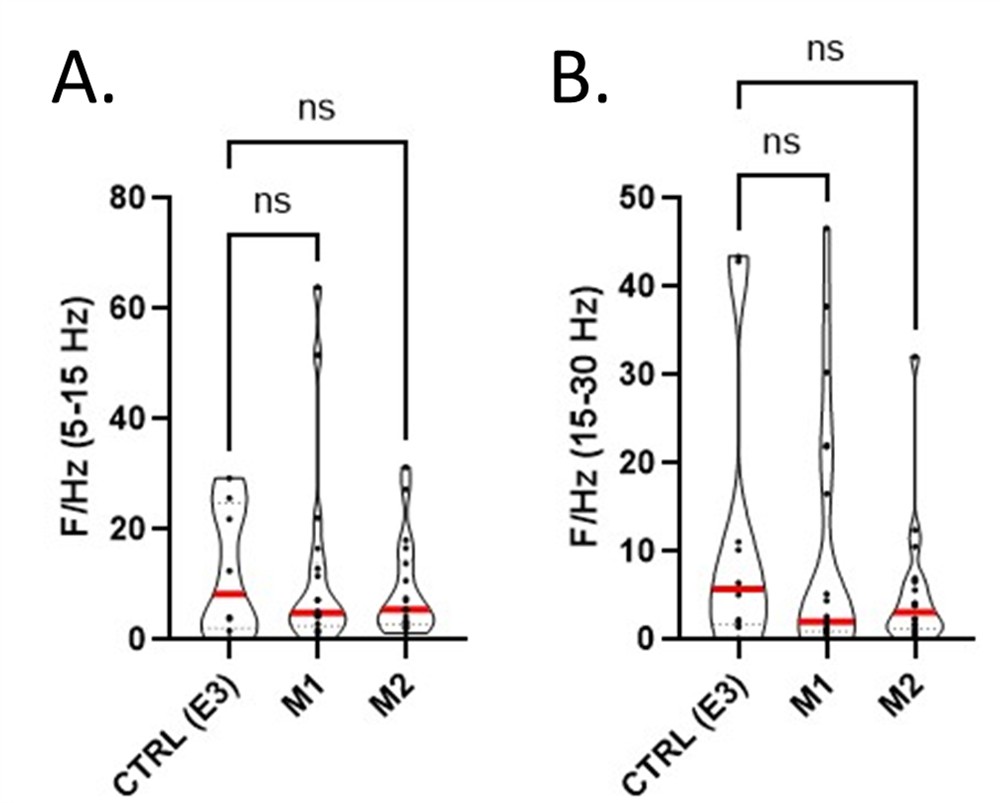

**Figure S4. Neurophysiological modifications exposed to bisphenol M1 and M2 mixtures.** Quantifications of brain voltages activities for the following frequencies intervals: A) 8-15 Hz and B) 15-30 Hz (n=10 zebrafish larvae for each condition, executed in duplicates, non-parametric test, Kruskal-Wallis test, Dunn’s multiple comparisons test, p< 0.05).

**Table S2**. Gene Ontology (GO) pathways enriched following M1 exposure and their associated genes, identified by gene set enrichment analysis (GSEA).

| ID | Description | NES | FDR(p adjusted value) | Genes implicated |
| --- | --- | --- | --- | --- |
| GO:0042254 | ribosome biogenesis | 2.68255980674827 | 1.14653846153846e-08 | imp3/rpl26/mrpl10/mrm3a/nle1/noc2l/rbis/urb2/exosc2/polr1c/rpp40/exosc6/abt1/mphosph6/snu13b/emg1/wdr18/znhit3/pwp2h/utp4/utp23/mrm3b/nop14/heatr1/mrto4/pin4/sbds/lsm6/ddx31/wdr12/nom1/nifk/utp11/tfb1m/ddx27/mrpl22/ngdn/pih1d1/wdr3/ddx51/heatr3/gnl3/mrpl20/rrp36/gar1/nvl/riok2/slx9/bxdc2/pdcd11/trmt112/nop58/dimt1l/bud23/rexo4/nip7/mettl16/mphosph10/wdr36/dkc1/npm1a/nsun5/rpusd2/ppan/mdn1/ddx49/rpl7l1/ddx52/bop1/abce1/nol6/sdad1/tbl3/nop10/ddx10/mrpl36/pak1ip1/rpl35/rbfa/zcchc4/rrp9/rps7/si:dkey-251i10.3/utp15/snu13a/rrp7a/nop56/nhp2/wdr75/ddx18/c1d/rcl1/wdr55/rrp8/tsr2/eif6/nat10/utp25/nop2/nop16/polr1d/mterf3/ebna1bp2/rps21/ftsj3/bysl/utp6/rrs1/gnl3l/rps8a/mpv17l2/rps14/nol10/mtrex/rpl5a/riok1/esf1/rpl35a/pop5/ran/rpf1/mrps11/gnl2/rpl14/rpf2/tfb2m/fbl/utp3/dhx37/rrp1/rps16/ddx54/gtpbp10/rpl7/tsr1/ddx28/znf622/malsu1/rps9/mak16/c1qbp/lsg1/naf1/kri1/rpl23a/urb1/exosc8/fcf1/pes/pop4/rps15/rpsa/mtg2/rrp15/rpl7a/mterf4/exosc7/dcaf13/rps5/rplp0/exosc9/tsr3 |
| GO:0006364 | rRNA processing | 2.64217411952833 | 1.14653846153846e-08 | imp3/mrm3a/exosc2/rpp40/exosc6/abt1/mphosph6/snu13b/emg1/wdr18/znhit3/pwp2h/utp4/utp23/mrm3b/heatr1/mrto4/pin4/lsm6/wdr12/nifk/utp11/tfb1m/ddx27/ngdn/pih1d1/wdr3/ddx51/gnl3/rrp36/gar1/riok2/slx9/pdcd11/trmt112/dimt1l/bud23/rexo4/mettl16/mphosph10/wdr36/dkc1/nsun5/rpusd2/ddx49/rpl7l1/ddx52/bop1/nol6/tbl3/nop10/ddx10/rpl35/rbfa/zcchc4/rrp9/rps7/si:dkey-251i10.3/utp15/snu13a/rrp7a/nhp2/wdr75/ddx18/c1d/rcl1/wdr55/rrp8/tsr2/eif6/nat10/utp25/nop2/ebna1bp2/rps21/ftsj3/bysl/utp6/rrs1/gnl3l/rps8a/rps14/nol10/mtrex/riok1/esf1/pop5/rpf1/mrps11/gnl2/rpf2/tfb2m/fbl/utp3/dhx37/rrp1/rps16/ddx54/rpl7/tsr1/mak16/naf1/kri1/urb1/exosc8/fcf1/pes/pop4 |
| GO:0034470 | ncRNA processing | 2.61312605149024 | 1.14653846153846e-08 | ints6/prkra/imp3/thumpd2/pus3/wdr4/qtrt2/trmu/mrm3a/mtfmt/trmt10c/exosc2/rpp21/rnf113a/rpp40/exosc6/abt1/mphosph6/snu13b/rpp30/emg1/henmt1/wdr18/tprkb/piwil2/znhit3/elp3/zgc:136564/pwp2h/elp5/utp4/trmt61a/spout1/zc3h7bb/tp53rk/utp23/mrm3b/trmt44/heatr1/mrto4/pin4/trmt6/exosc5/tyw3/rpusd4/adat1/tsen2/trmt12/lsm6/wdr12/nsun2/nifk/utp11/thumpd3/tfb1m/osgep/ddx27/urm1/ctu2/mettl3/ngdn/pih1d1/wdr3/srrt/ddx51/gnl3/rrp36/gar1/riok2/ints6l/slx9/pdcd11/trmt112/dimt1l/bud23/osgepl1/rexo4/ptcd1/mettl16/mphosph10/wdr36/dkc1/mettl2a/thada/pus1/nsun5/gtpbp3/rpusd2/ddx49/rpl7l1/ddx52/bop1/trmt10b/tdrd9/nol6/clp1/ctu1/tsen15/rtcb/tbl3/ints9/nop10/ddx10/rpl35/rbfa/zcchc4/rrp9/rpph1/rps7/si:dkey-251i10.3/utp15/snu13a/rrp7a/elp6/nhp2/wdr75/ddx18/c1d/rcl1/wdr55/rrp8/tsr2/trmt61b/elac1/trpt1/eif6/nat10/utp25/nop2/ebna1bp2/tut1/rps21/elp2/ftsj3/bysl/mettl1/pus7/utp6/rrs1/dtwd1/gnl3l/rps8a/rps14/nol10/rpp25l/thg1l/mtrex/riok1/sepsecs/esf1/eif1axb/pop5/cdk5rap1/zbtb8os/rpf1/mrps11/gnl2/rpf2/tfb2m/trmt2b/fbl/utp3/rpp38/dhx37/rtraf/rrp1/wdr6/ints11/rps16/ddx54/aars1/rpl7/tsr1/mak16/trmt5/trmt9b/naf1/kri1/urb1/snip1/exosc8/fcf1/pes/pop4 |
| GO:0016072 | rRNA metabolic process | 2.5944143069453 | 1.14653846153846e-08 | imp3/mrm3a/exosc2/rpp40/exosc6/abt1/mphosph6/snu13b/emg1/wdr18/znhit3/pwp2h/utp4/utp23/mrm3b/heatr1/mrto4/pin4/exosc5/lsm6/wdr12/nifk/utp11/tfb1m/ddx27/ngdn/pih1d1/wdr3/ddx51/gnl3/rrp36/gar1/riok2/slx9/pdcd11/trmt112/dimt1l/bud23/rexo4/mettl16/mphosph10/wdr36/dkc1/polr1e/nsun5/rpusd2/ddx49/rpl7l1/ddx52/bop1/nol6/tbl3/nop10/ddx10/rpl35/rbfa/zcchc4/rrp9/rps7/si:dkey-251i10.3/utp15/snu13a/rrp7a/nhp2/wdr75/ddx18/c1d/rcl1/wdr55/rrp8/tsr2/eif6/nat10/utp25/nop2/taf1b/ebna1bp2/rps21/ftsj3/bysl/dis3/utp6/rrs1/gnl3l/rps8a/rps14/nol10/mtrex/riok1/esf1/pop5/rpf1/mrps11/gnl2/rpf2/tfb2m/fbl/utp3/polr1a/dhx37/rrp1/rps16/ddx54/pelo/rpl7/tsr1/mak16/naf1/kri1/urb1/exosc8/fcf1/pes/pop4 |
| GO:0034660 | ncRNA metabolic process | 2.54552940229038 | 1.14653846153846e-08 | ints6/prkra/fars2/imp3/thumpd2/pus3/farsa/wdr4/qtrt2/trmu/mrm3a/dars2/wars2/mtfmt/trmt10c/exosc2/rpp21/rnf113a/rpp40/exosc6/dars1/abt1/mphosph6/snu13b/rpp30/emg1/henmt1/wdr18/tprkb/piwil2/znhit3/elp3/zgc:136564/pwp2h/elp5/utp4/trmt61a/spout1/zc3h7bb/tp53rk/utp23/mrm3b/trmt44/heatr1/mrto4/pin4/trmt6/exosc5/tyw3/lars1b/rpusd4/adat1/tsen2/trmt12/lsm6/wdr12/nsun2/nifk/utp11/thumpd3/si:ch211-256m1.8/tfb1m/mars2/osgep/ddx27/urm1/ctu2/mettl3/ngdn/pih1d1/wdr3/srrt/ddx51/snapc3/gnl3/rrp36/gar1/riok2/ints6l/slx9/pdcd11/trmt112/qrsl1/dimt1l/bud23/osgepl1/rexo4/ptcd1/mettl16/gatb/mphosph10/wdr36/dkc1/mettl2a/thada/polr1e/pus1/nsun5/gtpbp3/rpusd2/iars1/ddx49/hars/rpl7l1/ddx52/bop1/trmt10b/tdrd9/qars1/nol6/nars2/vars1/clp1/ctu1/tsen15/rtcb/tbl3/ints9/nop10/ddx10/rpl35/rbfa/zcchc4/rrp9/rpph1/rps7/si:dkey-251i10.3/utp15/snu13a/rrp7a/elp6/nhp2/wdr75/ddx18/yars1/c1d/rcl1/wdr55/rrp8/tsr2/trmt61b/elac1/rpap2/trpt1/rars1/eif6/nat10/utp25/nop2/taf1b/ebna1bp2/tut1/rps21/elp2/tars1/ftsj3/bysl/gars1/mettl1/pars2/dis3/pus7/iars2/utp6/rrs1/dtwd1/gnl3l/rps8a/rps14/nol10/rpp25l/thg1l/mtrex/riok1/sars1/sepsecs/esf1/eif1axb/pop5/vars2/mepceb/cdk5rap1/zbtb8os/rpf1/mrps11/gnl2/sars2/rpf2/tfb2m/trmt2b/fbl/utp3/polr1a/rpp38/dhx37/rtraf/rrp1/wdr6/ints11/rps16/ddx54/aars1/pelo/rpl7/tsr1/mak16/trmt5/trmt9b/naf1/kri1/dtd2/urb1/snip1/exosc8/fcf1/pes/pop4 |
| GO:0022613 | ribonucleoprotein complex biogenesis | 2.40988428678125 | 1.14653846153846e-08 | imp3/rpl26/mrpl10/mrm3a/nle1/noc2l/rbis/gemin5/urb2/exosc2/polr1c/rpp40/exosc6/abt1/mphosph6/snu13b/emg1/wdr18/znhit3/pwp2h/trnau1apa/utp4/utp23/mrm3b/nop14/heatr1/mrto4/pin4/sbds/snrpd2/lsm6/ddx31/wdr12/nom1/nifk/utp11/tfb1m/prmt7/ddx27/mrpl22/ngdn/pih1d1/gemin2/wdr3/ddx51/heatr3/gnl3/mrpl20/rrp36/gar1/nvl/snrpg/riok2/slx9/bxdc2/pdcd11/trmt112/nop58/dimt1l/bud23/prpf18/mcts1/rexo4/nip7/mettl16/mphosph10/wdr36/dkc1/npm1a/nsun5/rpusd2/ppan/mdn1/ddx49/rpl7l1/ddx52/bop1/abce1/nol6/clp1/sdad1/tbl3/nop10/ddx10/mrpl36/pak1ip1/rpl35/rbfa/zcchc4/rrp9/ptges3b/rps7/si:dkey-251i10.3/utp15/snu13a/rrp7a/nop56/eif3m/nhp2/wdr75/ddx18/c1d/nopchap1/snrpd3l/eif3ea/rcl1/wdr55/rrp8/tsr2/coil/eif6/snrpf/nat10/utp25/nop2/nop16/usp39/polr1d/mterf3/ebna1bp2/rps21/eif3jb/ftsj3/snrpe/bysl/utp6/rrs1/gnl3l/rps8a/mpv17l2/rpl38/rps14/nol10/shq1/mtrex/rpl5a/riok1/esf1/rpl35a/pop5/eif5/ran/eif3ha/rpf1/trnau1apb/mrps11/gnl2/scaf11/rpl14/rpf2/tfb2m/fbl/utp3/dhx37/rrp1/eif3s10/cdc73/rps16/ddx54/denr/gtpbp10/rpl7/tsr1/ddx28/znf622/malsu1/rps9/mak16/c1qbp/lsg1/naf1/kri1/rpl23a/urb1/exosc8/eif3s6ip/fcf1/pes/pop4/rps15/rpsa/eif2s2/mtg2/polr2d/rrp15/rpl7a/srpk1a/mterf4/exosc7/dcaf13/rps5/gemin8/rplp0/exosc9/tsr3/snrpd1/rps27.1/aar2/rpl10/rpl5b/eif3g/gtpbp4/eif3d/rpl6/yju2/noc4l |
| GO:0140053 | mitochondrial gene expression | 2.16306020123481 | 0.000153043817326382 | fastkd3/tefm/trmu/fastkd2/tbrg4/wars2/trmt10c/mrpl58/fastkd1/coa3a/mtg1/mrps18b/tfb1m/uqcc2/chchd1/tsfm/mtrf1l/qrsl1/gatb/gtpbp3/mtif2/hars/mrps24/rcc1l/tfam/trmt61b/mrpl47/mterf3/gars1/iars2/mrpl23/sars2/tfb2m/ptcd3/fastk/mrpl16 |
| GO:0043043 | peptide biosynthetic process | 2.02998011519709 | 1.14653846153846e-08 | dazl/fars2/farsa/zc3h12ab/dph7/rpl26/mrpl10/dars2/wars2/mtfmt/trmt10c/gemin5/exosc2/mrpl58/mrps5/dars1/rpp30/zar1/mrpl9/dcp1b/piwil2/coa3a/trnau1apa/mrpl35/mif4gdb/mrpl30/eif2ak1/dcps/mrpl17/dph5/lars1b/mtg1/mrps17/mrps18b/mrpl4/si:ch211-256m1.8/zgc:171480/mars2/uqcc2/chchd1/tsfm/gss/eif4ea/mrpl22/mettl3/ngdn/mtrf1l/faub/mrps18c/drg2/lsm1/mrpl27/mrpl20/dph6/magoh/eif1ad/eif2a/eefsec/qrsl1/mrpl51/mrps23/mcts1/ago3b/mettl16/gatb/mrpl18/mrps21/gtpbp3/mtif2/iars1/mrpl14/rpl12/hars/mrpl24/pdf/eif2b1/eif4e2rs1/abce1/eif2s1a/qars1/nars2/mrpl41/impact/vars1/mrpl34/fxr2/larp6a/mrps24/mrpl36/rpl35/cnot8/mrps12/mrpl13/rcc1l/eif5a2/rpl32/mrps2/rps7/mrpl12/brf1a/eif2b2/eif3m/eif4ebp3l/yars1/bdh2/eef1b2/eif3ea/mrpl11/rpl22l1/mrpl33/rpl34/rars1/eif6/dph2/ogfod1/gspt1/pde12/mrpl47/rps21/rps17/eef1a1l3/eif3jb/tars1/cnot6l/eef1da/eif4ebp1/gars1/mrps10/pars2/iars2/rc3h2/mrpl23/rpl29/zgc:171772/rpl37/rps8a/gatc/rpl38/rps3a/rps14/rps15a/srp9/larp4b/mrps30/rps20/rps29/tcea3/rpl5a/rpl15/sars1/sepsecs/mrpl3/eif1axb/rpl35a/vars2/eif5/rps18/eif3ha/zc3h15/mrpl32/mrpl37/trnau1apb/mrps11/rpl11/rpl14/rpl39/sars2/uba52/rpl36a/rps26/eif1b/ssb/igf2bp3/rpl24/eif4eb/ptcd3/rpl27/rpl23/eif3s10/rpl31/rps24/rps16/rwdd1/denr/aars1/rpl18/rpl13a/rpl22/mrps16/rbm24a/rplp2/mrpl16/mettl17/mtrfr/drg1/malsu1/rps9/rbm8a/rpl28/rack1/rplp1/rpl23a/eif3f/gspt1l/mrpl2/exosc8/larp1b/eif3s6ip/rps15/rpsa/eef1g/eif2s2/cnbpa/ftsj1/rps11/eif4e3/rplp2l/zgc:114188/polr2d/rpl17/cnot11/faua/pabpc1a/exosc7/rpl19/rps5/eif2s1b/rplp0/rpl18a/exosc9/tufm/rpl21/kars1/rpl4/eef2b/jmjd4/cnbpb/rps13/rpl8/eif5b/rps27.1/rpl10/rpl5b/eif3g/eif3d/mrps35/rpl6/gfm1/eif4e1c/rps3/eif1axa/cnot7/eif2b4/rps12/eef1a1l1/znf598/rpl9/cnot10/eef2a.1/rpl10a/cars1/rps6/rps27a/mars1/nck2a/rps27.2/rps28/mrps7/eif4bb/rpl36/rps23/eif2s3/gtpbp1l/nck2b/eif4g1a/cnot9/dcp2/rps19/dph1/ybx1/dhx29/eif3ja/mrpl15/eif4g3b/samd4a |
| GO:0006412 | translation | 2.02950881785419 | 1.14653846153846e-08 | dazl/fars2/farsa/zc3h12ab/dph7/rpl26/mrpl10/dars2/wars2/mtfmt/trmt10c/gemin5/exosc2/mrpl58/mrps5/dars1/rpp30/zar1/mrpl9/dcp1b/piwil2/coa3a/trnau1apa/mrpl35/mif4gdb/mrpl30/eif2ak1/dcps/mrpl17/dph5/lars1b/mtg1/mrps17/mrps18b/mrpl4/si:ch211-256m1.8/zgc:171480/mars2/uqcc2/chchd1/tsfm/eif4ea/mrpl22/mettl3/ngdn/mtrf1l/faub/mrps18c/drg2/lsm1/mrpl27/mrpl20/dph6/magoh/eif1ad/eif2a/eefsec/qrsl1/mrpl51/mrps23/mcts1/ago3b/mettl16/gatb/mrpl18/mrps21/gtpbp3/mtif2/iars1/mrpl14/rpl12/hars/mrpl24/pdf/eif2b1/eif4e2rs1/abce1/eif2s1a/qars1/nars2/mrpl41/impact/vars1/mrpl34/fxr2/larp6a/mrps24/mrpl36/rpl35/cnot8/mrps12/mrpl13/rcc1l/eif5a2/rpl32/mrps2/rps7/mrpl12/brf1a/eif2b2/eif3m/eif4ebp3l/yars1/eef1b2/eif3ea/mrpl11/rpl22l1/mrpl33/rpl34/rars1/eif6/dph2/ogfod1/gspt1/pde12/mrpl47/rps21/rps17/eef1a1l3/eif3jb/tars1/cnot6l/eef1da/eif4ebp1/gars1/mrps10/pars2/iars2/rc3h2/mrpl23/rpl29/zgc:171772/rpl37/rps8a/gatc/rpl38/rps3a/rps14/rps15a/srp9/larp4b/mrps30/rps20/rps29/tcea3/rpl5a/rpl15/sars1/sepsecs/mrpl3/eif1axb/rpl35a/vars2/eif5/rps18/eif3ha/zc3h15/mrpl32/mrpl37/trnau1apb/mrps11/rpl11/rpl14/rpl39/sars2/uba52/rpl36a/rps26/eif1b/ssb/igf2bp3/rpl24/eif4eb/ptcd3/rpl27/rpl23/eif3s10/rpl31/rps24/rps16/rwdd1/denr/aars1/rpl18/rpl13a/rpl22/mrps16/rbm24a/rplp2/mrpl16/mettl17/mtrfr/drg1/malsu1/rps9/rbm8a/rpl28/rack1/rplp1/rpl23a/eif3f/gspt1l/mrpl2/exosc8/larp1b/eif3s6ip/rps15/rpsa/eef1g/eif2s2/cnbpa/ftsj1/rps11/eif4e3/rplp2l/zgc:114188/polr2d/rpl17/cnot11/faua/pabpc1a/exosc7/rpl19/rps5/eif2s1b/rplp0/rpl18a/exosc9/tufm/rpl21/kars1/rpl4/eef2b/jmjd4/cnbpb/rps13/rpl8/eif5b/rps27.1/rpl10/rpl5b/eif3g/eif3d/mrps35/rpl6/gfm1/eif4e1c/rps3/eif1axa/cnot7/eif2b4/rps12/eef1a1l1/znf598/rpl9/cnot10/eef2a.1/rpl10a/cars1/rps6/rps27a/mars1/nck2a/rps27.2/rps28/mrps7/eif4bb/rpl36/rps23/eif2s3/gtpbp1l/nck2b/eif4g1a/cnot9/dcp2/rps19/dph1/ybx1/dhx29/eif3ja/mrpl15/eif4g3b/samd4a |
| GO:0045333 | cellular respiration | 1.99293254414966 | 4.41989406426173e-05 | ccn6/tefm/sfxn4/uqcc3/si:dkey-85n7.8/ugl/sdhda/ndufs8b/atpsckmt/mdh1b/mdh1ab/dhtkd1/wfs1b/uqcc2/idh3b/sdhaf2/atp5mf/atp5po/sdhdb/ndufb6/lyrm7/cox7b/uqcr10/ndufb8/mrpl24/etfdh/ndufa10/cox6c/cox4i1/ndufv1/ndufaf1/suclg2/si:dkey-190g11.3/antkmt/cox7c/ndufa7/atp5f1e/ndufa8/ndufc2/cox7a2a/sdhc/ogdhb/slc25a55b/slc25a12/coa6/sdha/ndufs6/pm20d1.1/cox6a1/uqcrq/lyrm5a/ndufs2/ndufb9/noa1/sucla2/idh1/uqcrfs1/ndufs8a/cox6a2/cox5ab/lyrm5b/uqcrh/ndufa5/ndufv2/idh3a/mdh2/idh2/cox7a2l/sdhb/dlst/uqcrb/ndufs7/cox5aa/cyc1/ND3/fh/cox5b2/ndufv3/ogdha/cox8a |
| GO:0006518 | peptide metabolic process | 1.98118588803671 | 1.14653846153846e-08 | dazl/fars2/farsa/zc3h12ab/dph7/parla/rpl26/mrpl10/dars2/wars2/mtfmt/trmt10c/gemin5/exosc2/mrpl58/mrps5/dars1/rpp30/zar1/erap2/mrpl9/dcp1b/piwil2/ggt1b/coa3a/trnau1apa/mrpl35/mif4gdb/mrpl30/eif2ak1/dcps/mrpl17/dph5/lars1b/mtg1/mrps17/mrps18b/cpz/mrpl4/si:ch211-256m1.8/zgc:171480/mars2/uqcc2/chchd1/tsfm/gss/eif4ea/mrpl22/mettl3/ngdn/mtrf1l/faub/mrps18c/drg2/lsm1/mrpl27/mrpl20/dph6/magoh/eif1ad/eif2a/eefsec/qrsl1/mrpl51/mrps23/mcts1/ago3b/mettl16/gatb/mrpl18/immp2l/mrps21/gtpbp3/mtif2/iars1/mrpl14/rpl12/hars/mrpl24/pdf/enpep/eif2b1/eif4e2rs1/abce1/eif2s1a/qars1/nars2/mrpl41/pcsk5b/impact/vars1/mrpl34/sec11a/nln/fxr2/larp6a/mrps24/mrpl36/rpl35/cnot8/mrps12/mrpl13/rcc1l/eif5a2/rpl32/mrps2/rps7/mrpl12/brf1a/eif2b2/eif3m/eif4ebp3l/yars1/bdh2/eef1b2/eif3ea/mrpl11/rpl22l1/mrpl33/rpl34/rars1/eif6/dph2/ogfod1/gspt1/pde12/sppl3/mrpl47/rps21/rps17/eef1a1l3/eif3jb/tars1/cnot6l/eef1da/eif4ebp1/gars1/mrps10/pars2/iars2/rc3h2/mrpl23/rpl29/zgc:171772/rpl37/rps8a/gatc/rpl38/spcs3/rps3a/rps14/rps15a/srp9/larp4b/mrps30/rps20/rps29/tcea3/rpl5a/rpl15/sars1/sepsecs/mrpl3/eif1axb/rpl35a/vars2/eif5/rps18/eif3ha/zc3h15/mrpl32/mrpl37/trnau1apb/mrps11/rpl11/rpl14/rpl39/sars2/uba52/rpl36a/rps26/eif1b/thop1/ssb/igf2bp3/rpl24/eif4eb/ptcd3/rpl27/rpl23/eif3s10/rpl31/rps24/rps16/rwdd1/denr/aars1/rpl18/rpl13a/rpl22/mrps16/mipepb/rbm24a/spcs1/rplp2/mrpl16/mettl17/mtrfr/drg1/malsu1/rps9/rbm8a/rpl28/rack1/rplp1/rpl23a/eif3f/gspt1l/mrpl2/exosc8/larp1b/eif3s6ip/rps15/rpsa/eef1g/eif2s2/cnbpa/ftsj1/rps11/eif4e3/rplp2l/zgc:114188/polr2d/rpl17/cnot11/faua/pabpc1a/exosc7/rpl19/rps5/eif2s1b/rplp0/rpl18a/exosc9/tufm/rpl21/kars1/rpl4/eef2b/jmjd4/cnbpb/rps13/rpl8/eif5b/rps27.1/rpl10/rpl5b/eif3g/eif3d/mrps35/rpl6/gfm1/ide/eif4e1c/rps3/eif1axa/cnot7/eif2b4/rps12/eef1a1l1/znf598/rpl9/cnot10/eef2a.1/rpl10a/spcs2/cars1/rps6/rps27a/mars1/nck2a/rps27.2/rps28/mrps7/eif4bb/rpl36/rps23/eif2s3/gtpbp1l/nck2b/lnpep/eif4g1a/cnot9/dcp2/rps19/dph1/ybx1/dhx29/eif3ja/mrpl15/eif4g3b/samd4a |
| GO:0007005 | mitochondrion organization | 1.97295174722294 | 2.83369625164005e-07 | si:dkey-32e23.4/bnip4/adck1/sqstm1/mtm1/mtx1b/mtx1a/rrm2b/sco2/si:dkey-82o10.4/vps13d/sdhaf1/uqcc3/sdhaf3/chchd4b/atp23/phb2b/timm21/ppargc1a/timm29/maip1/coa3a/sdhaf4/aifm1/ndufs8b/chchd4a/alkbh7/chchd6b/cox17/chchd7/coa4/mgme1/cox20/slc25a42/bcs1l/timm13/uqcc2/surf1/sdhaf2/timm50/spata18/lonp1/tmem126a/dnajc19/grpel1/bida/chchd3b/atp13a2/immp2l/wdr45b/lyrm7/uqcc6/cox19/etfa/cluha/dna2/coa5/snx7/apooa/hspd1/atpaf2/phb/ndufaf1/atg5/snx30/polg2/uqcc1/higd1a/chchd10/cox7a2a/primpol/atpaf1/higd2a/si:ch211-166a6.5/pet100/tomm40/apool/fam162a/agtpbp1/pmaip1/romo1/ndufa11/oxa1l/twnk/samm50/rpl37/ndufs8a/rhot1a/slc25a4/nubpl/ndufaf8/ttc19/dnajc11a/mfn1b/miga1/tmem70/dnlz/dnaja3a/mipepb/foxred1/cox7a2l/dlst/atg7/tafazzin/tomm40l/fis1 |
| GO:0002088 | lens development in camera-type eye | 1.96162227904054 | 0.000326438123176076 | fgf19/crygm2a/crybb2/crybb1l3/crygm5/crygm2d6/crygm2b/mipb/cryba1l2/crygm2d9/crygm2d15/dnajb1b/crygm2d12/crygm2d18/cryba2a/crygm2d17/crygm2d7/crygm2d1/lgsn/crygm2d5/crygm2f/cryba1l1/crybb1l2/dlx3b/crygm2d3/crygm2d20/crygm2d11/crygm2d10/crygm2d2/crygm2d21/cryba1a/dlx4b/mipa/lama1/crygm2d13/crygmxl2/crygn2/crygm2d16/cryba4/crygm2d8/tdrd7b/crygm2d14/cryba1b/crygn1/psmc3/foxe3/cryba2b/hsf4/unc45b/crygmx/crygmxl1/si:dkey-57a22.15/sox11b/crybb1l1/psmd6 |
| GO:0009145 | purine nucleoside triphosphate biosynthetic process | 1.93013788013767 | 0.00988355843878473 | nme9/uqcc3/atp5mc3a/atpsckmt/atp5mf/atp5pd/atp5po/atp5mea/atp5mc1/nme2b.2/nme6/atp5f1c/mrpl24/atp5meb/nme2a/si:dkey-190g11.3/antkmt/atp5f1e/nme3/atp5pf/atp5pb/atp5mc3b/nme7/atp5fa1/dlst/ATP6 |
| GO:0009206 | purine ribonucleoside triphosphate biosynthetic process | 1.93013788013767 | 0.00988355843878473 | nme9/uqcc3/atp5mc3a/atpsckmt/atp5mf/atp5pd/atp5po/atp5mea/atp5mc1/nme2b.2/nme6/atp5f1c/mrpl24/atp5meb/nme2a/si:dkey-190g11.3/antkmt/atp5f1e/nme3/atp5pf/atp5pb/atp5mc3b/nme7/atp5fa1/dlst/ATP6 |
| GO:0008202 | steroid metabolic process | 1.89342793339491 | 0.000798277428982877 | ch25h/sdr42e1/dhcr24/cyp46a1.1/ch25hl1.1/msmo1/amacr/nsdhl/cyp46a1.3/cyp39a1/cyp46a1.2/hmgcra/mvk/ebpl/hsd17b12b/bdh1/srd5a1/cyp51/gba/cyp21a2/cyp7a1/lipeb/slc27a2b/faxdc2/cyp46a1.4/apoea/cyp2r1/mvda/ch25hl2/fgfr4/lss/hmgcs1/ebp/cyp26c1/apoa1a/cyp27a1.4/hsd17b7/cyb5r2/cyp27a1.1/hsd17b14/rdh1/srd5a2b/erlin1/slc27a2a |
| GO:0006979 | response to oxidative stress | 1.83959019549846 | 0.00355248534040968 | bco2l/fancc/anxa1b/glrx2/keap1a/per2/prdx1/ptgs2a/sod2/park7/ptgs1/ptgs2b/lonp1/prdx3/si:ch211-120k19.1/trpm2/gpx4a/diabloa/impact/gpx1a/gpx9/hmox2a/gpx4b/gpx8/naprt/mpv17/brf2/sirt2/romo1/sod1/txndc12/ndufa6/mpv17l2/hmox1a/ryr1b/msraa |
| GO:0006260 | DNA replication | 1.7556677318185 | 0.00194590276805455 | dbf4/zranb3/polq/zgc:173742/wrn/lig4/pold3/rpa3/mcm8/wdr18/tonsl/terf1/rmi1/poln/ctc1/mgme1/rnaseh1/ssrp1a/gins3/rpa2/stn1/orc4/dna2/noc3l/mcm7/rfc5/gins2/orc5/tatdn1/pole2/polg2/rrm1/tipin/fen1/primpol/mcm3/rrm2/traip/orc1/lig1/igfbp2a/mcmbp/mcm2/wrnip1/twnk/recql/rad51/rfc2/mcm5/rpa1/esco2/cenps/mms22l/eme1/orc6/rbbp4/gins4/blm/rnaseh2a/cdc45/prim1/rfc3/helb/pcna/rbb4l/lrwd1/samhd1/orc3/dscc1/ssbp1/top1a/mcm4/pole |
| GO:0044743 | protein transmembrane import into intracellular organelle | 1.73136835035816 | 0.0470692918806812 | pex10/chchd4b/timm21/chchd4a/timm50/dnajc19/grpel1/hspd1/tomm40/romo1/pex1/dnlz/pex13/tomm40l/pex5/timm23a/tomm70a/tomm7/pex7/pam16/tomm20b/pex26 |
| GO:0050731 | positive regulation of peptidyl-tyrosine phosphorylation | 1.72535656967289 | 0.0678490288954661 | unc119a/epgn/arl2bp/btc/smtlb/hbegfa/zgc:153759 |
| GO:0034599 | cellular response to oxidative stress | 1.71630178467435 | 0.0639818065496334 | bco2l/fancc/anxa1b/glrx2/keap1a/prdx1/sod2/park7/lonp1/si:ch211-120k19.1/diabloa/impact/mpv17/brf2/sirt2/romo1/sod1/mpv17l2/ryr1b/msraa |
| GO:0042744 | hydrogen peroxide catabolic process | 1.71035395380329 | 0.0737625497040733 | prdx1/hbae5/zgc:163057/hbbe2/prdx3/hbae1.3/hbae1.1/hbae3/gpx1a |
| GO:0006974 | cellular response to DNA damage stimulus | 1.70302954476004 | 6.9422407694291e-06 | uba7/wdr76/hrob/mms19/ddias/cry5/slx1b/zswim7/fancc/zranb3/tdp2b/xpc/parpbp/rad51d/swsap1/chek2/cry-dash/si:ch211-127m7.2/msh3/palb2/polq/hic1/phkg1b/xndc1/neil1/rfwd3/rrm2b/ogg1/sfr1/wrn/fanci/rad54b/brip1/cpdp/lig4/pargl/mre11a/zgc:112496/rad51b/faap100/pold3/rpa3/atm/cdc7/mcm8/nthl1/mcm9/ddb2/ap5s1/ercc2/kmt5ab/aptx/nbn/tonsl/cep63/eya4/rmi1/alkbh7/poln/mnat1/si:dkey-74k8.3/mgme1/park7/dek/mpg/smyd2a/mettl3/spata18/polh/ssrp1a/tdp1/neil3/rpa2/stn1/poldip2/xrcc3/gtf2h3/trip13/zfyve26/fancg/msh2/dna2/mcm7/smarcad1a/rfc5/uba1/swi5/gins2/fxr2/xrcc6/rad51c/pole2/polg2/tipin/yars1/ercc6/ube2nb/prmt6/fen1/primpol/tigara/mcm3/unga/sem1/bcl2l16/rad23b/pmaip1/lig1/polr2i/bccip/nek4/vhl/mcm2/abraxas1/txndc12/rbbp8/ttf2/mlh1/rmi2/wrnip1/recql/poli/rad1/rad51/rfc2/mcm5/rpa1/ercc1/plrg1/smyd2b/cenps/psme4a/mms22l/eme1/cdca5/si:ch211-141o9.10/hltf |
| GO:0044283 | small molecule biosynthetic process | 1.67134004649573 | 0.000341676076873402 | gls2b/tecrl2b/pycr3/fads2/mcat/msmo1/sdsl/asns/gatm/chst15/elovl8a/gpib/anxa1b/prxl2b/mthfr/acacb/cyp39a1/elovl2/pycr1b/hmgcra/mvk/gch2/qdprb.1/otc/uros/ptgs2a/olah/hsd17b12b/glsl/coq5/aass/akr1a1b/srd5a3/moxd1l/pdss2/asnsd1/srr/cyp51/cyp7a1/adka/park7/lgsn/slc27a2b/apip/rfk/ptgs1/ptgs2b/mvda/pnpo/abhd2a/pcbd1/kmo/ptgesl/pah/elovl6l/fgfr4/tecra/asah2/lss/hmgcs1/ebp/apoa1a/cyp27a1.4/gch1/acmsd/hsd17b7/gnmt/coq4/pck2/elovl8b/sephs1/cdo1/ptges3b/gamt/cyp27a1.1/ndufab1b/fbp1b/impa1/bhmt/upb1/acot7/rgn/fa2h/slc27a2a/aprt/sptlc3/prtfdc1b/agxtb/g6pc3/hprt1/pdss1/ubiad1/coq6/cth/ndufa9a/qdpra |
| GO:0052646 | alditol phosphate metabolic process | 1.62136572553731 | 0.0983308153791125 | gpd1l/rfk/gpam/gpd1c/gpd1a/gpd2 |
| GO:0032259 | methylation | 1.61555246309617 | 0.00489199162953128 | hnmt/thumpd2/suv39h1a/zgc:113054/mettl21cb/wdr4/tpmt.2/etfbkmt/mrm3a/dnmt3bb.2/eef1akmt1/cskmt/trmt10c/dnmt3bb.3/emg1/ash2l/tyms/kmt5ab/henmt1/piwil2/coq5/trmt61a/vcpkmt/mrm3b/trmt44/atpsckmt/mettl4/trmt6/tyw3/trmt2a/dph5/trmt12/nsun2/eef1akmt2/thumpd3/tfb1m/prmt9/smyd2a/tpmt.1/hells/prmt7/mettl3/smyd1a/prmt3/trmt11/trmt112/dimt1l/bud23/ntmt1/mettl16/mettl21a/mettl2a/thada/nsun5/gtpbp3/amt/trmt10b/tdrd9/dnmt3bb.1/gnmt/zcchc4/gamt/cebpb/si:dkey-190g11.3/antkmt/cxxc1b/rrp8/mrm2/setd3/bhmt/prmt6/skic8/trmt61b/comta/lyn/unga/nop2/comtd1 |
| GO:0043436 | oxoacid metabolic process | 1.61491700112608 | 2.31796694375566e-05 | gls2b/tecrl2b/pycr3/fars2/fads2/ddo/mcat/farsa/amacr/sdsl/accs/etfbkmt/asns/si:dkey-238o13.4/gatm/elovl8a/gpib/anxa1b/acot20/prxl2b/dars2/acot17/wars2/mthfr/asah1a/acot11b/ugl/sdhaf3/acacb/cyp39a1/elovl2/hoga1/pycr1b/dars1/nit2/qdprb.1/gatd1/otc/abcd3b/uros/ptgs2a/olah/hsd17b12b/glsl/tdo2a/gcat/enosf1/uevld/aass/amdhd2/alkbh7/asnsd1/srr/mdh1b/mdh1ab/dhtkd1/marc1/lars1b/lpin1b/cyp7a1/tysnd1/park7/si:ch211-256m1.8/acaa1/tat/acads/lgsn/slc27a2b/mars2/apip/ptgs1/idh3b/ptgs2b/pnpo/tph1b/gpam/abhd2a/hacl1/hsd17b4/cpt2/adpgk2/kyat1/decr1/kmo/qrsl1/bpgm/ptgesl/gck/pah/elovl6l/si:ch211-212d10.2/gatb/zgc:162816/fgfr4/tecra/gtpbp3/asah2/iars1/amt/etfa/cpt1b/hars/acss1/gcshb/cmah/cyp26c1/ctps1a/acadm/qars1/nars2/vars1/hadhab/auh/bckdha/acmsd/gnmt/pck2/elovl8b/sephs1/cdo1/mrps36/ptges3b/eno1b/abcd3a/gamt/pgam2/gnpda1/ndufab1b/mccc2/pklr/yars1/aacs/hadh/gcdha/me1/bhmt/hpdb/upb1/acot7/rgn/rars1/tigara/fa2h/slc27a2a/kyat3.2/hibch/mpst/acss2/echdc2/eci1/hgd/agxtb/pm20d1.1/tars1/hadhaa/gars1/pars2/cth/qdpra/ptgr1.1/iars2/acsf2/aspg/gcsha/kynu/hadhb/acsl5/glud1b/got2b/hpdl/cpt1aa/hk2/idh1/acsl1b/sars1/ivd/acat2/vars2/acaa2/ehhadh/acot18/nags/amdhd1/sars2/phyh/ftcd/sgpl1/gip/slc25a44b/shmt1/pfas/acot15/mgst2/aldoab/elovl4b/idh3a/tecrb/aars1/ldha/elovl7a |
| GO:0044282 | small molecule catabolic process | 1.59271540808738 | 0.00441611574194559 | gls2b/cyp46a1.1/ddo/sdsl/etfbkmt/si:dkey-238o13.4/cyp46a1.3/bpnt1/cyp46a1.2/hoga1/qdprb.1/gatd1/abcd3b/glsl/tdo2a/gcat/enosf1/aass/srd5a3/amdhd2/srr/zgc:103586/rbks/miox/lpin1b/cyp7a1/tysnd1/acaa1/tat/acads/cyp46a1.4/apoea/uraha/sord/abhd2a/hacl1/hsd17b4/cpt2/decr1/kmo/pah/zgc:162816/amt/etfa/cpt1b/gcshb/cdaa/acadm/hadhab/auh/bckdha/pck2/cdo1/abcd3a/adh8b/gnpda1/mccc2/impa1/hadh/gcdha/hpdb/kyat3.2/hibch/echdc2/eci1/hgd/hadhaa/qdpra/gcsha/kynu/hadhb/glud1b/got2b/esd/cpt1aa/ivd/acat2/acaa2/ehhadh/amdhd1/phyh/ftcd/slc25a44b/shmt1 |
| GO:1901361 | organic cyclic compound catabolic process | 1.58649464452635 | 0.00292032014134612 | hnmt/nudt4b/cyp46a1.1/dazl/dnase1l1/blvra/fastkd3/fitm2/zc3h12ab/fastkd2/cyp46a1.3/tbrg4/nudt18/exosc2/cyp46a1.2/fastkd1/exosc6/qdprb.1/entpd3/dcp1b/pde4cb/tdo2a/nudt15/moxd1l/zgc:103586/dut/heatr1/mrto4/dcps/exosc5/cyp7a1/gda/acaa1/tat/dctpp1/cyp46a1.4/tent5bb/apoea/rnaseh1/mettl3/uraha/lsm1/magoh/kmo/lsm3/pah/rexo4/ago3b/mettl16/gtpbp3/xdh/cdaa/dna2/nudt4a/fxr2/cnot8/rnaset2l/entpd5b/dnase1l4.1/hsd17b14/hmox2a/fhit/hint1/dpys/hpdb/comta/fen1/kyat3.2/pde12/allc/hgd/cnot6l/edc3/nbas/dis3/qdpra/plex9.2/rc3h2/nudt12/kynu/tet2/syncripl/endog/hmox1a/mtrex/entpd5a |
| GO:0071826 | ribonucleoprotein complex subunit organization | 1.57792704614773 | 0.0130343312989884 | nle1/gemin5/zfand1/abt1/znhit3/pwp2h/trnau1apa/mrto4/snrpd2/prmt7/pih1d1/gemin2/mrpl20/snrpg/bxdc2/prpf18/mcts1/ppan/mdn1/clp1/ptges3b/rrp7a/eif3m/nopchap1/snrpd3l/eif3ea/coil/eif6/snrpf/usp39/eif3jb/snrpe/rpl38/rps14/shq1/rpl5a/eif5/eif3ha/trnau1apb/mrps11/scaf11/rpf2/eif3s10/cdc73/denr/ddx28/naf1/rpl23a/eif3s6ip/rps15/rpsa/eif2s2/polr2d/srpk1a/rps5/gemin8/rplp0/snrpd1/rps27.1/aar2/rpl10/rpl5b/eif3g/eif3d/rpl6/yju2/rps27.2/rps28/mrps7/gemin6/eif2s3/nudt21/dyrk3/lsm4/snrpc/rps19/eif3ja/eif3c |
| GO:0055088 | lipid homeostasis | 1.54238929377417 | 0.0961825238314319 | adck1/fitm2/zgc:162608/cyp39a1/spx/abcg4b/ormdl2/apobb.1/psmd10/apobb.2/cyp7a1/apoa4a/pnpla3/tlcd1/apoa4b.2/apoba/fabp2/angptl8 |
| GO:0140014 | mitotic nuclear division | 1.53553828880172 | 0.0229731377900925 | ncaph/epgn/btc/pdcd2/nde1/chtf8/si:ch211-244o22.2/cep192/sgo1/map10/mad1l1/incenp/gnl3/kif20bb/ndc80/anapc5/gcna/ncapg/bub3/nme6/mad2l1/reep3b/anapc15/birc5b/cdca8/esco1/ncapg2/abraxas2/ccnb1/ranbp1/aurkb/racgap1/anapc11/ube2s/zgc:86764/pmaip1/bccip/abraxas1/zw10/esco2/ncaph2/cenpk/spdl1/prc1b/ran/cdca5/gnl2/map9/mki67/kifc1/ube2ib/chmp7/kif11 |
| GO:0019439 | aromatic compound catabolic process | 1.4470736645875 | 0.0239204409208845 | hnmt/nudt4b/dazl/dnase1l1/blvra/fastkd3/fitm2/zc3h12ab/fastkd2/tbrg4/nudt18/exosc2/fastkd1/exosc6/qdprb.1/entpd3/dcp1b/pde4cb/tdo2a/nudt15/moxd1l/zgc:103586/dut/heatr1/mrto4/dcps/exosc5/gda/acaa1/tat/dctpp1/tent5bb/rnaseh1/mettl3/uraha/lsm1/magoh/kmo/lsm3/pah/rexo4/ago3b/mettl16/gtpbp3/xdh/cdaa/dna2/nudt4a/fxr2/cnot8/rnaset2l/entpd5b/dnase1l4.1/hmox2a/fhit/hint1/hpdb/comta/fen1/kyat3.2/pde12/hgd/cnot6l/edc3/nbas/dis3/qdpra/plex9.2/rc3h2/nudt12/kynu/tet2/syncripl/endog/hmox1a/mtrex/entpd5a/ephx1 |
| GO:0044270 | cellular nitrogen compound catabolic process | 1.42325048500041 | 0.0467129045300582 | hnmt/nudt4b/dazl/dnase1l1/blvra/fastkd3/fitm2/zc3h12ab/fastkd2/tbrg4/nudt18/exosc2/fastkd1/exosc6/entpd3/dcp1b/pde4cb/tdo2a/nudt15/zgc:103586/dut/heatr1/mrto4/dcps/exosc5/gda/dctpp1/tent5bb/rnaseh1/mettl3/uraha/lsm1/magoh/kmo/lsm3/rexo4/ago3b/mettl16/gtpbp3/xdh/cdaa/dna2/nudt4a/fxr2/cnot8/rnaset2l/entpd5b/dnase1l4.1/hmox2a/fhit/hint1/dpys/bhmt/fen1/pde12/cnot6l/edc3/nbas/dis3/plex9.2/rc3h2/nudt12/kynu/tet2/syncripl/endog/hmox1a/mtrex/entpd5a |
| GO:0007601 | visual perception | 1.4025303608205 | 0.0873969864431149 | unc119a/gprc5c/crygm2a/crybb2/crybb1l3/crygm5/grk7a/crygm2d6/crygm2b/cryba1l2/arr3b/crygm2d9/crygm2d15/crygm2d12/crygm2d18/cryba2a/opn1sw1/crygm2d17/crygm2d7/crygm2d1/arl6/crygm2d5/crygm2f/cryba1l1/pdcb/crybb1l2/crygm2d3/crygm2d20/crygm2d11/crygm2d10/crygm2d2/crygm2d21/cryba1a/opn1mw1/crygm2d13/crygmxl2/guca1b/crygn2/crygm2d16/arr3a/cryba4/crygm2d8/crygm2d14/cryba1b/crygn1/prph2a/lum/cryba2b/opn1sw2/crygmx/pdca/crygmxl1/si:dkey-57a22.15/crybb1l1/pde6a/gucy2f/grk7b/opn1lw2/nphp3/prph2lb/pde6d/crybgx/tmtops2a |
| GO:0099537 | trans-synaptic signaling | -2.69921105199218 | 1.14653846153846e-08 | dlg1b/dtnbp1b/ncdn/gabrr2a/gabrb1/cplx2b/drd4-rs/cacng8b/otofa/sv2bb/dlg3/cdh2/penka/cep89/chrnb5a/synj1/cart2/cspg5b/gria3a/dlgap1a/chrm2a/snap25a/napbb/penkb/glrbb/slc5a7a/napba/chata/spg11/apba1b/erc1a/grid1a/stxbp1b/cplx2a/clstn2b/cacnb1/gabrb3/chrng/hdc/sncb/glra1/drd3/cacng5a/si:ch211-81a5.8/syn1/erc2/gria1a/grm4/grm6b/stx3a/caskb/chrnb3a/ppp1r9ba/slc18a3a/snap25b/jph3b/mctp2b/grm8b/cart3/slc6a5/calb1/chrna3/htr1aa/calb2b/snap47/htr1d/pdyn/jph3a/cacnb2b/ptchd1/calb2a/syt7b/syn2a/cacng7b/chrna5/grin2aa/rims1b/shisa9a/clstn1/syt4/rab3ab/dlgap4b/lrrc4ca/chrna2a/gabra4/glra4b/nlgn4xa/gria2b/nsmfb/gria4a/slc17a6b/clstn2a/sncgb/htr3a/gabrb2a/lrrc4.1/cadpsb/grin1a/grm2a/ppp1r9bb/syt11a/unc13a/cacng2b/stxbp1a/syt1a/chrnb2/slc12a5b/chrna6/cacng3b/shisa7b/nlgn3a/gria3b/stx1b/dlg2/ppfia3/grin2ab/cacng2a/grm8a/lrrc4cb/atcaya/slc12a5a/chrna4b/grm2b/nlgn3b/cacnb4b/grid2/clstn3/syt2a/apba2b/grik1a/cspg5a/sv2a/pnocb/grin1b/baiap3/cacna1ab/grin2da/nlgn2a/nrxn2a/si:dkey-174m14.3/drd2a/rph3ab/dagla/grik5/htr1fa/shisa9b/gabrp/chrm4a/grin3bb/grin2bb/insyn1/grik4/gabrg2/grm1a/cacng8a/stx11a/gabrb4/dlgap3/htr7c/neto1l/nalcn/gria4b |
| GO:0007268 | chemical synaptic transmission | -2.68729704840182 | 1.14653846153846e-08 | dlg1b/dtnbp1b/ncdn/gabrr2a/gabrb1/cplx2b/drd4-rs/cacng8b/otofa/sv2bb/dlg3/cdh2/penka/cep89/chrnb5a/synj1/cart2/cspg5b/gria3a/dlgap1a/chrm2a/snap25a/napbb/penkb/glrbb/slc5a7a/napba/chata/spg11/apba1b/erc1a/grid1a/stxbp1b/cplx2a/clstn2b/cacnb1/gabrb3/chrng/hdc/sncb/glra1/drd3/cacng5a/si:ch211-81a5.8/syn1/erc2/gria1a/grm4/grm6b/stx3a/caskb/chrnb3a/ppp1r9ba/slc18a3a/snap25b/jph3b/mctp2b/grm8b/cart3/slc6a5/calb1/chrna3/htr1aa/calb2b/snap47/htr1d/pdyn/jph3a/cacnb2b/ptchd1/calb2a/syt7b/syn2a/cacng7b/chrna5/grin2aa/rims1b/shisa9a/clstn1/syt4/rab3ab/dlgap4b/lrrc4ca/chrna2a/gabra4/glra4b/nlgn4xa/gria2b/nsmfb/gria4a/slc17a6b/clstn2a/sncgb/htr3a/gabrb2a/lrrc4.1/cadpsb/grin1a/grm2a/ppp1r9bb/syt11a/unc13a/cacng2b/stxbp1a/syt1a/chrnb2/slc12a5b/chrna6/cacng3b/shisa7b/nlgn3a/gria3b/stx1b/dlg2/ppfia3/grin2ab/cacng2a/grm8a/lrrc4cb/atcaya/slc12a5a/chrna4b/grm2b/nlgn3b/cacnb4b/grid2/clstn3/syt2a/apba2b/grik1a/cspg5a/sv2a/pnocb/grin1b/baiap3/cacna1ab/grin2da/nlgn2a/nrxn2a/si:dkey-174m14.3/drd2a/rph3ab/grik5/htr1fa/shisa9b/gabrp/chrm4a/grin3bb/grin2bb/insyn1/grik4/gabrg2/grm1a/cacng8a/stx11a/gabrb4/dlgap3/htr7c/neto1l/nalcn/gria4b |
| GO:0098916 | anterograde trans-synaptic signaling | -2.68729704840182 | 1.14653846153846e-08 | dlg1b/dtnbp1b/ncdn/gabrr2a/gabrb1/cplx2b/drd4-rs/cacng8b/otofa/sv2bb/dlg3/cdh2/penka/cep89/chrnb5a/synj1/cart2/cspg5b/gria3a/dlgap1a/chrm2a/snap25a/napbb/penkb/glrbb/slc5a7a/napba/chata/spg11/apba1b/erc1a/grid1a/stxbp1b/cplx2a/clstn2b/cacnb1/gabrb3/chrng/hdc/sncb/glra1/drd3/cacng5a/si:ch211-81a5.8/syn1/erc2/gria1a/grm4/grm6b/stx3a/caskb/chrnb3a/ppp1r9ba/slc18a3a/snap25b/jph3b/mctp2b/grm8b/cart3/slc6a5/calb1/chrna3/htr1aa/calb2b/snap47/htr1d/pdyn/jph3a/cacnb2b/ptchd1/calb2a/syt7b/syn2a/cacng7b/chrna5/grin2aa/rims1b/shisa9a/clstn1/syt4/rab3ab/dlgap4b/lrrc4ca/chrna2a/gabra4/glra4b/nlgn4xa/gria2b/nsmfb/gria4a/slc17a6b/clstn2a/sncgb/htr3a/gabrb2a/lrrc4.1/cadpsb/grin1a/grm2a/ppp1r9bb/syt11a/unc13a/cacng2b/stxbp1a/syt1a/chrnb2/slc12a5b/chrna6/cacng3b/shisa7b/nlgn3a/gria3b/stx1b/dlg2/ppfia3/grin2ab/cacng2a/grm8a/lrrc4cb/atcaya/slc12a5a/chrna4b/grm2b/nlgn3b/cacnb4b/grid2/clstn3/syt2a/apba2b/grik1a/cspg5a/sv2a/pnocb/grin1b/baiap3/cacna1ab/grin2da/nlgn2a/nrxn2a/si:dkey-174m14.3/drd2a/rph3ab/grik5/htr1fa/shisa9b/gabrp/chrm4a/grin3bb/grin2bb/insyn1/grik4/gabrg2/grm1a/cacng8a/stx11a/gabrb4/dlgap3/htr7c/neto1l/nalcn/gria4b |
| GO:0050804 | modulation of chemical synaptic transmission | -2.68416739586281 | 1.14653846153846e-08 | erc1a/grid1a/cplx2a/clstn2b/drd3/cacng5a/si:ch211-81a5.8/erc2/gria1a/grm4/grm6b/caskb/ppp1r9ba/jph3b/mctp2b/grm8b/calb1/calb2b/jph3a/calb2a/cacng7b/grin2aa/rims1b/shisa9a/clstn1/dlgap4b/lrrc4ca/nlgn4xa/gria2b/nsmfb/gria4a/clstn2a/lrrc4.1/grm2a/ppp1r9bb/cacng2b/cacng3b/shisa7b/nlgn3a/gria3b/stx1b/grin2ab/cacng2a/grm8a/lrrc4cb/atcaya/grm2b/nlgn3b/grid2/clstn3/grik1a/baiap3/grin2da/nlgn2a/si:dkey-174m14.3/drd2a/grik5/shisa9b/grin3bb/grin2bb/grik4/grm1a/cacng8a/dlgap3/neto1l/nalcn/gria4b |
| GO:0099177 | regulation of trans-synaptic signaling | -2.68416739586281 | 1.14653846153846e-08 | erc1a/grid1a/cplx2a/clstn2b/drd3/cacng5a/si:ch211-81a5.8/erc2/gria1a/grm4/grm6b/caskb/ppp1r9ba/jph3b/mctp2b/grm8b/calb1/calb2b/jph3a/calb2a/cacng7b/grin2aa/rims1b/shisa9a/clstn1/dlgap4b/lrrc4ca/nlgn4xa/gria2b/nsmfb/gria4a/clstn2a/lrrc4.1/grm2a/ppp1r9bb/cacng2b/cacng3b/shisa7b/nlgn3a/gria3b/stx1b/grin2ab/cacng2a/grm8a/lrrc4cb/atcaya/grm2b/nlgn3b/grid2/clstn3/grik1a/baiap3/grin2da/nlgn2a/si:dkey-174m14.3/drd2a/grik5/shisa9b/grin3bb/grin2bb/grik4/grm1a/cacng8a/dlgap3/neto1l/nalcn/gria4b |
| GO:0099536 | synaptic signaling | -2.6833837570563 | 1.14653846153846e-08 | dlg1b/dtnbp1b/ncdn/gabrr2a/gabrb1/cplx2b/drd4-rs/cacng8b/otofa/sv2bb/dlg3/cdh2/penka/cep89/chrnb5a/synj1/cart2/cspg5b/gria3a/dlgap1a/chrm2a/snap25a/napbb/penkb/glrbb/slc5a7a/napba/chata/spg11/apba1b/erc1a/grid1a/stxbp1b/cplx2a/clstn2b/cacnb1/gabrb3/chrng/hdc/sncb/glra1/drd3/cacng5a/si:ch211-81a5.8/syn1/erc2/gria1a/grm4/grm6b/stx3a/caskb/chrnb3a/ppp1r9ba/slc18a3a/snap25b/jph3b/mctp2b/grm8b/cart3/slc6a5/calb1/chrna3/htr1aa/calb2b/snap47/htr1d/pdyn/jph3a/cacnb2b/ptchd1/calb2a/syt7b/syn2a/cacng7b/chrna5/grin2aa/rims1b/shisa9a/clstn1/syt4/rab3ab/dlgap4b/lrrc4ca/chrna2a/gabra4/glra4b/nlgn4xa/gria2b/nsmfb/gria4a/slc17a6b/clstn2a/sncgb/htr3a/gabrb2a/lrrc4.1/cadpsb/grin1a/grm2a/ppp1r9bb/syt11a/unc13a/cacng2b/stxbp1a/syt1a/chrnb2/slc12a5b/chrna6/cacng3b/shisa7b/nlgn3a/gria3b/stx1b/dlg2/ppfia3/grin2ab/cacng2a/grm8a/lrrc4cb/atcaya/slc12a5a/chrna4b/grm2b/nlgn3b/cacnb4b/grid2/clstn3/syt2a/apba2b/grik1a/cspg5a/sv2a/pnocb/grin1b/baiap3/cacna1ab/grin2da/nlgn2a/nrxn2a/si:dkey-174m14.3/drd2a/rph3ab/dagla/grik5/htr1fa/shisa9b/gabrp/chrm4a/grin3bb/grin2bb/insyn1/grik4/gabrg2/grm1a/cacng8a/stx11a/gabrb4/scn1lab/dlgap3/htr7c/neto1l/nalcn/gria4b |
| GO:0098742 | cell-cell adhesion via plasma-membrane adhesion molecules | -2.34722281326859 | 1.14653846153846e-08 | arvcfb/zeb1a/cdh2/prnprs3/ptk7a/pcdh2ab10/zgc:165604/si:ch211-186j3.6/tenm3/dchs1a/ret/pcdh8/ptprsa/clstn2b/dscama/pcdh17/pcdh2ab9/pecam1a/pcdh10b/pcdh7a/tenm2b/robo3/si:ch211-199f5.1/dsc2l/pcdh1g22/jag2b/nectin3b/tenm4/pcdh1g31/pcdh7b/pcdh1g29/pcdh2g13/pcdh1a/clstn1/pcdh19/cadm1b/lrrc4ca/pcdh1a4/pcdh2ab3/cadm3/dscamb/ptprdb/pcdh10a/pcdh11/clstn2a/lrrc4.1/igsf21b/robo4/fat3a/cadm4/si:ch211-214p13.3/cdh18a/si:dkey-237h12.3/lrrc4cb/igsf21a/tenm2a/esama/cdh13/clstn3/cdh15/cntn3a.1/si:ch73-233f7.1/pcdh1g32/cdh4/pcdh1g11/si:ch73-233f7.3/cdh7a/pcdh1g18/adgrl1a/pcdh1gc5 |
| GO:0050808 | synapse organization | -2.21669824394954 | 9.94928637272983e-08 | dbn1/ctnna2/cdh2/synj1/ptk7a/dbnlb/cbln2b/ephb3a/sez6a/tjp1b/ctbp2a/erc1a/fbxo45/gphnb/neurod2/ptprsa/clstn2b/cacnb1/sncb/erc2/tanc2a/ppfia4/gfpt1/slitrk4/gphna/cacnb2b/myo9ab/inpp5ka/caprin2/pcloa/clstn1/rab3ab/lrrc4ca/ptprdb/nlgn4xa/clstn2a/sncgb/sybu/lrrc4.1/arhgef9b/unc13a/dock10/ctnnd2b/nbeaa/nlgn3a/bsnb/cacna1da/ppfia3/lrrc4cb/epb41l3a/grna/bdnf/nlgn3b/ctnnd2a/cacnb4b/pclob/clstn3/slitrk2/sez6l2/nlgn2a/nrxn2a |
| GO:0048667 | cell morphogenesis involved in neuron differentiation | -2.08310048291656 | 1.14653846153846e-08 | plxnb1b/pik3cd/mycbp2/rb1/ptprfb/zgc:100846/ptprfa/isl2b/nrp2b/arhgap35a/rapgef1a/whrna/emb/pik3ca/rbpms2a/aplp2/mapk8ip3/npr1a/sema3fa/emx2/olfm1a/unc5b/nptna/vegfc/dbn1/kctd12.1/si:ch211-129c21.1/phox2bb/unk/ubap1/ctnna2/cntn4/cdh2/trarg1a/kalrnb/ulk2/ptk7a/dbnlb/efna1b/ephb3a/slc35b2/rbpms2b/fus/map1ab/dock7/foxb1a/nsfa/rab33ba/sipa1l1/nrp2a/ush1ga/pnpla6/kctd13/isl1a/fezf2/slit2/tenm3/rufy3/nfasca/hecw1b/sim1a/adcy8/pak1/spg11/ret/kif1b/arnt2/sema3bl/zdhhc17/plxna4/appa/nedd4l/arhgap35b/efna2a/dscama/apc/robo2/pcdh17/casp9/epha4a/hdc/anos1a/trioa/lhx9/epha7/rtn4r/cyfip2/dpysl3/draxinb/slit1b/robo3/usp33/map1sb/appb/tanc2a/dzip1/sema5a/kifbp/chad/robo1/tnr/whrnb/tenm4/hecw2a/epha4l/kif5c/slitrk4/islr2/nptnb/lhx2b/myo9ab/caprin2/kalrna/cdkl5/ephb2b/epha4b/clstn1/rac3a/brsk1b/aplp1/rab3ab/gap43/lrrc4ca/srrm4/lhx1a/map1aa/prmt8b/atl1/nkx2.2a/arhgef25b/dscamb/triob/tmem59l/myo10l3/nsmfb/slit3/dpysl5a/chst11/fzd3a/kif5aa/robo4/efnb3b/vax1/stat3/dock10/ctnnd2b/zgc:77784/sema4c/si:ch211-113g11.6/rac1l/elavl4/myo10l1/ntn1b/scn8aa/lrrc4cb/plppr4a/grna/bdnf/ctnnd2a/slitrk2/nkx6.1/ncam1b/cntn3a.1/nrxn2a/dagla/grxcr1a/efna3a/dpysl5b/thsd7aa/si:ch211-241e1.3/plxna3/l1camb/plxna1a/nrcama/sema6ba/sema3b |
| GO:0048858 | cell projection morphogenesis | -2.05831448361924 | 1.14653846153846e-08 | plxnb1b/tmem108/pik3cd/mycbp2/rb1/ptprfb/zgc:100846/ptprfa/spag6/isl2b/nrp2b/arhgap35a/rapgef1a/emb/pik3ca/rbpms2a/aplp2/mapk8ip3/npr1a/sema3fa/olfm1a/unc5b/nptna/vegfc/dbn1/kctd12.1/si:ch211-129c21.1/phox2bb/ubap1/ctnna2/cntn4/cdh2/trarg1a/kalrnb/ulk2/ptk7a/dbnlb/cspg5b/efna1b/ephb3a/slc35b2/rbpms2b/fus/map1ab/dock7/foxb1a/nsfa/rab33ba/sipa1l1/tnika/nrp2a/pnpla6/kctd13/isl1a/fezf2/mink1/slit2/tenm3/rufy3/nfasca/hecw1b/sim1a/adcy8/pak1/spg11/ret/kif1b/arnt2/sema3bl/zdhhc17/plxna4/appa/nedd4l/arhgap35b/efna2a/dscama/apc/robo2/pcdh17/casp9/epha4a/hdc/anos1a/trioa/lhx9/epha7/rtn4r/cyfip2/dpysl3/draxinb/slit1b/robo3/usp33/map1sb/appb/tanc2a/dzip1/nyap2b/sema5a/kifbp/chad/pacsin1a/robo1/tnr/tenm4/hecw2a/epha4l/kif5c/slitrk4/islr2/nptnb/lhx2b/myo9ab/caprin2/kalrna/cdkl5/ephb2b/epha4b/clstn1/rac3a/brsk1b/aplp1/rab3ab/gap43/lrrc4ca/srrm4/lhx1a/map1aa/prmt8b/atl1/nkx2.2a/arhgef25b/dscamb/triob/tmem59l/myo10l3/gas7a/nsmfb/slit3/dpysl5a/chst11/fzd3a/kif5aa/robo4/efnb3b/vax1/stat3/dock10/ctnnd2b/zgc:77784/sema4c/si:ch211-113g11.6/rac1l/elavl4/myo10l1/ntn1b/scn8aa/lrrc4cb/plppr4a/grna/bdnf/ctnnd2a/nrn1la/slitrk2/nkx6.1/cspg5a/ncam1b/cntn3a.1/nrxn2a/dagla/efna3a/dpysl5b/inab/thsd7aa/si:ch211-241e1.3/plxna3/l1camb/plxna1a/nrcama/sema6ba/sema3b |
| GO:0048812 | neuron projection morphogenesis | -2.04924995520432 | 1.14653846153846e-08 | plxnb1b/tmem108/pik3cd/mycbp2/rb1/ptprfb/zgc:100846/ptprfa/spag6/isl2b/nrp2b/arhgap35a/rapgef1a/emb/pik3ca/rbpms2a/aplp2/mapk8ip3/npr1a/sema3fa/olfm1a/unc5b/nptna/vegfc/dbn1/kctd12.1/si:ch211-129c21.1/phox2bb/ubap1/ctnna2/cntn4/cdh2/trarg1a/kalrnb/ulk2/ptk7a/dbnlb/efna1b/ephb3a/slc35b2/rbpms2b/fus/map1ab/dock7/foxb1a/nsfa/rab33ba/sipa1l1/tnika/nrp2a/pnpla6/kctd13/isl1a/fezf2/mink1/slit2/tenm3/rufy3/nfasca/hecw1b/sim1a/adcy8/pak1/spg11/ret/kif1b/arnt2/sema3bl/zdhhc17/plxna4/appa/nedd4l/arhgap35b/efna2a/dscama/apc/robo2/pcdh17/casp9/epha4a/hdc/anos1a/trioa/lhx9/epha7/rtn4r/cyfip2/dpysl3/draxinb/slit1b/robo3/usp33/map1sb/appb/tanc2a/dzip1/nyap2b/sema5a/kifbp/chad/pacsin1a/robo1/tnr/tenm4/hecw2a/epha4l/kif5c/slitrk4/islr2/nptnb/lhx2b/myo9ab/caprin2/kalrna/cdkl5/ephb2b/epha4b/clstn1/rac3a/brsk1b/aplp1/rab3ab/gap43/lrrc4ca/srrm4/lhx1a/map1aa/prmt8b/atl1/nkx2.2a/arhgef25b/dscamb/triob/tmem59l/myo10l3/gas7a/nsmfb/slit3/dpysl5a/chst11/fzd3a/kif5aa/robo4/efnb3b/vax1/stat3/dock10/ctnnd2b/zgc:77784/sema4c/si:ch211-113g11.6/rac1l/elavl4/myo10l1/ntn1b/scn8aa/lrrc4cb/plppr4a/grna/bdnf/ctnnd2a/nrn1la/slitrk2/nkx6.1/ncam1b/cntn3a.1/nrxn2a/dagla/efna3a/dpysl5b/inab/thsd7aa/si:ch211-241e1.3/plxna3/l1camb/plxna1a/nrcama/sema6ba/sema3b |
| GO:0120039 | plasma membrane bounded cell projection morphogenesis | -2.0490209096944 | 1.14653846153846e-08 | plxnb1b/tmem108/pik3cd/mycbp2/rb1/ptprfb/zgc:100846/ptprfa/spag6/isl2b/nrp2b/arhgap35a/rapgef1a/emb/pik3ca/rbpms2a/aplp2/mapk8ip3/npr1a/sema3fa/olfm1a/unc5b/nptna/vegfc/dbn1/kctd12.1/si:ch211-129c21.1/phox2bb/ubap1/ctnna2/cntn4/cdh2/trarg1a/kalrnb/ulk2/ptk7a/dbnlb/efna1b/ephb3a/slc35b2/rbpms2b/fus/map1ab/dock7/foxb1a/nsfa/rab33ba/sipa1l1/tnika/nrp2a/pnpla6/kctd13/isl1a/fezf2/mink1/slit2/tenm3/rufy3/nfasca/hecw1b/sim1a/adcy8/pak1/spg11/ret/kif1b/arnt2/sema3bl/zdhhc17/plxna4/appa/nedd4l/arhgap35b/efna2a/dscama/apc/robo2/pcdh17/casp9/epha4a/hdc/anos1a/trioa/lhx9/epha7/rtn4r/cyfip2/dpysl3/draxinb/slit1b/robo3/usp33/map1sb/appb/tanc2a/dzip1/nyap2b/sema5a/kifbp/chad/pacsin1a/robo1/tnr/tenm4/hecw2a/epha4l/kif5c/slitrk4/islr2/nptnb/lhx2b/myo9ab/caprin2/kalrna/cdkl5/ephb2b/epha4b/clstn1/rac3a/brsk1b/aplp1/rab3ab/gap43/lrrc4ca/srrm4/lhx1a/map1aa/prmt8b/atl1/nkx2.2a/arhgef25b/dscamb/triob/tmem59l/myo10l3/gas7a/nsmfb/slit3/dpysl5a/chst11/fzd3a/kif5aa/robo4/efnb3b/vax1/stat3/dock10/ctnnd2b/zgc:77784/sema4c/si:ch211-113g11.6/rac1l/elavl4/myo10l1/ntn1b/scn8aa/lrrc4cb/plppr4a/grna/bdnf/ctnnd2a/nrn1la/slitrk2/nkx6.1/ncam1b/cntn3a.1/nrxn2a/dagla/efna3a/dpysl5b/inab/thsd7aa/si:ch211-241e1.3/plxna3/l1camb/plxna1a/nrcama/sema6ba/sema3b |
| GO:0032990 | cell part morphogenesis | -2.03560178882976 | 1.14653846153846e-08 | plxnb1b/tmem108/pik3cd/mycbp2/rb1/ptprfb/zgc:100846/ptprfa/spag6/isl2b/nrp2b/arhgap35a/rapgef1a/emb/pik3ca/rbpms2a/aplp2/mapk8ip3/npr1a/sema3fa/olfm1a/unc5b/nptna/vegfc/dbn1/kctd12.1/si:ch211-129c21.1/phox2bb/ubap1/ctnna2/cntn4/cdh2/trarg1a/kalrnb/ulk2/ptk7a/dbnlb/supv3l1/cspg5b/efna1b/ephb3a/slc35b2/rbpms2b/fus/map1ab/dock7/foxb1a/nsfa/rab33ba/sipa1l1/tnika/nrp2a/pnpla6/kctd13/isl1a/fezf2/mink1/slit2/tenm3/rufy3/nfasca/hecw1b/sim1a/adcy8/pak1/spg11/ret/kif1b/arnt2/sema3bl/zdhhc17/plxna4/appa/nedd4l/arhgap35b/efna2a/dscama/apc/robo2/pcdh17/casp9/epha4a/hdc/anos1a/trioa/lhx9/epha7/rtn4r/cyfip2/dpysl3/draxinb/slit1b/robo3/usp33/map1sb/appb/tanc2a/dzip1/nyap2b/sema5a/kifbp/chad/pacsin1a/robo1/tnr/tenm4/hecw2a/epha4l/kif5c/slitrk4/islr2/nptnb/lhx2b/myo9ab/caprin2/kalrna/cdkl5/ephb2b/epha4b/clstn1/rac3a/brsk1b/aplp1/rab3ab/gap43/lrrc4ca/srrm4/lhx1a/map1aa/prmt8b/atl1/nkx2.2a/arhgef25b/dscamb/triob/tmem59l/myo10l3/gas7a/nsmfb/slit3/dpysl5a/chst11/fzd3a/kif5aa/robo4/efnb3b/vax1/stat3/dock10/ctnnd2b/zgc:77784/sema4c/si:ch211-113g11.6/rac1l/elavl4/myo10l1/ntn1b/scn8aa/lrrc4cb/plppr4a/grna/bdnf/ctnnd2a/nrn1la/slitrk2/nkx6.1/cspg5a/ncam1b/cntn3a.1/nrxn2a/dagla/efna3a/dpysl5b/inab/thsd7aa/si:ch211-241e1.3/plxna3/l1camb/plxna1a/nrcama/sema6ba/sema3b |
| GO:0000904 | cell morphogenesis involved in differentiation | -2.02983448887318 | 1.14653846153846e-08 | pik3cd/mycbp2/rb1/ptprfb/adgrg1/zgc:100846/ptprfa/isl2b/nrp2b/arhgap35a/rapgef1a/whrna/emb/pik3ca/erbb2/rbpms2a/aplp2/mapk8ip3/npr1a/sema3fa/emx2/olfm1a/unc5b/nptna/vegfc/dbn1/kctd12.1/si:ch211-129c21.1/phox2bb/unk/ubap1/ctnna2/cntn4/cdh2/trarg1a/kalrnb/ulk2/ptk7a/dbnlb/efna1b/rilpl2/ephb3a/slc35b2/rbpms2b/fus/map1ab/dock7/foxb1a/nsfa/rab33ba/sipa1l1/nrp2a/ush1ga/pnpla6/kctd13/isl1a/fezf2/slit2/tenm3/rufy3/nfasca/hecw1b/sim1a/adcy8/pak1/spg11/ret/kif1b/arnt2/sema3bl/zdhhc17/plxna4/appa/nedd4l/arhgap35b/efna2a/dscama/apc/robo2/pcdh17/casp9/epha4a/hdc/anos1a/trioa/lhx9/epha7/rtn4r/cyfip2/dpysl3/draxinb/slit1b/robo3/usp33/map1sb/appb/tanc2a/dzip1/sema5a/kifbp/chad/robo1/tnr/whrnb/tenm4/hecw2a/epha4l/kif5c/slitrk4/islr2/nptnb/lhx2b/tet3/myo9ab/caprin2/kalrna/cdkl5/ephb2b/epha4b/clstn1/rac3a/brsk1b/aplp1/rab3ab/gap43/lrrc4ca/srrm4/lhx1a/map1aa/prmt8b/atl1/nkx2.2a/arhgef25b/dscamb/triob/tmem59l/myo10l3/nsmfb/slit3/cxcl8a/dpysl5a/chst11/fzd3a/kif5aa/robo4/efnb3b/vax1/stat3/dock10/ctnnd2b/zgc:77784/sema4c/si:ch211-113g11.6/rac1l/elavl4/myo10l1/ntn1b/scn8aa/lrrc4cb/plppr4a/grna/bdnf/ctnnd2a/slitrk2/nkx6.1/ncam1b/cntn3a.1/nrxn2a/dagla/grxcr1a/efna3a/dpysl5b/thsd7aa/si:ch211-241e1.3/plxna3/l1camb/plxna1a/nrcama/sema6ba/sema3b |
| GO:0007409 | axonogenesis | -2.02083525635161 | 1.14653846153846e-08 | plxnb1b/pik3cd/mycbp2/rb1/ptprfb/zgc:100846/ptprfa/isl2b/nrp2b/arhgap35a/rapgef1a/emb/pik3ca/rbpms2a/aplp2/mapk8ip3/npr1a/sema3fa/olfm1a/unc5b/nptna/vegfc/dbn1/si:ch211-129c21.1/phox2bb/ubap1/ctnna2/cntn4/cdh2/trarg1a/kalrnb/ulk2/ptk7a/dbnlb/efna1b/ephb3a/slc35b2/rbpms2b/fus/map1ab/dock7/foxb1a/nsfa/rab33ba/nrp2a/pnpla6/kctd13/isl1a/fezf2/slit2/tenm3/rufy3/nfasca/sim1a/adcy8/pak1/spg11/ret/kif1b/arnt2/sema3bl/zdhhc17/plxna4/appa/arhgap35b/efna2a/dscama/apc/robo2/pcdh17/casp9/epha4a/hdc/anos1a/trioa/lhx9/epha7/rtn4r/cyfip2/dpysl3/draxinb/slit1b/robo3/usp33/map1sb/appb/dzip1/sema5a/kifbp/chad/robo1/tnr/tenm4/epha4l/kif5c/slitrk4/islr2/nptnb/lhx2b/myo9ab/kalrna/cdkl5/ephb2b/epha4b/clstn1/rac3a/brsk1b/aplp1/rab3ab/gap43/lrrc4ca/srrm4/lhx1a/map1aa/atl1/nkx2.2a/arhgef25b/dscamb/triob/tmem59l/myo10l3/nsmfb/slit3/dpysl5a/chst11/fzd3a/kif5aa/robo4/efnb3b/vax1/stat3/zgc:77784/sema4c/si:ch211-113g11.6/rac1l/elavl4/myo10l1/ntn1b/scn8aa/lrrc4cb/plppr4a/grna/bdnf/slitrk2/nkx6.1/ncam1b/cntn3a.1/nrxn2a/dagla/efna3a/dpysl5b/thsd7aa/si:ch211-241e1.3/plxna3/l1camb/plxna1a/nrcama/sema6ba/sema3b |
| GO:0001505 | regulation of neurotransmitter levels | -2.01367130950093 | 0.000141537068751352 | snap25a/napbb/slc5a7a/napba/chata/stxbp1b/cplx2a/comtb/si:ch211-81a5.8/dbh/syn1/stx3a/caskb/snap25b/mctp2b/snap47/syt7b/syn2a/rims1b/gad1b/syt4/rab3ab/moxd1/cadpsb/syt11a/unc13a/stxbp1a/syt1a/ppfia3/gad2/syt2a/baiap3/hrh3/rph3ab/th2/stx11a |
| GO:0061564 | axon development | -1.97936421112892 | 1.14653846153846e-08 | plxnb1b/pik3cd/mycbp2/rb1/ptprfb/zgc:100846/plp1b/ptprfa/isl2b/nrp2b/arhgap35a/rapgef1a/emb/pik3ca/erbb2/rbpms2a/aplp2/mapk8ip3/npr1a/sema3fa/olfm1a/unc5b/nptna/vegfc/dbn1/gpm6bb/si:ch211-129c21.1/phox2bb/ubap1/ctnna2/cntn4/cdh2/trarg1a/kalrnb/ulk2/ptk7a/dbnlb/nfia/efna1b/ephb3a/slc35b2/rbpms2b/fus/map1ab/dock7/foxb1a/nsfa/rab33ba/nrp2a/pnpla6/kctd13/isl1a/fezf2/gpm6ba/slit2/tenm3/rufy3/nfasca/sim1a/adcy8/pak1/spg11/ret/kif1b/arnt2/sema3bl/zdhhc17/plxna4/appa/arhgap35b/efna2a/dscama/apc/robo2/pcdh17/casp9/epha4a/hdc/anos1a/trioa/lhx9/epha7/rtn4r/cyfip2/dpysl3/draxinb/slit1b/robo3/usp33/map1sb/appb/dzip1/sema5a/kifbp/chad/il6st/robo1/tnr/tenm4/epha4l/kif5c/slitrk4/islr2/nptnb/lhx2b/myo9ab/kalrna/cdkl5/ephb2b/epha4b/ek1/clstn1/rac3a/brsk1b/aplp1/rab3ab/gap43/lrrc4ca/srrm4/lhx1a/map1aa/atl1/nkx2.2a/arhgef25b/dscamb/triob/tmem59l/myo10l3/nsmfb/slit3/dpysl5a/sarm1/chst11/fzd3a/kif5aa/robo4/efnb3b/vax1/stat3/zgc:77784/sema4c/si:ch211-113g11.6/rac1l/elavl4/myo10l1/ntn1b/scn8aa/lrrc4cb/plppr4a/grna/bdnf/slitrk2/nkx6.1/ncam1b/cntn3a.1/nrxn2a/dagla/efna3a/dpysl5b/thsd7aa/si:ch211-241e1.3/plxna3/l1camb/plxna1a/nrcama/sema6ba/sema3b |
| GO:0051049 | regulation of transport | -1.97152747166447 | 1.14653846153846e-08 | prnprs3/gem/kcne4/kcna1a/unc45a/dmxl2/mffa/scn5lab/mcoln1a/pxk/abcg1/kcnj13/rab33ba/fgf13b/cep290/c2cd5/cldnb/abca1a/hecw1b/cacna1ia/kctd7/nedd4l/cplx2a/chp2/kcnab1b/si:cabz01076231.1/cacnb1/bmp2k/cacna1fb/mcu/si:ch211-204c21.1/fgf13a/msnb/kcng1/cacng5a/cacna1ha/si:ch211-81a5.8/stim2b/cnstb/syt6b/stxbp5b/c2cd2l/caskb/fxyd6/crhbp/scn1ba/kcnab2a/pacsin1a/mctp2b/fgf12a/plcb3/rab3c/htr1aa/ank3a/dnajc5aa/hecw2a/ptprn2/rab3da/cacnb2b/syt7b/cacng7b/rab3b/rims1b/shisa9a/syt9b/zgc:91999/syt4/rab3ab/kcnj3b/kcnc2/kcnj19a/sgip1b/cadpsb/scn3b/rrad/syt13/stxbp5a/syt11a/cacng2b/kcnj1a.1/syt1a/cacng3b/shisa7b/syt9a/syt10/ptprna/kcnc1b/stx1b/cacna1da/stxbp4/anxa1c/scn8aa/cacng2a/rab44/stc1l/cacna1c/cacnb4b/fgf11b/cacna2d3/kcnj14/kcnj19b/syt2a/nkx6.1/kcnd3/il1rapl1a/syt3/cabp2a/baiap3/cacna1ab/kcnc1a/rph3ab/shisa9b/kcnd2/cnih2/npvf/kcnab2b/trh/cacng8a/cacna2d2b/scn1lab/kcnj5/syt6a/cacna2d4b/kcnh4a/kcnb1 |
| GO:0032879 | regulation of localization | -1.91598630922618 | 1.14653846153846e-08 | dtnbp1b/scn4ba/cplx2b/plcg1/gpr137ba/cacng8b/ptprnb/hip1rb/kcnh7/efcab7/scn8ab/aplnrb/adcyap1b/prnprs3/gem/kcne4/kcna1a/unc45a/letm1/dmxl2/mffa/scn5lab/mcoln1a/pxk/abcg1/kcnj13/rab33ba/fgf13b/cep290/c2cd5/cldnb/abca1a/hecw1b/cacna1ia/kctd7/nedd4l/cplx2a/chp2/plin2/kcnab1b/si:cabz01076231.1/cacnb1/bmp2k/cacna1fb/mcu/si:ch211-204c21.1/fgf13a/msnb/kcng1/plch1/cacng5a/cacna1ha/si:ch211-81a5.8/stim2b/cnstb/syt6b/stxbp5b/c2cd2l/caskb/fxyd6/crhbp/scn1ba/kcnab2a/pacsin1a/mctp2b/fgf12a/plcb3/rab3c/htr1aa/ank3a/dnajc5aa/hecw2a/wnt5b/ptprn2/rab3da/cacnb2b/plce1/plch2a/syt7b/cacng7b/rab3b/rims1b/shisa9a/syt9b/zgc:91999/syt4/gpc6a/rab3ab/kcnj3b/kcnc2/kcnj19a/sgip1b/zgc:77486/cadpsb/scn3b/rrad/syt13/stxbp5a/syt11a/cacng2b/kcnj1a.1/syt1a/gpc5b/cacng3b/shisa7b/syt9a/syt10/ptprna/kcnc1b/stx1b/cacna1da/stxbp4/anxa1c/scn8aa/cacng2a/rab44/stc1l/cacna1c/cacnb4b/fgf11b/cacna2d3/kcnj14/kcnj19b/syt2a/nkx6.1/kcnd3/il1rapl1a/syt3/cabp2a/baiap3/cacna1ab/kcnc1a/rph3ab/shisa9b/kcnd2/si:ch211-13f8.1/cnih2/npvf/kcnab2b/trh/cacng8a/cacna2d2b/scn1lab/kcnj5/syt6a/cacna2d4b/kcnh4a/kcnb1 |
| GO:1901701 | cellular response to oxygen-containing compound | -1.89692649413327 | 1.56824905636127e-05 | zgc:172120/pik3r3b/myd88/carf/stat6/nr4a2b/cpeb4a/chrng/glra1/syt6b/il13/c2cd2l/phip/htr1aa/ptger4c/ticam1/htr1d/ptprn2/tet3/gnao1b/inpp5ka/syt7b/syt9b/syt4/insig1/esr2b/ptger4a/cpne4a/vtcn1/cxcl8a/cxcl8b.3/nr4a2a/syt13/syt11a/stat3/nsg2/syt1a/sstr1a/syt9a/syt10/gnao1a/ptprna/cpne2/stxbp4/adgrv1/anxa1c/nr4a3/inpp5kb/syt2a/syt3/cpne7/irs2b/drd2a/htr1fa/chrm4a/gnb5a/cpne8/syt6a/il1b/gper1/cxcl18b |
| GO:0048640 | negative regulation of developmental growth | -1.88113815885922 | 0.0105693423015173 | sema6d/sema6e/dusp6/sema3fa/si:ch211-129c21.1/ulk2/slit2/sema3bl/robo2/epha4a/wwc1/sema5a/epha4b/slit3/sema4c/si:ch211-113g11.6/wwc3/sema6ba/sema3b |
| GO:0007019 | microtubule depolymerization | -1.8750187848337 | 0.021391144294225 | map1ab/stmn4/apc/map1b/stmn2a/map1sb/stmn2b/stmn3/map1aa/camsap1a/stmn4l/apc2 |
| GO:0032409 | regulation of transporter activity | -1.86261008963063 | 0.00378678429698509 | cabp1a/cabp5a/scn4ba/cacng8b/prnprs3/gem/kcne4/fgf13b/hecw1b/kctd7/nedd4l/chp2/kcnab1b/si:cabz01076231.1/cacnb1/fgf13a/kcng1/cacng5a/stim2b/fxyd6/fgf12a/ank3a/hecw2a/cacnb2b/cacng7b/shisa9a/rrad/cacng2b/cacng3b/shisa7b/cacng2a/cacnb4b/fgf11b/cabp2a/shisa9b/cnih2/cacng8a |
| GO:0003310 | pancreatic A cell differentiation | -1.86173412990953 | 0.00748525766259099 | isl1a/pax6b/insm1b/nkx6.2/myt1b/nkx6.1/arxa |
| GO:0036269 | swimming behavior | -1.85563104344825 | 0.0118323095206783 | dmrt3a/myo9aa/sin3b/panx1b/kif2a/gdpd5a/zgc:100846/ca10a/klhl41a/spg11/scdb/micall2b/ca10b/myo9ab/cdkl5/grna/tcap/cacna1ab/dagla/syngap1b/gnb5a/scn1lab/dnmt3aa/oxt |
| GO:0019226 | transmission of nerve impulse | -1.84484111835001 | 0.0190096470355517 | cacng8b/scn8ab/cacna1ia/cacng5a/cacna1ha/ank3a/cacng7b/cacng2b/cacng3b/scn8aa/cacng2a/kcnab2b/cacng8a/scn1lab |
| GO:0021545 | cranial nerve development | -1.80828563846919 | 0.00979204145463202 | rb1/adgrg1/klf6a/hgfa/mib1/erbb2/nes/tuba1a/nsfa/six3a/slit2/kif1b/chn1/robo2/klf7b/slit1b/six3b/kifbp/diras1a/lhx2b/vax1/mdga2a/cdh4/foxi1 |
| GO:0007186 | G protein-coupled receptor signaling pathway | -1.79358601966978 | 1.60681800279223e-06 | penka/gpr27/rgs18/trpm1b/gpr78a/cart2/opn1mw2/gnb1a/gpr182/npy/chrm2a/rgrb/penkb/gng3/glrbb/adcy2a/gal/adcy6b/adcy8/adgre15/ptgir/f2rl2/rgra/grm5a/gpr22a/gnal/lpar1/or95a1/gpr35.1/s1pr1/glra1/ccl38.6/gnai2b/tmem116/drd3/oprd1b/aplnr2/gpr22b/usp33/grm4/grm6b/npffl/lmbrd2b/rgs7a/ccl34b.4/grm8b/cart3/gpr174/htr1aa/ptger4c/tafa5a/gpr61l/adgrf3b/pde6ga/htr1d/pdyn/gnb2/si:ch211-117l17.6/gnao1b/adgra2/xcr1a.1/gnb3a/ccl44/gnb4b/agtr2/rgs9b/mrap2a/gpr63/ptger4a/adra2da/opn6b/prlhr2a/adcy6a/nmur3/adcy2b/adgrl3.1/adcy1a/si:dkey-6n21.13/grin1a/grm2a/tas1r2.1/sstr1a/gabbr1b/pth2/adra2b/trpm1a/git2b/tmtops2b/gnao1a/gpr85/gpr158b/adgrv1/ccl35.1/si:dkey-172j4.3/adcy5/grm8a/tshba/adgra1a/tafa5b/grm2b/prlhr2b/gpr75/vipr1b/adgrb3/pnocb/adgre5b.2/hrh3/adrb1/rgs11/drd2a/dagla/adgrb1a/htr1fa/npb/si:ch211-132p1.2/kctd8/chrm4a/ltb4r2a/crhr1/gpr158a/npvf/ltb4r/grm1a/adcyap1r1a/pomca/adgrg3/gnb5a/rxfp1/tac3b/htr7c/adgrl1a/adgrb1b/gper1/or92a7/p2ry13 |
| GO:0010721 | negative regulation of cell development | -1.7837780436552 | 0.00979204145463202 | sema3fa/gpr137ba/si:ch211-129c21.1/irf2bp2b/spi1b/ulk2/slit2/sema3bl/tmem178a/robo2/epha4a/bmp15/sema5a/epha4b/slit3/sema4c/si:ch211-113g11.6/anxa1c/mmp9/sema6ba/gper1/sema3b |
| GO:0006935 | chemotaxis | -1.78340911781747 | 1.1812068993636e-05 | unc5b/nptna/vegfc/tnfrsf1a/si:ch211-129c21.1/cntn4/cdh2/kalrnb/maco1a/tdgf1/ptk7a/efna1b/ephb3a/slc35b2/rbpms2b/rab33ba/nrp2a/fezf2/slit2/tenm3/maco1b/nfasca/sim1a/adcy8/kif1b/arnt2/sema3bl/myd88/plxna4/efna2a/dscama/apc/robo2/pcdh17/hdc/anos1a/trioa/s1pr1/lhx9/epha7/ccl38.6/rtn4r/cyfip2/dpysl3/draxinb/slit1b/robo3/usp33/aanat2/dock4b/dzip1/trim33/ccl34b.4/sema5a/chad/vegfd/dock4/robo1/tnr/tenm4/epha4l/kif5c/islr2/nptnb/lhx2b/kalrna/xcr1a.1/ccl44/ephb2b/rac3a/gap43/lhx1a/atl1/nkx2.2a/arhgef25b/dscamb/triob/slit3/cxcl8a/dpysl5a/cxcl8b.3/chst11/fzd3a/kif5aa/si:dkey-12h9.6/robo4/efnb3b/vax1/stat3/zgc:77784/sema4c/si:ch211-113g11.6/rac1l/anxa1c/ntn1b/scn8aa/ccl35.1/cxcl8b.1/ncam1b/cntn3a.1/nrxn2a/efna3a/dpysl5b/si:ch211-241e1.3/plxna3/l1camb/plxna1a/nrcama/sema6ba/il1b/sema3b/cxcl18b |
| GO:0042330 | taxis | -1.78340911781747 | 1.1812068993636e-05 | unc5b/nptna/vegfc/tnfrsf1a/si:ch211-129c21.1/cntn4/cdh2/kalrnb/maco1a/tdgf1/ptk7a/efna1b/ephb3a/slc35b2/rbpms2b/rab33ba/nrp2a/fezf2/slit2/tenm3/maco1b/nfasca/sim1a/adcy8/kif1b/arnt2/sema3bl/myd88/plxna4/efna2a/dscama/apc/robo2/pcdh17/hdc/anos1a/trioa/s1pr1/lhx9/epha7/ccl38.6/rtn4r/cyfip2/dpysl3/draxinb/slit1b/robo3/usp33/aanat2/dock4b/dzip1/trim33/ccl34b.4/sema5a/chad/vegfd/dock4/robo1/tnr/tenm4/epha4l/kif5c/islr2/nptnb/lhx2b/kalrna/xcr1a.1/ccl44/ephb2b/rac3a/gap43/lhx1a/atl1/nkx2.2a/arhgef25b/dscamb/triob/slit3/cxcl8a/dpysl5a/cxcl8b.3/chst11/fzd3a/kif5aa/si:dkey-12h9.6/robo4/efnb3b/vax1/stat3/zgc:77784/sema4c/si:ch211-113g11.6/rac1l/anxa1c/ntn1b/scn8aa/ccl35.1/cxcl8b.1/ncam1b/cntn3a.1/nrxn2a/efna3a/dpysl5b/si:ch211-241e1.3/plxna3/l1camb/plxna1a/nrcama/sema6ba/il1b/sema3b/cxcl18b |
| GO:0098953 | receptor diffusion trapping | -1.77120547755989 | 0.0221781634063151 | cacng8b/gphnb/cacng5a/gphna/cacng7b/cacng2b/cacng3b/cacng2a/cacng8a |
| GO:0098970 | postsynaptic neurotransmitter receptor diffusion trapping | -1.77120547755989 | 0.0221781634063151 | cacng8b/gphnb/cacng5a/gphna/cacng7b/cacng2b/cacng3b/cacng2a/cacng8a |
| GO:0099628 | neurotransmitter receptor diffusion trapping | -1.77120547755989 | 0.0221781634063151 | cacng8b/gphnb/cacng5a/gphna/cacng7b/cacng2b/cacng3b/cacng2a/cacng8a |
| GO:0071867 | response to monoamine | -1.76002500271274 | 0.0460556112254136 | htr1aa/htr1d/gnao1b/nsg2/gnao1a/drd2a/htr1fa/gnb5a |
| GO:0071869 | response to catecholamine | -1.76002500271274 | 0.0460556112254136 | htr1aa/htr1d/gnao1b/nsg2/gnao1a/drd2a/htr1fa/gnb5a |
| GO:0051960 | regulation of nervous system development | -1.75557639124564 | 0.00248666027456893 | nlk1/si:zfos-741a10.3/plxnb1b/hes6/neurod6a/adnpb/sema3fa/btbd6a/her12/dbn1/si:ch211-129c21.1/lef1/ulk2/dbnlb/ephb3a/dhrs12la/slit2/ctbp2a/sema3bl/plxna4/clstn2b/bhlhe40/robo2/epha4a/neurod4/gata3/her13/gbx2/tnfrsf21/sema5a/foxo3a/tnr/tenm4/slitrk4/caprin2/zbtb16a/cdkl5/epha4b/clstn1/nsmfb/slit3/clstn2a/elavl3/stat3/sema4c/si:ch211-113g11.6/gbx1/grna/bdnf/clstn3/slitrk2/dmbx1a/plxna3/l1camb/mmp9/plxna1a/sema6ba/sema3b |
| GO:0097120 | receptor localization to synapse | -1.75544036908873 | 0.0351674145187552 | dlg1b/cacng8b/grip2b/gphnb/cacng5a/stx3a/anks1b/kif5c/gphna/cacng7b/kif5aa/cacng2b/cacng3b/stx1b/cacng2a/rab44/cacng8a |
| GO:0031532 | actin cytoskeleton reorganization | -1.74088656656218 | 0.0402190977319739 | klf6a/anxa1a/baiap2a/sipa1l1/mink1/nedd9/thsd7bb/carmil2/parvg/carmil3/plek/thsd7ba/anxa1c/plek2/phactr1/thsd7aa |
| GO:0040013 | negative regulation of locomotion | -1.73929262815316 | 0.018759055936431 | sema3fa/lrch1/si:ch211-129c21.1/lbx2/prox1a/miip/lef1/phf8/srgap3/sema3bl/robo2/anos1a/rap2b/sema5a/srgap1b/sinhcaf/arhgap4b/sema4c/si:ch211-113g11.6/rtn4rl2a/sema6ba/sema3b |
| GO:0031110 | regulation of microtubule polymerization or depolymerization | -1.72739295421889 | 0.0437181181328161 | ska3/cdk5rap2/stmn1b/map1sa/camsap2a/camsap1b/slain2/map1ab/traf3ip1/zgc:66447/clasp2/mapre3b/stmn4/apc/map1b/stmn2a/map1sb/stmn2b/stmn3/clip1a/map1aa/camsap1a/stmn4l/apc2 |
| GO:0040011 | locomotion | -1.71881331498307 | 4.74228301214316e-06 | plcg1/unc5b/nptna/lrch1/vegfc/tnfrsf1a/si:ch211-129c21.1/lbx2/cntn4/cdh2/prox1a/kalrnb/miip/maco1a/lef1/tdgf1/foxj1a/ptk7a/efna1b/ephb3a/slc35b2/gnb1a/rbpms2b/sst7/slkb/rab33ba/nrp2a/fezf2/phf8/slit2/tenm3/maco1b/srgap3/nfasca/sim1a/adcy8/kif1b/arnt2/sema3bl/myd88/plxna4/efna2a/dscama/apc/robo2/pcdh17/hdc/asap1a/arhgef39/anos1a/trioa/s1pr1/lhx9/epha7/hace1/peak1/ccl38.6/rtn4r/cyfip2/dpysl3/draxinb/gata3/slit1b/robo3/usp33/tek/aanat2/dock4b/dzip1/rap2b/trim33/ccl34b.4/sema5a/chad/vegfd/pdgfbb/dock4/robo1/tnr/tenm4/epha4l/kif5c/islr2/nptnb/htr1d/lhx2b/kalrna/xcr1a.1/ccl44/ephb2b/sst1.1/srgap1b/rac3a/gap43/lhx1a/atl1/nkx2.2a/arhgef25b/dscamb/amot/triob/nsmfb/slit3/cxcl8a/dpysl5a/cxcl8b.3/chst11/fzd3a/kif5aa/si:dkey-12h9.6/sinhcaf/robo4/efnb3b/vax1/arhgap4b/stat3/dock10/zgc:77784/sema4c/adam8a/si:ch211-113g11.6/rac1l/anxa1c/ntn1b/scn8aa/ccl35.1/cxcl8b.1/mdga2a/zgc:153631/asap1b/ncam1b/cntn3a.1/nrxn2a/klhl40b/efna3a/chrm4a/dpysl5b/si:ch211-241e1.3/plxna3/l1camb/plxna1a/nrcama/rtn4rl2a/sema6ba/il1b/sema3b/cxcl18b |
| GO:0006959 | humoral immune response | -1.69323952530372 | 0.0594675488145613 | c4/tnfrsf21/c1qa/tfe3a/cfb/cxcl8b.3/c6/susd4/c7b/c4b/c3a.2/itln2/cxcl18b |
| GO:0007420 | brain development | -1.67227058783588 | 2.31796694375566e-05 | her12/wnt3/macroh2a2/marcksa/kctd12.1/daam1a/pknox1.1/tgif1/ctnna2/cntn4/cdh2/adcyap1b/prox1a/zfhx3b/prnprs3/sbno2a/lef1/tdgf1/ulk2/tsc22d3/bag6l/pou3f3a/atg4da/creb1a/pafah1b1a/birc6-as2/sox14/foxb1a/nsfa/zfhx3a/kdm6al/slc2a3a/cep290/kctd13/fezf2/phf8/six3a/nfasca/sim1a/pax6b/arnt2/raraa/grhl3/tbr1b/kirrel3l/cars2/ptprsa/kdm7ab/nr4a2b/ralgapa1/apc/epha4a/hdc/anos1a/lhx9/neurod4/pcdh10b/pbx1a/mvp/six3b/mdkb/si:ch73-119p20.1/gbx2/dzip1/lgi1a/pbx1b/epb41l5/auts2a/pbx3b/pcare1/hecw2a/znf503/meis2a/wnt5b/lhx6a/irx7/lhx2b/brf1b/rbfox1/tcf7l2/nkx2.1/pcdh19/srrm4/lhx1a/asphd1/pou4f2/prmt8b/tbxta/aatkb/rtn1b/sp5l/nr2e3/tbcd/has2/lhx5/fzd3a/pou3f2b/nr4a2a/efnb3b/ndrg4/vax1/pax2a/git2b/gbx1/ntn1b/pou3f2a/bdnf/atp1a3a/sez6l2/mdga2a/rx3/dmbx1a/otpb/six6a/cdk5r2a/arxa/nkx2.4a/syngap1b/gnrh2/l1camb/cnnm2a/otpa/kcnb1/nrcama/gper1/bicdl1 |
| GO:0060322 | head development | -1.66225293057786 | 4.42238345198842e-05 | her12/wnt3/macroh2a2/marcksa/kctd12.1/daam1a/pknox1.1/tgif1/ctnna2/cldn7a/cntn4/cdh2/adcyap1b/prox1a/zfhx3b/prnprs3/sbno2a/lef1/tdgf1/ulk2/tsc22d3/bag6l/pou3f3a/atg4da/creb1a/pafah1b1a/birc6-as2/sox14/foxb1a/nsfa/zfhx3a/kdm6al/slc2a3a/cep290/kctd13/fezf2/phf8/six3a/nfasca/sim1a/pax6b/arnt2/raraa/grhl3/tbr1b/kirrel3l/cars2/ptprsa/kdm7ab/nr4a2b/ralgapa1/apc/epha4a/hdc/wdr11/anos1a/lhx9/neurod4/pcdh10b/pbx1a/mvp/six3b/mdkb/si:ch73-119p20.1/gbx2/dzip1/lgi1a/pbx1b/epb41l5/auts2a/pbx3b/pcare1/hecw2a/znf503/meis2a/wnt5b/lhx6a/irx7/lhx2b/brf1b/rbfox1/tcf7l2/nkx2.1/pcdh19/srrm4/lhx1a/asphd1/pou4f2/prmt8b/tbxta/aatkb/rtn1b/sp5l/nr2e3/tbcd/has2/lhx5/fzd3a/pou3f2b/nr4a2a/efnb3b/ndrg4/vax1/pax2a/git2b/gbx1/ntn1b/chd5/pou3f2a/bdnf/atp1a3a/sez6l2/mdga2a/specc1/rx3/dmbx1a/otpb/six6a/cdk5r2a/arxa/nkx2.4a/syngap1b/gnrh2/l1camb/cnnm2a/otpa/kcnb1/nrcama/gper1/bicdl1 |
| GO:0045732 | positive regulation of protein catabolic process | -1.66175262841073 | 0.0607582432020156 | oaz1a/csnk1db/tmem259/oaz2b/socs5b/prkn/apc/rnf217/si:ch211-278j3.3/trib1/fzr1b/rnf144ab/csnk1e/adam8a/klhl40b/apc2/rnf144aa |
| GO:1901797 | negative regulation of signal transduction by p53 class mediator | -1.65413798029984 | 0.0829144295829561 | cd74b/cd74a/bcor/vax1/bcl6aa |
| GO:0006688 | glycosphingolipid biosynthetic process | -1.64952029857802 | 0.097836707744085 | b4galt6/b4galt3/st6galnac3 |
| GO:0031644 | regulation of nervous system process | -1.64521543139582 | 0.0906720571933235 | dlgap1a/tnfrsf21/tenm4/dlgap4b/si:dkey-174m14.3/kcnab2b/dlgap3/neto1l |
| GO:0060560 | developmental growth involved in morphogenesis | -1.64279725335724 | 0.0128583805736124 | plxnb1b/tmem108/pik3cd/mycbp2/spag6/cbsb/smad2/pik3ca/zeb2a/mapk8ip3/sema3fa/dbn1/si:ch211-129c21.1/ubap1/cdh2/trarg1a/tmem67/ulk2/ptk7a/dbnlb/creb1a/fus/isl1a/slit2/ppp1cb/spg11/ret/kif1b/sema3bl/bbs1/robo2/pcdh17/casp9/epha4a/robo3/appb/sema5a/wnt5b/myo9ab/cdkl5/epha4b/clstn1/tbxta/zeb2b/tmem59l/myo10l3/nsmfb/slit3/stat3/sema4c/si:ch211-113g11.6/myo10l1/ntn1b/grna/bdnf/nrn1la/nkx6.1/nrxn2a/thsd7aa/plxna3/l1camb/sema6ba/sema3b |
| GO:0016049 | cell growth | -1.63666909504427 | 0.0141269580867656 | plxnb1b/tmem108/pik3cd/mycbp2/spag6/ewsr1a/pik3ca/mapk8ip3/sema3fa/dbn1/si:ch211-129c21.1/ubap1/trarg1a/ulk2/dbnlb/supv3l1/dnph1/fus/isl1a/slit2/spg11/ret/kif1b/sema3bl/robo2/pcdh17/casp9/epha4a/robo3/appb/sema5a/wnt5b/myo9ab/cdkl5/epha4b/clstn1/tmem59l/myo10l3/nsmfb/slit3/sema4c/si:ch211-113g11.6/myo10l1/ntn1b/epb41l3a/grna/bdnf/nrn1la/nkx6.1/nrxn2a/thsd7aa/plxna3/l1camb/sema6ba/sema3b |
| GO:0034643 | establishment of mitochondrion localization. microtubule-mediated | -1.63631487834659 | 0.0931198545493737 | rhot1b/trak1a/sybu/kif5aa/rhot2 |
| GO:0047497 | mitochondrion transport along microtubule | -1.63631487834659 | 0.0931198545493737 | rhot1b/trak1a/sybu/kif5aa/rhot2 |
| GO:0051654 | establishment of mitochondrion localization | -1.63631487834659 | 0.0931198545493737 | rhot1b/trak1a/sybu/kif5aa/rhot2 |
| GO:0032102 | negative regulation of response to external stimulus | -1.63570938930065 | 0.0540332674423811 | sema3bl/gramd4b/robo2/epha4a/il13/nr1h5/sema5a/cart3/ptger4c/irak3/epha4b/gramd4a/ptger4a/sema4c/si:ch211-113g11.6/susd4/grna/serpine1/sc:d189/sema6ba/sema3b |
| GO:0048885 | neuromast deposition | -1.63446228315307 | 0.0960340406478379 | lbx2/prox1a/lef1/phf8/anos1a/rtn4rl2a |
| GO:0071407 | cellular response to organic cyclic compound | -1.62675574144756 | 0.0543657683428525 | abhd2b/htr1aa/htr1d/gnao1b/insig1/esr2b/cxcl8a/nsg2/sstr1a/gnao1a/anxa1c/drd2a/htr1fa/gnb5a/gper1 |
| GO:0022603 | regulation of anatomical structure morphogenesis | -1.62242060545465 | 0.00196692398869123 | plcd4a/vegfab/stab1/hif1ab/amotl2b/sema6d/ptenb/alcama/sema6e/ptk2ab/thbs2b/brwd1/rspo3/tspan18a/gata2a/zeb1b/plxnb1b/pik3cd/disc1/mia3/anxa1a/fn1b/arhgap35a/pik3ca/flot2a/sema3fa/camsap1b/plcg1/unc5b/vegfc/dbn1/aplnrb/kctd12.1/si:ch211-129c21.1/rhogd/zeb1a/lbx2/cdh2/fmnl2b/ulk2/elmo1/marchf5/ptk7a/dbnlb/dab2ipb/lurap1/ephb3a/mffa/sipa1l1/tnika/vat1/palm1b/slit2/mief2/rufy3/hecw1b/pak1/sema3bl/plxna4/nedd4l/arhgap35b/robo2/casp9/epha4a/s1pr1/msnb/tanc2a/phip/sema5a/vegfd/tenm4/hecw2a/wnt5b/caprin2/cdkl5/epha4b/myo10/clstn1/rac3a/lrrc4ca/lhx1a/nsmfb/slit3/basp1/stat3/prkd1/sema4c/si:ch211-113g11.6/rac1l/gbx1/anxa1c/lrrc4cb/hif1aa/bdnf/serpine1/adgrb3/scg2b/gdf3/adgrb1a/thbs1a/dab2ipa/plxna3/l1camb/eve1/plxna1a/adgrb1b/sema6ba/sema3b |
| GO:0000381 | regulation of alternative mRNA splicing. via spliceosome | -1.62121505428978 | 0.0619371464051369 | hnrnpa0a/rbm38/rbm25a/celf2/rbfox3a/rbfox3b/celf6/rbfox1l/rbfox1/srrm4/nova2/celf3a/khdrbs2/celf5a/celf4 |
| GO:0001933 | negative regulation of protein phosphorylation | -1.61410605566215 | 0.0735990755145654 | spry4/aida/dtnbp1b/ptprc/ldlrad4a/camk2n2/cbl/camk2n1a/cblc/prkn/sh3bp5la/sh3bp5b/si:dkey-175m17.7/ibtk/camk2n1/irak3/inpp5ka/cadm4/inpp5kb/smcr8b |
| GO:0008361 | regulation of cell size | -1.59323427771673 | 0.0658674351521172 | arhgap35a/pik3ca/sema3fa/arhgap5/dbn1/si:ch211-129c21.1/ulk2/dbnlb/zgc:153039/slit2/sema3bl/arhgap35b/robo2/casp9/epha4a/sema5a/piezo1/cdkl5/epha4b/clstn1/nsmfb/slit3/si:ch211-178n15.1/stat3/slc12a5b/sema4c/si:ch211-113g11.6/ntn1b/slc12a5a/bdnf/nkx6.1/plxna3/l1camb/sema6ba/sema3b |
| GO:0043087 | regulation of GTPase activity | -1.57086875382947 | 0.0297635976408028 | tbc1d9/rapgef1a/flcn/sipa1l2/rasgef1ba/iqsec1b/ralgapa2/rasa4/gpr137ba/rgl1/nf1a/usp6nl/adcyap1b/vav2/agap1/rapgef5b/rapgef2a/si:dkey-91m11.5/dab2ipb/rgl3a/ephb3a/rapgef2b/dock7/rapgef4b/sipa1l1/gapvd1/rasal2/plxna4/chn1/ralgapa1/ccpg1/asap1a/rasgef1bb/ccl38.6/bcr/rgs7a/ccl34b.4/si:ch211-288d18.1/evi5l/tbc1d30/elmod1/vav3b/myo9ab/ccl44/ek1/rgs9b/sgsm3/syngap1a/arap3/rap1gapa/unm_sa1614/ralgps1/dock10/agap2/ccl35.1/asap1b/rasgrf2b/rgs11/syngap1b/dab2ipa/plxna3/rasgrf1/plxna1a/arhgap25 |
| GO:0002252 | immune effector process | -1.52782636998837 | 0.0607582432020156 | cd74b/c4/c1qa/ticam1/cd74a/cfb/cadm1b/cxcl8a/c6/si:ch211-196f2.6/susd4/anxa1c/zgc:153631/c7b/c4b/zgc:123107/c3a.2/csf2rb/mmp9/mhc2d4b |
| GO:0001755 | neural crest cell migration | -1.51280994298295 | 0.0960340406478379 | hif1ab/sema6d/pax6a/med13b/sema6e/lrp5/disc1/nrp2b/erbb2/pdgfrb/sema3fa/radil/si:ch211-129c21.1/cdh2/lef1/dmxl2/pax3a/nrp2a/fscn1a/pak1/ret/pax6b/sema3bl/mmp17b/wdr11/sema5a/zeb2b/sh3pxd2aa/sema4c/si:ch211-113g11.6/rx3/foxi1/sema6ba/sema3b |
| GO:0051240 | positive regulation of multicellular organismal process | -1.44230693231483 | 0.0657742225763037 | gata2a/slc27a1b/nlk1/si:zfos-741a10.3/plxnb1b/anxa1a/avpr1ab/neurod6a/tnfsf13b/nr2c2/cactin/apln/tlr5b/olfm1a/vegfc/dbn1/aplnrb/lef1/elmo1/dbnlb/lurap1/ephb3a/npy/tapt1b/dhrs12la/loxl2a/zgc:172120/plxna4/clstn2b/cd74b/neurod4/gata3/aanat2/vegfd/enpp5/tenm4/ticam1/slitrk4/caprin2/zbtb16a/cdkl5/clstn1/cd74a/lhx1a/vtcn1/nsmfb/clstn2a/basp1/stat3/si:ch211-196f2.6/anxa1c/grna/bdnf/zbtb16b/clstn3/slitrk2/zgc:153631/adrb1/npvf/plxna3/l1camb/zgc:123107/plxna1a/panx2/gper1/mhc2d4b |
| GO:0030334 | regulation of cell migration | -1.43156853115565 | 0.0696903189146766 | plcg1/lrch1/vegfc/si:ch211-129c21.1/lbx2/prox1a/miip/lef1/gnb1a/sst7/slkb/phf8/srgap3/sema3bl/plxna4/asap1a/arhgef39/anos1a/s1pr1/hace1/peak1/dpysl3/gata3/tek/rap2b/sema5a/vegfd/pdgfbb/sst1.1/srgap1b/amot/nsmfb/sinhcaf/robo4/arhgap4b/dock10/sema4c/adam8a/si:ch211-113g11.6/anxa1c/mdga2a/zgc:153631/asap1b/si:ch211-241e1.3/plxna3/plxna1a/rtn4rl2a/sema6ba/sema3b |
| GO:0009719 | response to endogenous stimulus | -1.32546647522578 | 0.0666420813824336 | tbx16/irs1/skia/igfbp3/pik3r3a/gsk3ba/rbm14b/gna11b/slc39a14/slc27a1b/esr2a/zyx/smad7/tll1/ltbp3/tacr1a/spry4/dusp6/trim71/anxa1a/avpr1ab/nrp2b/nr1d1/smad2/grb10a/flcn/smurf1/zeb2a/asic4b/fcho1/skib/rgmb/fgfbp1a/irs2a/drd4-rs/nptna/vegfc/fstl4/si:rp71-46j2.7/ldb2a/timp2a/tgif1/ldlrad4a/caly/fstl3/castor2/ppp4cb/zdhhc13/tsc22d3/il13ra1/dab2ipb/lifrb/golm1/rerea/chrm2a/slc2a3a/nrp2a/pnpla6/asip1/tcf21/grem2b/pik3r3b/raraa/thrb/zdhhc17/bmpr1ab/smad3b/stat6/gpr22a/nr4a2b/cpeb4a/chrng/glra1/sall1a/gata3/bmp15/dzip1/fabp4b/phip/crhbp/sp9/snx25/inhbab/vegfd/fgf12a/abhd2b/htr1aa/bmpr2a/scube1/ptger4c/ccbe1/stat1b/nrros/htr1d/lemd3/gnao1b/inpp5ka/smad6b/insig1/esr2b/ptger4a/atl1/tbxta/inhbaa/asic2/timp4.3/has2/adgrl3.1/kcp/grem1a/nr4a2a/bmpr2b/stat3/nsg2/wbp2/gnao1a/stxbp4/anxa1c/nr4a3/bdnf/elapor2b/inpp5kb/ntrk3a/frs3/irs2b/gdf3/drd2a/ntrk3b/htr1fa/chrm4a/trh/dio2/dab2ipa/gnb5a/rxfp1/c3a.2/sostdc1b/mmp9/timp2b/hpxa |
| ID | Description | NES | p.adjust | core_enrichment |
| GO:0042254 | ribosome biogenesis | 2.68255980674827 | 1.14653846153846e-08 | imp3/rpl26/mrpl10/mrm3a/nle1/noc2l/rbis/urb2/exosc2/polr1c/rpp40/exosc6/abt1/mphosph6/snu13b/emg1/wdr18/znhit3/pwp2h/utp4/utp23/mrm3b/nop14/heatr1/mrto4/pin4/sbds/lsm6/ddx31/wdr12/nom1/nifk/utp11/tfb1m/ddx27/mrpl22/ngdn/pih1d1/wdr3/ddx51/heatr3/gnl3/mrpl20/rrp36/gar1/nvl/riok2/slx9/bxdc2/pdcd11/trmt112/nop58/dimt1l/bud23/rexo4/nip7/mettl16/mphosph10/wdr36/dkc1/npm1a/nsun5/rpusd2/ppan/mdn1/ddx49/rpl7l1/ddx52/bop1/abce1/nol6/sdad1/tbl3/nop10/ddx10/mrpl36/pak1ip1/rpl35/rbfa/zcchc4/rrp9/rps7/si:dkey-251i10.3/utp15/snu13a/rrp7a/nop56/nhp2/wdr75/ddx18/c1d/rcl1/wdr55/rrp8/tsr2/eif6/nat10/utp25/nop2/nop16/polr1d/mterf3/ebna1bp2/rps21/ftsj3/bysl/utp6/rrs1/gnl3l/rps8a/mpv17l2/rps14/nol10/mtrex/rpl5a/riok1/esf1/rpl35a/pop5/ran/rpf1/mrps11/gnl2/rpl14/rpf2/tfb2m/fbl/utp3/dhx37/rrp1/rps16/ddx54/gtpbp10/rpl7/tsr1/ddx28/znf622/malsu1/rps9/mak16/c1qbp/lsg1/naf1/kri1/rpl23a/urb1/exosc8/fcf1/pes/pop4/rps15/rpsa/mtg2/rrp15/rpl7a/mterf4/exosc7/dcaf13/rps5/rplp0/exosc9/tsr3 |
| GO:0006364 | rRNA processing | 2.64217411952833 | 1.14653846153846e-08 | imp3/mrm3a/exosc2/rpp40/exosc6/abt1/mphosph6/snu13b/emg1/wdr18/znhit3/pwp2h/utp4/utp23/mrm3b/heatr1/mrto4/pin4/lsm6/wdr12/nifk/utp11/tfb1m/ddx27/ngdn/pih1d1/wdr3/ddx51/gnl3/rrp36/gar1/riok2/slx9/pdcd11/trmt112/dimt1l/bud23/rexo4/mettl16/mphosph10/wdr36/dkc1/nsun5/rpusd2/ddx49/rpl7l1/ddx52/bop1/nol6/tbl3/nop10/ddx10/rpl35/rbfa/zcchc4/rrp9/rps7/si:dkey-251i10.3/utp15/snu13a/rrp7a/nhp2/wdr75/ddx18/c1d/rcl1/wdr55/rrp8/tsr2/eif6/nat10/utp25/nop2/ebna1bp2/rps21/ftsj3/bysl/utp6/rrs1/gnl3l/rps8a/rps14/nol10/mtrex/riok1/esf1/pop5/rpf1/mrps11/gnl2/rpf2/tfb2m/fbl/utp3/dhx37/rrp1/rps16/ddx54/rpl7/tsr1/mak16/naf1/kri1/urb1/exosc8/fcf1/pes/pop4 |
| GO:0034470 | ncRNA processing | 2.61312605149024 | 1.14653846153846e-08 | ints6/prkra/imp3/thumpd2/pus3/wdr4/qtrt2/trmu/mrm3a/mtfmt/trmt10c/exosc2/rpp21/rnf113a/rpp40/exosc6/abt1/mphosph6/snu13b/rpp30/emg1/henmt1/wdr18/tprkb/piwil2/znhit3/elp3/zgc:136564/pwp2h/elp5/utp4/trmt61a/spout1/zc3h7bb/tp53rk/utp23/mrm3b/trmt44/heatr1/mrto4/pin4/trmt6/exosc5/tyw3/rpusd4/adat1/tsen2/trmt12/lsm6/wdr12/nsun2/nifk/utp11/thumpd3/tfb1m/osgep/ddx27/urm1/ctu2/mettl3/ngdn/pih1d1/wdr3/srrt/ddx51/gnl3/rrp36/gar1/riok2/ints6l/slx9/pdcd11/trmt112/dimt1l/bud23/osgepl1/rexo4/ptcd1/mettl16/mphosph10/wdr36/dkc1/mettl2a/thada/pus1/nsun5/gtpbp3/rpusd2/ddx49/rpl7l1/ddx52/bop1/trmt10b/tdrd9/nol6/clp1/ctu1/tsen15/rtcb/tbl3/ints9/nop10/ddx10/rpl35/rbfa/zcchc4/rrp9/rpph1/rps7/si:dkey-251i10.3/utp15/snu13a/rrp7a/elp6/nhp2/wdr75/ddx18/c1d/rcl1/wdr55/rrp8/tsr2/trmt61b/elac1/trpt1/eif6/nat10/utp25/nop2/ebna1bp2/tut1/rps21/elp2/ftsj3/bysl/mettl1/pus7/utp6/rrs1/dtwd1/gnl3l/rps8a/rps14/nol10/rpp25l/thg1l/mtrex/riok1/sepsecs/esf1/eif1axb/pop5/cdk5rap1/zbtb8os/rpf1/mrps11/gnl2/rpf2/tfb2m/trmt2b/fbl/utp3/rpp38/dhx37/rtraf/rrp1/wdr6/ints11/rps16/ddx54/aars1/rpl7/tsr1/mak16/trmt5/trmt9b/naf1/kri1/urb1/snip1/exosc8/fcf1/pes/pop4 |
| GO:0016072 | rRNA metabolic process | 2.5944143069453 | 1.14653846153846e-08 | imp3/mrm3a/exosc2/rpp40/exosc6/abt1/mphosph6/snu13b/emg1/wdr18/znhit3/pwp2h/utp4/utp23/mrm3b/heatr1/mrto4/pin4/exosc5/lsm6/wdr12/nifk/utp11/tfb1m/ddx27/ngdn/pih1d1/wdr3/ddx51/gnl3/rrp36/gar1/riok2/slx9/pdcd11/trmt112/dimt1l/bud23/rexo4/mettl16/mphosph10/wdr36/dkc1/polr1e/nsun5/rpusd2/ddx49/rpl7l1/ddx52/bop1/nol6/tbl3/nop10/ddx10/rpl35/rbfa/zcchc4/rrp9/rps7/si:dkey-251i10.3/utp15/snu13a/rrp7a/nhp2/wdr75/ddx18/c1d/rcl1/wdr55/rrp8/tsr2/eif6/nat10/utp25/nop2/taf1b/ebna1bp2/rps21/ftsj3/bysl/dis3/utp6/rrs1/gnl3l/rps8a/rps14/nol10/mtrex/riok1/esf1/pop5/rpf1/mrps11/gnl2/rpf2/tfb2m/fbl/utp3/polr1a/dhx37/rrp1/rps16/ddx54/pelo/rpl7/tsr1/mak16/naf1/kri1/urb1/exosc8/fcf1/pes/pop4 |
| GO:0034660 | ncRNA metabolic process | 2.54552940229038 | 1.14653846153846e-08 | ints6/prkra/fars2/imp3/thumpd2/pus3/farsa/wdr4/qtrt2/trmu/mrm3a/dars2/wars2/mtfmt/trmt10c/exosc2/rpp21/rnf113a/rpp40/exosc6/dars1/abt1/mphosph6/snu13b/rpp30/emg1/henmt1/wdr18/tprkb/piwil2/znhit3/elp3/zgc:136564/pwp2h/elp5/utp4/trmt61a/spout1/zc3h7bb/tp53rk/utp23/mrm3b/trmt44/heatr1/mrto4/pin4/trmt6/exosc5/tyw3/lars1b/rpusd4/adat1/tsen2/trmt12/lsm6/wdr12/nsun2/nifk/utp11/thumpd3/si:ch211-256m1.8/tfb1m/mars2/osgep/ddx27/urm1/ctu2/mettl3/ngdn/pih1d1/wdr3/srrt/ddx51/snapc3/gnl3/rrp36/gar1/riok2/ints6l/slx9/pdcd11/trmt112/qrsl1/dimt1l/bud23/osgepl1/rexo4/ptcd1/mettl16/gatb/mphosph10/wdr36/dkc1/mettl2a/thada/polr1e/pus1/nsun5/gtpbp3/rpusd2/iars1/ddx49/hars/rpl7l1/ddx52/bop1/trmt10b/tdrd9/qars1/nol6/nars2/vars1/clp1/ctu1/tsen15/rtcb/tbl3/ints9/nop10/ddx10/rpl35/rbfa/zcchc4/rrp9/rpph1/rps7/si:dkey-251i10.3/utp15/snu13a/rrp7a/elp6/nhp2/wdr75/ddx18/yars1/c1d/rcl1/wdr55/rrp8/tsr2/trmt61b/elac1/rpap2/trpt1/rars1/eif6/nat10/utp25/nop2/taf1b/ebna1bp2/tut1/rps21/elp2/tars1/ftsj3/bysl/gars1/mettl1/pars2/dis3/pus7/iars2/utp6/rrs1/dtwd1/gnl3l/rps8a/rps14/nol10/rpp25l/thg1l/mtrex/riok1/sars1/sepsecs/esf1/eif1axb/pop5/vars2/mepceb/cdk5rap1/zbtb8os/rpf1/mrps11/gnl2/sars2/rpf2/tfb2m/trmt2b/fbl/utp3/polr1a/rpp38/dhx37/rtraf/rrp1/wdr6/ints11/rps16/ddx54/aars1/pelo/rpl7/tsr1/mak16/trmt5/trmt9b/naf1/kri1/dtd2/urb1/snip1/exosc8/fcf1/pes/pop4 |
| GO:0022613 | ribonucleoprotein complex biogenesis | 2.40988428678125 | 1.14653846153846e-08 | imp3/rpl26/mrpl10/mrm3a/nle1/noc2l/rbis/gemin5/urb2/exosc2/polr1c/rpp40/exosc6/abt1/mphosph6/snu13b/emg1/wdr18/znhit3/pwp2h/trnau1apa/utp4/utp23/mrm3b/nop14/heatr1/mrto4/pin4/sbds/snrpd2/lsm6/ddx31/wdr12/nom1/nifk/utp11/tfb1m/prmt7/ddx27/mrpl22/ngdn/pih1d1/gemin2/wdr3/ddx51/heatr3/gnl3/mrpl20/rrp36/gar1/nvl/snrpg/riok2/slx9/bxdc2/pdcd11/trmt112/nop58/dimt1l/bud23/prpf18/mcts1/rexo4/nip7/mettl16/mphosph10/wdr36/dkc1/npm1a/nsun5/rpusd2/ppan/mdn1/ddx49/rpl7l1/ddx52/bop1/abce1/nol6/clp1/sdad1/tbl3/nop10/ddx10/mrpl36/pak1ip1/rpl35/rbfa/zcchc4/rrp9/ptges3b/rps7/si:dkey-251i10.3/utp15/snu13a/rrp7a/nop56/eif3m/nhp2/wdr75/ddx18/c1d/nopchap1/snrpd3l/eif3ea/rcl1/wdr55/rrp8/tsr2/coil/eif6/snrpf/nat10/utp25/nop2/nop16/usp39/polr1d/mterf3/ebna1bp2/rps21/eif3jb/ftsj3/snrpe/bysl/utp6/rrs1/gnl3l/rps8a/mpv17l2/rpl38/rps14/nol10/shq1/mtrex/rpl5a/riok1/esf1/rpl35a/pop5/eif5/ran/eif3ha/rpf1/trnau1apb/mrps11/gnl2/scaf11/rpl14/rpf2/tfb2m/fbl/utp3/dhx37/rrp1/eif3s10/cdc73/rps16/ddx54/denr/gtpbp10/rpl7/tsr1/ddx28/znf622/malsu1/rps9/mak16/c1qbp/lsg1/naf1/kri1/rpl23a/urb1/exosc8/eif3s6ip/fcf1/pes/pop4/rps15/rpsa/eif2s2/mtg2/polr2d/rrp15/rpl7a/srpk1a/mterf4/exosc7/dcaf13/rps5/gemin8/rplp0/exosc9/tsr3/snrpd1/rps27.1/aar2/rpl10/rpl5b/eif3g/gtpbp4/eif3d/rpl6/yju2/noc4l |
| GO:0140053 | mitochondrial gene expression | 2.16306020123481 | 0.000153043817326382 | fastkd3/tefm/trmu/fastkd2/tbrg4/wars2/trmt10c/mrpl58/fastkd1/coa3a/mtg1/mrps18b/tfb1m/uqcc2/chchd1/tsfm/mtrf1l/qrsl1/gatb/gtpbp3/mtif2/hars/mrps24/rcc1l/tfam/trmt61b/mrpl47/mterf3/gars1/iars2/mrpl23/sars2/tfb2m/ptcd3/fastk/mrpl16 |
| GO:0043043 | peptide biosynthetic process | 2.02998011519709 | 1.14653846153846e-08 | dazl/fars2/farsa/zc3h12ab/dph7/rpl26/mrpl10/dars2/wars2/mtfmt/trmt10c/gemin5/exosc2/mrpl58/mrps5/dars1/rpp30/zar1/mrpl9/dcp1b/piwil2/coa3a/trnau1apa/mrpl35/mif4gdb/mrpl30/eif2ak1/dcps/mrpl17/dph5/lars1b/mtg1/mrps17/mrps18b/mrpl4/si:ch211-256m1.8/zgc:171480/mars2/uqcc2/chchd1/tsfm/gss/eif4ea/mrpl22/mettl3/ngdn/mtrf1l/faub/mrps18c/drg2/lsm1/mrpl27/mrpl20/dph6/magoh/eif1ad/eif2a/eefsec/qrsl1/mrpl51/mrps23/mcts1/ago3b/mettl16/gatb/mrpl18/mrps21/gtpbp3/mtif2/iars1/mrpl14/rpl12/hars/mrpl24/pdf/eif2b1/eif4e2rs1/abce1/eif2s1a/qars1/nars2/mrpl41/impact/vars1/mrpl34/fxr2/larp6a/mrps24/mrpl36/rpl35/cnot8/mrps12/mrpl13/rcc1l/eif5a2/rpl32/mrps2/rps7/mrpl12/brf1a/eif2b2/eif3m/eif4ebp3l/yars1/bdh2/eef1b2/eif3ea/mrpl11/rpl22l1/mrpl33/rpl34/rars1/eif6/dph2/ogfod1/gspt1/pde12/mrpl47/rps21/rps17/eef1a1l3/eif3jb/tars1/cnot6l/eef1da/eif4ebp1/gars1/mrps10/pars2/iars2/rc3h2/mrpl23/rpl29/zgc:171772/rpl37/rps8a/gatc/rpl38/rps3a/rps14/rps15a/srp9/larp4b/mrps30/rps20/rps29/tcea3/rpl5a/rpl15/sars1/sepsecs/mrpl3/eif1axb/rpl35a/vars2/eif5/rps18/eif3ha/zc3h15/mrpl32/mrpl37/trnau1apb/mrps11/rpl11/rpl14/rpl39/sars2/uba52/rpl36a/rps26/eif1b/ssb/igf2bp3/rpl24/eif4eb/ptcd3/rpl27/rpl23/eif3s10/rpl31/rps24/rps16/rwdd1/denr/aars1/rpl18/rpl13a/rpl22/mrps16/rbm24a/rplp2/mrpl16/mettl17/mtrfr/drg1/malsu1/rps9/rbm8a/rpl28/rack1/rplp1/rpl23a/eif3f/gspt1l/mrpl2/exosc8/larp1b/eif3s6ip/rps15/rpsa/eef1g/eif2s2/cnbpa/ftsj1/rps11/eif4e3/rplp2l/zgc:114188/polr2d/rpl17/cnot11/faua/pabpc1a/exosc7/rpl19/rps5/eif2s1b/rplp0/rpl18a/exosc9/tufm/rpl21/kars1/rpl4/eef2b/jmjd4/cnbpb/rps13/rpl8/eif5b/rps27.1/rpl10/rpl5b/eif3g/eif3d/mrps35/rpl6/gfm1/eif4e1c/rps3/eif1axa/cnot7/eif2b4/rps12/eef1a1l1/znf598/rpl9/cnot10/eef2a.1/rpl10a/cars1/rps6/rps27a/mars1/nck2a/rps27.2/rps28/mrps7/eif4bb/rpl36/rps23/eif2s3/gtpbp1l/nck2b/eif4g1a/cnot9/dcp2/rps19/dph1/ybx1/dhx29/eif3ja/mrpl15/eif4g3b/samd4a |
| GO:0006412 | translation | 2.02950881785419 | 1.14653846153846e-08 | dazl/fars2/farsa/zc3h12ab/dph7/rpl26/mrpl10/dars2/wars2/mtfmt/trmt10c/gemin5/exosc2/mrpl58/mrps5/dars1/rpp30/zar1/mrpl9/dcp1b/piwil2/coa3a/trnau1apa/mrpl35/mif4gdb/mrpl30/eif2ak1/dcps/mrpl17/dph5/lars1b/mtg1/mrps17/mrps18b/mrpl4/si:ch211-256m1.8/zgc:171480/mars2/uqcc2/chchd1/tsfm/eif4ea/mrpl22/mettl3/ngdn/mtrf1l/faub/mrps18c/drg2/lsm1/mrpl27/mrpl20/dph6/magoh/eif1ad/eif2a/eefsec/qrsl1/mrpl51/mrps23/mcts1/ago3b/mettl16/gatb/mrpl18/mrps21/gtpbp3/mtif2/iars1/mrpl14/rpl12/hars/mrpl24/pdf/eif2b1/eif4e2rs1/abce1/eif2s1a/qars1/nars2/mrpl41/impact/vars1/mrpl34/fxr2/larp6a/mrps24/mrpl36/rpl35/cnot8/mrps12/mrpl13/rcc1l/eif5a2/rpl32/mrps2/rps7/mrpl12/brf1a/eif2b2/eif3m/eif4ebp3l/yars1/eef1b2/eif3ea/mrpl11/rpl22l1/mrpl33/rpl34/rars1/eif6/dph2/ogfod1/gspt1/pde12/mrpl47/rps21/rps17/eef1a1l3/eif3jb/tars1/cnot6l/eef1da/eif4ebp1/gars1/mrps10/pars2/iars2/rc3h2/mrpl23/rpl29/zgc:171772/rpl37/rps8a/gatc/rpl38/rps3a/rps14/rps15a/srp9/larp4b/mrps30/rps20/rps29/tcea3/rpl5a/rpl15/sars1/sepsecs/mrpl3/eif1axb/rpl35a/vars2/eif5/rps18/eif3ha/zc3h15/mrpl32/mrpl37/trnau1apb/mrps11/rpl11/rpl14/rpl39/sars2/uba52/rpl36a/rps26/eif1b/ssb/igf2bp3/rpl24/eif4eb/ptcd3/rpl27/rpl23/eif3s10/rpl31/rps24/rps16/rwdd1/denr/aars1/rpl18/rpl13a/rpl22/mrps16/rbm24a/rplp2/mrpl16/mettl17/mtrfr/drg1/malsu1/rps9/rbm8a/rpl28/rack1/rplp1/rpl23a/eif3f/gspt1l/mrpl2/exosc8/larp1b/eif3s6ip/rps15/rpsa/eef1g/eif2s2/cnbpa/ftsj1/rps11/eif4e3/rplp2l/zgc:114188/polr2d/rpl17/cnot11/faua/pabpc1a/exosc7/rpl19/rps5/eif2s1b/rplp0/rpl18a/exosc9/tufm/rpl21/kars1/rpl4/eef2b/jmjd4/cnbpb/rps13/rpl8/eif5b/rps27.1/rpl10/rpl5b/eif3g/eif3d/mrps35/rpl6/gfm1/eif4e1c/rps3/eif1axa/cnot7/eif2b4/rps12/eef1a1l1/znf598/rpl9/cnot10/eef2a.1/rpl10a/cars1/rps6/rps27a/mars1/nck2a/rps27.2/rps28/mrps7/eif4bb/rpl36/rps23/eif2s3/gtpbp1l/nck2b/eif4g1a/cnot9/dcp2/rps19/dph1/ybx1/dhx29/eif3ja/mrpl15/eif4g3b/samd4a |
| GO:0045333 | cellular respiration | 1.99293254414966 | 4.41989406426173e-05 | ccn6/tefm/sfxn4/uqcc3/si:dkey-85n7.8/ugl/sdhda/ndufs8b/atpsckmt/mdh1b/mdh1ab/dhtkd1/wfs1b/uqcc2/idh3b/sdhaf2/atp5mf/atp5po/sdhdb/ndufb6/lyrm7/cox7b/uqcr10/ndufb8/mrpl24/etfdh/ndufa10/cox6c/cox4i1/ndufv1/ndufaf1/suclg2/si:dkey-190g11.3/antkmt/cox7c/ndufa7/atp5f1e/ndufa8/ndufc2/cox7a2a/sdhc/ogdhb/slc25a55b/slc25a12/coa6/sdha/ndufs6/pm20d1.1/cox6a1/uqcrq/lyrm5a/ndufs2/ndufb9/noa1/sucla2/idh1/uqcrfs1/ndufs8a/cox6a2/cox5ab/lyrm5b/uqcrh/ndufa5/ndufv2/idh3a/mdh2/idh2/cox7a2l/sdhb/dlst/uqcrb/ndufs7/cox5aa/cyc1/ND3/fh/cox5b2/ndufv3/ogdha/cox8a |
| GO:0006518 | peptide metabolic process | 1.98118588803671 | 1.14653846153846e-08 | dazl/fars2/farsa/zc3h12ab/dph7/parla/rpl26/mrpl10/dars2/wars2/mtfmt/trmt10c/gemin5/exosc2/mrpl58/mrps5/dars1/rpp30/zar1/erap2/mrpl9/dcp1b/piwil2/ggt1b/coa3a/trnau1apa/mrpl35/mif4gdb/mrpl30/eif2ak1/dcps/mrpl17/dph5/lars1b/mtg1/mrps17/mrps18b/cpz/mrpl4/si:ch211-256m1.8/zgc:171480/mars2/uqcc2/chchd1/tsfm/gss/eif4ea/mrpl22/mettl3/ngdn/mtrf1l/faub/mrps18c/drg2/lsm1/mrpl27/mrpl20/dph6/magoh/eif1ad/eif2a/eefsec/qrsl1/mrpl51/mrps23/mcts1/ago3b/mettl16/gatb/mrpl18/immp2l/mrps21/gtpbp3/mtif2/iars1/mrpl14/rpl12/hars/mrpl24/pdf/enpep/eif2b1/eif4e2rs1/abce1/eif2s1a/qars1/nars2/mrpl41/pcsk5b/impact/vars1/mrpl34/sec11a/nln/fxr2/larp6a/mrps24/mrpl36/rpl35/cnot8/mrps12/mrpl13/rcc1l/eif5a2/rpl32/mrps2/rps7/mrpl12/brf1a/eif2b2/eif3m/eif4ebp3l/yars1/bdh2/eef1b2/eif3ea/mrpl11/rpl22l1/mrpl33/rpl34/rars1/eif6/dph2/ogfod1/gspt1/pde12/sppl3/mrpl47/rps21/rps17/eef1a1l3/eif3jb/tars1/cnot6l/eef1da/eif4ebp1/gars1/mrps10/pars2/iars2/rc3h2/mrpl23/rpl29/zgc:171772/rpl37/rps8a/gatc/rpl38/spcs3/rps3a/rps14/rps15a/srp9/larp4b/mrps30/rps20/rps29/tcea3/rpl5a/rpl15/sars1/sepsecs/mrpl3/eif1axb/rpl35a/vars2/eif5/rps18/eif3ha/zc3h15/mrpl32/mrpl37/trnau1apb/mrps11/rpl11/rpl14/rpl39/sars2/uba52/rpl36a/rps26/eif1b/thop1/ssb/igf2bp3/rpl24/eif4eb/ptcd3/rpl27/rpl23/eif3s10/rpl31/rps24/rps16/rwdd1/denr/aars1/rpl18/rpl13a/rpl22/mrps16/mipepb/rbm24a/spcs1/rplp2/mrpl16/mettl17/mtrfr/drg1/malsu1/rps9/rbm8a/rpl28/rack1/rplp1/rpl23a/eif3f/gspt1l/mrpl2/exosc8/larp1b/eif3s6ip/rps15/rpsa/eef1g/eif2s2/cnbpa/ftsj1/rps11/eif4e3/rplp2l/zgc:114188/polr2d/rpl17/cnot11/faua/pabpc1a/exosc7/rpl19/rps5/eif2s1b/rplp0/rpl18a/exosc9/tufm/rpl21/kars1/rpl4/eef2b/jmjd4/cnbpb/rps13/rpl8/eif5b/rps27.1/rpl10/rpl5b/eif3g/eif3d/mrps35/rpl6/gfm1/ide/eif4e1c/rps3/eif1axa/cnot7/eif2b4/rps12/eef1a1l1/znf598/rpl9/cnot10/eef2a.1/rpl10a/spcs2/cars1/rps6/rps27a/mars1/nck2a/rps27.2/rps28/mrps7/eif4bb/rpl36/rps23/eif2s3/gtpbp1l/nck2b/lnpep/eif4g1a/cnot9/dcp2/rps19/dph1/ybx1/dhx29/eif3ja/mrpl15/eif4g3b/samd4a |
| GO:0007005 | mitochondrion organization | 1.97295174722294 | 2.83369625164005e-07 | si:dkey-32e23.4/bnip4/adck1/sqstm1/mtm1/mtx1b/mtx1a/rrm2b/sco2/si:dkey-82o10.4/vps13d/sdhaf1/uqcc3/sdhaf3/chchd4b/atp23/phb2b/timm21/ppargc1a/timm29/maip1/coa3a/sdhaf4/aifm1/ndufs8b/chchd4a/alkbh7/chchd6b/cox17/chchd7/coa4/mgme1/cox20/slc25a42/bcs1l/timm13/uqcc2/surf1/sdhaf2/timm50/spata18/lonp1/tmem126a/dnajc19/grpel1/bida/chchd3b/atp13a2/immp2l/wdr45b/lyrm7/uqcc6/cox19/etfa/cluha/dna2/coa5/snx7/apooa/hspd1/atpaf2/phb/ndufaf1/atg5/snx30/polg2/uqcc1/higd1a/chchd10/cox7a2a/primpol/atpaf1/higd2a/si:ch211-166a6.5/pet100/tomm40/apool/fam162a/agtpbp1/pmaip1/romo1/ndufa11/oxa1l/twnk/samm50/rpl37/ndufs8a/rhot1a/slc25a4/nubpl/ndufaf8/ttc19/dnajc11a/mfn1b/miga1/tmem70/dnlz/dnaja3a/mipepb/foxred1/cox7a2l/dlst/atg7/tafazzin/tomm40l/fis1 |
| GO:0002088 | lens development in camera-type eye | 1.96162227904054 | 0.000326438123176076 | fgf19/crygm2a/crybb2/crybb1l3/crygm5/crygm2d6/crygm2b/mipb/cryba1l2/crygm2d9/crygm2d15/dnajb1b/crygm2d12/crygm2d18/cryba2a/crygm2d17/crygm2d7/crygm2d1/lgsn/crygm2d5/crygm2f/cryba1l1/crybb1l2/dlx3b/crygm2d3/crygm2d20/crygm2d11/crygm2d10/crygm2d2/crygm2d21/cryba1a/dlx4b/mipa/lama1/crygm2d13/crygmxl2/crygn2/crygm2d16/cryba4/crygm2d8/tdrd7b/crygm2d14/cryba1b/crygn1/psmc3/foxe3/cryba2b/hsf4/unc45b/crygmx/crygmxl1/si:dkey-57a22.15/sox11b/crybb1l1/psmd6 |
| GO:0009145 | purine nucleoside triphosphate biosynthetic process | 1.93013788013767 | 0.00988355843878473 | nme9/uqcc3/atp5mc3a/atpsckmt/atp5mf/atp5pd/atp5po/atp5mea/atp5mc1/nme2b.2/nme6/atp5f1c/mrpl24/atp5meb/nme2a/si:dkey-190g11.3/antkmt/atp5f1e/nme3/atp5pf/atp5pb/atp5mc3b/nme7/atp5fa1/dlst/ATP6 |
| GO:0009206 | purine ribonucleoside triphosphate biosynthetic process | 1.93013788013767 | 0.00988355843878473 | nme9/uqcc3/atp5mc3a/atpsckmt/atp5mf/atp5pd/atp5po/atp5mea/atp5mc1/nme2b.2/nme6/atp5f1c/mrpl24/atp5meb/nme2a/si:dkey-190g11.3/antkmt/atp5f1e/nme3/atp5pf/atp5pb/atp5mc3b/nme7/atp5fa1/dlst/ATP6 |
| GO:0008202 | steroid metabolic process | 1.89342793339491 | 0.000798277428982877 | ch25h/sdr42e1/dhcr24/cyp46a1.1/ch25hl1.1/msmo1/amacr/nsdhl/cyp46a1.3/cyp39a1/cyp46a1.2/hmgcra/mvk/ebpl/hsd17b12b/bdh1/srd5a1/cyp51/gba/cyp21a2/cyp7a1/lipeb/slc27a2b/faxdc2/cyp46a1.4/apoea/cyp2r1/mvda/ch25hl2/fgfr4/lss/hmgcs1/ebp/cyp26c1/apoa1a/cyp27a1.4/hsd17b7/cyb5r2/cyp27a1.1/hsd17b14/rdh1/srd5a2b/erlin1/slc27a2a |
| GO:0006979 | response to oxidative stress | 1.83959019549846 | 0.00355248534040968 | bco2l/fancc/anxa1b/glrx2/keap1a/per2/prdx1/ptgs2a/sod2/park7/ptgs1/ptgs2b/lonp1/prdx3/si:ch211-120k19.1/trpm2/gpx4a/diabloa/impact/gpx1a/gpx9/hmox2a/gpx4b/gpx8/naprt/mpv17/brf2/sirt2/romo1/sod1/txndc12/ndufa6/mpv17l2/hmox1a/ryr1b/msraa |
| GO:0006260 | DNA replication | 1.7556677318185 | 0.00194590276805455 | dbf4/zranb3/polq/zgc:173742/wrn/lig4/pold3/rpa3/mcm8/wdr18/tonsl/terf1/rmi1/poln/ctc1/mgme1/rnaseh1/ssrp1a/gins3/rpa2/stn1/orc4/dna2/noc3l/mcm7/rfc5/gins2/orc5/tatdn1/pole2/polg2/rrm1/tipin/fen1/primpol/mcm3/rrm2/traip/orc1/lig1/igfbp2a/mcmbp/mcm2/wrnip1/twnk/recql/rad51/rfc2/mcm5/rpa1/esco2/cenps/mms22l/eme1/orc6/rbbp4/gins4/blm/rnaseh2a/cdc45/prim1/rfc3/helb/pcna/rbb4l/lrwd1/samhd1/orc3/dscc1/ssbp1/top1a/mcm4/pole |
| GO:0044743 | protein transmembrane import into intracellular organelle | 1.73136835035816 | 0.0470692918806812 | pex10/chchd4b/timm21/chchd4a/timm50/dnajc19/grpel1/hspd1/tomm40/romo1/pex1/dnlz/pex13/tomm40l/pex5/timm23a/tomm70a/tomm7/pex7/pam16/tomm20b/pex26 |
| GO:0050731 | positive regulation of peptidyl-tyrosine phosphorylation | 1.72535656967289 | 0.0678490288954661 | unc119a/epgn/arl2bp/btc/smtlb/hbegfa/zgc:153759 |
| GO:0034599 | cellular response to oxidative stress | 1.71630178467435 | 0.0639818065496334 | bco2l/fancc/anxa1b/glrx2/keap1a/prdx1/sod2/park7/lonp1/si:ch211-120k19.1/diabloa/impact/mpv17/brf2/sirt2/romo1/sod1/mpv17l2/ryr1b/msraa |
| GO:0042744 | hydrogen peroxide catabolic process | 1.71035395380329 | 0.0737625497040733 | prdx1/hbae5/zgc:163057/hbbe2/prdx3/hbae1.3/hbae1.1/hbae3/gpx1a |
| GO:0006974 | cellular response to DNA damage stimulus | 1.70302954476004 | 6.9422407694291e-06 | uba7/wdr76/hrob/mms19/ddias/cry5/slx1b/zswim7/fancc/zranb3/tdp2b/xpc/parpbp/rad51d/swsap1/chek2/cry-dash/si:ch211-127m7.2/msh3/palb2/polq/hic1/phkg1b/xndc1/neil1/rfwd3/rrm2b/ogg1/sfr1/wrn/fanci/rad54b/brip1/cpdp/lig4/pargl/mre11a/zgc:112496/rad51b/faap100/pold3/rpa3/atm/cdc7/mcm8/nthl1/mcm9/ddb2/ap5s1/ercc2/kmt5ab/aptx/nbn/tonsl/cep63/eya4/rmi1/alkbh7/poln/mnat1/si:dkey-74k8.3/mgme1/park7/dek/mpg/smyd2a/mettl3/spata18/polh/ssrp1a/tdp1/neil3/rpa2/stn1/poldip2/xrcc3/gtf2h3/trip13/zfyve26/fancg/msh2/dna2/mcm7/smarcad1a/rfc5/uba1/swi5/gins2/fxr2/xrcc6/rad51c/pole2/polg2/tipin/yars1/ercc6/ube2nb/prmt6/fen1/primpol/tigara/mcm3/unga/sem1/bcl2l16/rad23b/pmaip1/lig1/polr2i/bccip/nek4/vhl/mcm2/abraxas1/txndc12/rbbp8/ttf2/mlh1/rmi2/wrnip1/recql/poli/rad1/rad51/rfc2/mcm5/rpa1/ercc1/plrg1/smyd2b/cenps/psme4a/mms22l/eme1/cdca5/si:ch211-141o9.10/hltf |
| GO:0044283 | small molecule biosynthetic process | 1.67134004649573 | 0.000341676076873402 | gls2b/tecrl2b/pycr3/fads2/mcat/msmo1/sdsl/asns/gatm/chst15/elovl8a/gpib/anxa1b/prxl2b/mthfr/acacb/cyp39a1/elovl2/pycr1b/hmgcra/mvk/gch2/qdprb.1/otc/uros/ptgs2a/olah/hsd17b12b/glsl/coq5/aass/akr1a1b/srd5a3/moxd1l/pdss2/asnsd1/srr/cyp51/cyp7a1/adka/park7/lgsn/slc27a2b/apip/rfk/ptgs1/ptgs2b/mvda/pnpo/abhd2a/pcbd1/kmo/ptgesl/pah/elovl6l/fgfr4/tecra/asah2/lss/hmgcs1/ebp/apoa1a/cyp27a1.4/gch1/acmsd/hsd17b7/gnmt/coq4/pck2/elovl8b/sephs1/cdo1/ptges3b/gamt/cyp27a1.1/ndufab1b/fbp1b/impa1/bhmt/upb1/acot7/rgn/fa2h/slc27a2a/aprt/sptlc3/prtfdc1b/agxtb/g6pc3/hprt1/pdss1/ubiad1/coq6/cth/ndufa9a/qdpra |
| GO:0052646 | alditol phosphate metabolic process | 1.62136572553731 | 0.0983308153791125 | gpd1l/rfk/gpam/gpd1c/gpd1a/gpd2 |
| GO:0032259 | methylation | 1.61555246309617 | 0.00489199162953128 | hnmt/thumpd2/suv39h1a/zgc:113054/mettl21cb/wdr4/tpmt.2/etfbkmt/mrm3a/dnmt3bb.2/eef1akmt1/cskmt/trmt10c/dnmt3bb.3/emg1/ash2l/tyms/kmt5ab/henmt1/piwil2/coq5/trmt61a/vcpkmt/mrm3b/trmt44/atpsckmt/mettl4/trmt6/tyw3/trmt2a/dph5/trmt12/nsun2/eef1akmt2/thumpd3/tfb1m/prmt9/smyd2a/tpmt.1/hells/prmt7/mettl3/smyd1a/prmt3/trmt11/trmt112/dimt1l/bud23/ntmt1/mettl16/mettl21a/mettl2a/thada/nsun5/gtpbp3/amt/trmt10b/tdrd9/dnmt3bb.1/gnmt/zcchc4/gamt/cebpb/si:dkey-190g11.3/antkmt/cxxc1b/rrp8/mrm2/setd3/bhmt/prmt6/skic8/trmt61b/comta/lyn/unga/nop2/comtd1 |
| GO:0043436 | oxoacid metabolic process | 1.61491700112608 | 2.31796694375566e-05 | gls2b/tecrl2b/pycr3/fars2/fads2/ddo/mcat/farsa/amacr/sdsl/accs/etfbkmt/asns/si:dkey-238o13.4/gatm/elovl8a/gpib/anxa1b/acot20/prxl2b/dars2/acot17/wars2/mthfr/asah1a/acot11b/ugl/sdhaf3/acacb/cyp39a1/elovl2/hoga1/pycr1b/dars1/nit2/qdprb.1/gatd1/otc/abcd3b/uros/ptgs2a/olah/hsd17b12b/glsl/tdo2a/gcat/enosf1/uevld/aass/amdhd2/alkbh7/asnsd1/srr/mdh1b/mdh1ab/dhtkd1/marc1/lars1b/lpin1b/cyp7a1/tysnd1/park7/si:ch211-256m1.8/acaa1/tat/acads/lgsn/slc27a2b/mars2/apip/ptgs1/idh3b/ptgs2b/pnpo/tph1b/gpam/abhd2a/hacl1/hsd17b4/cpt2/adpgk2/kyat1/decr1/kmo/qrsl1/bpgm/ptgesl/gck/pah/elovl6l/si:ch211-212d10.2/gatb/zgc:162816/fgfr4/tecra/gtpbp3/asah2/iars1/amt/etfa/cpt1b/hars/acss1/gcshb/cmah/cyp26c1/ctps1a/acadm/qars1/nars2/vars1/hadhab/auh/bckdha/acmsd/gnmt/pck2/elovl8b/sephs1/cdo1/mrps36/ptges3b/eno1b/abcd3a/gamt/pgam2/gnpda1/ndufab1b/mccc2/pklr/yars1/aacs/hadh/gcdha/me1/bhmt/hpdb/upb1/acot7/rgn/rars1/tigara/fa2h/slc27a2a/kyat3.2/hibch/mpst/acss2/echdc2/eci1/hgd/agxtb/pm20d1.1/tars1/hadhaa/gars1/pars2/cth/qdpra/ptgr1.1/iars2/acsf2/aspg/gcsha/kynu/hadhb/acsl5/glud1b/got2b/hpdl/cpt1aa/hk2/idh1/acsl1b/sars1/ivd/acat2/vars2/acaa2/ehhadh/acot18/nags/amdhd1/sars2/phyh/ftcd/sgpl1/gip/slc25a44b/shmt1/pfas/acot15/mgst2/aldoab/elovl4b/idh3a/tecrb/aars1/ldha/elovl7a |
| GO:0044282 | small molecule catabolic process | 1.59271540808738 | 0.00441611574194559 | gls2b/cyp46a1.1/ddo/sdsl/etfbkmt/si:dkey-238o13.4/cyp46a1.3/bpnt1/cyp46a1.2/hoga1/qdprb.1/gatd1/abcd3b/glsl/tdo2a/gcat/enosf1/aass/srd5a3/amdhd2/srr/zgc:103586/rbks/miox/lpin1b/cyp7a1/tysnd1/acaa1/tat/acads/cyp46a1.4/apoea/uraha/sord/abhd2a/hacl1/hsd17b4/cpt2/decr1/kmo/pah/zgc:162816/amt/etfa/cpt1b/gcshb/cdaa/acadm/hadhab/auh/bckdha/pck2/cdo1/abcd3a/adh8b/gnpda1/mccc2/impa1/hadh/gcdha/hpdb/kyat3.2/hibch/echdc2/eci1/hgd/hadhaa/qdpra/gcsha/kynu/hadhb/glud1b/got2b/esd/cpt1aa/ivd/acat2/acaa2/ehhadh/amdhd1/phyh/ftcd/slc25a44b/shmt1 |
| GO:1901361 | organic cyclic compound catabolic process | 1.58649464452635 | 0.00292032014134612 | hnmt/nudt4b/cyp46a1.1/dazl/dnase1l1/blvra/fastkd3/fitm2/zc3h12ab/fastkd2/cyp46a1.3/tbrg4/nudt18/exosc2/cyp46a1.2/fastkd1/exosc6/qdprb.1/entpd3/dcp1b/pde4cb/tdo2a/nudt15/moxd1l/zgc:103586/dut/heatr1/mrto4/dcps/exosc5/cyp7a1/gda/acaa1/tat/dctpp1/cyp46a1.4/tent5bb/apoea/rnaseh1/mettl3/uraha/lsm1/magoh/kmo/lsm3/pah/rexo4/ago3b/mettl16/gtpbp3/xdh/cdaa/dna2/nudt4a/fxr2/cnot8/rnaset2l/entpd5b/dnase1l4.1/hsd17b14/hmox2a/fhit/hint1/dpys/hpdb/comta/fen1/kyat3.2/pde12/allc/hgd/cnot6l/edc3/nbas/dis3/qdpra/plex9.2/rc3h2/nudt12/kynu/tet2/syncripl/endog/hmox1a/mtrex/entpd5a |
| GO:0071826 | ribonucleoprotein complex subunit organization | 1.57792704614773 | 0.0130343312989884 | nle1/gemin5/zfand1/abt1/znhit3/pwp2h/trnau1apa/mrto4/snrpd2/prmt7/pih1d1/gemin2/mrpl20/snrpg/bxdc2/prpf18/mcts1/ppan/mdn1/clp1/ptges3b/rrp7a/eif3m/nopchap1/snrpd3l/eif3ea/coil/eif6/snrpf/usp39/eif3jb/snrpe/rpl38/rps14/shq1/rpl5a/eif5/eif3ha/trnau1apb/mrps11/scaf11/rpf2/eif3s10/cdc73/denr/ddx28/naf1/rpl23a/eif3s6ip/rps15/rpsa/eif2s2/polr2d/srpk1a/rps5/gemin8/rplp0/snrpd1/rps27.1/aar2/rpl10/rpl5b/eif3g/eif3d/rpl6/yju2/rps27.2/rps28/mrps7/gemin6/eif2s3/nudt21/dyrk3/lsm4/snrpc/rps19/eif3ja/eif3c |
| GO:0055088 | lipid homeostasis | 1.54238929377417 | 0.0961825238314319 | adck1/fitm2/zgc:162608/cyp39a1/spx/abcg4b/ormdl2/apobb.1/psmd10/apobb.2/cyp7a1/apoa4a/pnpla3/tlcd1/apoa4b.2/apoba/fabp2/angptl8 |
| GO:0140014 | mitotic nuclear division | 1.53553828880172 | 0.0229731377900925 | ncaph/epgn/btc/pdcd2/nde1/chtf8/si:ch211-244o22.2/cep192/sgo1/map10/mad1l1/incenp/gnl3/kif20bb/ndc80/anapc5/gcna/ncapg/bub3/nme6/mad2l1/reep3b/anapc15/birc5b/cdca8/esco1/ncapg2/abraxas2/ccnb1/ranbp1/aurkb/racgap1/anapc11/ube2s/zgc:86764/pmaip1/bccip/abraxas1/zw10/esco2/ncaph2/cenpk/spdl1/prc1b/ran/cdca5/gnl2/map9/mki67/kifc1/ube2ib/chmp7/kif11 |
| GO:0019439 | aromatic compound catabolic process | 1.4470736645875 | 0.0239204409208845 | hnmt/nudt4b/dazl/dnase1l1/blvra/fastkd3/fitm2/zc3h12ab/fastkd2/tbrg4/nudt18/exosc2/fastkd1/exosc6/qdprb.1/entpd3/dcp1b/pde4cb/tdo2a/nudt15/moxd1l/zgc:103586/dut/heatr1/mrto4/dcps/exosc5/gda/acaa1/tat/dctpp1/tent5bb/rnaseh1/mettl3/uraha/lsm1/magoh/kmo/lsm3/pah/rexo4/ago3b/mettl16/gtpbp3/xdh/cdaa/dna2/nudt4a/fxr2/cnot8/rnaset2l/entpd5b/dnase1l4.1/hmox2a/fhit/hint1/hpdb/comta/fen1/kyat3.2/pde12/hgd/cnot6l/edc3/nbas/dis3/qdpra/plex9.2/rc3h2/nudt12/kynu/tet2/syncripl/endog/hmox1a/mtrex/entpd5a/ephx1 |
| GO:0044270 | cellular nitrogen compound catabolic process | 1.42325048500041 | 0.0467129045300582 | hnmt/nudt4b/dazl/dnase1l1/blvra/fastkd3/fitm2/zc3h12ab/fastkd2/tbrg4/nudt18/exosc2/fastkd1/exosc6/entpd3/dcp1b/pde4cb/tdo2a/nudt15/zgc:103586/dut/heatr1/mrto4/dcps/exosc5/gda/dctpp1/tent5bb/rnaseh1/mettl3/uraha/lsm1/magoh/kmo/lsm3/rexo4/ago3b/mettl16/gtpbp3/xdh/cdaa/dna2/nudt4a/fxr2/cnot8/rnaset2l/entpd5b/dnase1l4.1/hmox2a/fhit/hint1/dpys/bhmt/fen1/pde12/cnot6l/edc3/nbas/dis3/plex9.2/rc3h2/nudt12/kynu/tet2/syncripl/endog/hmox1a/mtrex/entpd5a |
| GO:0007601 | visual perception | 1.4025303608205 | 0.0873969864431149 | unc119a/gprc5c/crygm2a/crybb2/crybb1l3/crygm5/grk7a/crygm2d6/crygm2b/cryba1l2/arr3b/crygm2d9/crygm2d15/crygm2d12/crygm2d18/cryba2a/opn1sw1/crygm2d17/crygm2d7/crygm2d1/arl6/crygm2d5/crygm2f/cryba1l1/pdcb/crybb1l2/crygm2d3/crygm2d20/crygm2d11/crygm2d10/crygm2d2/crygm2d21/cryba1a/opn1mw1/crygm2d13/crygmxl2/guca1b/crygn2/crygm2d16/arr3a/cryba4/crygm2d8/crygm2d14/cryba1b/crygn1/prph2a/lum/cryba2b/opn1sw2/crygmx/pdca/crygmxl1/si:dkey-57a22.15/crybb1l1/pde6a/gucy2f/grk7b/opn1lw2/nphp3/prph2lb/pde6d/crybgx/tmtops2a |
| GO:0099537 | trans-synaptic signaling | -2.69921105199218 | 1.14653846153846e-08 | dlg1b/dtnbp1b/ncdn/gabrr2a/gabrb1/cplx2b/drd4-rs/cacng8b/otofa/sv2bb/dlg3/cdh2/penka/cep89/chrnb5a/synj1/cart2/cspg5b/gria3a/dlgap1a/chrm2a/snap25a/napbb/penkb/glrbb/slc5a7a/napba/chata/spg11/apba1b/erc1a/grid1a/stxbp1b/cplx2a/clstn2b/cacnb1/gabrb3/chrng/hdc/sncb/glra1/drd3/cacng5a/si:ch211-81a5.8/syn1/erc2/gria1a/grm4/grm6b/stx3a/caskb/chrnb3a/ppp1r9ba/slc18a3a/snap25b/jph3b/mctp2b/grm8b/cart3/slc6a5/calb1/chrna3/htr1aa/calb2b/snap47/htr1d/pdyn/jph3a/cacnb2b/ptchd1/calb2a/syt7b/syn2a/cacng7b/chrna5/grin2aa/rims1b/shisa9a/clstn1/syt4/rab3ab/dlgap4b/lrrc4ca/chrna2a/gabra4/glra4b/nlgn4xa/gria2b/nsmfb/gria4a/slc17a6b/clstn2a/sncgb/htr3a/gabrb2a/lrrc4.1/cadpsb/grin1a/grm2a/ppp1r9bb/syt11a/unc13a/cacng2b/stxbp1a/syt1a/chrnb2/slc12a5b/chrna6/cacng3b/shisa7b/nlgn3a/gria3b/stx1b/dlg2/ppfia3/grin2ab/cacng2a/grm8a/lrrc4cb/atcaya/slc12a5a/chrna4b/grm2b/nlgn3b/cacnb4b/grid2/clstn3/syt2a/apba2b/grik1a/cspg5a/sv2a/pnocb/grin1b/baiap3/cacna1ab/grin2da/nlgn2a/nrxn2a/si:dkey-174m14.3/drd2a/rph3ab/dagla/grik5/htr1fa/shisa9b/gabrp/chrm4a/grin3bb/grin2bb/insyn1/grik4/gabrg2/grm1a/cacng8a/stx11a/gabrb4/dlgap3/htr7c/neto1l/nalcn/gria4b |
| GO:0007268 | chemical synaptic transmission | -2.68729704840182 | 1.14653846153846e-08 | dlg1b/dtnbp1b/ncdn/gabrr2a/gabrb1/cplx2b/drd4-rs/cacng8b/otofa/sv2bb/dlg3/cdh2/penka/cep89/chrnb5a/synj1/cart2/cspg5b/gria3a/dlgap1a/chrm2a/snap25a/napbb/penkb/glrbb/slc5a7a/napba/chata/spg11/apba1b/erc1a/grid1a/stxbp1b/cplx2a/clstn2b/cacnb1/gabrb3/chrng/hdc/sncb/glra1/drd3/cacng5a/si:ch211-81a5.8/syn1/erc2/gria1a/grm4/grm6b/stx3a/caskb/chrnb3a/ppp1r9ba/slc18a3a/snap25b/jph3b/mctp2b/grm8b/cart3/slc6a5/calb1/chrna3/htr1aa/calb2b/snap47/htr1d/pdyn/jph3a/cacnb2b/ptchd1/calb2a/syt7b/syn2a/cacng7b/chrna5/grin2aa/rims1b/shisa9a/clstn1/syt4/rab3ab/dlgap4b/lrrc4ca/chrna2a/gabra4/glra4b/nlgn4xa/gria2b/nsmfb/gria4a/slc17a6b/clstn2a/sncgb/htr3a/gabrb2a/lrrc4.1/cadpsb/grin1a/grm2a/ppp1r9bb/syt11a/unc13a/cacng2b/stxbp1a/syt1a/chrnb2/slc12a5b/chrna6/cacng3b/shisa7b/nlgn3a/gria3b/stx1b/dlg2/ppfia3/grin2ab/cacng2a/grm8a/lrrc4cb/atcaya/slc12a5a/chrna4b/grm2b/nlgn3b/cacnb4b/grid2/clstn3/syt2a/apba2b/grik1a/cspg5a/sv2a/pnocb/grin1b/baiap3/cacna1ab/grin2da/nlgn2a/nrxn2a/si:dkey-174m14.3/drd2a/rph3ab/grik5/htr1fa/shisa9b/gabrp/chrm4a/grin3bb/grin2bb/insyn1/grik4/gabrg2/grm1a/cacng8a/stx11a/gabrb4/dlgap3/htr7c/neto1l/nalcn/gria4b |
| GO:0098916 | anterograde trans-synaptic signaling | -2.68729704840182 | 1.14653846153846e-08 | dlg1b/dtnbp1b/ncdn/gabrr2a/gabrb1/cplx2b/drd4-rs/cacng8b/otofa/sv2bb/dlg3/cdh2/penka/cep89/chrnb5a/synj1/cart2/cspg5b/gria3a/dlgap1a/chrm2a/snap25a/napbb/penkb/glrbb/slc5a7a/napba/chata/spg11/apba1b/erc1a/grid1a/stxbp1b/cplx2a/clstn2b/cacnb1/gabrb3/chrng/hdc/sncb/glra1/drd3/cacng5a/si:ch211-81a5.8/syn1/erc2/gria1a/grm4/grm6b/stx3a/caskb/chrnb3a/ppp1r9ba/slc18a3a/snap25b/jph3b/mctp2b/grm8b/cart3/slc6a5/calb1/chrna3/htr1aa/calb2b/snap47/htr1d/pdyn/jph3a/cacnb2b/ptchd1/calb2a/syt7b/syn2a/cacng7b/chrna5/grin2aa/rims1b/shisa9a/clstn1/syt4/rab3ab/dlgap4b/lrrc4ca/chrna2a/gabra4/glra4b/nlgn4xa/gria2b/nsmfb/gria4a/slc17a6b/clstn2a/sncgb/htr3a/gabrb2a/lrrc4.1/cadpsb/grin1a/grm2a/ppp1r9bb/syt11a/unc13a/cacng2b/stxbp1a/syt1a/chrnb2/slc12a5b/chrna6/cacng3b/shisa7b/nlgn3a/gria3b/stx1b/dlg2/ppfia3/grin2ab/cacng2a/grm8a/lrrc4cb/atcaya/slc12a5a/chrna4b/grm2b/nlgn3b/cacnb4b/grid2/clstn3/syt2a/apba2b/grik1a/cspg5a/sv2a/pnocb/grin1b/baiap3/cacna1ab/grin2da/nlgn2a/nrxn2a/si:dkey-174m14.3/drd2a/rph3ab/grik5/htr1fa/shisa9b/gabrp/chrm4a/grin3bb/grin2bb/insyn1/grik4/gabrg2/grm1a/cacng8a/stx11a/gabrb4/dlgap3/htr7c/neto1l/nalcn/gria4b |
| GO:0050804 | modulation of chemical synaptic transmission | -2.68416739586281 | 1.14653846153846e-08 | erc1a/grid1a/cplx2a/clstn2b/drd3/cacng5a/si:ch211-81a5.8/erc2/gria1a/grm4/grm6b/caskb/ppp1r9ba/jph3b/mctp2b/grm8b/calb1/calb2b/jph3a/calb2a/cacng7b/grin2aa/rims1b/shisa9a/clstn1/dlgap4b/lrrc4ca/nlgn4xa/gria2b/nsmfb/gria4a/clstn2a/lrrc4.1/grm2a/ppp1r9bb/cacng2b/cacng3b/shisa7b/nlgn3a/gria3b/stx1b/grin2ab/cacng2a/grm8a/lrrc4cb/atcaya/grm2b/nlgn3b/grid2/clstn3/grik1a/baiap3/grin2da/nlgn2a/si:dkey-174m14.3/drd2a/grik5/shisa9b/grin3bb/grin2bb/grik4/grm1a/cacng8a/dlgap3/neto1l/nalcn/gria4b |
| GO:0099177 | regulation of trans-synaptic signaling | -2.68416739586281 | 1.14653846153846e-08 | erc1a/grid1a/cplx2a/clstn2b/drd3/cacng5a/si:ch211-81a5.8/erc2/gria1a/grm4/grm6b/caskb/ppp1r9ba/jph3b/mctp2b/grm8b/calb1/calb2b/jph3a/calb2a/cacng7b/grin2aa/rims1b/shisa9a/clstn1/dlgap4b/lrrc4ca/nlgn4xa/gria2b/nsmfb/gria4a/clstn2a/lrrc4.1/grm2a/ppp1r9bb/cacng2b/cacng3b/shisa7b/nlgn3a/gria3b/stx1b/grin2ab/cacng2a/grm8a/lrrc4cb/atcaya/grm2b/nlgn3b/grid2/clstn3/grik1a/baiap3/grin2da/nlgn2a/si:dkey-174m14.3/drd2a/grik5/shisa9b/grin3bb/grin2bb/grik4/grm1a/cacng8a/dlgap3/neto1l/nalcn/gria4b |
| GO:0099536 | synaptic signaling | -2.6833837570563 | 1.14653846153846e-08 | dlg1b/dtnbp1b/ncdn/gabrr2a/gabrb1/cplx2b/drd4-rs/cacng8b/otofa/sv2bb/dlg3/cdh2/penka/cep89/chrnb5a/synj1/cart2/cspg5b/gria3a/dlgap1a/chrm2a/snap25a/napbb/penkb/glrbb/slc5a7a/napba/chata/spg11/apba1b/erc1a/grid1a/stxbp1b/cplx2a/clstn2b/cacnb1/gabrb3/chrng/hdc/sncb/glra1/drd3/cacng5a/si:ch211-81a5.8/syn1/erc2/gria1a/grm4/grm6b/stx3a/caskb/chrnb3a/ppp1r9ba/slc18a3a/snap25b/jph3b/mctp2b/grm8b/cart3/slc6a5/calb1/chrna3/htr1aa/calb2b/snap47/htr1d/pdyn/jph3a/cacnb2b/ptchd1/calb2a/syt7b/syn2a/cacng7b/chrna5/grin2aa/rims1b/shisa9a/clstn1/syt4/rab3ab/dlgap4b/lrrc4ca/chrna2a/gabra4/glra4b/nlgn4xa/gria2b/nsmfb/gria4a/slc17a6b/clstn2a/sncgb/htr3a/gabrb2a/lrrc4.1/cadpsb/grin1a/grm2a/ppp1r9bb/syt11a/unc13a/cacng2b/stxbp1a/syt1a/chrnb2/slc12a5b/chrna6/cacng3b/shisa7b/nlgn3a/gria3b/stx1b/dlg2/ppfia3/grin2ab/cacng2a/grm8a/lrrc4cb/atcaya/slc12a5a/chrna4b/grm2b/nlgn3b/cacnb4b/grid2/clstn3/syt2a/apba2b/grik1a/cspg5a/sv2a/pnocb/grin1b/baiap3/cacna1ab/grin2da/nlgn2a/nrxn2a/si:dkey-174m14.3/drd2a/rph3ab/dagla/grik5/htr1fa/shisa9b/gabrp/chrm4a/grin3bb/grin2bb/insyn1/grik4/gabrg2/grm1a/cacng8a/stx11a/gabrb4/scn1lab/dlgap3/htr7c/neto1l/nalcn/gria4b |
| GO:0098742 | cell-cell adhesion via plasma-membrane adhesion molecules | -2.34722281326859 | 1.14653846153846e-08 | arvcfb/zeb1a/cdh2/prnprs3/ptk7a/pcdh2ab10/zgc:165604/si:ch211-186j3.6/tenm3/dchs1a/ret/pcdh8/ptprsa/clstn2b/dscama/pcdh17/pcdh2ab9/pecam1a/pcdh10b/pcdh7a/tenm2b/robo3/si:ch211-199f5.1/dsc2l/pcdh1g22/jag2b/nectin3b/tenm4/pcdh1g31/pcdh7b/pcdh1g29/pcdh2g13/pcdh1a/clstn1/pcdh19/cadm1b/lrrc4ca/pcdh1a4/pcdh2ab3/cadm3/dscamb/ptprdb/pcdh10a/pcdh11/clstn2a/lrrc4.1/igsf21b/robo4/fat3a/cadm4/si:ch211-214p13.3/cdh18a/si:dkey-237h12.3/lrrc4cb/igsf21a/tenm2a/esama/cdh13/clstn3/cdh15/cntn3a.1/si:ch73-233f7.1/pcdh1g32/cdh4/pcdh1g11/si:ch73-233f7.3/cdh7a/pcdh1g18/adgrl1a/pcdh1gc5 |
| GO:0050808 | synapse organization | -2.21669824394954 | 9.94928637272983e-08 | dbn1/ctnna2/cdh2/synj1/ptk7a/dbnlb/cbln2b/ephb3a/sez6a/tjp1b/ctbp2a/erc1a/fbxo45/gphnb/neurod2/ptprsa/clstn2b/cacnb1/sncb/erc2/tanc2a/ppfia4/gfpt1/slitrk4/gphna/cacnb2b/myo9ab/inpp5ka/caprin2/pcloa/clstn1/rab3ab/lrrc4ca/ptprdb/nlgn4xa/clstn2a/sncgb/sybu/lrrc4.1/arhgef9b/unc13a/dock10/ctnnd2b/nbeaa/nlgn3a/bsnb/cacna1da/ppfia3/lrrc4cb/epb41l3a/grna/bdnf/nlgn3b/ctnnd2a/cacnb4b/pclob/clstn3/slitrk2/sez6l2/nlgn2a/nrxn2a |
| GO:0048667 | cell morphogenesis involved in neuron differentiation | -2.08310048291656 | 1.14653846153846e-08 | plxnb1b/pik3cd/mycbp2/rb1/ptprfb/zgc:100846/ptprfa/isl2b/nrp2b/arhgap35a/rapgef1a/whrna/emb/pik3ca/rbpms2a/aplp2/mapk8ip3/npr1a/sema3fa/emx2/olfm1a/unc5b/nptna/vegfc/dbn1/kctd12.1/si:ch211-129c21.1/phox2bb/unk/ubap1/ctnna2/cntn4/cdh2/trarg1a/kalrnb/ulk2/ptk7a/dbnlb/efna1b/ephb3a/slc35b2/rbpms2b/fus/map1ab/dock7/foxb1a/nsfa/rab33ba/sipa1l1/nrp2a/ush1ga/pnpla6/kctd13/isl1a/fezf2/slit2/tenm3/rufy3/nfasca/hecw1b/sim1a/adcy8/pak1/spg11/ret/kif1b/arnt2/sema3bl/zdhhc17/plxna4/appa/nedd4l/arhgap35b/efna2a/dscama/apc/robo2/pcdh17/casp9/epha4a/hdc/anos1a/trioa/lhx9/epha7/rtn4r/cyfip2/dpysl3/draxinb/slit1b/robo3/usp33/map1sb/appb/tanc2a/dzip1/sema5a/kifbp/chad/robo1/tnr/whrnb/tenm4/hecw2a/epha4l/kif5c/slitrk4/islr2/nptnb/lhx2b/myo9ab/caprin2/kalrna/cdkl5/ephb2b/epha4b/clstn1/rac3a/brsk1b/aplp1/rab3ab/gap43/lrrc4ca/srrm4/lhx1a/map1aa/prmt8b/atl1/nkx2.2a/arhgef25b/dscamb/triob/tmem59l/myo10l3/nsmfb/slit3/dpysl5a/chst11/fzd3a/kif5aa/robo4/efnb3b/vax1/stat3/dock10/ctnnd2b/zgc:77784/sema4c/si:ch211-113g11.6/rac1l/elavl4/myo10l1/ntn1b/scn8aa/lrrc4cb/plppr4a/grna/bdnf/ctnnd2a/slitrk2/nkx6.1/ncam1b/cntn3a.1/nrxn2a/dagla/grxcr1a/efna3a/dpysl5b/thsd7aa/si:ch211-241e1.3/plxna3/l1camb/plxna1a/nrcama/sema6ba/sema3b |
| GO:0048858 | cell projection morphogenesis | -2.05831448361924 | 1.14653846153846e-08 | plxnb1b/tmem108/pik3cd/mycbp2/rb1/ptprfb/zgc:100846/ptprfa/spag6/isl2b/nrp2b/arhgap35a/rapgef1a/emb/pik3ca/rbpms2a/aplp2/mapk8ip3/npr1a/sema3fa/olfm1a/unc5b/nptna/vegfc/dbn1/kctd12.1/si:ch211-129c21.1/phox2bb/ubap1/ctnna2/cntn4/cdh2/trarg1a/kalrnb/ulk2/ptk7a/dbnlb/cspg5b/efna1b/ephb3a/slc35b2/rbpms2b/fus/map1ab/dock7/foxb1a/nsfa/rab33ba/sipa1l1/tnika/nrp2a/pnpla6/kctd13/isl1a/fezf2/mink1/slit2/tenm3/rufy3/nfasca/hecw1b/sim1a/adcy8/pak1/spg11/ret/kif1b/arnt2/sema3bl/zdhhc17/plxna4/appa/nedd4l/arhgap35b/efna2a/dscama/apc/robo2/pcdh17/casp9/epha4a/hdc/anos1a/trioa/lhx9/epha7/rtn4r/cyfip2/dpysl3/draxinb/slit1b/robo3/usp33/map1sb/appb/tanc2a/dzip1/nyap2b/sema5a/kifbp/chad/pacsin1a/robo1/tnr/tenm4/hecw2a/epha4l/kif5c/slitrk4/islr2/nptnb/lhx2b/myo9ab/caprin2/kalrna/cdkl5/ephb2b/epha4b/clstn1/rac3a/brsk1b/aplp1/rab3ab/gap43/lrrc4ca/srrm4/lhx1a/map1aa/prmt8b/atl1/nkx2.2a/arhgef25b/dscamb/triob/tmem59l/myo10l3/gas7a/nsmfb/slit3/dpysl5a/chst11/fzd3a/kif5aa/robo4/efnb3b/vax1/stat3/dock10/ctnnd2b/zgc:77784/sema4c/si:ch211-113g11.6/rac1l/elavl4/myo10l1/ntn1b/scn8aa/lrrc4cb/plppr4a/grna/bdnf/ctnnd2a/nrn1la/slitrk2/nkx6.1/cspg5a/ncam1b/cntn3a.1/nrxn2a/dagla/efna3a/dpysl5b/inab/thsd7aa/si:ch211-241e1.3/plxna3/l1camb/plxna1a/nrcama/sema6ba/sema3b |
| GO:0048812 | neuron projection morphogenesis | -2.04924995520432 | 1.14653846153846e-08 | plxnb1b/tmem108/pik3cd/mycbp2/rb1/ptprfb/zgc:100846/ptprfa/spag6/isl2b/nrp2b/arhgap35a/rapgef1a/emb/pik3ca/rbpms2a/aplp2/mapk8ip3/npr1a/sema3fa/olfm1a/unc5b/nptna/vegfc/dbn1/kctd12.1/si:ch211-129c21.1/phox2bb/ubap1/ctnna2/cntn4/cdh2/trarg1a/kalrnb/ulk2/ptk7a/dbnlb/efna1b/ephb3a/slc35b2/rbpms2b/fus/map1ab/dock7/foxb1a/nsfa/rab33ba/sipa1l1/tnika/nrp2a/pnpla6/kctd13/isl1a/fezf2/mink1/slit2/tenm3/rufy3/nfasca/hecw1b/sim1a/adcy8/pak1/spg11/ret/kif1b/arnt2/sema3bl/zdhhc17/plxna4/appa/nedd4l/arhgap35b/efna2a/dscama/apc/robo2/pcdh17/casp9/epha4a/hdc/anos1a/trioa/lhx9/epha7/rtn4r/cyfip2/dpysl3/draxinb/slit1b/robo3/usp33/map1sb/appb/tanc2a/dzip1/nyap2b/sema5a/kifbp/chad/pacsin1a/robo1/tnr/tenm4/hecw2a/epha4l/kif5c/slitrk4/islr2/nptnb/lhx2b/myo9ab/caprin2/kalrna/cdkl5/ephb2b/epha4b/clstn1/rac3a/brsk1b/aplp1/rab3ab/gap43/lrrc4ca/srrm4/lhx1a/map1aa/prmt8b/atl1/nkx2.2a/arhgef25b/dscamb/triob/tmem59l/myo10l3/gas7a/nsmfb/slit3/dpysl5a/chst11/fzd3a/kif5aa/robo4/efnb3b/vax1/stat3/dock10/ctnnd2b/zgc:77784/sema4c/si:ch211-113g11.6/rac1l/elavl4/myo10l1/ntn1b/scn8aa/lrrc4cb/plppr4a/grna/bdnf/ctnnd2a/nrn1la/slitrk2/nkx6.1/ncam1b/cntn3a.1/nrxn2a/dagla/efna3a/dpysl5b/inab/thsd7aa/si:ch211-241e1.3/plxna3/l1camb/plxna1a/nrcama/sema6ba/sema3b |
| GO:0120039 | plasma membrane bounded cell projection morphogenesis | -2.0490209096944 | 1.14653846153846e-08 | plxnb1b/tmem108/pik3cd/mycbp2/rb1/ptprfb/zgc:100846/ptprfa/spag6/isl2b/nrp2b/arhgap35a/rapgef1a/emb/pik3ca/rbpms2a/aplp2/mapk8ip3/npr1a/sema3fa/olfm1a/unc5b/nptna/vegfc/dbn1/kctd12.1/si:ch211-129c21.1/phox2bb/ubap1/ctnna2/cntn4/cdh2/trarg1a/kalrnb/ulk2/ptk7a/dbnlb/efna1b/ephb3a/slc35b2/rbpms2b/fus/map1ab/dock7/foxb1a/nsfa/rab33ba/sipa1l1/tnika/nrp2a/pnpla6/kctd13/isl1a/fezf2/mink1/slit2/tenm3/rufy3/nfasca/hecw1b/sim1a/adcy8/pak1/spg11/ret/kif1b/arnt2/sema3bl/zdhhc17/plxna4/appa/nedd4l/arhgap35b/efna2a/dscama/apc/robo2/pcdh17/casp9/epha4a/hdc/anos1a/trioa/lhx9/epha7/rtn4r/cyfip2/dpysl3/draxinb/slit1b/robo3/usp33/map1sb/appb/tanc2a/dzip1/nyap2b/sema5a/kifbp/chad/pacsin1a/robo1/tnr/tenm4/hecw2a/epha4l/kif5c/slitrk4/islr2/nptnb/lhx2b/myo9ab/caprin2/kalrna/cdkl5/ephb2b/epha4b/clstn1/rac3a/brsk1b/aplp1/rab3ab/gap43/lrrc4ca/srrm4/lhx1a/map1aa/prmt8b/atl1/nkx2.2a/arhgef25b/dscamb/triob/tmem59l/myo10l3/gas7a/nsmfb/slit3/dpysl5a/chst11/fzd3a/kif5aa/robo4/efnb3b/vax1/stat3/dock10/ctnnd2b/zgc:77784/sema4c/si:ch211-113g11.6/rac1l/elavl4/myo10l1/ntn1b/scn8aa/lrrc4cb/plppr4a/grna/bdnf/ctnnd2a/nrn1la/slitrk2/nkx6.1/ncam1b/cntn3a.1/nrxn2a/dagla/efna3a/dpysl5b/inab/thsd7aa/si:ch211-241e1.3/plxna3/l1camb/plxna1a/nrcama/sema6ba/sema3b |
| GO:0032990 | cell part morphogenesis | -2.03560178882976 | 1.14653846153846e-08 | plxnb1b/tmem108/pik3cd/mycbp2/rb1/ptprfb/zgc:100846/ptprfa/spag6/isl2b/nrp2b/arhgap35a/rapgef1a/emb/pik3ca/rbpms2a/aplp2/mapk8ip3/npr1a/sema3fa/olfm1a/unc5b/nptna/vegfc/dbn1/kctd12.1/si:ch211-129c21.1/phox2bb/ubap1/ctnna2/cntn4/cdh2/trarg1a/kalrnb/ulk2/ptk7a/dbnlb/supv3l1/cspg5b/efna1b/ephb3a/slc35b2/rbpms2b/fus/map1ab/dock7/foxb1a/nsfa/rab33ba/sipa1l1/tnika/nrp2a/pnpla6/kctd13/isl1a/fezf2/mink1/slit2/tenm3/rufy3/nfasca/hecw1b/sim1a/adcy8/pak1/spg11/ret/kif1b/arnt2/sema3bl/zdhhc17/plxna4/appa/nedd4l/arhgap35b/efna2a/dscama/apc/robo2/pcdh17/casp9/epha4a/hdc/anos1a/trioa/lhx9/epha7/rtn4r/cyfip2/dpysl3/draxinb/slit1b/robo3/usp33/map1sb/appb/tanc2a/dzip1/nyap2b/sema5a/kifbp/chad/pacsin1a/robo1/tnr/tenm4/hecw2a/epha4l/kif5c/slitrk4/islr2/nptnb/lhx2b/myo9ab/caprin2/kalrna/cdkl5/ephb2b/epha4b/clstn1/rac3a/brsk1b/aplp1/rab3ab/gap43/lrrc4ca/srrm4/lhx1a/map1aa/prmt8b/atl1/nkx2.2a/arhgef25b/dscamb/triob/tmem59l/myo10l3/gas7a/nsmfb/slit3/dpysl5a/chst11/fzd3a/kif5aa/robo4/efnb3b/vax1/stat3/dock10/ctnnd2b/zgc:77784/sema4c/si:ch211-113g11.6/rac1l/elavl4/myo10l1/ntn1b/scn8aa/lrrc4cb/plppr4a/grna/bdnf/ctnnd2a/nrn1la/slitrk2/nkx6.1/cspg5a/ncam1b/cntn3a.1/nrxn2a/dagla/efna3a/dpysl5b/inab/thsd7aa/si:ch211-241e1.3/plxna3/l1camb/plxna1a/nrcama/sema6ba/sema3b |
| GO:0000904 | cell morphogenesis involved in differentiation | -2.02983448887318 | 1.14653846153846e-08 | pik3cd/mycbp2/rb1/ptprfb/adgrg1/zgc:100846/ptprfa/isl2b/nrp2b/arhgap35a/rapgef1a/whrna/emb/pik3ca/erbb2/rbpms2a/aplp2/mapk8ip3/npr1a/sema3fa/emx2/olfm1a/unc5b/nptna/vegfc/dbn1/kctd12.1/si:ch211-129c21.1/phox2bb/unk/ubap1/ctnna2/cntn4/cdh2/trarg1a/kalrnb/ulk2/ptk7a/dbnlb/efna1b/rilpl2/ephb3a/slc35b2/rbpms2b/fus/map1ab/dock7/foxb1a/nsfa/rab33ba/sipa1l1/nrp2a/ush1ga/pnpla6/kctd13/isl1a/fezf2/slit2/tenm3/rufy3/nfasca/hecw1b/sim1a/adcy8/pak1/spg11/ret/kif1b/arnt2/sema3bl/zdhhc17/plxna4/appa/nedd4l/arhgap35b/efna2a/dscama/apc/robo2/pcdh17/casp9/epha4a/hdc/anos1a/trioa/lhx9/epha7/rtn4r/cyfip2/dpysl3/draxinb/slit1b/robo3/usp33/map1sb/appb/tanc2a/dzip1/sema5a/kifbp/chad/robo1/tnr/whrnb/tenm4/hecw2a/epha4l/kif5c/slitrk4/islr2/nptnb/lhx2b/tet3/myo9ab/caprin2/kalrna/cdkl5/ephb2b/epha4b/clstn1/rac3a/brsk1b/aplp1/rab3ab/gap43/lrrc4ca/srrm4/lhx1a/map1aa/prmt8b/atl1/nkx2.2a/arhgef25b/dscamb/triob/tmem59l/myo10l3/nsmfb/slit3/cxcl8a/dpysl5a/chst11/fzd3a/kif5aa/robo4/efnb3b/vax1/stat3/dock10/ctnnd2b/zgc:77784/sema4c/si:ch211-113g11.6/rac1l/elavl4/myo10l1/ntn1b/scn8aa/lrrc4cb/plppr4a/grna/bdnf/ctnnd2a/slitrk2/nkx6.1/ncam1b/cntn3a.1/nrxn2a/dagla/grxcr1a/efna3a/dpysl5b/thsd7aa/si:ch211-241e1.3/plxna3/l1camb/plxna1a/nrcama/sema6ba/sema3b |
| GO:0007409 | axonogenesis | -2.02083525635161 | 1.14653846153846e-08 | plxnb1b/pik3cd/mycbp2/rb1/ptprfb/zgc:100846/ptprfa/isl2b/nrp2b/arhgap35a/rapgef1a/emb/pik3ca/rbpms2a/aplp2/mapk8ip3/npr1a/sema3fa/olfm1a/unc5b/nptna/vegfc/dbn1/si:ch211-129c21.1/phox2bb/ubap1/ctnna2/cntn4/cdh2/trarg1a/kalrnb/ulk2/ptk7a/dbnlb/efna1b/ephb3a/slc35b2/rbpms2b/fus/map1ab/dock7/foxb1a/nsfa/rab33ba/nrp2a/pnpla6/kctd13/isl1a/fezf2/slit2/tenm3/rufy3/nfasca/sim1a/adcy8/pak1/spg11/ret/kif1b/arnt2/sema3bl/zdhhc17/plxna4/appa/arhgap35b/efna2a/dscama/apc/robo2/pcdh17/casp9/epha4a/hdc/anos1a/trioa/lhx9/epha7/rtn4r/cyfip2/dpysl3/draxinb/slit1b/robo3/usp33/map1sb/appb/dzip1/sema5a/kifbp/chad/robo1/tnr/tenm4/epha4l/kif5c/slitrk4/islr2/nptnb/lhx2b/myo9ab/kalrna/cdkl5/ephb2b/epha4b/clstn1/rac3a/brsk1b/aplp1/rab3ab/gap43/lrrc4ca/srrm4/lhx1a/map1aa/atl1/nkx2.2a/arhgef25b/dscamb/triob/tmem59l/myo10l3/nsmfb/slit3/dpysl5a/chst11/fzd3a/kif5aa/robo4/efnb3b/vax1/stat3/zgc:77784/sema4c/si:ch211-113g11.6/rac1l/elavl4/myo10l1/ntn1b/scn8aa/lrrc4cb/plppr4a/grna/bdnf/slitrk2/nkx6.1/ncam1b/cntn3a.1/nrxn2a/dagla/efna3a/dpysl5b/thsd7aa/si:ch211-241e1.3/plxna3/l1camb/plxna1a/nrcama/sema6ba/sema3b |
| GO:0001505 | regulation of neurotransmitter levels | -2.01367130950093 | 0.000141537068751352 | snap25a/napbb/slc5a7a/napba/chata/stxbp1b/cplx2a/comtb/si:ch211-81a5.8/dbh/syn1/stx3a/caskb/snap25b/mctp2b/snap47/syt7b/syn2a/rims1b/gad1b/syt4/rab3ab/moxd1/cadpsb/syt11a/unc13a/stxbp1a/syt1a/ppfia3/gad2/syt2a/baiap3/hrh3/rph3ab/th2/stx11a |
| GO:0061564 | axon development | -1.97936421112892 | 1.14653846153846e-08 | plxnb1b/pik3cd/mycbp2/rb1/ptprfb/zgc:100846/plp1b/ptprfa/isl2b/nrp2b/arhgap35a/rapgef1a/emb/pik3ca/erbb2/rbpms2a/aplp2/mapk8ip3/npr1a/sema3fa/olfm1a/unc5b/nptna/vegfc/dbn1/gpm6bb/si:ch211-129c21.1/phox2bb/ubap1/ctnna2/cntn4/cdh2/trarg1a/kalrnb/ulk2/ptk7a/dbnlb/nfia/efna1b/ephb3a/slc35b2/rbpms2b/fus/map1ab/dock7/foxb1a/nsfa/rab33ba/nrp2a/pnpla6/kctd13/isl1a/fezf2/gpm6ba/slit2/tenm3/rufy3/nfasca/sim1a/adcy8/pak1/spg11/ret/kif1b/arnt2/sema3bl/zdhhc17/plxna4/appa/arhgap35b/efna2a/dscama/apc/robo2/pcdh17/casp9/epha4a/hdc/anos1a/trioa/lhx9/epha7/rtn4r/cyfip2/dpysl3/draxinb/slit1b/robo3/usp33/map1sb/appb/dzip1/sema5a/kifbp/chad/il6st/robo1/tnr/tenm4/epha4l/kif5c/slitrk4/islr2/nptnb/lhx2b/myo9ab/kalrna/cdkl5/ephb2b/epha4b/ek1/clstn1/rac3a/brsk1b/aplp1/rab3ab/gap43/lrrc4ca/srrm4/lhx1a/map1aa/atl1/nkx2.2a/arhgef25b/dscamb/triob/tmem59l/myo10l3/nsmfb/slit3/dpysl5a/sarm1/chst11/fzd3a/kif5aa/robo4/efnb3b/vax1/stat3/zgc:77784/sema4c/si:ch211-113g11.6/rac1l/elavl4/myo10l1/ntn1b/scn8aa/lrrc4cb/plppr4a/grna/bdnf/slitrk2/nkx6.1/ncam1b/cntn3a.1/nrxn2a/dagla/efna3a/dpysl5b/thsd7aa/si:ch211-241e1.3/plxna3/l1camb/plxna1a/nrcama/sema6ba/sema3b |
| GO:0051049 | regulation of transport | -1.97152747166447 | 1.14653846153846e-08 | prnprs3/gem/kcne4/kcna1a/unc45a/dmxl2/mffa/scn5lab/mcoln1a/pxk/abcg1/kcnj13/rab33ba/fgf13b/cep290/c2cd5/cldnb/abca1a/hecw1b/cacna1ia/kctd7/nedd4l/cplx2a/chp2/kcnab1b/si:cabz01076231.1/cacnb1/bmp2k/cacna1fb/mcu/si:ch211-204c21.1/fgf13a/msnb/kcng1/cacng5a/cacna1ha/si:ch211-81a5.8/stim2b/cnstb/syt6b/stxbp5b/c2cd2l/caskb/fxyd6/crhbp/scn1ba/kcnab2a/pacsin1a/mctp2b/fgf12a/plcb3/rab3c/htr1aa/ank3a/dnajc5aa/hecw2a/ptprn2/rab3da/cacnb2b/syt7b/cacng7b/rab3b/rims1b/shisa9a/syt9b/zgc:91999/syt4/rab3ab/kcnj3b/kcnc2/kcnj19a/sgip1b/cadpsb/scn3b/rrad/syt13/stxbp5a/syt11a/cacng2b/kcnj1a.1/syt1a/cacng3b/shisa7b/syt9a/syt10/ptprna/kcnc1b/stx1b/cacna1da/stxbp4/anxa1c/scn8aa/cacng2a/rab44/stc1l/cacna1c/cacnb4b/fgf11b/cacna2d3/kcnj14/kcnj19b/syt2a/nkx6.1/kcnd3/il1rapl1a/syt3/cabp2a/baiap3/cacna1ab/kcnc1a/rph3ab/shisa9b/kcnd2/cnih2/npvf/kcnab2b/trh/cacng8a/cacna2d2b/scn1lab/kcnj5/syt6a/cacna2d4b/kcnh4a/kcnb1 |
| GO:0032879 | regulation of localization | -1.91598630922618 | 1.14653846153846e-08 | dtnbp1b/scn4ba/cplx2b/plcg1/gpr137ba/cacng8b/ptprnb/hip1rb/kcnh7/efcab7/scn8ab/aplnrb/adcyap1b/prnprs3/gem/kcne4/kcna1a/unc45a/letm1/dmxl2/mffa/scn5lab/mcoln1a/pxk/abcg1/kcnj13/rab33ba/fgf13b/cep290/c2cd5/cldnb/abca1a/hecw1b/cacna1ia/kctd7/nedd4l/cplx2a/chp2/plin2/kcnab1b/si:cabz01076231.1/cacnb1/bmp2k/cacna1fb/mcu/si:ch211-204c21.1/fgf13a/msnb/kcng1/plch1/cacng5a/cacna1ha/si:ch211-81a5.8/stim2b/cnstb/syt6b/stxbp5b/c2cd2l/caskb/fxyd6/crhbp/scn1ba/kcnab2a/pacsin1a/mctp2b/fgf12a/plcb3/rab3c/htr1aa/ank3a/dnajc5aa/hecw2a/wnt5b/ptprn2/rab3da/cacnb2b/plce1/plch2a/syt7b/cacng7b/rab3b/rims1b/shisa9a/syt9b/zgc:91999/syt4/gpc6a/rab3ab/kcnj3b/kcnc2/kcnj19a/sgip1b/zgc:77486/cadpsb/scn3b/rrad/syt13/stxbp5a/syt11a/cacng2b/kcnj1a.1/syt1a/gpc5b/cacng3b/shisa7b/syt9a/syt10/ptprna/kcnc1b/stx1b/cacna1da/stxbp4/anxa1c/scn8aa/cacng2a/rab44/stc1l/cacna1c/cacnb4b/fgf11b/cacna2d3/kcnj14/kcnj19b/syt2a/nkx6.1/kcnd3/il1rapl1a/syt3/cabp2a/baiap3/cacna1ab/kcnc1a/rph3ab/shisa9b/kcnd2/si:ch211-13f8.1/cnih2/npvf/kcnab2b/trh/cacng8a/cacna2d2b/scn1lab/kcnj5/syt6a/cacna2d4b/kcnh4a/kcnb1 |
| GO:1901701 | cellular response to oxygen-containing compound | -1.89692649413327 | 1.56824905636127e-05 | zgc:172120/pik3r3b/myd88/carf/stat6/nr4a2b/cpeb4a/chrng/glra1/syt6b/il13/c2cd2l/phip/htr1aa/ptger4c/ticam1/htr1d/ptprn2/tet3/gnao1b/inpp5ka/syt7b/syt9b/syt4/insig1/esr2b/ptger4a/cpne4a/vtcn1/cxcl8a/cxcl8b.3/nr4a2a/syt13/syt11a/stat3/nsg2/syt1a/sstr1a/syt9a/syt10/gnao1a/ptprna/cpne2/stxbp4/adgrv1/anxa1c/nr4a3/inpp5kb/syt2a/syt3/cpne7/irs2b/drd2a/htr1fa/chrm4a/gnb5a/cpne8/syt6a/il1b/gper1/cxcl18b |
| GO:0048640 | negative regulation of developmental growth | -1.88113815885922 | 0.0105693423015173 | sema6d/sema6e/dusp6/sema3fa/si:ch211-129c21.1/ulk2/slit2/sema3bl/robo2/epha4a/wwc1/sema5a/epha4b/slit3/sema4c/si:ch211-113g11.6/wwc3/sema6ba/sema3b |
| GO:0007019 | microtubule depolymerization | -1.8750187848337 | 0.021391144294225 | map1ab/stmn4/apc/map1b/stmn2a/map1sb/stmn2b/stmn3/map1aa/camsap1a/stmn4l/apc2 |
| GO:0032409 | regulation of transporter activity | -1.86261008963063 | 0.00378678429698509 | cabp1a/cabp5a/scn4ba/cacng8b/prnprs3/gem/kcne4/fgf13b/hecw1b/kctd7/nedd4l/chp2/kcnab1b/si:cabz01076231.1/cacnb1/fgf13a/kcng1/cacng5a/stim2b/fxyd6/fgf12a/ank3a/hecw2a/cacnb2b/cacng7b/shisa9a/rrad/cacng2b/cacng3b/shisa7b/cacng2a/cacnb4b/fgf11b/cabp2a/shisa9b/cnih2/cacng8a |
| GO:0003310 | pancreatic A cell differentiation | -1.86173412990953 | 0.00748525766259099 | isl1a/pax6b/insm1b/nkx6.2/myt1b/nkx6.1/arxa |
| GO:0036269 | swimming behavior | -1.85563104344825 | 0.0118323095206783 | dmrt3a/myo9aa/sin3b/panx1b/kif2a/gdpd5a/zgc:100846/ca10a/klhl41a/spg11/scdb/micall2b/ca10b/myo9ab/cdkl5/grna/tcap/cacna1ab/dagla/syngap1b/gnb5a/scn1lab/dnmt3aa/oxt |
| GO:0019226 | transmission of nerve impulse | -1.84484111835001 | 0.0190096470355517 | cacng8b/scn8ab/cacna1ia/cacng5a/cacna1ha/ank3a/cacng7b/cacng2b/cacng3b/scn8aa/cacng2a/kcnab2b/cacng8a/scn1lab |
| GO:0021545 | cranial nerve development | -1.80828563846919 | 0.00979204145463202 | rb1/adgrg1/klf6a/hgfa/mib1/erbb2/nes/tuba1a/nsfa/six3a/slit2/kif1b/chn1/robo2/klf7b/slit1b/six3b/kifbp/diras1a/lhx2b/vax1/mdga2a/cdh4/foxi1 |
| GO:0007186 | G protein-coupled receptor signaling pathway | -1.79358601966978 | 1.60681800279223e-06 | penka/gpr27/rgs18/trpm1b/gpr78a/cart2/opn1mw2/gnb1a/gpr182/npy/chrm2a/rgrb/penkb/gng3/glrbb/adcy2a/gal/adcy6b/adcy8/adgre15/ptgir/f2rl2/rgra/grm5a/gpr22a/gnal/lpar1/or95a1/gpr35.1/s1pr1/glra1/ccl38.6/gnai2b/tmem116/drd3/oprd1b/aplnr2/gpr22b/usp33/grm4/grm6b/npffl/lmbrd2b/rgs7a/ccl34b.4/grm8b/cart3/gpr174/htr1aa/ptger4c/tafa5a/gpr61l/adgrf3b/pde6ga/htr1d/pdyn/gnb2/si:ch211-117l17.6/gnao1b/adgra2/xcr1a.1/gnb3a/ccl44/gnb4b/agtr2/rgs9b/mrap2a/gpr63/ptger4a/adra2da/opn6b/prlhr2a/adcy6a/nmur3/adcy2b/adgrl3.1/adcy1a/si:dkey-6n21.13/grin1a/grm2a/tas1r2.1/sstr1a/gabbr1b/pth2/adra2b/trpm1a/git2b/tmtops2b/gnao1a/gpr85/gpr158b/adgrv1/ccl35.1/si:dkey-172j4.3/adcy5/grm8a/tshba/adgra1a/tafa5b/grm2b/prlhr2b/gpr75/vipr1b/adgrb3/pnocb/adgre5b.2/hrh3/adrb1/rgs11/drd2a/dagla/adgrb1a/htr1fa/npb/si:ch211-132p1.2/kctd8/chrm4a/ltb4r2a/crhr1/gpr158a/npvf/ltb4r/grm1a/adcyap1r1a/pomca/adgrg3/gnb5a/rxfp1/tac3b/htr7c/adgrl1a/adgrb1b/gper1/or92a7/p2ry13 |
| GO:0010721 | negative regulation of cell development | -1.7837780436552 | 0.00979204145463202 | sema3fa/gpr137ba/si:ch211-129c21.1/irf2bp2b/spi1b/ulk2/slit2/sema3bl/tmem178a/robo2/epha4a/bmp15/sema5a/epha4b/slit3/sema4c/si:ch211-113g11.6/anxa1c/mmp9/sema6ba/gper1/sema3b |
| GO:0006935 | chemotaxis | -1.78340911781747 | 1.1812068993636e-05 | unc5b/nptna/vegfc/tnfrsf1a/si:ch211-129c21.1/cntn4/cdh2/kalrnb/maco1a/tdgf1/ptk7a/efna1b/ephb3a/slc35b2/rbpms2b/rab33ba/nrp2a/fezf2/slit2/tenm3/maco1b/nfasca/sim1a/adcy8/kif1b/arnt2/sema3bl/myd88/plxna4/efna2a/dscama/apc/robo2/pcdh17/hdc/anos1a/trioa/s1pr1/lhx9/epha7/ccl38.6/rtn4r/cyfip2/dpysl3/draxinb/slit1b/robo3/usp33/aanat2/dock4b/dzip1/trim33/ccl34b.4/sema5a/chad/vegfd/dock4/robo1/tnr/tenm4/epha4l/kif5c/islr2/nptnb/lhx2b/kalrna/xcr1a.1/ccl44/ephb2b/rac3a/gap43/lhx1a/atl1/nkx2.2a/arhgef25b/dscamb/triob/slit3/cxcl8a/dpysl5a/cxcl8b.3/chst11/fzd3a/kif5aa/si:dkey-12h9.6/robo4/efnb3b/vax1/stat3/zgc:77784/sema4c/si:ch211-113g11.6/rac1l/anxa1c/ntn1b/scn8aa/ccl35.1/cxcl8b.1/ncam1b/cntn3a.1/nrxn2a/efna3a/dpysl5b/si:ch211-241e1.3/plxna3/l1camb/plxna1a/nrcama/sema6ba/il1b/sema3b/cxcl18b |
| GO:0042330 | taxis | -1.78340911781747 | 1.1812068993636e-05 | unc5b/nptna/vegfc/tnfrsf1a/si:ch211-129c21.1/cntn4/cdh2/kalrnb/maco1a/tdgf1/ptk7a/efna1b/ephb3a/slc35b2/rbpms2b/rab33ba/nrp2a/fezf2/slit2/tenm3/maco1b/nfasca/sim1a/adcy8/kif1b/arnt2/sema3bl/myd88/plxna4/efna2a/dscama/apc/robo2/pcdh17/hdc/anos1a/trioa/s1pr1/lhx9/epha7/ccl38.6/rtn4r/cyfip2/dpysl3/draxinb/slit1b/robo3/usp33/aanat2/dock4b/dzip1/trim33/ccl34b.4/sema5a/chad/vegfd/dock4/robo1/tnr/tenm4/epha4l/kif5c/islr2/nptnb/lhx2b/kalrna/xcr1a.1/ccl44/ephb2b/rac3a/gap43/lhx1a/atl1/nkx2.2a/arhgef25b/dscamb/triob/slit3/cxcl8a/dpysl5a/cxcl8b.3/chst11/fzd3a/kif5aa/si:dkey-12h9.6/robo4/efnb3b/vax1/stat3/zgc:77784/sema4c/si:ch211-113g11.6/rac1l/anxa1c/ntn1b/scn8aa/ccl35.1/cxcl8b.1/ncam1b/cntn3a.1/nrxn2a/efna3a/dpysl5b/si:ch211-241e1.3/plxna3/l1camb/plxna1a/nrcama/sema6ba/il1b/sema3b/cxcl18b |
| GO:0098953 | receptor diffusion trapping | -1.77120547755989 | 0.0221781634063151 | cacng8b/gphnb/cacng5a/gphna/cacng7b/cacng2b/cacng3b/cacng2a/cacng8a |
| GO:0098970 | postsynaptic neurotransmitter receptor diffusion trapping | -1.77120547755989 | 0.0221781634063151 | cacng8b/gphnb/cacng5a/gphna/cacng7b/cacng2b/cacng3b/cacng2a/cacng8a |
| GO:0099628 | neurotransmitter receptor diffusion trapping | -1.77120547755989 | 0.0221781634063151 | cacng8b/gphnb/cacng5a/gphna/cacng7b/cacng2b/cacng3b/cacng2a/cacng8a |
| GO:0071867 | response to monoamine | -1.76002500271274 | 0.0460556112254136 | htr1aa/htr1d/gnao1b/nsg2/gnao1a/drd2a/htr1fa/gnb5a |
| GO:0071869 | response to catecholamine | -1.76002500271274 | 0.0460556112254136 | htr1aa/htr1d/gnao1b/nsg2/gnao1a/drd2a/htr1fa/gnb5a |
| GO:0051960 | regulation of nervous system development | -1.75557639124564 | 0.00248666027456893 | nlk1/si:zfos-741a10.3/plxnb1b/hes6/neurod6a/adnpb/sema3fa/btbd6a/her12/dbn1/si:ch211-129c21.1/lef1/ulk2/dbnlb/ephb3a/dhrs12la/slit2/ctbp2a/sema3bl/plxna4/clstn2b/bhlhe40/robo2/epha4a/neurod4/gata3/her13/gbx2/tnfrsf21/sema5a/foxo3a/tnr/tenm4/slitrk4/caprin2/zbtb16a/cdkl5/epha4b/clstn1/nsmfb/slit3/clstn2a/elavl3/stat3/sema4c/si:ch211-113g11.6/gbx1/grna/bdnf/clstn3/slitrk2/dmbx1a/plxna3/l1camb/mmp9/plxna1a/sema6ba/sema3b |
| GO:0097120 | receptor localization to synapse | -1.75544036908873 | 0.0351674145187552 | dlg1b/cacng8b/grip2b/gphnb/cacng5a/stx3a/anks1b/kif5c/gphna/cacng7b/kif5aa/cacng2b/cacng3b/stx1b/cacng2a/rab44/cacng8a |
| GO:0031532 | actin cytoskeleton reorganization | -1.74088656656218 | 0.0402190977319739 | klf6a/anxa1a/baiap2a/sipa1l1/mink1/nedd9/thsd7bb/carmil2/parvg/carmil3/plek/thsd7ba/anxa1c/plek2/phactr1/thsd7aa |
| GO:0040013 | negative regulation of locomotion | -1.73929262815316 | 0.018759055936431 | sema3fa/lrch1/si:ch211-129c21.1/lbx2/prox1a/miip/lef1/phf8/srgap3/sema3bl/robo2/anos1a/rap2b/sema5a/srgap1b/sinhcaf/arhgap4b/sema4c/si:ch211-113g11.6/rtn4rl2a/sema6ba/sema3b |
| GO:0031110 | regulation of microtubule polymerization or depolymerization | -1.72739295421889 | 0.0437181181328161 | ska3/cdk5rap2/stmn1b/map1sa/camsap2a/camsap1b/slain2/map1ab/traf3ip1/zgc:66447/clasp2/mapre3b/stmn4/apc/map1b/stmn2a/map1sb/stmn2b/stmn3/clip1a/map1aa/camsap1a/stmn4l/apc2 |
| GO:0040011 | locomotion | -1.71881331498307 | 4.74228301214316e-06 | plcg1/unc5b/nptna/lrch1/vegfc/tnfrsf1a/si:ch211-129c21.1/lbx2/cntn4/cdh2/prox1a/kalrnb/miip/maco1a/lef1/tdgf1/foxj1a/ptk7a/efna1b/ephb3a/slc35b2/gnb1a/rbpms2b/sst7/slkb/rab33ba/nrp2a/fezf2/phf8/slit2/tenm3/maco1b/srgap3/nfasca/sim1a/adcy8/kif1b/arnt2/sema3bl/myd88/plxna4/efna2a/dscama/apc/robo2/pcdh17/hdc/asap1a/arhgef39/anos1a/trioa/s1pr1/lhx9/epha7/hace1/peak1/ccl38.6/rtn4r/cyfip2/dpysl3/draxinb/gata3/slit1b/robo3/usp33/tek/aanat2/dock4b/dzip1/rap2b/trim33/ccl34b.4/sema5a/chad/vegfd/pdgfbb/dock4/robo1/tnr/tenm4/epha4l/kif5c/islr2/nptnb/htr1d/lhx2b/kalrna/xcr1a.1/ccl44/ephb2b/sst1.1/srgap1b/rac3a/gap43/lhx1a/atl1/nkx2.2a/arhgef25b/dscamb/amot/triob/nsmfb/slit3/cxcl8a/dpysl5a/cxcl8b.3/chst11/fzd3a/kif5aa/si:dkey-12h9.6/sinhcaf/robo4/efnb3b/vax1/arhgap4b/stat3/dock10/zgc:77784/sema4c/adam8a/si:ch211-113g11.6/rac1l/anxa1c/ntn1b/scn8aa/ccl35.1/cxcl8b.1/mdga2a/zgc:153631/asap1b/ncam1b/cntn3a.1/nrxn2a/klhl40b/efna3a/chrm4a/dpysl5b/si:ch211-241e1.3/plxna3/l1camb/plxna1a/nrcama/rtn4rl2a/sema6ba/il1b/sema3b/cxcl18b |
| GO:0006959 | humoral immune response | -1.69323952530372 | 0.0594675488145613 | c4/tnfrsf21/c1qa/tfe3a/cfb/cxcl8b.3/c6/susd4/c7b/c4b/c3a.2/itln2/cxcl18b |
| GO:0007420 | brain development | -1.67227058783588 | 2.31796694375566e-05 | her12/wnt3/macroh2a2/marcksa/kctd12.1/daam1a/pknox1.1/tgif1/ctnna2/cntn4/cdh2/adcyap1b/prox1a/zfhx3b/prnprs3/sbno2a/lef1/tdgf1/ulk2/tsc22d3/bag6l/pou3f3a/atg4da/creb1a/pafah1b1a/birc6-as2/sox14/foxb1a/nsfa/zfhx3a/kdm6al/slc2a3a/cep290/kctd13/fezf2/phf8/six3a/nfasca/sim1a/pax6b/arnt2/raraa/grhl3/tbr1b/kirrel3l/cars2/ptprsa/kdm7ab/nr4a2b/ralgapa1/apc/epha4a/hdc/anos1a/lhx9/neurod4/pcdh10b/pbx1a/mvp/six3b/mdkb/si:ch73-119p20.1/gbx2/dzip1/lgi1a/pbx1b/epb41l5/auts2a/pbx3b/pcare1/hecw2a/znf503/meis2a/wnt5b/lhx6a/irx7/lhx2b/brf1b/rbfox1/tcf7l2/nkx2.1/pcdh19/srrm4/lhx1a/asphd1/pou4f2/prmt8b/tbxta/aatkb/rtn1b/sp5l/nr2e3/tbcd/has2/lhx5/fzd3a/pou3f2b/nr4a2a/efnb3b/ndrg4/vax1/pax2a/git2b/gbx1/ntn1b/pou3f2a/bdnf/atp1a3a/sez6l2/mdga2a/rx3/dmbx1a/otpb/six6a/cdk5r2a/arxa/nkx2.4a/syngap1b/gnrh2/l1camb/cnnm2a/otpa/kcnb1/nrcama/gper1/bicdl1 |
| GO:0060322 | head development | -1.66225293057786 | 4.42238345198842e-05 | her12/wnt3/macroh2a2/marcksa/kctd12.1/daam1a/pknox1.1/tgif1/ctnna2/cldn7a/cntn4/cdh2/adcyap1b/prox1a/zfhx3b/prnprs3/sbno2a/lef1/tdgf1/ulk2/tsc22d3/bag6l/pou3f3a/atg4da/creb1a/pafah1b1a/birc6-as2/sox14/foxb1a/nsfa/zfhx3a/kdm6al/slc2a3a/cep290/kctd13/fezf2/phf8/six3a/nfasca/sim1a/pax6b/arnt2/raraa/grhl3/tbr1b/kirrel3l/cars2/ptprsa/kdm7ab/nr4a2b/ralgapa1/apc/epha4a/hdc/wdr11/anos1a/lhx9/neurod4/pcdh10b/pbx1a/mvp/six3b/mdkb/si:ch73-119p20.1/gbx2/dzip1/lgi1a/pbx1b/epb41l5/auts2a/pbx3b/pcare1/hecw2a/znf503/meis2a/wnt5b/lhx6a/irx7/lhx2b/brf1b/rbfox1/tcf7l2/nkx2.1/pcdh19/srrm4/lhx1a/asphd1/pou4f2/prmt8b/tbxta/aatkb/rtn1b/sp5l/nr2e3/tbcd/has2/lhx5/fzd3a/pou3f2b/nr4a2a/efnb3b/ndrg4/vax1/pax2a/git2b/gbx1/ntn1b/chd5/pou3f2a/bdnf/atp1a3a/sez6l2/mdga2a/specc1/rx3/dmbx1a/otpb/six6a/cdk5r2a/arxa/nkx2.4a/syngap1b/gnrh2/l1camb/cnnm2a/otpa/kcnb1/nrcama/gper1/bicdl1 |
| GO:0045732 | positive regulation of protein catabolic process | -1.66175262841073 | 0.0607582432020156 | oaz1a/csnk1db/tmem259/oaz2b/socs5b/prkn/apc/rnf217/si:ch211-278j3.3/trib1/fzr1b/rnf144ab/csnk1e/adam8a/klhl40b/apc2/rnf144aa |
| GO:1901797 | negative regulation of signal transduction by p53 class mediator | -1.65413798029984 | 0.0829144295829561 | cd74b/cd74a/bcor/vax1/bcl6aa |
| GO:0006688 | glycosphingolipid biosynthetic process | -1.64952029857802 | 0.097836707744085 | b4galt6/b4galt3/st6galnac3 |
| GO:0031644 | regulation of nervous system process | -1.64521543139582 | 0.0906720571933235 | dlgap1a/tnfrsf21/tenm4/dlgap4b/si:dkey-174m14.3/kcnab2b/dlgap3/neto1l |
| GO:0060560 | developmental growth involved in morphogenesis | -1.64279725335724 | 0.0128583805736124 | plxnb1b/tmem108/pik3cd/mycbp2/spag6/cbsb/smad2/pik3ca/zeb2a/mapk8ip3/sema3fa/dbn1/si:ch211-129c21.1/ubap1/cdh2/trarg1a/tmem67/ulk2/ptk7a/dbnlb/creb1a/fus/isl1a/slit2/ppp1cb/spg11/ret/kif1b/sema3bl/bbs1/robo2/pcdh17/casp9/epha4a/robo3/appb/sema5a/wnt5b/myo9ab/cdkl5/epha4b/clstn1/tbxta/zeb2b/tmem59l/myo10l3/nsmfb/slit3/stat3/sema4c/si:ch211-113g11.6/myo10l1/ntn1b/grna/bdnf/nrn1la/nkx6.1/nrxn2a/thsd7aa/plxna3/l1camb/sema6ba/sema3b |
| GO:0016049 | cell growth | -1.63666909504427 | 0.0141269580867656 | plxnb1b/tmem108/pik3cd/mycbp2/spag6/ewsr1a/pik3ca/mapk8ip3/sema3fa/dbn1/si:ch211-129c21.1/ubap1/trarg1a/ulk2/dbnlb/supv3l1/dnph1/fus/isl1a/slit2/spg11/ret/kif1b/sema3bl/robo2/pcdh17/casp9/epha4a/robo3/appb/sema5a/wnt5b/myo9ab/cdkl5/epha4b/clstn1/tmem59l/myo10l3/nsmfb/slit3/sema4c/si:ch211-113g11.6/myo10l1/ntn1b/epb41l3a/grna/bdnf/nrn1la/nkx6.1/nrxn2a/thsd7aa/plxna3/l1camb/sema6ba/sema3b |
| GO:0034643 | establishment of mitochondrion localization. microtubule-mediated | -1.63631487834659 | 0.0931198545493737 | rhot1b/trak1a/sybu/kif5aa/rhot2 |
| GO:0047497 | mitochondrion transport along microtubule | -1.63631487834659 | 0.0931198545493737 | rhot1b/trak1a/sybu/kif5aa/rhot2 |
| GO:0051654 | establishment of mitochondrion localization | -1.63631487834659 | 0.0931198545493737 | rhot1b/trak1a/sybu/kif5aa/rhot2 |
| GO:0032102 | negative regulation of response to external stimulus | -1.63570938930065 | 0.0540332674423811 | sema3bl/gramd4b/robo2/epha4a/il13/nr1h5/sema5a/cart3/ptger4c/irak3/epha4b/gramd4a/ptger4a/sema4c/si:ch211-113g11.6/susd4/grna/serpine1/sc:d189/sema6ba/sema3b |
| GO:0048885 | neuromast deposition | -1.63446228315307 | 0.0960340406478379 | lbx2/prox1a/lef1/phf8/anos1a/rtn4rl2a |
| GO:0071407 | cellular response to organic cyclic compound | -1.62675574144756 | 0.0543657683428525 | abhd2b/htr1aa/htr1d/gnao1b/insig1/esr2b/cxcl8a/nsg2/sstr1a/gnao1a/anxa1c/drd2a/htr1fa/gnb5a/gper1 |
| GO:0022603 | regulation of anatomical structure morphogenesis | -1.62242060545465 | 0.00196692398869123 | plcd4a/vegfab/stab1/hif1ab/amotl2b/sema6d/ptenb/alcama/sema6e/ptk2ab/thbs2b/brwd1/rspo3/tspan18a/gata2a/zeb1b/plxnb1b/pik3cd/disc1/mia3/anxa1a/fn1b/arhgap35a/pik3ca/flot2a/sema3fa/camsap1b/plcg1/unc5b/vegfc/dbn1/aplnrb/kctd12.1/si:ch211-129c21.1/rhogd/zeb1a/lbx2/cdh2/fmnl2b/ulk2/elmo1/marchf5/ptk7a/dbnlb/dab2ipb/lurap1/ephb3a/mffa/sipa1l1/tnika/vat1/palm1b/slit2/mief2/rufy3/hecw1b/pak1/sema3bl/plxna4/nedd4l/arhgap35b/robo2/casp9/epha4a/s1pr1/msnb/tanc2a/phip/sema5a/vegfd/tenm4/hecw2a/wnt5b/caprin2/cdkl5/epha4b/myo10/clstn1/rac3a/lrrc4ca/lhx1a/nsmfb/slit3/basp1/stat3/prkd1/sema4c/si:ch211-113g11.6/rac1l/gbx1/anxa1c/lrrc4cb/hif1aa/bdnf/serpine1/adgrb3/scg2b/gdf3/adgrb1a/thbs1a/dab2ipa/plxna3/l1camb/eve1/plxna1a/adgrb1b/sema6ba/sema3b |
| GO:0000381 | regulation of alternative mRNA splicing. via spliceosome | -1.62121505428978 | 0.0619371464051369 | hnrnpa0a/rbm38/rbm25a/celf2/rbfox3a/rbfox3b/celf6/rbfox1l/rbfox1/srrm4/nova2/celf3a/khdrbs2/celf5a/celf4 |
| GO:0001933 | negative regulation of protein phosphorylation | -1.61410605566215 | 0.0735990755145654 | spry4/aida/dtnbp1b/ptprc/ldlrad4a/camk2n2/cbl/camk2n1a/cblc/prkn/sh3bp5la/sh3bp5b/si:dkey-175m17.7/ibtk/camk2n1/irak3/inpp5ka/cadm4/inpp5kb/smcr8b |
| GO:0008361 | regulation of cell size | -1.59323427771673 | 0.0658674351521172 | arhgap35a/pik3ca/sema3fa/arhgap5/dbn1/si:ch211-129c21.1/ulk2/dbnlb/zgc:153039/slit2/sema3bl/arhgap35b/robo2/casp9/epha4a/sema5a/piezo1/cdkl5/epha4b/clstn1/nsmfb/slit3/si:ch211-178n15.1/stat3/slc12a5b/sema4c/si:ch211-113g11.6/ntn1b/slc12a5a/bdnf/nkx6.1/plxna3/l1camb/sema6ba/sema3b |
| GO:0043087 | regulation of GTPase activity | -1.57086875382947 | 0.0297635976408028 | tbc1d9/rapgef1a/flcn/sipa1l2/rasgef1ba/iqsec1b/ralgapa2/rasa4/gpr137ba/rgl1/nf1a/usp6nl/adcyap1b/vav2/agap1/rapgef5b/rapgef2a/si:dkey-91m11.5/dab2ipb/rgl3a/ephb3a/rapgef2b/dock7/rapgef4b/sipa1l1/gapvd1/rasal2/plxna4/chn1/ralgapa1/ccpg1/asap1a/rasgef1bb/ccl38.6/bcr/rgs7a/ccl34b.4/si:ch211-288d18.1/evi5l/tbc1d30/elmod1/vav3b/myo9ab/ccl44/ek1/rgs9b/sgsm3/syngap1a/arap3/rap1gapa/unm_sa1614/ralgps1/dock10/agap2/ccl35.1/asap1b/rasgrf2b/rgs11/syngap1b/dab2ipa/plxna3/rasgrf1/plxna1a/arhgap25 |
| GO:0002252 | immune effector process | -1.52782636998837 | 0.0607582432020156 | cd74b/c4/c1qa/ticam1/cd74a/cfb/cadm1b/cxcl8a/c6/si:ch211-196f2.6/susd4/anxa1c/zgc:153631/c7b/c4b/zgc:123107/c3a.2/csf2rb/mmp9/mhc2d4b |
| GO:0001755 | neural crest cell migration | -1.51280994298295 | 0.0960340406478379 | hif1ab/sema6d/pax6a/med13b/sema6e/lrp5/disc1/nrp2b/erbb2/pdgfrb/sema3fa/radil/si:ch211-129c21.1/cdh2/lef1/dmxl2/pax3a/nrp2a/fscn1a/pak1/ret/pax6b/sema3bl/mmp17b/wdr11/sema5a/zeb2b/sh3pxd2aa/sema4c/si:ch211-113g11.6/rx3/foxi1/sema6ba/sema3b |
| GO:0051240 | positive regulation of multicellular organismal process | -1.44230693231483 | 0.0657742225763037 | gata2a/slc27a1b/nlk1/si:zfos-741a10.3/plxnb1b/anxa1a/avpr1ab/neurod6a/tnfsf13b/nr2c2/cactin/apln/tlr5b/olfm1a/vegfc/dbn1/aplnrb/lef1/elmo1/dbnlb/lurap1/ephb3a/npy/tapt1b/dhrs12la/loxl2a/zgc:172120/plxna4/clstn2b/cd74b/neurod4/gata3/aanat2/vegfd/enpp5/tenm4/ticam1/slitrk4/caprin2/zbtb16a/cdkl5/clstn1/cd74a/lhx1a/vtcn1/nsmfb/clstn2a/basp1/stat3/si:ch211-196f2.6/anxa1c/grna/bdnf/zbtb16b/clstn3/slitrk2/zgc:153631/adrb1/npvf/plxna3/l1camb/zgc:123107/plxna1a/panx2/gper1/mhc2d4b |
| GO:0030334 | regulation of cell migration | -1.43156853115565 | 0.0696903189146766 | plcg1/lrch1/vegfc/si:ch211-129c21.1/lbx2/prox1a/miip/lef1/gnb1a/sst7/slkb/phf8/srgap3/sema3bl/plxna4/asap1a/arhgef39/anos1a/s1pr1/hace1/peak1/dpysl3/gata3/tek/rap2b/sema5a/vegfd/pdgfbb/sst1.1/srgap1b/amot/nsmfb/sinhcaf/robo4/arhgap4b/dock10/sema4c/adam8a/si:ch211-113g11.6/anxa1c/mdga2a/zgc:153631/asap1b/si:ch211-241e1.3/plxna3/plxna1a/rtn4rl2a/sema6ba/sema3b |
| GO:0009719 | response to endogenous stimulus | -1.32546647522578 | 0.0666420813824336 | tbx16/irs1/skia/igfbp3/pik3r3a/gsk3ba/rbm14b/gna11b/slc39a14/slc27a1b/esr2a/zyx/smad7/tll1/ltbp3/tacr1a/spry4/dusp6/trim71/anxa1a/avpr1ab/nrp2b/nr1d1/smad2/grb10a/flcn/smurf1/zeb2a/asic4b/fcho1/skib/rgmb/fgfbp1a/irs2a/drd4-rs/nptna/vegfc/fstl4/si:rp71-46j2.7/ldb2a/timp2a/tgif1/ldlrad4a/caly/fstl3/castor2/ppp4cb/zdhhc13/tsc22d3/il13ra1/dab2ipb/lifrb/golm1/rerea/chrm2a/slc2a3a/nrp2a/pnpla6/asip1/tcf21/grem2b/pik3r3b/raraa/thrb/zdhhc17/bmpr1ab/smad3b/stat6/gpr22a/nr4a2b/cpeb4a/chrng/glra1/sall1a/gata3/bmp15/dzip1/fabp4b/phip/crhbp/sp9/snx25/inhbab/vegfd/fgf12a/abhd2b/htr1aa/bmpr2a/scube1/ptger4c/ccbe1/stat1b/nrros/htr1d/lemd3/gnao1b/inpp5ka/smad6b/insig1/esr2b/ptger4a/atl1/tbxta/inhbaa/asic2/timp4.3/has2/adgrl3.1/kcp/grem1a/nr4a2a/bmpr2b/stat3/nsg2/wbp2/gnao1a/stxbp4/anxa1c/nr4a3/bdnf/elapor2b/inpp5kb/ntrk3a/frs3/irs2b/gdf3/drd2a/ntrk3b/htr1fa/chrm4a/trh/dio2/dab2ipa/gnb5a/rxfp1/c3a.2/sostdc1b/mmp9/timp2b/hpxa |

**Table S3**. Gene Ontology (GO) pathways enriched following M2 exposure and their associated genes, identified by gene set enrichment analysis (GSEA).

| ID | Description | NES | FDR(p adjusted value) | Genes implicated |
| --- | --- | --- | --- | --- |
| GO:0042254 | ribosome biogenesis | 2.68 | 1.14653846153846e-08 | imp3/rpl26/mrpl10/mrm3a/nle1/noc2l/rbis/urb2/exosc2/polr1c/rpp40/exosc6/abt1/mphosph6/snu13b/emg1/wdr18/znhit3/pwp2h/utp4/utp23/mrm3b/nop14/heatr1/mrto4/pin4/sbds/lsm6/ddx31/wdr12/nom1/nifk/utp11/tfb1m/ddx27/mrpl22/ngdn/pih1d1/wdr3/ddx51/heatr3/gnl3/mrpl20/rrp36/gar1/nvl/riok2/slx9/bxdc2/pdcd11/trmt112/nop58/dimt1l/bud23/rexo4/nip7/mettl16/mphosph10/wdr36/dkc1/npm1a/nsun5/rpusd2/ppan/mdn1/ddx49/rpl7l1/ddx52/bop1/abce1/nol6/sdad1/tbl3/nop10/ddx10/mrpl36/pak1ip1/rpl35/rbfa/zcchc4/rrp9/rps7/si:dkey-251i10.3/utp15/snu13a/rrp7a/nop56/nhp2/wdr75/ddx18/c1d/rcl1/wdr55/rrp8/tsr2/eif6/nat10/utp25/nop2/nop16/polr1d/mterf3/ebna1bp2/rps21/ftsj3/bysl/utp6/rrs1/gnl3l/rps8a/mpv17l2/rps14/nol10/mtrex/rpl5a/riok1/esf1/rpl35a/pop5/ran/rpf1/mrps11/gnl2/rpl14/rpf2/tfb2m/fbl/utp3/dhx37/rrp1/rps16/ddx54/gtpbp10/rpl7/tsr1/ddx28/znf622/malsu1/rps9/mak16/c1qbp/lsg1/naf1/kri1/rpl23a/urb1/exosc8/fcf1/pes/pop4/rps15/rpsa/mtg2/rrp15/rpl7a/mterf4/exosc7/dcaf13/rps5/rplp0/exosc9/tsr3 |
| GO:0006364 | rRNA processing | 2.64217411952833 | 1.14653846153846e-08 | imp3/mrm3a/exosc2/rpp40/exosc6/abt1/mphosph6/snu13b/emg1/wdr18/znhit3/pwp2h/utp4/utp23/mrm3b/heatr1/mrto4/pin4/lsm6/wdr12/nifk/utp11/tfb1m/ddx27/ngdn/pih1d1/wdr3/ddx51/gnl3/rrp36/gar1/riok2/slx9/pdcd11/trmt112/dimt1l/bud23/rexo4/mettl16/mphosph10/wdr36/dkc1/nsun5/rpusd2/ddx49/rpl7l1/ddx52/bop1/nol6/tbl3/nop10/ddx10/rpl35/rbfa/zcchc4/rrp9/rps7/si:dkey-251i10.3/utp15/snu13a/rrp7a/nhp2/wdr75/ddx18/c1d/rcl1/wdr55/rrp8/tsr2/eif6/nat10/utp25/nop2/ebna1bp2/rps21/ftsj3/bysl/utp6/rrs1/gnl3l/rps8a/rps14/nol10/mtrex/riok1/esf1/pop5/rpf1/mrps11/gnl2/rpf2/tfb2m/fbl/utp3/dhx37/rrp1/rps16/ddx54/rpl7/tsr1/mak16/naf1/kri1/urb1/exosc8/fcf1/pes/pop4 |
| GO:0034470 | ncRNA processing | 2.61312605149024 | 1.14653846153846e-08 | ints6/prkra/imp3/thumpd2/pus3/wdr4/qtrt2/trmu/mrm3a/mtfmt/trmt10c/exosc2/rpp21/rnf113a/rpp40/exosc6/abt1/mphosph6/snu13b/rpp30/emg1/henmt1/wdr18/tprkb/piwil2/znhit3/elp3/zgc:136564/pwp2h/elp5/utp4/trmt61a/spout1/zc3h7bb/tp53rk/utp23/mrm3b/trmt44/heatr1/mrto4/pin4/trmt6/exosc5/tyw3/rpusd4/adat1/tsen2/trmt12/lsm6/wdr12/nsun2/nifk/utp11/thumpd3/tfb1m/osgep/ddx27/urm1/ctu2/mettl3/ngdn/pih1d1/wdr3/srrt/ddx51/gnl3/rrp36/gar1/riok2/ints6l/slx9/pdcd11/trmt112/dimt1l/bud23/osgepl1/rexo4/ptcd1/mettl16/mphosph10/wdr36/dkc1/mettl2a/thada/pus1/nsun5/gtpbp3/rpusd2/ddx49/rpl7l1/ddx52/bop1/trmt10b/tdrd9/nol6/clp1/ctu1/tsen15/rtcb/tbl3/ints9/nop10/ddx10/rpl35/rbfa/zcchc4/rrp9/rpph1/rps7/si:dkey-251i10.3/utp15/snu13a/rrp7a/elp6/nhp2/wdr75/ddx18/c1d/rcl1/wdr55/rrp8/tsr2/trmt61b/elac1/trpt1/eif6/nat10/utp25/nop2/ebna1bp2/tut1/rps21/elp2/ftsj3/bysl/mettl1/pus7/utp6/rrs1/dtwd1/gnl3l/rps8a/rps14/nol10/rpp25l/thg1l/mtrex/riok1/sepsecs/esf1/eif1axb/pop5/cdk5rap1/zbtb8os/rpf1/mrps11/gnl2/rpf2/tfb2m/trmt2b/fbl/utp3/rpp38/dhx37/rtraf/rrp1/wdr6/ints11/rps16/ddx54/aars1/rpl7/tsr1/mak16/trmt5/trmt9b/naf1/kri1/urb1/snip1/exosc8/fcf1/pes/pop4 |
| GO:0016072 | rRNA metabolic process | 2.5944143069453 | 1.14653846153846e-08 | imp3/mrm3a/exosc2/rpp40/exosc6/abt1/mphosph6/snu13b/emg1/wdr18/znhit3/pwp2h/utp4/utp23/mrm3b/heatr1/mrto4/pin4/exosc5/lsm6/wdr12/nifk/utp11/tfb1m/ddx27/ngdn/pih1d1/wdr3/ddx51/gnl3/rrp36/gar1/riok2/slx9/pdcd11/trmt112/dimt1l/bud23/rexo4/mettl16/mphosph10/wdr36/dkc1/polr1e/nsun5/rpusd2/ddx49/rpl7l1/ddx52/bop1/nol6/tbl3/nop10/ddx10/rpl35/rbfa/zcchc4/rrp9/rps7/si:dkey-251i10.3/utp15/snu13a/rrp7a/nhp2/wdr75/ddx18/c1d/rcl1/wdr55/rrp8/tsr2/eif6/nat10/utp25/nop2/taf1b/ebna1bp2/rps21/ftsj3/bysl/dis3/utp6/rrs1/gnl3l/rps8a/rps14/nol10/mtrex/riok1/esf1/pop5/rpf1/mrps11/gnl2/rpf2/tfb2m/fbl/utp3/polr1a/dhx37/rrp1/rps16/ddx54/pelo/rpl7/tsr1/mak16/naf1/kri1/urb1/exosc8/fcf1/pes/pop4 |
| GO:0034660 | ncRNA metabolic process | 2.54552940229038 | 1.14653846153846e-08 | ints6/prkra/fars2/imp3/thumpd2/pus3/farsa/wdr4/qtrt2/trmu/mrm3a/dars2/wars2/mtfmt/trmt10c/exosc2/rpp21/rnf113a/rpp40/exosc6/dars1/abt1/mphosph6/snu13b/rpp30/emg1/henmt1/wdr18/tprkb/piwil2/znhit3/elp3/zgc:136564/pwp2h/elp5/utp4/trmt61a/spout1/zc3h7bb/tp53rk/utp23/mrm3b/trmt44/heatr1/mrto4/pin4/trmt6/exosc5/tyw3/lars1b/rpusd4/adat1/tsen2/trmt12/lsm6/wdr12/nsun2/nifk/utp11/thumpd3/si:ch211-256m1.8/tfb1m/mars2/osgep/ddx27/urm1/ctu2/mettl3/ngdn/pih1d1/wdr3/srrt/ddx51/snapc3/gnl3/rrp36/gar1/riok2/ints6l/slx9/pdcd11/trmt112/qrsl1/dimt1l/bud23/osgepl1/rexo4/ptcd1/mettl16/gatb/mphosph10/wdr36/dkc1/mettl2a/thada/polr1e/pus1/nsun5/gtpbp3/rpusd2/iars1/ddx49/hars/rpl7l1/ddx52/bop1/trmt10b/tdrd9/qars1/nol6/nars2/vars1/clp1/ctu1/tsen15/rtcb/tbl3/ints9/nop10/ddx10/rpl35/rbfa/zcchc4/rrp9/rpph1/rps7/si:dkey-251i10.3/utp15/snu13a/rrp7a/elp6/nhp2/wdr75/ddx18/yars1/c1d/rcl1/wdr55/rrp8/tsr2/trmt61b/elac1/rpap2/trpt1/rars1/eif6/nat10/utp25/nop2/taf1b/ebna1bp2/tut1/rps21/elp2/tars1/ftsj3/bysl/gars1/mettl1/pars2/dis3/pus7/iars2/utp6/rrs1/dtwd1/gnl3l/rps8a/rps14/nol10/rpp25l/thg1l/mtrex/riok1/sars1/sepsecs/esf1/eif1axb/pop5/vars2/mepceb/cdk5rap1/zbtb8os/rpf1/mrps11/gnl2/sars2/rpf2/tfb2m/trmt2b/fbl/utp3/polr1a/rpp38/dhx37/rtraf/rrp1/wdr6/ints11/rps16/ddx54/aars1/pelo/rpl7/tsr1/mak16/trmt5/trmt9b/naf1/kri1/dtd2/urb1/snip1/exosc8/fcf1/pes/pop4 |
| GO:0022613 | ribonucleoprotein complex biogenesis | 2.40988428678125 | 1.14653846153846e-08 | imp3/rpl26/mrpl10/mrm3a/nle1/noc2l/rbis/gemin5/urb2/exosc2/polr1c/rpp40/exosc6/abt1/mphosph6/snu13b/emg1/wdr18/znhit3/pwp2h/trnau1apa/utp4/utp23/mrm3b/nop14/heatr1/mrto4/pin4/sbds/snrpd2/lsm6/ddx31/wdr12/nom1/nifk/utp11/tfb1m/prmt7/ddx27/mrpl22/ngdn/pih1d1/gemin2/wdr3/ddx51/heatr3/gnl3/mrpl20/rrp36/gar1/nvl/snrpg/riok2/slx9/bxdc2/pdcd11/trmt112/nop58/dimt1l/bud23/prpf18/mcts1/rexo4/nip7/mettl16/mphosph10/wdr36/dkc1/npm1a/nsun5/rpusd2/ppan/mdn1/ddx49/rpl7l1/ddx52/bop1/abce1/nol6/clp1/sdad1/tbl3/nop10/ddx10/mrpl36/pak1ip1/rpl35/rbfa/zcchc4/rrp9/ptges3b/rps7/si:dkey-251i10.3/utp15/snu13a/rrp7a/nop56/eif3m/nhp2/wdr75/ddx18/c1d/nopchap1/snrpd3l/eif3ea/rcl1/wdr55/rrp8/tsr2/coil/eif6/snrpf/nat10/utp25/nop2/nop16/usp39/polr1d/mterf3/ebna1bp2/rps21/eif3jb/ftsj3/snrpe/bysl/utp6/rrs1/gnl3l/rps8a/mpv17l2/rpl38/rps14/nol10/shq1/mtrex/rpl5a/riok1/esf1/rpl35a/pop5/eif5/ran/eif3ha/rpf1/trnau1apb/mrps11/gnl2/scaf11/rpl14/rpf2/tfb2m/fbl/utp3/dhx37/rrp1/eif3s10/cdc73/rps16/ddx54/denr/gtpbp10/rpl7/tsr1/ddx28/znf622/malsu1/rps9/mak16/c1qbp/lsg1/naf1/kri1/rpl23a/urb1/exosc8/eif3s6ip/fcf1/pes/pop4/rps15/rpsa/eif2s2/mtg2/polr2d/rrp15/rpl7a/srpk1a/mterf4/exosc7/dcaf13/rps5/gemin8/rplp0/exosc9/tsr3/snrpd1/rps27.1/aar2/rpl10/rpl5b/eif3g/gtpbp4/eif3d/rpl6/yju2/noc4l |
| GO:0140053 | mitochondrial gene expression | 2.16306020123481 | 0.000153043817326382 | fastkd3/tefm/trmu/fastkd2/tbrg4/wars2/trmt10c/mrpl58/fastkd1/coa3a/mtg1/mrps18b/tfb1m/uqcc2/chchd1/tsfm/mtrf1l/qrsl1/gatb/gtpbp3/mtif2/hars/mrps24/rcc1l/tfam/trmt61b/mrpl47/mterf3/gars1/iars2/mrpl23/sars2/tfb2m/ptcd3/fastk/mrpl16 |
| GO:0043043 | peptide biosynthetic process | 2.02998011519709 | 1.14653846153846e-08 | dazl/fars2/farsa/zc3h12ab/dph7/rpl26/mrpl10/dars2/wars2/mtfmt/trmt10c/gemin5/exosc2/mrpl58/mrps5/dars1/rpp30/zar1/mrpl9/dcp1b/piwil2/coa3a/trnau1apa/mrpl35/mif4gdb/mrpl30/eif2ak1/dcps/mrpl17/dph5/lars1b/mtg1/mrps17/mrps18b/mrpl4/si:ch211-256m1.8/zgc:171480/mars2/uqcc2/chchd1/tsfm/gss/eif4ea/mrpl22/mettl3/ngdn/mtrf1l/faub/mrps18c/drg2/lsm1/mrpl27/mrpl20/dph6/magoh/eif1ad/eif2a/eefsec/qrsl1/mrpl51/mrps23/mcts1/ago3b/mettl16/gatb/mrpl18/mrps21/gtpbp3/mtif2/iars1/mrpl14/rpl12/hars/mrpl24/pdf/eif2b1/eif4e2rs1/abce1/eif2s1a/qars1/nars2/mrpl41/impact/vars1/mrpl34/fxr2/larp6a/mrps24/mrpl36/rpl35/cnot8/mrps12/mrpl13/rcc1l/eif5a2/rpl32/mrps2/rps7/mrpl12/brf1a/eif2b2/eif3m/eif4ebp3l/yars1/bdh2/eef1b2/eif3ea/mrpl11/rpl22l1/mrpl33/rpl34/rars1/eif6/dph2/ogfod1/gspt1/pde12/mrpl47/rps21/rps17/eef1a1l3/eif3jb/tars1/cnot6l/eef1da/eif4ebp1/gars1/mrps10/pars2/iars2/rc3h2/mrpl23/rpl29/zgc:171772/rpl37/rps8a/gatc/rpl38/rps3a/rps14/rps15a/srp9/larp4b/mrps30/rps20/rps29/tcea3/rpl5a/rpl15/sars1/sepsecs/mrpl3/eif1axb/rpl35a/vars2/eif5/rps18/eif3ha/zc3h15/mrpl32/mrpl37/trnau1apb/mrps11/rpl11/rpl14/rpl39/sars2/uba52/rpl36a/rps26/eif1b/ssb/igf2bp3/rpl24/eif4eb/ptcd3/rpl27/rpl23/eif3s10/rpl31/rps24/rps16/rwdd1/denr/aars1/rpl18/rpl13a/rpl22/mrps16/rbm24a/rplp2/mrpl16/mettl17/mtrfr/drg1/malsu1/rps9/rbm8a/rpl28/rack1/rplp1/rpl23a/eif3f/gspt1l/mrpl2/exosc8/larp1b/eif3s6ip/rps15/rpsa/eef1g/eif2s2/cnbpa/ftsj1/rps11/eif4e3/rplp2l/zgc:114188/polr2d/rpl17/cnot11/faua/pabpc1a/exosc7/rpl19/rps5/eif2s1b/rplp0/rpl18a/exosc9/tufm/rpl21/kars1/rpl4/eef2b/jmjd4/cnbpb/rps13/rpl8/eif5b/rps27.1/rpl10/rpl5b/eif3g/eif3d/mrps35/rpl6/gfm1/eif4e1c/rps3/eif1axa/cnot7/eif2b4/rps12/eef1a1l1/znf598/rpl9/cnot10/eef2a.1/rpl10a/cars1/rps6/rps27a/mars1/nck2a/rps27.2/rps28/mrps7/eif4bb/rpl36/rps23/eif2s3/gtpbp1l/nck2b/eif4g1a/cnot9/dcp2/rps19/dph1/ybx1/dhx29/eif3ja/mrpl15/eif4g3b/samd4a |
| GO:0006412 | translation | 2.02950881785419 | 1.14653846153846e-08 | dazl/fars2/farsa/zc3h12ab/dph7/rpl26/mrpl10/dars2/wars2/mtfmt/trmt10c/gemin5/exosc2/mrpl58/mrps5/dars1/rpp30/zar1/mrpl9/dcp1b/piwil2/coa3a/trnau1apa/mrpl35/mif4gdb/mrpl30/eif2ak1/dcps/mrpl17/dph5/lars1b/mtg1/mrps17/mrps18b/mrpl4/si:ch211-256m1.8/zgc:171480/mars2/uqcc2/chchd1/tsfm/eif4ea/mrpl22/mettl3/ngdn/mtrf1l/faub/mrps18c/drg2/lsm1/mrpl27/mrpl20/dph6/magoh/eif1ad/eif2a/eefsec/qrsl1/mrpl51/mrps23/mcts1/ago3b/mettl16/gatb/mrpl18/mrps21/gtpbp3/mtif2/iars1/mrpl14/rpl12/hars/mrpl24/pdf/eif2b1/eif4e2rs1/abce1/eif2s1a/qars1/nars2/mrpl41/impact/vars1/mrpl34/fxr2/larp6a/mrps24/mrpl36/rpl35/cnot8/mrps12/mrpl13/rcc1l/eif5a2/rpl32/mrps2/rps7/mrpl12/brf1a/eif2b2/eif3m/eif4ebp3l/yars1/eef1b2/eif3ea/mrpl11/rpl22l1/mrpl33/rpl34/rars1/eif6/dph2/ogfod1/gspt1/pde12/mrpl47/rps21/rps17/eef1a1l3/eif3jb/tars1/cnot6l/eef1da/eif4ebp1/gars1/mrps10/pars2/iars2/rc3h2/mrpl23/rpl29/zgc:171772/rpl37/rps8a/gatc/rpl38/rps3a/rps14/rps15a/srp9/larp4b/mrps30/rps20/rps29/tcea3/rpl5a/rpl15/sars1/sepsecs/mrpl3/eif1axb/rpl35a/vars2/eif5/rps18/eif3ha/zc3h15/mrpl32/mrpl37/trnau1apb/mrps11/rpl11/rpl14/rpl39/sars2/uba52/rpl36a/rps26/eif1b/ssb/igf2bp3/rpl24/eif4eb/ptcd3/rpl27/rpl23/eif3s10/rpl31/rps24/rps16/rwdd1/denr/aars1/rpl18/rpl13a/rpl22/mrps16/rbm24a/rplp2/mrpl16/mettl17/mtrfr/drg1/malsu1/rps9/rbm8a/rpl28/rack1/rplp1/rpl23a/eif3f/gspt1l/mrpl2/exosc8/larp1b/eif3s6ip/rps15/rpsa/eef1g/eif2s2/cnbpa/ftsj1/rps11/eif4e3/rplp2l/zgc:114188/polr2d/rpl17/cnot11/faua/pabpc1a/exosc7/rpl19/rps5/eif2s1b/rplp0/rpl18a/exosc9/tufm/rpl21/kars1/rpl4/eef2b/jmjd4/cnbpb/rps13/rpl8/eif5b/rps27.1/rpl10/rpl5b/eif3g/eif3d/mrps35/rpl6/gfm1/eif4e1c/rps3/eif1axa/cnot7/eif2b4/rps12/eef1a1l1/znf598/rpl9/cnot10/eef2a.1/rpl10a/cars1/rps6/rps27a/mars1/nck2a/rps27.2/rps28/mrps7/eif4bb/rpl36/rps23/eif2s3/gtpbp1l/nck2b/eif4g1a/cnot9/dcp2/rps19/dph1/ybx1/dhx29/eif3ja/mrpl15/eif4g3b/samd4a |
| GO:0045333 | cellular respiration | 1.99293254414966 | 4.41989406426173e-05 | ccn6/tefm/sfxn4/uqcc3/si:dkey-85n7.8/ugl/sdhda/ndufs8b/atpsckmt/mdh1b/mdh1ab/dhtkd1/wfs1b/uqcc2/idh3b/sdhaf2/atp5mf/atp5po/sdhdb/ndufb6/lyrm7/cox7b/uqcr10/ndufb8/mrpl24/etfdh/ndufa10/cox6c/cox4i1/ndufv1/ndufaf1/suclg2/si:dkey-190g11.3/antkmt/cox7c/ndufa7/atp5f1e/ndufa8/ndufc2/cox7a2a/sdhc/ogdhb/slc25a55b/slc25a12/coa6/sdha/ndufs6/pm20d1.1/cox6a1/uqcrq/lyrm5a/ndufs2/ndufb9/noa1/sucla2/idh1/uqcrfs1/ndufs8a/cox6a2/cox5ab/lyrm5b/uqcrh/ndufa5/ndufv2/idh3a/mdh2/idh2/cox7a2l/sdhb/dlst/uqcrb/ndufs7/cox5aa/cyc1/ND3/fh/cox5b2/ndufv3/ogdha/cox8a |
| GO:0006518 | peptide metabolic process | 1.98118588803671 | 1.14653846153846e-08 | dazl/fars2/farsa/zc3h12ab/dph7/parla/rpl26/mrpl10/dars2/wars2/mtfmt/trmt10c/gemin5/exosc2/mrpl58/mrps5/dars1/rpp30/zar1/erap2/mrpl9/dcp1b/piwil2/ggt1b/coa3a/trnau1apa/mrpl35/mif4gdb/mrpl30/eif2ak1/dcps/mrpl17/dph5/lars1b/mtg1/mrps17/mrps18b/cpz/mrpl4/si:ch211-256m1.8/zgc:171480/mars2/uqcc2/chchd1/tsfm/gss/eif4ea/mrpl22/mettl3/ngdn/mtrf1l/faub/mrps18c/drg2/lsm1/mrpl27/mrpl20/dph6/magoh/eif1ad/eif2a/eefsec/qrsl1/mrpl51/mrps23/mcts1/ago3b/mettl16/gatb/mrpl18/immp2l/mrps21/gtpbp3/mtif2/iars1/mrpl14/rpl12/hars/mrpl24/pdf/enpep/eif2b1/eif4e2rs1/abce1/eif2s1a/qars1/nars2/mrpl41/pcsk5b/impact/vars1/mrpl34/sec11a/nln/fxr2/larp6a/mrps24/mrpl36/rpl35/cnot8/mrps12/mrpl13/rcc1l/eif5a2/rpl32/mrps2/rps7/mrpl12/brf1a/eif2b2/eif3m/eif4ebp3l/yars1/bdh2/eef1b2/eif3ea/mrpl11/rpl22l1/mrpl33/rpl34/rars1/eif6/dph2/ogfod1/gspt1/pde12/sppl3/mrpl47/rps21/rps17/eef1a1l3/eif3jb/tars1/cnot6l/eef1da/eif4ebp1/gars1/mrps10/pars2/iars2/rc3h2/mrpl23/rpl29/zgc:171772/rpl37/rps8a/gatc/rpl38/spcs3/rps3a/rps14/rps15a/srp9/larp4b/mrps30/rps20/rps29/tcea3/rpl5a/rpl15/sars1/sepsecs/mrpl3/eif1axb/rpl35a/vars2/eif5/rps18/eif3ha/zc3h15/mrpl32/mrpl37/trnau1apb/mrps11/rpl11/rpl14/rpl39/sars2/uba52/rpl36a/rps26/eif1b/thop1/ssb/igf2bp3/rpl24/eif4eb/ptcd3/rpl27/rpl23/eif3s10/rpl31/rps24/rps16/rwdd1/denr/aars1/rpl18/rpl13a/rpl22/mrps16/mipepb/rbm24a/spcs1/rplp2/mrpl16/mettl17/mtrfr/drg1/malsu1/rps9/rbm8a/rpl28/rack1/rplp1/rpl23a/eif3f/gspt1l/mrpl2/exosc8/larp1b/eif3s6ip/rps15/rpsa/eef1g/eif2s2/cnbpa/ftsj1/rps11/eif4e3/rplp2l/zgc:114188/polr2d/rpl17/cnot11/faua/pabpc1a/exosc7/rpl19/rps5/eif2s1b/rplp0/rpl18a/exosc9/tufm/rpl21/kars1/rpl4/eef2b/jmjd4/cnbpb/rps13/rpl8/eif5b/rps27.1/rpl10/rpl5b/eif3g/eif3d/mrps35/rpl6/gfm1/ide/eif4e1c/rps3/eif1axa/cnot7/eif2b4/rps12/eef1a1l1/znf598/rpl9/cnot10/eef2a.1/rpl10a/spcs2/cars1/rps6/rps27a/mars1/nck2a/rps27.2/rps28/mrps7/eif4bb/rpl36/rps23/eif2s3/gtpbp1l/nck2b/lnpep/eif4g1a/cnot9/dcp2/rps19/dph1/ybx1/dhx29/eif3ja/mrpl15/eif4g3b/samd4a |
| GO:0007005 | mitochondrion organization | 1.97295174722294 | 2.83369625164005e-07 | si:dkey-32e23.4/bnip4/adck1/sqstm1/mtm1/mtx1b/mtx1a/rrm2b/sco2/si:dkey-82o10.4/vps13d/sdhaf1/uqcc3/sdhaf3/chchd4b/atp23/phb2b/timm21/ppargc1a/timm29/maip1/coa3a/sdhaf4/aifm1/ndufs8b/chchd4a/alkbh7/chchd6b/cox17/chchd7/coa4/mgme1/cox20/slc25a42/bcs1l/timm13/uqcc2/surf1/sdhaf2/timm50/spata18/lonp1/tmem126a/dnajc19/grpel1/bida/chchd3b/atp13a2/immp2l/wdr45b/lyrm7/uqcc6/cox19/etfa/cluha/dna2/coa5/snx7/apooa/hspd1/atpaf2/phb/ndufaf1/atg5/snx30/polg2/uqcc1/higd1a/chchd10/cox7a2a/primpol/atpaf1/higd2a/si:ch211-166a6.5/pet100/tomm40/apool/fam162a/agtpbp1/pmaip1/romo1/ndufa11/oxa1l/twnk/samm50/rpl37/ndufs8a/rhot1a/slc25a4/nubpl/ndufaf8/ttc19/dnajc11a/mfn1b/miga1/tmem70/dnlz/dnaja3a/mipepb/foxred1/cox7a2l/dlst/atg7/tafazzin/tomm40l/fis1 |
| GO:0002088 | lens development in camera-type eye | 1.96162227904054 | 0.000326438123176076 | fgf19/crygm2a/crybb2/crybb1l3/crygm5/crygm2d6/crygm2b/mipb/cryba1l2/crygm2d9/crygm2d15/dnajb1b/crygm2d12/crygm2d18/cryba2a/crygm2d17/crygm2d7/crygm2d1/lgsn/crygm2d5/crygm2f/cryba1l1/crybb1l2/dlx3b/crygm2d3/crygm2d20/crygm2d11/crygm2d10/crygm2d2/crygm2d21/cryba1a/dlx4b/mipa/lama1/crygm2d13/crygmxl2/crygn2/crygm2d16/cryba4/crygm2d8/tdrd7b/crygm2d14/cryba1b/crygn1/psmc3/foxe3/cryba2b/hsf4/unc45b/crygmx/crygmxl1/si:dkey-57a22.15/sox11b/crybb1l1/psmd6 |
| GO:0009145 | purine nucleoside triphosphate biosynthetic process | 1.93013788013767 | 0.00988355843878473 | nme9/uqcc3/atp5mc3a/atpsckmt/atp5mf/atp5pd/atp5po/atp5mea/atp5mc1/nme2b.2/nme6/atp5f1c/mrpl24/atp5meb/nme2a/si:dkey-190g11.3/antkmt/atp5f1e/nme3/atp5pf/atp5pb/atp5mc3b/nme7/atp5fa1/dlst/ATP6 |
| GO:0009206 | purine ribonucleoside triphosphate biosynthetic process | 1.93013788013767 | 0.00988355843878473 | nme9/uqcc3/atp5mc3a/atpsckmt/atp5mf/atp5pd/atp5po/atp5mea/atp5mc1/nme2b.2/nme6/atp5f1c/mrpl24/atp5meb/nme2a/si:dkey-190g11.3/antkmt/atp5f1e/nme3/atp5pf/atp5pb/atp5mc3b/nme7/atp5fa1/dlst/ATP6 |
| GO:0008202 | steroid metabolic process | 1.89342793339491 | 0.000798277428982877 | ch25h/sdr42e1/dhcr24/cyp46a1.1/ch25hl1.1/msmo1/amacr/nsdhl/cyp46a1.3/cyp39a1/cyp46a1.2/hmgcra/mvk/ebpl/hsd17b12b/bdh1/srd5a1/cyp51/gba/cyp21a2/cyp7a1/lipeb/slc27a2b/faxdc2/cyp46a1.4/apoea/cyp2r1/mvda/ch25hl2/fgfr4/lss/hmgcs1/ebp/cyp26c1/apoa1a/cyp27a1.4/hsd17b7/cyb5r2/cyp27a1.1/hsd17b14/rdh1/srd5a2b/erlin1/slc27a2a |
| GO:0006979 | response to oxidative stress | 1.83959019549846 | 0.00355248534040968 | bco2l/fancc/anxa1b/glrx2/keap1a/per2/prdx1/ptgs2a/sod2/park7/ptgs1/ptgs2b/lonp1/prdx3/si:ch211-120k19.1/trpm2/gpx4a/diabloa/impact/gpx1a/gpx9/hmox2a/gpx4b/gpx8/naprt/mpv17/brf2/sirt2/romo1/sod1/txndc12/ndufa6/mpv17l2/hmox1a/ryr1b/msraa |
| GO:0006260 | DNA replication | 1.7556677318185 | 0.00194590276805455 | dbf4/zranb3/polq/zgc:173742/wrn/lig4/pold3/rpa3/mcm8/wdr18/tonsl/terf1/rmi1/poln/ctc1/mgme1/rnaseh1/ssrp1a/gins3/rpa2/stn1/orc4/dna2/noc3l/mcm7/rfc5/gins2/orc5/tatdn1/pole2/polg2/rrm1/tipin/fen1/primpol/mcm3/rrm2/traip/orc1/lig1/igfbp2a/mcmbp/mcm2/wrnip1/twnk/recql/rad51/rfc2/mcm5/rpa1/esco2/cenps/mms22l/eme1/orc6/rbbp4/gins4/blm/rnaseh2a/cdc45/prim1/rfc3/helb/pcna/rbb4l/lrwd1/samhd1/orc3/dscc1/ssbp1/top1a/mcm4/pole |
| GO:0044743 | protein transmembrane import into intracellular organelle | 1.73136835035816 | 0.0470692918806812 | pex10/chchd4b/timm21/chchd4a/timm50/dnajc19/grpel1/hspd1/tomm40/romo1/pex1/dnlz/pex13/tomm40l/pex5/timm23a/tomm70a/tomm7/pex7/pam16/tomm20b/pex26 |
| GO:0050731 | positive regulation of peptidyl-tyrosine phosphorylation | 1.72535656967289 | 0.0678490288954661 | unc119a/epgn/arl2bp/btc/smtlb/hbegfa/zgc:153759 |
| GO:0034599 | cellular response to oxidative stress | 1.71630178467435 | 0.0639818065496334 | bco2l/fancc/anxa1b/glrx2/keap1a/prdx1/sod2/park7/lonp1/si:ch211-120k19.1/diabloa/impact/mpv17/brf2/sirt2/romo1/sod1/mpv17l2/ryr1b/msraa |
| GO:0042744 | hydrogen peroxide catabolic process | 1.71035395380329 | 0.0737625497040733 | prdx1/hbae5/zgc:163057/hbbe2/prdx3/hbae1.3/hbae1.1/hbae3/gpx1a |
| GO:0006974 | cellular response to DNA damage stimulus | 1.70302954476004 | 6.9422407694291e-06 | uba7/wdr76/hrob/mms19/ddias/cry5/slx1b/zswim7/fancc/zranb3/tdp2b/xpc/parpbp/rad51d/swsap1/chek2/cry-dash/si:ch211-127m7.2/msh3/palb2/polq/hic1/phkg1b/xndc1/neil1/rfwd3/rrm2b/ogg1/sfr1/wrn/fanci/rad54b/brip1/cpdp/lig4/pargl/mre11a/zgc:112496/rad51b/faap100/pold3/rpa3/atm/cdc7/mcm8/nthl1/mcm9/ddb2/ap5s1/ercc2/kmt5ab/aptx/nbn/tonsl/cep63/eya4/rmi1/alkbh7/poln/mnat1/si:dkey-74k8.3/mgme1/park7/dek/mpg/smyd2a/mettl3/spata18/polh/ssrp1a/tdp1/neil3/rpa2/stn1/poldip2/xrcc3/gtf2h3/trip13/zfyve26/fancg/msh2/dna2/mcm7/smarcad1a/rfc5/uba1/swi5/gins2/fxr2/xrcc6/rad51c/pole2/polg2/tipin/yars1/ercc6/ube2nb/prmt6/fen1/primpol/tigara/mcm3/unga/sem1/bcl2l16/rad23b/pmaip1/lig1/polr2i/bccip/nek4/vhl/mcm2/abraxas1/txndc12/rbbp8/ttf2/mlh1/rmi2/wrnip1/recql/poli/rad1/rad51/rfc2/mcm5/rpa1/ercc1/plrg1/smyd2b/cenps/psme4a/mms22l/eme1/cdca5/si:ch211-141o9.10/hltf |
| GO:0044283 | small molecule biosynthetic process | 1.67134004649573 | 0.000341676076873402 | gls2b/tecrl2b/pycr3/fads2/mcat/msmo1/sdsl/asns/gatm/chst15/elovl8a/gpib/anxa1b/prxl2b/mthfr/acacb/cyp39a1/elovl2/pycr1b/hmgcra/mvk/gch2/qdprb.1/otc/uros/ptgs2a/olah/hsd17b12b/glsl/coq5/aass/akr1a1b/srd5a3/moxd1l/pdss2/asnsd1/srr/cyp51/cyp7a1/adka/park7/lgsn/slc27a2b/apip/rfk/ptgs1/ptgs2b/mvda/pnpo/abhd2a/pcbd1/kmo/ptgesl/pah/elovl6l/fgfr4/tecra/asah2/lss/hmgcs1/ebp/apoa1a/cyp27a1.4/gch1/acmsd/hsd17b7/gnmt/coq4/pck2/elovl8b/sephs1/cdo1/ptges3b/gamt/cyp27a1.1/ndufab1b/fbp1b/impa1/bhmt/upb1/acot7/rgn/fa2h/slc27a2a/aprt/sptlc3/prtfdc1b/agxtb/g6pc3/hprt1/pdss1/ubiad1/coq6/cth/ndufa9a/qdpra |
| GO:0052646 | alditol phosphate metabolic process | 1.62136572553731 | 0.0983308153791125 | gpd1l/rfk/gpam/gpd1c/gpd1a/gpd2 |
| GO:0032259 | methylation | 1.61555246309617 | 0.00489199162953128 | hnmt/thumpd2/suv39h1a/zgc:113054/mettl21cb/wdr4/tpmt.2/etfbkmt/mrm3a/dnmt3bb.2/eef1akmt1/cskmt/trmt10c/dnmt3bb.3/emg1/ash2l/tyms/kmt5ab/henmt1/piwil2/coq5/trmt61a/vcpkmt/mrm3b/trmt44/atpsckmt/mettl4/trmt6/tyw3/trmt2a/dph5/trmt12/nsun2/eef1akmt2/thumpd3/tfb1m/prmt9/smyd2a/tpmt.1/hells/prmt7/mettl3/smyd1a/prmt3/trmt11/trmt112/dimt1l/bud23/ntmt1/mettl16/mettl21a/mettl2a/thada/nsun5/gtpbp3/amt/trmt10b/tdrd9/dnmt3bb.1/gnmt/zcchc4/gamt/cebpb/si:dkey-190g11.3/antkmt/cxxc1b/rrp8/mrm2/setd3/bhmt/prmt6/skic8/trmt61b/comta/lyn/unga/nop2/comtd1 |
| GO:0043436 | oxoacid metabolic process | 1.61491700112608 | 2.31796694375566e-05 | gls2b/tecrl2b/pycr3/fars2/fads2/ddo/mcat/farsa/amacr/sdsl/accs/etfbkmt/asns/si:dkey-238o13.4/gatm/elovl8a/gpib/anxa1b/acot20/prxl2b/dars2/acot17/wars2/mthfr/asah1a/acot11b/ugl/sdhaf3/acacb/cyp39a1/elovl2/hoga1/pycr1b/dars1/nit2/qdprb.1/gatd1/otc/abcd3b/uros/ptgs2a/olah/hsd17b12b/glsl/tdo2a/gcat/enosf1/uevld/aass/amdhd2/alkbh7/asnsd1/srr/mdh1b/mdh1ab/dhtkd1/marc1/lars1b/lpin1b/cyp7a1/tysnd1/park7/si:ch211-256m1.8/acaa1/tat/acads/lgsn/slc27a2b/mars2/apip/ptgs1/idh3b/ptgs2b/pnpo/tph1b/gpam/abhd2a/hacl1/hsd17b4/cpt2/adpgk2/kyat1/decr1/kmo/qrsl1/bpgm/ptgesl/gck/pah/elovl6l/si:ch211-212d10.2/gatb/zgc:162816/fgfr4/tecra/gtpbp3/asah2/iars1/amt/etfa/cpt1b/hars/acss1/gcshb/cmah/cyp26c1/ctps1a/acadm/qars1/nars2/vars1/hadhab/auh/bckdha/acmsd/gnmt/pck2/elovl8b/sephs1/cdo1/mrps36/ptges3b/eno1b/abcd3a/gamt/pgam2/gnpda1/ndufab1b/mccc2/pklr/yars1/aacs/hadh/gcdha/me1/bhmt/hpdb/upb1/acot7/rgn/rars1/tigara/fa2h/slc27a2a/kyat3.2/hibch/mpst/acss2/echdc2/eci1/hgd/agxtb/pm20d1.1/tars1/hadhaa/gars1/pars2/cth/qdpra/ptgr1.1/iars2/acsf2/aspg/gcsha/kynu/hadhb/acsl5/glud1b/got2b/hpdl/cpt1aa/hk2/idh1/acsl1b/sars1/ivd/acat2/vars2/acaa2/ehhadh/acot18/nags/amdhd1/sars2/phyh/ftcd/sgpl1/gip/slc25a44b/shmt1/pfas/acot15/mgst2/aldoab/elovl4b/idh3a/tecrb/aars1/ldha/elovl7a |
| GO:0044282 | small molecule catabolic process | 1.59271540808738 | 0.00441611574194559 | gls2b/cyp46a1.1/ddo/sdsl/etfbkmt/si:dkey-238o13.4/cyp46a1.3/bpnt1/cyp46a1.2/hoga1/qdprb.1/gatd1/abcd3b/glsl/tdo2a/gcat/enosf1/aass/srd5a3/amdhd2/srr/zgc:103586/rbks/miox/lpin1b/cyp7a1/tysnd1/acaa1/tat/acads/cyp46a1.4/apoea/uraha/sord/abhd2a/hacl1/hsd17b4/cpt2/decr1/kmo/pah/zgc:162816/amt/etfa/cpt1b/gcshb/cdaa/acadm/hadhab/auh/bckdha/pck2/cdo1/abcd3a/adh8b/gnpda1/mccc2/impa1/hadh/gcdha/hpdb/kyat3.2/hibch/echdc2/eci1/hgd/hadhaa/qdpra/gcsha/kynu/hadhb/glud1b/got2b/esd/cpt1aa/ivd/acat2/acaa2/ehhadh/amdhd1/phyh/ftcd/slc25a44b/shmt1 |
| GO:1901361 | organic cyclic compound catabolic process | 1.58649464452635 | 0.00292032014134612 | hnmt/nudt4b/cyp46a1.1/dazl/dnase1l1/blvra/fastkd3/fitm2/zc3h12ab/fastkd2/cyp46a1.3/tbrg4/nudt18/exosc2/cyp46a1.2/fastkd1/exosc6/qdprb.1/entpd3/dcp1b/pde4cb/tdo2a/nudt15/moxd1l/zgc:103586/dut/heatr1/mrto4/dcps/exosc5/cyp7a1/gda/acaa1/tat/dctpp1/cyp46a1.4/tent5bb/apoea/rnaseh1/mettl3/uraha/lsm1/magoh/kmo/lsm3/pah/rexo4/ago3b/mettl16/gtpbp3/xdh/cdaa/dna2/nudt4a/fxr2/cnot8/rnaset2l/entpd5b/dnase1l4.1/hsd17b14/hmox2a/fhit/hint1/dpys/hpdb/comta/fen1/kyat3.2/pde12/allc/hgd/cnot6l/edc3/nbas/dis3/qdpra/plex9.2/rc3h2/nudt12/kynu/tet2/syncripl/endog/hmox1a/mtrex/entpd5a |
| GO:0071826 | ribonucleoprotein complex subunit organization | 1.57792704614773 | 0.0130343312989884 | nle1/gemin5/zfand1/abt1/znhit3/pwp2h/trnau1apa/mrto4/snrpd2/prmt7/pih1d1/gemin2/mrpl20/snrpg/bxdc2/prpf18/mcts1/ppan/mdn1/clp1/ptges3b/rrp7a/eif3m/nopchap1/snrpd3l/eif3ea/coil/eif6/snrpf/usp39/eif3jb/snrpe/rpl38/rps14/shq1/rpl5a/eif5/eif3ha/trnau1apb/mrps11/scaf11/rpf2/eif3s10/cdc73/denr/ddx28/naf1/rpl23a/eif3s6ip/rps15/rpsa/eif2s2/polr2d/srpk1a/rps5/gemin8/rplp0/snrpd1/rps27.1/aar2/rpl10/rpl5b/eif3g/eif3d/rpl6/yju2/rps27.2/rps28/mrps7/gemin6/eif2s3/nudt21/dyrk3/lsm4/snrpc/rps19/eif3ja/eif3c |
| GO:0055088 | lipid homeostasis | 1.54238929377417 | 0.0961825238314319 | adck1/fitm2/zgc:162608/cyp39a1/spx/abcg4b/ormdl2/apobb.1/psmd10/apobb.2/cyp7a1/apoa4a/pnpla3/tlcd1/apoa4b.2/apoba/fabp2/angptl8 |
| GO:0140014 | mitotic nuclear division | 1.53553828880172 | 0.0229731377900925 | ncaph/epgn/btc/pdcd2/nde1/chtf8/si:ch211-244o22.2/cep192/sgo1/map10/mad1l1/incenp/gnl3/kif20bb/ndc80/anapc5/gcna/ncapg/bub3/nme6/mad2l1/reep3b/anapc15/birc5b/cdca8/esco1/ncapg2/abraxas2/ccnb1/ranbp1/aurkb/racgap1/anapc11/ube2s/zgc:86764/pmaip1/bccip/abraxas1/zw10/esco2/ncaph2/cenpk/spdl1/prc1b/ran/cdca5/gnl2/map9/mki67/kifc1/ube2ib/chmp7/kif11 |
| GO:0019439 | aromatic compound catabolic process | 1.4470736645875 | 0.0239204409208845 | hnmt/nudt4b/dazl/dnase1l1/blvra/fastkd3/fitm2/zc3h12ab/fastkd2/tbrg4/nudt18/exosc2/fastkd1/exosc6/qdprb.1/entpd3/dcp1b/pde4cb/tdo2a/nudt15/moxd1l/zgc:103586/dut/heatr1/mrto4/dcps/exosc5/gda/acaa1/tat/dctpp1/tent5bb/rnaseh1/mettl3/uraha/lsm1/magoh/kmo/lsm3/pah/rexo4/ago3b/mettl16/gtpbp3/xdh/cdaa/dna2/nudt4a/fxr2/cnot8/rnaset2l/entpd5b/dnase1l4.1/hmox2a/fhit/hint1/hpdb/comta/fen1/kyat3.2/pde12/hgd/cnot6l/edc3/nbas/dis3/qdpra/plex9.2/rc3h2/nudt12/kynu/tet2/syncripl/endog/hmox1a/mtrex/entpd5a/ephx1 |
| GO:0044270 | cellular nitrogen compound catabolic process | 1.42325048500041 | 0.0467129045300582 | hnmt/nudt4b/dazl/dnase1l1/blvra/fastkd3/fitm2/zc3h12ab/fastkd2/tbrg4/nudt18/exosc2/fastkd1/exosc6/entpd3/dcp1b/pde4cb/tdo2a/nudt15/zgc:103586/dut/heatr1/mrto4/dcps/exosc5/gda/dctpp1/tent5bb/rnaseh1/mettl3/uraha/lsm1/magoh/kmo/lsm3/rexo4/ago3b/mettl16/gtpbp3/xdh/cdaa/dna2/nudt4a/fxr2/cnot8/rnaset2l/entpd5b/dnase1l4.1/hmox2a/fhit/hint1/dpys/bhmt/fen1/pde12/cnot6l/edc3/nbas/dis3/plex9.2/rc3h2/nudt12/kynu/tet2/syncripl/endog/hmox1a/mtrex/entpd5a |
| GO:0007601 | visual perception | 1.4025303608205 | 0.0873969864431149 | unc119a/gprc5c/crygm2a/crybb2/crybb1l3/crygm5/grk7a/crygm2d6/crygm2b/cryba1l2/arr3b/crygm2d9/crygm2d15/crygm2d12/crygm2d18/cryba2a/opn1sw1/crygm2d17/crygm2d7/crygm2d1/arl6/crygm2d5/crygm2f/cryba1l1/pdcb/crybb1l2/crygm2d3/crygm2d20/crygm2d11/crygm2d10/crygm2d2/crygm2d21/cryba1a/opn1mw1/crygm2d13/crygmxl2/guca1b/crygn2/crygm2d16/arr3a/cryba4/crygm2d8/crygm2d14/cryba1b/crygn1/prph2a/lum/cryba2b/opn1sw2/crygmx/pdca/crygmxl1/si:dkey-57a22.15/crybb1l1/pde6a/gucy2f/grk7b/opn1lw2/nphp3/prph2lb/pde6d/crybgx/tmtops2a |
| GO:0099537 | trans-synaptic signaling | -2.69921105199218 | 1.14653846153846e-08 | dlg1b/dtnbp1b/ncdn/gabrr2a/gabrb1/cplx2b/drd4-rs/cacng8b/otofa/sv2bb/dlg3/cdh2/penka/cep89/chrnb5a/synj1/cart2/cspg5b/gria3a/dlgap1a/chrm2a/snap25a/napbb/penkb/glrbb/slc5a7a/napba/chata/spg11/apba1b/erc1a/grid1a/stxbp1b/cplx2a/clstn2b/cacnb1/gabrb3/chrng/hdc/sncb/glra1/drd3/cacng5a/si:ch211-81a5.8/syn1/erc2/gria1a/grm4/grm6b/stx3a/caskb/chrnb3a/ppp1r9ba/slc18a3a/snap25b/jph3b/mctp2b/grm8b/cart3/slc6a5/calb1/chrna3/htr1aa/calb2b/snap47/htr1d/pdyn/jph3a/cacnb2b/ptchd1/calb2a/syt7b/syn2a/cacng7b/chrna5/grin2aa/rims1b/shisa9a/clstn1/syt4/rab3ab/dlgap4b/lrrc4ca/chrna2a/gabra4/glra4b/nlgn4xa/gria2b/nsmfb/gria4a/slc17a6b/clstn2a/sncgb/htr3a/gabrb2a/lrrc4.1/cadpsb/grin1a/grm2a/ppp1r9bb/syt11a/unc13a/cacng2b/stxbp1a/syt1a/chrnb2/slc12a5b/chrna6/cacng3b/shisa7b/nlgn3a/gria3b/stx1b/dlg2/ppfia3/grin2ab/cacng2a/grm8a/lrrc4cb/atcaya/slc12a5a/chrna4b/grm2b/nlgn3b/cacnb4b/grid2/clstn3/syt2a/apba2b/grik1a/cspg5a/sv2a/pnocb/grin1b/baiap3/cacna1ab/grin2da/nlgn2a/nrxn2a/si:dkey-174m14.3/drd2a/rph3ab/dagla/grik5/htr1fa/shisa9b/gabrp/chrm4a/grin3bb/grin2bb/insyn1/grik4/gabrg2/grm1a/cacng8a/stx11a/gabrb4/dlgap3/htr7c/neto1l/nalcn/gria4b |
| GO:0007268 | chemical synaptic transmission | -2.68729704840182 | 1.14653846153846e-08 | dlg1b/dtnbp1b/ncdn/gabrr2a/gabrb1/cplx2b/drd4-rs/cacng8b/otofa/sv2bb/dlg3/cdh2/penka/cep89/chrnb5a/synj1/cart2/cspg5b/gria3a/dlgap1a/chrm2a/snap25a/napbb/penkb/glrbb/slc5a7a/napba/chata/spg11/apba1b/erc1a/grid1a/stxbp1b/cplx2a/clstn2b/cacnb1/gabrb3/chrng/hdc/sncb/glra1/drd3/cacng5a/si:ch211-81a5.8/syn1/erc2/gria1a/grm4/grm6b/stx3a/caskb/chrnb3a/ppp1r9ba/slc18a3a/snap25b/jph3b/mctp2b/grm8b/cart3/slc6a5/calb1/chrna3/htr1aa/calb2b/snap47/htr1d/pdyn/jph3a/cacnb2b/ptchd1/calb2a/syt7b/syn2a/cacng7b/chrna5/grin2aa/rims1b/shisa9a/clstn1/syt4/rab3ab/dlgap4b/lrrc4ca/chrna2a/gabra4/glra4b/nlgn4xa/gria2b/nsmfb/gria4a/slc17a6b/clstn2a/sncgb/htr3a/gabrb2a/lrrc4.1/cadpsb/grin1a/grm2a/ppp1r9bb/syt11a/unc13a/cacng2b/stxbp1a/syt1a/chrnb2/slc12a5b/chrna6/cacng3b/shisa7b/nlgn3a/gria3b/stx1b/dlg2/ppfia3/grin2ab/cacng2a/grm8a/lrrc4cb/atcaya/slc12a5a/chrna4b/grm2b/nlgn3b/cacnb4b/grid2/clstn3/syt2a/apba2b/grik1a/cspg5a/sv2a/pnocb/grin1b/baiap3/cacna1ab/grin2da/nlgn2a/nrxn2a/si:dkey-174m14.3/drd2a/rph3ab/grik5/htr1fa/shisa9b/gabrp/chrm4a/grin3bb/grin2bb/insyn1/grik4/gabrg2/grm1a/cacng8a/stx11a/gabrb4/dlgap3/htr7c/neto1l/nalcn/gria4b |
| GO:0098916 | anterograde trans-synaptic signaling | -2.68729704840182 | 1.14653846153846e-08 | dlg1b/dtnbp1b/ncdn/gabrr2a/gabrb1/cplx2b/drd4-rs/cacng8b/otofa/sv2bb/dlg3/cdh2/penka/cep89/chrnb5a/synj1/cart2/cspg5b/gria3a/dlgap1a/chrm2a/snap25a/napbb/penkb/glrbb/slc5a7a/napba/chata/spg11/apba1b/erc1a/grid1a/stxbp1b/cplx2a/clstn2b/cacnb1/gabrb3/chrng/hdc/sncb/glra1/drd3/cacng5a/si:ch211-81a5.8/syn1/erc2/gria1a/grm4/grm6b/stx3a/caskb/chrnb3a/ppp1r9ba/slc18a3a/snap25b/jph3b/mctp2b/grm8b/cart3/slc6a5/calb1/chrna3/htr1aa/calb2b/snap47/htr1d/pdyn/jph3a/cacnb2b/ptchd1/calb2a/syt7b/syn2a/cacng7b/chrna5/grin2aa/rims1b/shisa9a/clstn1/syt4/rab3ab/dlgap4b/lrrc4ca/chrna2a/gabra4/glra4b/nlgn4xa/gria2b/nsmfb/gria4a/slc17a6b/clstn2a/sncgb/htr3a/gabrb2a/lrrc4.1/cadpsb/grin1a/grm2a/ppp1r9bb/syt11a/unc13a/cacng2b/stxbp1a/syt1a/chrnb2/slc12a5b/chrna6/cacng3b/shisa7b/nlgn3a/gria3b/stx1b/dlg2/ppfia3/grin2ab/cacng2a/grm8a/lrrc4cb/atcaya/slc12a5a/chrna4b/grm2b/nlgn3b/cacnb4b/grid2/clstn3/syt2a/apba2b/grik1a/cspg5a/sv2a/pnocb/grin1b/baiap3/cacna1ab/grin2da/nlgn2a/nrxn2a/si:dkey-174m14.3/drd2a/rph3ab/grik5/htr1fa/shisa9b/gabrp/chrm4a/grin3bb/grin2bb/insyn1/grik4/gabrg2/grm1a/cacng8a/stx11a/gabrb4/dlgap3/htr7c/neto1l/nalcn/gria4b |
| GO:0050804 | modulation of chemical synaptic transmission | -2.68416739586281 | 1.14653846153846e-08 | erc1a/grid1a/cplx2a/clstn2b/drd3/cacng5a/si:ch211-81a5.8/erc2/gria1a/grm4/grm6b/caskb/ppp1r9ba/jph3b/mctp2b/grm8b/calb1/calb2b/jph3a/calb2a/cacng7b/grin2aa/rims1b/shisa9a/clstn1/dlgap4b/lrrc4ca/nlgn4xa/gria2b/nsmfb/gria4a/clstn2a/lrrc4.1/grm2a/ppp1r9bb/cacng2b/cacng3b/shisa7b/nlgn3a/gria3b/stx1b/grin2ab/cacng2a/grm8a/lrrc4cb/atcaya/grm2b/nlgn3b/grid2/clstn3/grik1a/baiap3/grin2da/nlgn2a/si:dkey-174m14.3/drd2a/grik5/shisa9b/grin3bb/grin2bb/grik4/grm1a/cacng8a/dlgap3/neto1l/nalcn/gria4b |
| GO:0099177 | regulation of trans-synaptic signaling | -2.68416739586281 | 1.14653846153846e-08 | erc1a/grid1a/cplx2a/clstn2b/drd3/cacng5a/si:ch211-81a5.8/erc2/gria1a/grm4/grm6b/caskb/ppp1r9ba/jph3b/mctp2b/grm8b/calb1/calb2b/jph3a/calb2a/cacng7b/grin2aa/rims1b/shisa9a/clstn1/dlgap4b/lrrc4ca/nlgn4xa/gria2b/nsmfb/gria4a/clstn2a/lrrc4.1/grm2a/ppp1r9bb/cacng2b/cacng3b/shisa7b/nlgn3a/gria3b/stx1b/grin2ab/cacng2a/grm8a/lrrc4cb/atcaya/grm2b/nlgn3b/grid2/clstn3/grik1a/baiap3/grin2da/nlgn2a/si:dkey-174m14.3/drd2a/grik5/shisa9b/grin3bb/grin2bb/grik4/grm1a/cacng8a/dlgap3/neto1l/nalcn/gria4b |
| GO:0099536 | synaptic signaling | -2.6833837570563 | 1.14653846153846e-08 | dlg1b/dtnbp1b/ncdn/gabrr2a/gabrb1/cplx2b/drd4-rs/cacng8b/otofa/sv2bb/dlg3/cdh2/penka/cep89/chrnb5a/synj1/cart2/cspg5b/gria3a/dlgap1a/chrm2a/snap25a/napbb/penkb/glrbb/slc5a7a/napba/chata/spg11/apba1b/erc1a/grid1a/stxbp1b/cplx2a/clstn2b/cacnb1/gabrb3/chrng/hdc/sncb/glra1/drd3/cacng5a/si:ch211-81a5.8/syn1/erc2/gria1a/grm4/grm6b/stx3a/caskb/chrnb3a/ppp1r9ba/slc18a3a/snap25b/jph3b/mctp2b/grm8b/cart3/slc6a5/calb1/chrna3/htr1aa/calb2b/snap47/htr1d/pdyn/jph3a/cacnb2b/ptchd1/calb2a/syt7b/syn2a/cacng7b/chrna5/grin2aa/rims1b/shisa9a/clstn1/syt4/rab3ab/dlgap4b/lrrc4ca/chrna2a/gabra4/glra4b/nlgn4xa/gria2b/nsmfb/gria4a/slc17a6b/clstn2a/sncgb/htr3a/gabrb2a/lrrc4.1/cadpsb/grin1a/grm2a/ppp1r9bb/syt11a/unc13a/cacng2b/stxbp1a/syt1a/chrnb2/slc12a5b/chrna6/cacng3b/shisa7b/nlgn3a/gria3b/stx1b/dlg2/ppfia3/grin2ab/cacng2a/grm8a/lrrc4cb/atcaya/slc12a5a/chrna4b/grm2b/nlgn3b/cacnb4b/grid2/clstn3/syt2a/apba2b/grik1a/cspg5a/sv2a/pnocb/grin1b/baiap3/cacna1ab/grin2da/nlgn2a/nrxn2a/si:dkey-174m14.3/drd2a/rph3ab/dagla/grik5/htr1fa/shisa9b/gabrp/chrm4a/grin3bb/grin2bb/insyn1/grik4/gabrg2/grm1a/cacng8a/stx11a/gabrb4/scn1lab/dlgap3/htr7c/neto1l/nalcn/gria4b |
| GO:0098742 | cell-cell adhesion via plasma-membrane adhesion molecules | -2.34722281326859 | 1.14653846153846e-08 | arvcfb/zeb1a/cdh2/prnprs3/ptk7a/pcdh2ab10/zgc:165604/si:ch211-186j3.6/tenm3/dchs1a/ret/pcdh8/ptprsa/clstn2b/dscama/pcdh17/pcdh2ab9/pecam1a/pcdh10b/pcdh7a/tenm2b/robo3/si:ch211-199f5.1/dsc2l/pcdh1g22/jag2b/nectin3b/tenm4/pcdh1g31/pcdh7b/pcdh1g29/pcdh2g13/pcdh1a/clstn1/pcdh19/cadm1b/lrrc4ca/pcdh1a4/pcdh2ab3/cadm3/dscamb/ptprdb/pcdh10a/pcdh11/clstn2a/lrrc4.1/igsf21b/robo4/fat3a/cadm4/si:ch211-214p13.3/cdh18a/si:dkey-237h12.3/lrrc4cb/igsf21a/tenm2a/esama/cdh13/clstn3/cdh15/cntn3a.1/si:ch73-233f7.1/pcdh1g32/cdh4/pcdh1g11/si:ch73-233f7.3/cdh7a/pcdh1g18/adgrl1a/pcdh1gc5 |
| GO:0050808 | synapse organization | -2.21669824394954 | 9.94928637272983e-08 | dbn1/ctnna2/cdh2/synj1/ptk7a/dbnlb/cbln2b/ephb3a/sez6a/tjp1b/ctbp2a/erc1a/fbxo45/gphnb/neurod2/ptprsa/clstn2b/cacnb1/sncb/erc2/tanc2a/ppfia4/gfpt1/slitrk4/gphna/cacnb2b/myo9ab/inpp5ka/caprin2/pcloa/clstn1/rab3ab/lrrc4ca/ptprdb/nlgn4xa/clstn2a/sncgb/sybu/lrrc4.1/arhgef9b/unc13a/dock10/ctnnd2b/nbeaa/nlgn3a/bsnb/cacna1da/ppfia3/lrrc4cb/epb41l3a/grna/bdnf/nlgn3b/ctnnd2a/cacnb4b/pclob/clstn3/slitrk2/sez6l2/nlgn2a/nrxn2a |
| GO:0048667 | cell morphogenesis involved in neuron differentiation | -2.08310048291656 | 1.14653846153846e-08 | plxnb1b/pik3cd/mycbp2/rb1/ptprfb/zgc:100846/ptprfa/isl2b/nrp2b/arhgap35a/rapgef1a/whrna/emb/pik3ca/rbpms2a/aplp2/mapk8ip3/npr1a/sema3fa/emx2/olfm1a/unc5b/nptna/vegfc/dbn1/kctd12.1/si:ch211-129c21.1/phox2bb/unk/ubap1/ctnna2/cntn4/cdh2/trarg1a/kalrnb/ulk2/ptk7a/dbnlb/efna1b/ephb3a/slc35b2/rbpms2b/fus/map1ab/dock7/foxb1a/nsfa/rab33ba/sipa1l1/nrp2a/ush1ga/pnpla6/kctd13/isl1a/fezf2/slit2/tenm3/rufy3/nfasca/hecw1b/sim1a/adcy8/pak1/spg11/ret/kif1b/arnt2/sema3bl/zdhhc17/plxna4/appa/nedd4l/arhgap35b/efna2a/dscama/apc/robo2/pcdh17/casp9/epha4a/hdc/anos1a/trioa/lhx9/epha7/rtn4r/cyfip2/dpysl3/draxinb/slit1b/robo3/usp33/map1sb/appb/tanc2a/dzip1/sema5a/kifbp/chad/robo1/tnr/whrnb/tenm4/hecw2a/epha4l/kif5c/slitrk4/islr2/nptnb/lhx2b/myo9ab/caprin2/kalrna/cdkl5/ephb2b/epha4b/clstn1/rac3a/brsk1b/aplp1/rab3ab/gap43/lrrc4ca/srrm4/lhx1a/map1aa/prmt8b/atl1/nkx2.2a/arhgef25b/dscamb/triob/tmem59l/myo10l3/nsmfb/slit3/dpysl5a/chst11/fzd3a/kif5aa/robo4/efnb3b/vax1/stat3/dock10/ctnnd2b/zgc:77784/sema4c/si:ch211-113g11.6/rac1l/elavl4/myo10l1/ntn1b/scn8aa/lrrc4cb/plppr4a/grna/bdnf/ctnnd2a/slitrk2/nkx6.1/ncam1b/cntn3a.1/nrxn2a/dagla/grxcr1a/efna3a/dpysl5b/thsd7aa/si:ch211-241e1.3/plxna3/l1camb/plxna1a/nrcama/sema6ba/sema3b |
| GO:0048858 | cell projection morphogenesis | -2.05831448361924 | 1.14653846153846e-08 | plxnb1b/tmem108/pik3cd/mycbp2/rb1/ptprfb/zgc:100846/ptprfa/spag6/isl2b/nrp2b/arhgap35a/rapgef1a/emb/pik3ca/rbpms2a/aplp2/mapk8ip3/npr1a/sema3fa/olfm1a/unc5b/nptna/vegfc/dbn1/kctd12.1/si:ch211-129c21.1/phox2bb/ubap1/ctnna2/cntn4/cdh2/trarg1a/kalrnb/ulk2/ptk7a/dbnlb/cspg5b/efna1b/ephb3a/slc35b2/rbpms2b/fus/map1ab/dock7/foxb1a/nsfa/rab33ba/sipa1l1/tnika/nrp2a/pnpla6/kctd13/isl1a/fezf2/mink1/slit2/tenm3/rufy3/nfasca/hecw1b/sim1a/adcy8/pak1/spg11/ret/kif1b/arnt2/sema3bl/zdhhc17/plxna4/appa/nedd4l/arhgap35b/efna2a/dscama/apc/robo2/pcdh17/casp9/epha4a/hdc/anos1a/trioa/lhx9/epha7/rtn4r/cyfip2/dpysl3/draxinb/slit1b/robo3/usp33/map1sb/appb/tanc2a/dzip1/nyap2b/sema5a/kifbp/chad/pacsin1a/robo1/tnr/tenm4/hecw2a/epha4l/kif5c/slitrk4/islr2/nptnb/lhx2b/myo9ab/caprin2/kalrna/cdkl5/ephb2b/epha4b/clstn1/rac3a/brsk1b/aplp1/rab3ab/gap43/lrrc4ca/srrm4/lhx1a/map1aa/prmt8b/atl1/nkx2.2a/arhgef25b/dscamb/triob/tmem59l/myo10l3/gas7a/nsmfb/slit3/dpysl5a/chst11/fzd3a/kif5aa/robo4/efnb3b/vax1/stat3/dock10/ctnnd2b/zgc:77784/sema4c/si:ch211-113g11.6/rac1l/elavl4/myo10l1/ntn1b/scn8aa/lrrc4cb/plppr4a/grna/bdnf/ctnnd2a/nrn1la/slitrk2/nkx6.1/cspg5a/ncam1b/cntn3a.1/nrxn2a/dagla/efna3a/dpysl5b/inab/thsd7aa/si:ch211-241e1.3/plxna3/l1camb/plxna1a/nrcama/sema6ba/sema3b |
| GO:0048812 | neuron projection morphogenesis | -2.04924995520432 | 1.14653846153846e-08 | plxnb1b/tmem108/pik3cd/mycbp2/rb1/ptprfb/zgc:100846/ptprfa/spag6/isl2b/nrp2b/arhgap35a/rapgef1a/emb/pik3ca/rbpms2a/aplp2/mapk8ip3/npr1a/sema3fa/olfm1a/unc5b/nptna/vegfc/dbn1/kctd12.1/si:ch211-129c21.1/phox2bb/ubap1/ctnna2/cntn4/cdh2/trarg1a/kalrnb/ulk2/ptk7a/dbnlb/efna1b/ephb3a/slc35b2/rbpms2b/fus/map1ab/dock7/foxb1a/nsfa/rab33ba/sipa1l1/tnika/nrp2a/pnpla6/kctd13/isl1a/fezf2/mink1/slit2/tenm3/rufy3/nfasca/hecw1b/sim1a/adcy8/pak1/spg11/ret/kif1b/arnt2/sema3bl/zdhhc17/plxna4/appa/nedd4l/arhgap35b/efna2a/dscama/apc/robo2/pcdh17/casp9/epha4a/hdc/anos1a/trioa/lhx9/epha7/rtn4r/cyfip2/dpysl3/draxinb/slit1b/robo3/usp33/map1sb/appb/tanc2a/dzip1/nyap2b/sema5a/kifbp/chad/pacsin1a/robo1/tnr/tenm4/hecw2a/epha4l/kif5c/slitrk4/islr2/nptnb/lhx2b/myo9ab/caprin2/kalrna/cdkl5/ephb2b/epha4b/clstn1/rac3a/brsk1b/aplp1/rab3ab/gap43/lrrc4ca/srrm4/lhx1a/map1aa/prmt8b/atl1/nkx2.2a/arhgef25b/dscamb/triob/tmem59l/myo10l3/gas7a/nsmfb/slit3/dpysl5a/chst11/fzd3a/kif5aa/robo4/efnb3b/vax1/stat3/dock10/ctnnd2b/zgc:77784/sema4c/si:ch211-113g11.6/rac1l/elavl4/myo10l1/ntn1b/scn8aa/lrrc4cb/plppr4a/grna/bdnf/ctnnd2a/nrn1la/slitrk2/nkx6.1/ncam1b/cntn3a.1/nrxn2a/dagla/efna3a/dpysl5b/inab/thsd7aa/si:ch211-241e1.3/plxna3/l1camb/plxna1a/nrcama/sema6ba/sema3b |
| GO:0120039 | plasma membrane bounded cell projection morphogenesis | -2.0490209096944 | 1.14653846153846e-08 | plxnb1b/tmem108/pik3cd/mycbp2/rb1/ptprfb/zgc:100846/ptprfa/spag6/isl2b/nrp2b/arhgap35a/rapgef1a/emb/pik3ca/rbpms2a/aplp2/mapk8ip3/npr1a/sema3fa/olfm1a/unc5b/nptna/vegfc/dbn1/kctd12.1/si:ch211-129c21.1/phox2bb/ubap1/ctnna2/cntn4/cdh2/trarg1a/kalrnb/ulk2/ptk7a/dbnlb/efna1b/ephb3a/slc35b2/rbpms2b/fus/map1ab/dock7/foxb1a/nsfa/rab33ba/sipa1l1/tnika/nrp2a/pnpla6/kctd13/isl1a/fezf2/mink1/slit2/tenm3/rufy3/nfasca/hecw1b/sim1a/adcy8/pak1/spg11/ret/kif1b/arnt2/sema3bl/zdhhc17/plxna4/appa/nedd4l/arhgap35b/efna2a/dscama/apc/robo2/pcdh17/casp9/epha4a/hdc/anos1a/trioa/lhx9/epha7/rtn4r/cyfip2/dpysl3/draxinb/slit1b/robo3/usp33/map1sb/appb/tanc2a/dzip1/nyap2b/sema5a/kifbp/chad/pacsin1a/robo1/tnr/tenm4/hecw2a/epha4l/kif5c/slitrk4/islr2/nptnb/lhx2b/myo9ab/caprin2/kalrna/cdkl5/ephb2b/epha4b/clstn1/rac3a/brsk1b/aplp1/rab3ab/gap43/lrrc4ca/srrm4/lhx1a/map1aa/prmt8b/atl1/nkx2.2a/arhgef25b/dscamb/triob/tmem59l/myo10l3/gas7a/nsmfb/slit3/dpysl5a/chst11/fzd3a/kif5aa/robo4/efnb3b/vax1/stat3/dock10/ctnnd2b/zgc:77784/sema4c/si:ch211-113g11.6/rac1l/elavl4/myo10l1/ntn1b/scn8aa/lrrc4cb/plppr4a/grna/bdnf/ctnnd2a/nrn1la/slitrk2/nkx6.1/ncam1b/cntn3a.1/nrxn2a/dagla/efna3a/dpysl5b/inab/thsd7aa/si:ch211-241e1.3/plxna3/l1camb/plxna1a/nrcama/sema6ba/sema3b |
| GO:0032990 | cell part morphogenesis | -2.03560178882976 | 1.14653846153846e-08 | plxnb1b/tmem108/pik3cd/mycbp2/rb1/ptprfb/zgc:100846/ptprfa/spag6/isl2b/nrp2b/arhgap35a/rapgef1a/emb/pik3ca/rbpms2a/aplp2/mapk8ip3/npr1a/sema3fa/olfm1a/unc5b/nptna/vegfc/dbn1/kctd12.1/si:ch211-129c21.1/phox2bb/ubap1/ctnna2/cntn4/cdh2/trarg1a/kalrnb/ulk2/ptk7a/dbnlb/supv3l1/cspg5b/efna1b/ephb3a/slc35b2/rbpms2b/fus/map1ab/dock7/foxb1a/nsfa/rab33ba/sipa1l1/tnika/nrp2a/pnpla6/kctd13/isl1a/fezf2/mink1/slit2/tenm3/rufy3/nfasca/hecw1b/sim1a/adcy8/pak1/spg11/ret/kif1b/arnt2/sema3bl/zdhhc17/plxna4/appa/nedd4l/arhgap35b/efna2a/dscama/apc/robo2/pcdh17/casp9/epha4a/hdc/anos1a/trioa/lhx9/epha7/rtn4r/cyfip2/dpysl3/draxinb/slit1b/robo3/usp33/map1sb/appb/tanc2a/dzip1/nyap2b/sema5a/kifbp/chad/pacsin1a/robo1/tnr/tenm4/hecw2a/epha4l/kif5c/slitrk4/islr2/nptnb/lhx2b/myo9ab/caprin2/kalrna/cdkl5/ephb2b/epha4b/clstn1/rac3a/brsk1b/aplp1/rab3ab/gap43/lrrc4ca/srrm4/lhx1a/map1aa/prmt8b/atl1/nkx2.2a/arhgef25b/dscamb/triob/tmem59l/myo10l3/gas7a/nsmfb/slit3/dpysl5a/chst11/fzd3a/kif5aa/robo4/efnb3b/vax1/stat3/dock10/ctnnd2b/zgc:77784/sema4c/si:ch211-113g11.6/rac1l/elavl4/myo10l1/ntn1b/scn8aa/lrrc4cb/plppr4a/grna/bdnf/ctnnd2a/nrn1la/slitrk2/nkx6.1/cspg5a/ncam1b/cntn3a.1/nrxn2a/dagla/efna3a/dpysl5b/inab/thsd7aa/si:ch211-241e1.3/plxna3/l1camb/plxna1a/nrcama/sema6ba/sema3b |
| GO:0000904 | cell morphogenesis involved in differentiation | -2.02983448887318 | 1.14653846153846e-08 | pik3cd/mycbp2/rb1/ptprfb/adgrg1/zgc:100846/ptprfa/isl2b/nrp2b/arhgap35a/rapgef1a/whrna/emb/pik3ca/erbb2/rbpms2a/aplp2/mapk8ip3/npr1a/sema3fa/emx2/olfm1a/unc5b/nptna/vegfc/dbn1/kctd12.1/si:ch211-129c21.1/phox2bb/unk/ubap1/ctnna2/cntn4/cdh2/trarg1a/kalrnb/ulk2/ptk7a/dbnlb/efna1b/rilpl2/ephb3a/slc35b2/rbpms2b/fus/map1ab/dock7/foxb1a/nsfa/rab33ba/sipa1l1/nrp2a/ush1ga/pnpla6/kctd13/isl1a/fezf2/slit2/tenm3/rufy3/nfasca/hecw1b/sim1a/adcy8/pak1/spg11/ret/kif1b/arnt2/sema3bl/zdhhc17/plxna4/appa/nedd4l/arhgap35b/efna2a/dscama/apc/robo2/pcdh17/casp9/epha4a/hdc/anos1a/trioa/lhx9/epha7/rtn4r/cyfip2/dpysl3/draxinb/slit1b/robo3/usp33/map1sb/appb/tanc2a/dzip1/sema5a/kifbp/chad/robo1/tnr/whrnb/tenm4/hecw2a/epha4l/kif5c/slitrk4/islr2/nptnb/lhx2b/tet3/myo9ab/caprin2/kalrna/cdkl5/ephb2b/epha4b/clstn1/rac3a/brsk1b/aplp1/rab3ab/gap43/lrrc4ca/srrm4/lhx1a/map1aa/prmt8b/atl1/nkx2.2a/arhgef25b/dscamb/triob/tmem59l/myo10l3/nsmfb/slit3/cxcl8a/dpysl5a/chst11/fzd3a/kif5aa/robo4/efnb3b/vax1/stat3/dock10/ctnnd2b/zgc:77784/sema4c/si:ch211-113g11.6/rac1l/elavl4/myo10l1/ntn1b/scn8aa/lrrc4cb/plppr4a/grna/bdnf/ctnnd2a/slitrk2/nkx6.1/ncam1b/cntn3a.1/nrxn2a/dagla/grxcr1a/efna3a/dpysl5b/thsd7aa/si:ch211-241e1.3/plxna3/l1camb/plxna1a/nrcama/sema6ba/sema3b |
| GO:0007409 | axonogenesis | -2.02083525635161 | 1.14653846153846e-08 | plxnb1b/pik3cd/mycbp2/rb1/ptprfb/zgc:100846/ptprfa/isl2b/nrp2b/arhgap35a/rapgef1a/emb/pik3ca/rbpms2a/aplp2/mapk8ip3/npr1a/sema3fa/olfm1a/unc5b/nptna/vegfc/dbn1/si:ch211-129c21.1/phox2bb/ubap1/ctnna2/cntn4/cdh2/trarg1a/kalrnb/ulk2/ptk7a/dbnlb/efna1b/ephb3a/slc35b2/rbpms2b/fus/map1ab/dock7/foxb1a/nsfa/rab33ba/nrp2a/pnpla6/kctd13/isl1a/fezf2/slit2/tenm3/rufy3/nfasca/sim1a/adcy8/pak1/spg11/ret/kif1b/arnt2/sema3bl/zdhhc17/plxna4/appa/arhgap35b/efna2a/dscama/apc/robo2/pcdh17/casp9/epha4a/hdc/anos1a/trioa/lhx9/epha7/rtn4r/cyfip2/dpysl3/draxinb/slit1b/robo3/usp33/map1sb/appb/dzip1/sema5a/kifbp/chad/robo1/tnr/tenm4/epha4l/kif5c/slitrk4/islr2/nptnb/lhx2b/myo9ab/kalrna/cdkl5/ephb2b/epha4b/clstn1/rac3a/brsk1b/aplp1/rab3ab/gap43/lrrc4ca/srrm4/lhx1a/map1aa/atl1/nkx2.2a/arhgef25b/dscamb/triob/tmem59l/myo10l3/nsmfb/slit3/dpysl5a/chst11/fzd3a/kif5aa/robo4/efnb3b/vax1/stat3/zgc:77784/sema4c/si:ch211-113g11.6/rac1l/elavl4/myo10l1/ntn1b/scn8aa/lrrc4cb/plppr4a/grna/bdnf/slitrk2/nkx6.1/ncam1b/cntn3a.1/nrxn2a/dagla/efna3a/dpysl5b/thsd7aa/si:ch211-241e1.3/plxna3/l1camb/plxna1a/nrcama/sema6ba/sema3b |
| GO:0001505 | regulation of neurotransmitter levels | -2.01367130950093 | 0.000141537068751352 | snap25a/napbb/slc5a7a/napba/chata/stxbp1b/cplx2a/comtb/si:ch211-81a5.8/dbh/syn1/stx3a/caskb/snap25b/mctp2b/snap47/syt7b/syn2a/rims1b/gad1b/syt4/rab3ab/moxd1/cadpsb/syt11a/unc13a/stxbp1a/syt1a/ppfia3/gad2/syt2a/baiap3/hrh3/rph3ab/th2/stx11a |
| GO:0061564 | axon development | -1.97936421112892 | 1.14653846153846e-08 | plxnb1b/pik3cd/mycbp2/rb1/ptprfb/zgc:100846/plp1b/ptprfa/isl2b/nrp2b/arhgap35a/rapgef1a/emb/pik3ca/erbb2/rbpms2a/aplp2/mapk8ip3/npr1a/sema3fa/olfm1a/unc5b/nptna/vegfc/dbn1/gpm6bb/si:ch211-129c21.1/phox2bb/ubap1/ctnna2/cntn4/cdh2/trarg1a/kalrnb/ulk2/ptk7a/dbnlb/nfia/efna1b/ephb3a/slc35b2/rbpms2b/fus/map1ab/dock7/foxb1a/nsfa/rab33ba/nrp2a/pnpla6/kctd13/isl1a/fezf2/gpm6ba/slit2/tenm3/rufy3/nfasca/sim1a/adcy8/pak1/spg11/ret/kif1b/arnt2/sema3bl/zdhhc17/plxna4/appa/arhgap35b/efna2a/dscama/apc/robo2/pcdh17/casp9/epha4a/hdc/anos1a/trioa/lhx9/epha7/rtn4r/cyfip2/dpysl3/draxinb/slit1b/robo3/usp33/map1sb/appb/dzip1/sema5a/kifbp/chad/il6st/robo1/tnr/tenm4/epha4l/kif5c/slitrk4/islr2/nptnb/lhx2b/myo9ab/kalrna/cdkl5/ephb2b/epha4b/ek1/clstn1/rac3a/brsk1b/aplp1/rab3ab/gap43/lrrc4ca/srrm4/lhx1a/map1aa/atl1/nkx2.2a/arhgef25b/dscamb/triob/tmem59l/myo10l3/nsmfb/slit3/dpysl5a/sarm1/chst11/fzd3a/kif5aa/robo4/efnb3b/vax1/stat3/zgc:77784/sema4c/si:ch211-113g11.6/rac1l/elavl4/myo10l1/ntn1b/scn8aa/lrrc4cb/plppr4a/grna/bdnf/slitrk2/nkx6.1/ncam1b/cntn3a.1/nrxn2a/dagla/efna3a/dpysl5b/thsd7aa/si:ch211-241e1.3/plxna3/l1camb/plxna1a/nrcama/sema6ba/sema3b |
| GO:0051049 | regulation of transport | -1.97152747166447 | 1.14653846153846e-08 | prnprs3/gem/kcne4/kcna1a/unc45a/dmxl2/mffa/scn5lab/mcoln1a/pxk/abcg1/kcnj13/rab33ba/fgf13b/cep290/c2cd5/cldnb/abca1a/hecw1b/cacna1ia/kctd7/nedd4l/cplx2a/chp2/kcnab1b/si:cabz01076231.1/cacnb1/bmp2k/cacna1fb/mcu/si:ch211-204c21.1/fgf13a/msnb/kcng1/cacng5a/cacna1ha/si:ch211-81a5.8/stim2b/cnstb/syt6b/stxbp5b/c2cd2l/caskb/fxyd6/crhbp/scn1ba/kcnab2a/pacsin1a/mctp2b/fgf12a/plcb3/rab3c/htr1aa/ank3a/dnajc5aa/hecw2a/ptprn2/rab3da/cacnb2b/syt7b/cacng7b/rab3b/rims1b/shisa9a/syt9b/zgc:91999/syt4/rab3ab/kcnj3b/kcnc2/kcnj19a/sgip1b/cadpsb/scn3b/rrad/syt13/stxbp5a/syt11a/cacng2b/kcnj1a.1/syt1a/cacng3b/shisa7b/syt9a/syt10/ptprna/kcnc1b/stx1b/cacna1da/stxbp4/anxa1c/scn8aa/cacng2a/rab44/stc1l/cacna1c/cacnb4b/fgf11b/cacna2d3/kcnj14/kcnj19b/syt2a/nkx6.1/kcnd3/il1rapl1a/syt3/cabp2a/baiap3/cacna1ab/kcnc1a/rph3ab/shisa9b/kcnd2/cnih2/npvf/kcnab2b/trh/cacng8a/cacna2d2b/scn1lab/kcnj5/syt6a/cacna2d4b/kcnh4a/kcnb1 |
| GO:0032879 | regulation of localization | -1.91598630922618 | 1.14653846153846e-08 | dtnbp1b/scn4ba/cplx2b/plcg1/gpr137ba/cacng8b/ptprnb/hip1rb/kcnh7/efcab7/scn8ab/aplnrb/adcyap1b/prnprs3/gem/kcne4/kcna1a/unc45a/letm1/dmxl2/mffa/scn5lab/mcoln1a/pxk/abcg1/kcnj13/rab33ba/fgf13b/cep290/c2cd5/cldnb/abca1a/hecw1b/cacna1ia/kctd7/nedd4l/cplx2a/chp2/plin2/kcnab1b/si:cabz01076231.1/cacnb1/bmp2k/cacna1fb/mcu/si:ch211-204c21.1/fgf13a/msnb/kcng1/plch1/cacng5a/cacna1ha/si:ch211-81a5.8/stim2b/cnstb/syt6b/stxbp5b/c2cd2l/caskb/fxyd6/crhbp/scn1ba/kcnab2a/pacsin1a/mctp2b/fgf12a/plcb3/rab3c/htr1aa/ank3a/dnajc5aa/hecw2a/wnt5b/ptprn2/rab3da/cacnb2b/plce1/plch2a/syt7b/cacng7b/rab3b/rims1b/shisa9a/syt9b/zgc:91999/syt4/gpc6a/rab3ab/kcnj3b/kcnc2/kcnj19a/sgip1b/zgc:77486/cadpsb/scn3b/rrad/syt13/stxbp5a/syt11a/cacng2b/kcnj1a.1/syt1a/gpc5b/cacng3b/shisa7b/syt9a/syt10/ptprna/kcnc1b/stx1b/cacna1da/stxbp4/anxa1c/scn8aa/cacng2a/rab44/stc1l/cacna1c/cacnb4b/fgf11b/cacna2d3/kcnj14/kcnj19b/syt2a/nkx6.1/kcnd3/il1rapl1a/syt3/cabp2a/baiap3/cacna1ab/kcnc1a/rph3ab/shisa9b/kcnd2/si:ch211-13f8.1/cnih2/npvf/kcnab2b/trh/cacng8a/cacna2d2b/scn1lab/kcnj5/syt6a/cacna2d4b/kcnh4a/kcnb1 |
| GO:1901701 | cellular response to oxygen-containing compound | -1.89692649413327 | 1.56824905636127e-05 | zgc:172120/pik3r3b/myd88/carf/stat6/nr4a2b/cpeb4a/chrng/glra1/syt6b/il13/c2cd2l/phip/htr1aa/ptger4c/ticam1/htr1d/ptprn2/tet3/gnao1b/inpp5ka/syt7b/syt9b/syt4/insig1/esr2b/ptger4a/cpne4a/vtcn1/cxcl8a/cxcl8b.3/nr4a2a/syt13/syt11a/stat3/nsg2/syt1a/sstr1a/syt9a/syt10/gnao1a/ptprna/cpne2/stxbp4/adgrv1/anxa1c/nr4a3/inpp5kb/syt2a/syt3/cpne7/irs2b/drd2a/htr1fa/chrm4a/gnb5a/cpne8/syt6a/il1b/gper1/cxcl18b |
| GO:0048640 | negative regulation of developmental growth | -1.88113815885922 | 0.0105693423015173 | sema6d/sema6e/dusp6/sema3fa/si:ch211-129c21.1/ulk2/slit2/sema3bl/robo2/epha4a/wwc1/sema5a/epha4b/slit3/sema4c/si:ch211-113g11.6/wwc3/sema6ba/sema3b |
| GO:0007019 | microtubule depolymerization | -1.8750187848337 | 0.021391144294225 | map1ab/stmn4/apc/map1b/stmn2a/map1sb/stmn2b/stmn3/map1aa/camsap1a/stmn4l/apc2 |
| GO:0032409 | regulation of transporter activity | -1.86261008963063 | 0.00378678429698509 | cabp1a/cabp5a/scn4ba/cacng8b/prnprs3/gem/kcne4/fgf13b/hecw1b/kctd7/nedd4l/chp2/kcnab1b/si:cabz01076231.1/cacnb1/fgf13a/kcng1/cacng5a/stim2b/fxyd6/fgf12a/ank3a/hecw2a/cacnb2b/cacng7b/shisa9a/rrad/cacng2b/cacng3b/shisa7b/cacng2a/cacnb4b/fgf11b/cabp2a/shisa9b/cnih2/cacng8a |
| GO:0003310 | pancreatic A cell differentiation | -1.86173412990953 | 0.00748525766259099 | isl1a/pax6b/insm1b/nkx6.2/myt1b/nkx6.1/arxa |
| GO:0036269 | swimming behavior | -1.85563104344825 | 0.0118323095206783 | dmrt3a/myo9aa/sin3b/panx1b/kif2a/gdpd5a/zgc:100846/ca10a/klhl41a/spg11/scdb/micall2b/ca10b/myo9ab/cdkl5/grna/tcap/cacna1ab/dagla/syngap1b/gnb5a/scn1lab/dnmt3aa/oxt |
| GO:0019226 | transmission of nerve impulse | -1.84484111835001 | 0.0190096470355517 | cacng8b/scn8ab/cacna1ia/cacng5a/cacna1ha/ank3a/cacng7b/cacng2b/cacng3b/scn8aa/cacng2a/kcnab2b/cacng8a/scn1lab |
| GO:0021545 | cranial nerve development | -1.80828563846919 | 0.00979204145463202 | rb1/adgrg1/klf6a/hgfa/mib1/erbb2/nes/tuba1a/nsfa/six3a/slit2/kif1b/chn1/robo2/klf7b/slit1b/six3b/kifbp/diras1a/lhx2b/vax1/mdga2a/cdh4/foxi1 |
| GO:0007186 | G protein-coupled receptor signaling pathway | -1.79358601966978 | 1.60681800279223e-06 | penka/gpr27/rgs18/trpm1b/gpr78a/cart2/opn1mw2/gnb1a/gpr182/npy/chrm2a/rgrb/penkb/gng3/glrbb/adcy2a/gal/adcy6b/adcy8/adgre15/ptgir/f2rl2/rgra/grm5a/gpr22a/gnal/lpar1/or95a1/gpr35.1/s1pr1/glra1/ccl38.6/gnai2b/tmem116/drd3/oprd1b/aplnr2/gpr22b/usp33/grm4/grm6b/npffl/lmbrd2b/rgs7a/ccl34b.4/grm8b/cart3/gpr174/htr1aa/ptger4c/tafa5a/gpr61l/adgrf3b/pde6ga/htr1d/pdyn/gnb2/si:ch211-117l17.6/gnao1b/adgra2/xcr1a.1/gnb3a/ccl44/gnb4b/agtr2/rgs9b/mrap2a/gpr63/ptger4a/adra2da/opn6b/prlhr2a/adcy6a/nmur3/adcy2b/adgrl3.1/adcy1a/si:dkey-6n21.13/grin1a/grm2a/tas1r2.1/sstr1a/gabbr1b/pth2/adra2b/trpm1a/git2b/tmtops2b/gnao1a/gpr85/gpr158b/adgrv1/ccl35.1/si:dkey-172j4.3/adcy5/grm8a/tshba/adgra1a/tafa5b/grm2b/prlhr2b/gpr75/vipr1b/adgrb3/pnocb/adgre5b.2/hrh3/adrb1/rgs11/drd2a/dagla/adgrb1a/htr1fa/npb/si:ch211-132p1.2/kctd8/chrm4a/ltb4r2a/crhr1/gpr158a/npvf/ltb4r/grm1a/adcyap1r1a/pomca/adgrg3/gnb5a/rxfp1/tac3b/htr7c/adgrl1a/adgrb1b/gper1/or92a7/p2ry13 |
| GO:0010721 | negative regulation of cell development | -1.7837780436552 | 0.00979204145463202 | sema3fa/gpr137ba/si:ch211-129c21.1/irf2bp2b/spi1b/ulk2/slit2/sema3bl/tmem178a/robo2/epha4a/bmp15/sema5a/epha4b/slit3/sema4c/si:ch211-113g11.6/anxa1c/mmp9/sema6ba/gper1/sema3b |
| GO:0006935 | chemotaxis | -1.78340911781747 | 1.1812068993636e-05 | unc5b/nptna/vegfc/tnfrsf1a/si:ch211-129c21.1/cntn4/cdh2/kalrnb/maco1a/tdgf1/ptk7a/efna1b/ephb3a/slc35b2/rbpms2b/rab33ba/nrp2a/fezf2/slit2/tenm3/maco1b/nfasca/sim1a/adcy8/kif1b/arnt2/sema3bl/myd88/plxna4/efna2a/dscama/apc/robo2/pcdh17/hdc/anos1a/trioa/s1pr1/lhx9/epha7/ccl38.6/rtn4r/cyfip2/dpysl3/draxinb/slit1b/robo3/usp33/aanat2/dock4b/dzip1/trim33/ccl34b.4/sema5a/chad/vegfd/dock4/robo1/tnr/tenm4/epha4l/kif5c/islr2/nptnb/lhx2b/kalrna/xcr1a.1/ccl44/ephb2b/rac3a/gap43/lhx1a/atl1/nkx2.2a/arhgef25b/dscamb/triob/slit3/cxcl8a/dpysl5a/cxcl8b.3/chst11/fzd3a/kif5aa/si:dkey-12h9.6/robo4/efnb3b/vax1/stat3/zgc:77784/sema4c/si:ch211-113g11.6/rac1l/anxa1c/ntn1b/scn8aa/ccl35.1/cxcl8b.1/ncam1b/cntn3a.1/nrxn2a/efna3a/dpysl5b/si:ch211-241e1.3/plxna3/l1camb/plxna1a/nrcama/sema6ba/il1b/sema3b/cxcl18b |
| GO:0042330 | taxis | -1.78340911781747 | 1.1812068993636e-05 | unc5b/nptna/vegfc/tnfrsf1a/si:ch211-129c21.1/cntn4/cdh2/kalrnb/maco1a/tdgf1/ptk7a/efna1b/ephb3a/slc35b2/rbpms2b/rab33ba/nrp2a/fezf2/slit2/tenm3/maco1b/nfasca/sim1a/adcy8/kif1b/arnt2/sema3bl/myd88/plxna4/efna2a/dscama/apc/robo2/pcdh17/hdc/anos1a/trioa/s1pr1/lhx9/epha7/ccl38.6/rtn4r/cyfip2/dpysl3/draxinb/slit1b/robo3/usp33/aanat2/dock4b/dzip1/trim33/ccl34b.4/sema5a/chad/vegfd/dock4/robo1/tnr/tenm4/epha4l/kif5c/islr2/nptnb/lhx2b/kalrna/xcr1a.1/ccl44/ephb2b/rac3a/gap43/lhx1a/atl1/nkx2.2a/arhgef25b/dscamb/triob/slit3/cxcl8a/dpysl5a/cxcl8b.3/chst11/fzd3a/kif5aa/si:dkey-12h9.6/robo4/efnb3b/vax1/stat3/zgc:77784/sema4c/si:ch211-113g11.6/rac1l/anxa1c/ntn1b/scn8aa/ccl35.1/cxcl8b.1/ncam1b/cntn3a.1/nrxn2a/efna3a/dpysl5b/si:ch211-241e1.3/plxna3/l1camb/plxna1a/nrcama/sema6ba/il1b/sema3b/cxcl18b |
| GO:0098953 | receptor diffusion trapping | -1.77120547755989 | 0.0221781634063151 | cacng8b/gphnb/cacng5a/gphna/cacng7b/cacng2b/cacng3b/cacng2a/cacng8a |
| GO:0098970 | postsynaptic neurotransmitter receptor diffusion trapping | -1.77120547755989 | 0.0221781634063151 | cacng8b/gphnb/cacng5a/gphna/cacng7b/cacng2b/cacng3b/cacng2a/cacng8a |
| GO:0099628 | neurotransmitter receptor diffusion trapping | -1.77120547755989 | 0.0221781634063151 | cacng8b/gphnb/cacng5a/gphna/cacng7b/cacng2b/cacng3b/cacng2a/cacng8a |
| GO:0071867 | response to monoamine | -1.76002500271274 | 0.0460556112254136 | htr1aa/htr1d/gnao1b/nsg2/gnao1a/drd2a/htr1fa/gnb5a |
| GO:0071869 | response to catecholamine | -1.76002500271274 | 0.0460556112254136 | htr1aa/htr1d/gnao1b/nsg2/gnao1a/drd2a/htr1fa/gnb5a |
| GO:0051960 | regulation of nervous system development | -1.75557639124564 | 0.00248666027456893 | nlk1/si:zfos-741a10.3/plxnb1b/hes6/neurod6a/adnpb/sema3fa/btbd6a/her12/dbn1/si:ch211-129c21.1/lef1/ulk2/dbnlb/ephb3a/dhrs12la/slit2/ctbp2a/sema3bl/plxna4/clstn2b/bhlhe40/robo2/epha4a/neurod4/gata3/her13/gbx2/tnfrsf21/sema5a/foxo3a/tnr/tenm4/slitrk4/caprin2/zbtb16a/cdkl5/epha4b/clstn1/nsmfb/slit3/clstn2a/elavl3/stat3/sema4c/si:ch211-113g11.6/gbx1/grna/bdnf/clstn3/slitrk2/dmbx1a/plxna3/l1camb/mmp9/plxna1a/sema6ba/sema3b |
| GO:0097120 | receptor localization to synapse | -1.75544036908873 | 0.0351674145187552 | dlg1b/cacng8b/grip2b/gphnb/cacng5a/stx3a/anks1b/kif5c/gphna/cacng7b/kif5aa/cacng2b/cacng3b/stx1b/cacng2a/rab44/cacng8a |
| GO:0031532 | actin cytoskeleton reorganization | -1.74088656656218 | 0.0402190977319739 | klf6a/anxa1a/baiap2a/sipa1l1/mink1/nedd9/thsd7bb/carmil2/parvg/carmil3/plek/thsd7ba/anxa1c/plek2/phactr1/thsd7aa |
| GO:0040013 | negative regulation of locomotion | -1.73929262815316 | 0.018759055936431 | sema3fa/lrch1/si:ch211-129c21.1/lbx2/prox1a/miip/lef1/phf8/srgap3/sema3bl/robo2/anos1a/rap2b/sema5a/srgap1b/sinhcaf/arhgap4b/sema4c/si:ch211-113g11.6/rtn4rl2a/sema6ba/sema3b |
| GO:0031110 | regulation of microtubule polymerization or depolymerization | -1.72739295421889 | 0.0437181181328161 | ska3/cdk5rap2/stmn1b/map1sa/camsap2a/camsap1b/slain2/map1ab/traf3ip1/zgc:66447/clasp2/mapre3b/stmn4/apc/map1b/stmn2a/map1sb/stmn2b/stmn3/clip1a/map1aa/camsap1a/stmn4l/apc2 |
| GO:0040011 | locomotion | -1.71881331498307 | 4.74228301214316e-06 | plcg1/unc5b/nptna/lrch1/vegfc/tnfrsf1a/si:ch211-129c21.1/lbx2/cntn4/cdh2/prox1a/kalrnb/miip/maco1a/lef1/tdgf1/foxj1a/ptk7a/efna1b/ephb3a/slc35b2/gnb1a/rbpms2b/sst7/slkb/rab33ba/nrp2a/fezf2/phf8/slit2/tenm3/maco1b/srgap3/nfasca/sim1a/adcy8/kif1b/arnt2/sema3bl/myd88/plxna4/efna2a/dscama/apc/robo2/pcdh17/hdc/asap1a/arhgef39/anos1a/trioa/s1pr1/lhx9/epha7/hace1/peak1/ccl38.6/rtn4r/cyfip2/dpysl3/draxinb/gata3/slit1b/robo3/usp33/tek/aanat2/dock4b/dzip1/rap2b/trim33/ccl34b.4/sema5a/chad/vegfd/pdgfbb/dock4/robo1/tnr/tenm4/epha4l/kif5c/islr2/nptnb/htr1d/lhx2b/kalrna/xcr1a.1/ccl44/ephb2b/sst1.1/srgap1b/rac3a/gap43/lhx1a/atl1/nkx2.2a/arhgef25b/dscamb/amot/triob/nsmfb/slit3/cxcl8a/dpysl5a/cxcl8b.3/chst11/fzd3a/kif5aa/si:dkey-12h9.6/sinhcaf/robo4/efnb3b/vax1/arhgap4b/stat3/dock10/zgc:77784/sema4c/adam8a/si:ch211-113g11.6/rac1l/anxa1c/ntn1b/scn8aa/ccl35.1/cxcl8b.1/mdga2a/zgc:153631/asap1b/ncam1b/cntn3a.1/nrxn2a/klhl40b/efna3a/chrm4a/dpysl5b/si:ch211-241e1.3/plxna3/l1camb/plxna1a/nrcama/rtn4rl2a/sema6ba/il1b/sema3b/cxcl18b |
| GO:0006959 | humoral immune response | -1.69323952530372 | 0.0594675488145613 | c4/tnfrsf21/c1qa/tfe3a/cfb/cxcl8b.3/c6/susd4/c7b/c4b/c3a.2/itln2/cxcl18b |
| GO:0007420 | brain development | -1.67227058783588 | 2.31796694375566e-05 | her12/wnt3/macroh2a2/marcksa/kctd12.1/daam1a/pknox1.1/tgif1/ctnna2/cntn4/cdh2/adcyap1b/prox1a/zfhx3b/prnprs3/sbno2a/lef1/tdgf1/ulk2/tsc22d3/bag6l/pou3f3a/atg4da/creb1a/pafah1b1a/birc6-as2/sox14/foxb1a/nsfa/zfhx3a/kdm6al/slc2a3a/cep290/kctd13/fezf2/phf8/six3a/nfasca/sim1a/pax6b/arnt2/raraa/grhl3/tbr1b/kirrel3l/cars2/ptprsa/kdm7ab/nr4a2b/ralgapa1/apc/epha4a/hdc/anos1a/lhx9/neurod4/pcdh10b/pbx1a/mvp/six3b/mdkb/si:ch73-119p20.1/gbx2/dzip1/lgi1a/pbx1b/epb41l5/auts2a/pbx3b/pcare1/hecw2a/znf503/meis2a/wnt5b/lhx6a/irx7/lhx2b/brf1b/rbfox1/tcf7l2/nkx2.1/pcdh19/srrm4/lhx1a/asphd1/pou4f2/prmt8b/tbxta/aatkb/rtn1b/sp5l/nr2e3/tbcd/has2/lhx5/fzd3a/pou3f2b/nr4a2a/efnb3b/ndrg4/vax1/pax2a/git2b/gbx1/ntn1b/pou3f2a/bdnf/atp1a3a/sez6l2/mdga2a/rx3/dmbx1a/otpb/six6a/cdk5r2a/arxa/nkx2.4a/syngap1b/gnrh2/l1camb/cnnm2a/otpa/kcnb1/nrcama/gper1/bicdl1 |
| GO:0060322 | head development | -1.66225293057786 | 4.42238345198842e-05 | her12/wnt3/macroh2a2/marcksa/kctd12.1/daam1a/pknox1.1/tgif1/ctnna2/cldn7a/cntn4/cdh2/adcyap1b/prox1a/zfhx3b/prnprs3/sbno2a/lef1/tdgf1/ulk2/tsc22d3/bag6l/pou3f3a/atg4da/creb1a/pafah1b1a/birc6-as2/sox14/foxb1a/nsfa/zfhx3a/kdm6al/slc2a3a/cep290/kctd13/fezf2/phf8/six3a/nfasca/sim1a/pax6b/arnt2/raraa/grhl3/tbr1b/kirrel3l/cars2/ptprsa/kdm7ab/nr4a2b/ralgapa1/apc/epha4a/hdc/wdr11/anos1a/lhx9/neurod4/pcdh10b/pbx1a/mvp/six3b/mdkb/si:ch73-119p20.1/gbx2/dzip1/lgi1a/pbx1b/epb41l5/auts2a/pbx3b/pcare1/hecw2a/znf503/meis2a/wnt5b/lhx6a/irx7/lhx2b/brf1b/rbfox1/tcf7l2/nkx2.1/pcdh19/srrm4/lhx1a/asphd1/pou4f2/prmt8b/tbxta/aatkb/rtn1b/sp5l/nr2e3/tbcd/has2/lhx5/fzd3a/pou3f2b/nr4a2a/efnb3b/ndrg4/vax1/pax2a/git2b/gbx1/ntn1b/chd5/pou3f2a/bdnf/atp1a3a/sez6l2/mdga2a/specc1/rx3/dmbx1a/otpb/six6a/cdk5r2a/arxa/nkx2.4a/syngap1b/gnrh2/l1camb/cnnm2a/otpa/kcnb1/nrcama/gper1/bicdl1 |
| GO:0045732 | positive regulation of protein catabolic process | -1.66175262841073 | 0.0607582432020156 | oaz1a/csnk1db/tmem259/oaz2b/socs5b/prkn/apc/rnf217/si:ch211-278j3.3/trib1/fzr1b/rnf144ab/csnk1e/adam8a/klhl40b/apc2/rnf144aa |
| GO:1901797 | negative regulation of signal transduction by p53 class mediator | -1.65413798029984 | 0.0829144295829561 | cd74b/cd74a/bcor/vax1/bcl6aa |
| GO:0006688 | glycosphingolipid biosynthetic process | -1.64952029857802 | 0.097836707744085 | b4galt6/b4galt3/st6galnac3 |
| GO:0031644 | regulation of nervous system process | -1.64521543139582 | 0.0906720571933235 | dlgap1a/tnfrsf21/tenm4/dlgap4b/si:dkey-174m14.3/kcnab2b/dlgap3/neto1l |
| GO:0060560 | developmental growth involved in morphogenesis | -1.64279725335724 | 0.0128583805736124 | plxnb1b/tmem108/pik3cd/mycbp2/spag6/cbsb/smad2/pik3ca/zeb2a/mapk8ip3/sema3fa/dbn1/si:ch211-129c21.1/ubap1/cdh2/trarg1a/tmem67/ulk2/ptk7a/dbnlb/creb1a/fus/isl1a/slit2/ppp1cb/spg11/ret/kif1b/sema3bl/bbs1/robo2/pcdh17/casp9/epha4a/robo3/appb/sema5a/wnt5b/myo9ab/cdkl5/epha4b/clstn1/tbxta/zeb2b/tmem59l/myo10l3/nsmfb/slit3/stat3/sema4c/si:ch211-113g11.6/myo10l1/ntn1b/grna/bdnf/nrn1la/nkx6.1/nrxn2a/thsd7aa/plxna3/l1camb/sema6ba/sema3b |
| GO:0016049 | cell growth | -1.63666909504427 | 0.0141269580867656 | plxnb1b/tmem108/pik3cd/mycbp2/spag6/ewsr1a/pik3ca/mapk8ip3/sema3fa/dbn1/si:ch211-129c21.1/ubap1/trarg1a/ulk2/dbnlb/supv3l1/dnph1/fus/isl1a/slit2/spg11/ret/kif1b/sema3bl/robo2/pcdh17/casp9/epha4a/robo3/appb/sema5a/wnt5b/myo9ab/cdkl5/epha4b/clstn1/tmem59l/myo10l3/nsmfb/slit3/sema4c/si:ch211-113g11.6/myo10l1/ntn1b/epb41l3a/grna/bdnf/nrn1la/nkx6.1/nrxn2a/thsd7aa/plxna3/l1camb/sema6ba/sema3b |
| GO:0034643 | establishment of mitochondrion localization. microtubule-mediated | -1.63631487834659 | 0.0931198545493737 | rhot1b/trak1a/sybu/kif5aa/rhot2 |
| GO:0047497 | mitochondrion transport along microtubule | -1.63631487834659 | 0.0931198545493737 | rhot1b/trak1a/sybu/kif5aa/rhot2 |
| GO:0051654 | establishment of mitochondrion localization | -1.63631487834659 | 0.0931198545493737 | rhot1b/trak1a/sybu/kif5aa/rhot2 |
| GO:0032102 | negative regulation of response to external stimulus | -1.63570938930065 | 0.0540332674423811 | sema3bl/gramd4b/robo2/epha4a/il13/nr1h5/sema5a/cart3/ptger4c/irak3/epha4b/gramd4a/ptger4a/sema4c/si:ch211-113g11.6/susd4/grna/serpine1/sc:d189/sema6ba/sema3b |
| GO:0048885 | neuromast deposition | -1.63446228315307 | 0.0960340406478379 | lbx2/prox1a/lef1/phf8/anos1a/rtn4rl2a |
| GO:0071407 | cellular response to organic cyclic compound | -1.62675574144756 | 0.0543657683428525 | abhd2b/htr1aa/htr1d/gnao1b/insig1/esr2b/cxcl8a/nsg2/sstr1a/gnao1a/anxa1c/drd2a/htr1fa/gnb5a/gper1 |
| GO:0022603 | regulation of anatomical structure morphogenesis | -1.62242060545465 | 0.00196692398869123 | plcd4a/vegfab/stab1/hif1ab/amotl2b/sema6d/ptenb/alcama/sema6e/ptk2ab/thbs2b/brwd1/rspo3/tspan18a/gata2a/zeb1b/plxnb1b/pik3cd/disc1/mia3/anxa1a/fn1b/arhgap35a/pik3ca/flot2a/sema3fa/camsap1b/plcg1/unc5b/vegfc/dbn1/aplnrb/kctd12.1/si:ch211-129c21.1/rhogd/zeb1a/lbx2/cdh2/fmnl2b/ulk2/elmo1/marchf5/ptk7a/dbnlb/dab2ipb/lurap1/ephb3a/mffa/sipa1l1/tnika/vat1/palm1b/slit2/mief2/rufy3/hecw1b/pak1/sema3bl/plxna4/nedd4l/arhgap35b/robo2/casp9/epha4a/s1pr1/msnb/tanc2a/phip/sema5a/vegfd/tenm4/hecw2a/wnt5b/caprin2/cdkl5/epha4b/myo10/clstn1/rac3a/lrrc4ca/lhx1a/nsmfb/slit3/basp1/stat3/prkd1/sema4c/si:ch211-113g11.6/rac1l/gbx1/anxa1c/lrrc4cb/hif1aa/bdnf/serpine1/adgrb3/scg2b/gdf3/adgrb1a/thbs1a/dab2ipa/plxna3/l1camb/eve1/plxna1a/adgrb1b/sema6ba/sema3b |
| GO:0000381 | regulation of alternative mRNA splicing. via spliceosome | -1.62121505428978 | 0.0619371464051369 | hnrnpa0a/rbm38/rbm25a/celf2/rbfox3a/rbfox3b/celf6/rbfox1l/rbfox1/srrm4/nova2/celf3a/khdrbs2/celf5a/celf4 |
| GO:0001933 | negative regulation of protein phosphorylation | -1.61410605566215 | 0.0735990755145654 | spry4/aida/dtnbp1b/ptprc/ldlrad4a/camk2n2/cbl/camk2n1a/cblc/prkn/sh3bp5la/sh3bp5b/si:dkey-175m17.7/ibtk/camk2n1/irak3/inpp5ka/cadm4/inpp5kb/smcr8b |
| GO:0008361 | regulation of cell size | -1.59323427771673 | 0.0658674351521172 | arhgap35a/pik3ca/sema3fa/arhgap5/dbn1/si:ch211-129c21.1/ulk2/dbnlb/zgc:153039/slit2/sema3bl/arhgap35b/robo2/casp9/epha4a/sema5a/piezo1/cdkl5/epha4b/clstn1/nsmfb/slit3/si:ch211-178n15.1/stat3/slc12a5b/sema4c/si:ch211-113g11.6/ntn1b/slc12a5a/bdnf/nkx6.1/plxna3/l1camb/sema6ba/sema3b |
| GO:0043087 | regulation of GTPase activity | -1.57086875382947 | 0.0297635976408028 | tbc1d9/rapgef1a/flcn/sipa1l2/rasgef1ba/iqsec1b/ralgapa2/rasa4/gpr137ba/rgl1/nf1a/usp6nl/adcyap1b/vav2/agap1/rapgef5b/rapgef2a/si:dkey-91m11.5/dab2ipb/rgl3a/ephb3a/rapgef2b/dock7/rapgef4b/sipa1l1/gapvd1/rasal2/plxna4/chn1/ralgapa1/ccpg1/asap1a/rasgef1bb/ccl38.6/bcr/rgs7a/ccl34b.4/si:ch211-288d18.1/evi5l/tbc1d30/elmod1/vav3b/myo9ab/ccl44/ek1/rgs9b/sgsm3/syngap1a/arap3/rap1gapa/unm_sa1614/ralgps1/dock10/agap2/ccl35.1/asap1b/rasgrf2b/rgs11/syngap1b/dab2ipa/plxna3/rasgrf1/plxna1a/arhgap25 |
| GO:0002252 | immune effector process | -1.52782636998837 | 0.0607582432020156 | cd74b/c4/c1qa/ticam1/cd74a/cfb/cadm1b/cxcl8a/c6/si:ch211-196f2.6/susd4/anxa1c/zgc:153631/c7b/c4b/zgc:123107/c3a.2/csf2rb/mmp9/mhc2d4b |
| GO:0001755 | neural crest cell migration | -1.51280994298295 | 0.0960340406478379 | hif1ab/sema6d/pax6a/med13b/sema6e/lrp5/disc1/nrp2b/erbb2/pdgfrb/sema3fa/radil/si:ch211-129c21.1/cdh2/lef1/dmxl2/pax3a/nrp2a/fscn1a/pak1/ret/pax6b/sema3bl/mmp17b/wdr11/sema5a/zeb2b/sh3pxd2aa/sema4c/si:ch211-113g11.6/rx3/foxi1/sema6ba/sema3b |
| GO:0051240 | positive regulation of multicellular organismal process | -1.44230693231483 | 0.0657742225763037 | gata2a/slc27a1b/nlk1/si:zfos-741a10.3/plxnb1b/anxa1a/avpr1ab/neurod6a/tnfsf13b/nr2c2/cactin/apln/tlr5b/olfm1a/vegfc/dbn1/aplnrb/lef1/elmo1/dbnlb/lurap1/ephb3a/npy/tapt1b/dhrs12la/loxl2a/zgc:172120/plxna4/clstn2b/cd74b/neurod4/gata3/aanat2/vegfd/enpp5/tenm4/ticam1/slitrk4/caprin2/zbtb16a/cdkl5/clstn1/cd74a/lhx1a/vtcn1/nsmfb/clstn2a/basp1/stat3/si:ch211-196f2.6/anxa1c/grna/bdnf/zbtb16b/clstn3/slitrk2/zgc:153631/adrb1/npvf/plxna3/l1camb/zgc:123107/plxna1a/panx2/gper1/mhc2d4b |
| GO:0030334 | regulation of cell migration | -1.43156853115565 | 0.0696903189146766 | plcg1/lrch1/vegfc/si:ch211-129c21.1/lbx2/prox1a/miip/lef1/gnb1a/sst7/slkb/phf8/srgap3/sema3bl/plxna4/asap1a/arhgef39/anos1a/s1pr1/hace1/peak1/dpysl3/gata3/tek/rap2b/sema5a/vegfd/pdgfbb/sst1.1/srgap1b/amot/nsmfb/sinhcaf/robo4/arhgap4b/dock10/sema4c/adam8a/si:ch211-113g11.6/anxa1c/mdga2a/zgc:153631/asap1b/si:ch211-241e1.3/plxna3/plxna1a/rtn4rl2a/sema6ba/sema3b |
| GO:0009719 | response to endogenous stimulus | -1.32546647522578 | 0.0666420813824336 | tbx16/irs1/skia/igfbp3/pik3r3a/gsk3ba/rbm14b/gna11b/slc39a14/slc27a1b/esr2a/zyx/smad7/tll1/ltbp3/tacr1a/spry4/dusp6/trim71/anxa1a/avpr1ab/nrp2b/nr1d1/smad2/grb10a/flcn/smurf1/zeb2a/asic4b/fcho1/skib/rgmb/fgfbp1a/irs2a/drd4-rs/nptna/vegfc/fstl4/si:rp71-46j2.7/ldb2a/timp2a/tgif1/ldlrad4a/caly/fstl3/castor2/ppp4cb/zdhhc13/tsc22d3/il13ra1/dab2ipb/lifrb/golm1/rerea/chrm2a/slc2a3a/nrp2a/pnpla6/asip1/tcf21/grem2b/pik3r3b/raraa/thrb/zdhhc17/bmpr1ab/smad3b/stat6/gpr22a/nr4a2b/cpeb4a/chrng/glra1/sall1a/gata3/bmp15/dzip1/fabp4b/phip/crhbp/sp9/snx25/inhbab/vegfd/fgf12a/abhd2b/htr1aa/bmpr2a/scube1/ptger4c/ccbe1/stat1b/nrros/htr1d/lemd3/gnao1b/inpp5ka/smad6b/insig1/esr2b/ptger4a/atl1/tbxta/inhbaa/asic2/timp4.3/has2/adgrl3.1/kcp/grem1a/nr4a2a/bmpr2b/stat3/nsg2/wbp2/gnao1a/stxbp4/anxa1c/nr4a3/bdnf/elapor2b/inpp5kb/ntrk3a/frs3/irs2b/gdf3/drd2a/ntrk3b/htr1fa/chrm4a/trh/dio2/dab2ipa/gnb5a/rxfp1/c3a.2/sostdc1b/mmp9/timp2b/hpxa |

**Table S4.** Enriched Kyoto Encyclopedia of Genes and Genomes (KEGG) pathways and their associated genes following M1 exposure, identified by gene set enrichment analysis (GSEA).

| ID | Description | NES | FDR (p adjusted value) | Genes implicated |
| --- | --- | --- | --- | --- |
| dre03010 | Ribosome | 2.39091223023189 | 6.16666666666667e-09 | zgc:158463/mrpl49/mrpl10/mrps5/mrpl4/mrps33/mrps18b/rpl26/mrpl27/mrpl42/mrpl40/mrpl39/mrpl14/mrpl1/rpl12/mrpl35/mrpl17/mrps18c/rps25/mrpl44/mrps12/rps7/rpl29/mrpl30/rpl38/mrps26/rps21/mrpl23/rpl32/rpl11/mrpl13/mrpl9/mrpl48/mrps28/mrps25/mrpl12/rpl34/mrps2/rps17/rps8a/mrps27/mrpl51/mrpl47/mrpl36/mrpl18/rps15a/mrps22/rpl22l1/mrpl33/mrps10/mrps31/rps3a/rps24/rpl36a/rpl14/zgc:171772/rpl35/rpl37/faub/rps11/rpl5a/zgc:171480/rpl13a/mrpl34/uba52/mrpl11/mrpl2/mrps14/rpl22/mrpl46/mrps35/rpl7/rpl24/mrpl24/chchd1/mrpl41/rpl6/mrpl54/rps18/rpl35a/mrps6/rpl30/mrps21/rpl39/rps29/rps27a/mrpl20/mrps16/mrps17/rpl23a/mrpl53/rpl21/rpl36/mrpl3/rpl5b/mrpl22/rpl27/rplp1/rpl23/mrpl15/rps15/rpsa/rps14/mrpl19/rpl17/mrpl45/mrpl32/rsl24d1/rps20/rpl19/rpl15/mrps15/rplp2/rpl4/rps27.1/rack1/mrpl43/rps16/rpl31/rps26/rpl10/rpl28/faua/mrpl37/rps10/mrps24/rpl18/rps5/rps27l/mrps11/rpl7a/mrpl52/rpl9/rps11b/rplp2l |
| dre00860 | Porphyrin metabolism | 2.09426267112894 | 0.000319865137159646 | fthl29/fthl28/fthl31/blvra/fthl30/ugt5b1/ppox/ugt5c2/urod/hmox1a/hccsa.1/hmox2a/ugt5g1/blvrb/fech/hephl1a/ugt5e1/fxn/fthl27/ugt5a2/mmab/ugt5b4 |
| dre00120 | Primary bile acid biosynthesis | 2.0733417350763 | 0.000664048141002421 | amacr/ch25hl1.1/cyp46a1.2/cyp7a1b/cyp39a1/ch25h/cyp46a1.4/cyp46a1.3/cyp27a1.1/cyp46a1.1/cyp27a3 |
| dre03008 | Ribosome biogenesis in eukaryotes | 1.95637425837524 | 0.00114869587315338 | nmd3/zgc:158463/nxt2/imp3/zgc:86811/tbl3/smfn/mphosph10/gnl3/snu13b/rpp30/mdn1/snu13a/wdr3/nol6/rpp40/pwp2h/wdr36/sbds/lsg1/heatr1/riok2/nop58/emg1/gnl2/fcf1/rrp7a/nop56/wdr75/si:dkey-251i10.3/bms1/nat10/rexo1/gar1/nxf1b/dkc1/fbl/nop10/utp4 |
| dre00983 | Drug metabolism - other enzymes | 1.90714576252407 | 0.00114869587315338 | rrm2b/dpyda.1/ugt5b1/tk2/ugt5c2/zgc:103601/gsta.2/nme4/tpmt.2/gstp1.2/ugt5g1/si:dkey-78a14.4/gstm.2/ugt5e1/dut/nme2a/gstt2/zgc:103586/uckl1a/tpmt.1/cdab/gstt1a/ugt5a2/hprt1/itpa/impdh2/ugt5b4/rrm1/rrm2/upp2/nme7/nme3/uckl1b/gstt1b/cdaa/nme2b.2/mgst2/xdh/uck1/gstm.1 |
| dre04216 | Ferroptosis | 1.90274562050419 | 0.00304766869057691 | fthl29/fthl28/fthl31/fthl30/tfr1b/map1lc3c/slc3a2b/hmox1a |
| dre00100 | Steroid biosynthesis | 1.85355866469734 | 0.0183999269399779 | dhcr24/nsdhl/sc5d/soat2/msmo1/cyp24a1/cel.2/ebp/cyp51/lss/cel.1/lipf/hsd17b7 |
| dre04981 | Folate transport and metabolism | 1.81174568388793 | 0.0336508116431211 | abcc4/abcg2a/abcc6b.2/abcg2b/slc22a6l/wu:fb13g09/abcc1/mthfs/mtr/mthfd1a/zgc:172121/dhfr/bhmt/folr/mthfr/tyms |
| dre00533 | Glycosaminoglycan biosynthesis - keratan sulfate | 1.76217978284825 | 0.0400906255215409 | b4galt3/si:dkey-160o24.3/b4galt1l/b4galt4 |
| dre03440 | Homologous recombination | 1.75475710765219 | 0.0400906255215409 | blm/nbn/eme1/rbbp8/palb2/rad54l/rad50/mus81/abraxas1/pold3/brip1/rpa3/bard1/mre11a/rad51d/rad51b/sem1/atm/rpa2/pold4/rad54b/rad51c |
| dre00240 | Pyrimidine metabolism | 1.74852490158296 | 0.015407731275394 | rrm2b/dpyda.1/entpd6/tk2/nt5c2b/nme4/nt5c2l1/entpd8/entpd3/dctd/dut/nme2a/zgc:103586/uckl1a/entpd1/cdab/rrm1/rrm2/upp2/dctpp1/zgc:110540/nme7/nme3/nudt2/uckl1b/dhodh/zgc:136858/cdaa/nme2b.2/uck1/tyms/ctps1a/nt5c1aa/nme6 |
| dre00970 | Aminoacyl-tRNA biosynthesis | 1.74532545298755 | 0.0400906255215409 | fars2/farsa/dars1/aars2/dars2/mtfmt/qars1/nars2/yars1/tars1/yars2/qrsl1/pars2/wars2/gars1/hars/lars1b/rars1/vars1/gatc/sars1/mars1/iars2/cars1 |
| dre00500 | Starch and sucrose metabolism | 1.73336246829957 | 0.0400906255215409 | gys2/gyg2/gpib/gba3/hk2/amy2al1/pgm2l1/g6pc1a.1/si/pygb/amy2a/g6pc3/aglb/gbe1a/amy2al2 |
| dre00670 | One carbon pool by folate | 1.67677064432588 | 0.0661919744710782 | dmgdh/aldh1l2/mat2aa/gnmt/gart/mthfs/mtr/mthfd1a/mat1a/zgc:172121/ahcy/dhfr/bhmt/atic/gcshb/mthfr/cbsa/aldh7a1/tyms/cth/shmt1 |
| dre01240 | Biosynthesis of cofactors | 1.60930418218394 | 0.015407731275394 | ido1/si:dkey-94e7.2/ugt5b1/ppox/ugt5c2/urod/bco1l/gch2/zgc:86811/LOC103911725/adsl/akr1a1b/mat2aa/nme4/coq4/tdo2b/ak8/akr1a1a/flad1/ugt5g1/rgn/aldh2.2/si:ch211-107o10.3/fech/coq7/adss2/coq5/ugt5e1/nme2a/ppcs/gch1/coq2/rfk/dhrs3a/nqo1/mthfd1a/ugt5a2/si:dkey-23o4.6/lipt1/mmab/mat1a/nfs1/ugt5b4/hpdb/psat1/nadka/pmm2/nme7/nme3/hmbsa/bco1/dhfr/dhodh/coq6/hmbsb/aldh3a2a/nme2b.2/ggh/ak3/pnpo/coq3/ctps1a/alpi.1/spra/tdo2a/nme6/uros/alas2/gss/gusb/shmt1/ugt5b2/naprt/adss1/zgc:112332 |
| dre01232 | Nucleotide metabolism | 1.56350519578115 | 0.0400906255215409 | rrm2b/entpd6/tk2/zgc:86811/nt5c2b/adsl/nme4/dguok/nt5c2l1/ak8/guk1b/pnp6/adka/entpd8/entpd3/adss2/dctd/dut/nme2a/zgc:103586/uckl1a/entpd1/pnp5b/cdab/hprt1/itpa/impdh2/rrm1/rrm2/upp2/dctpp1/zgc:110540/nme7/nme3/gda/uckl1b/cdaa/nme2b.2/xdh/uck1/ak3/tyms/ctps1a/impdh1b/aprt/nt5c1aa/nme6 |
| dre00190 | Oxidative phosphorylation | 1.46009325668473 | 0.0779574444123974 | sdhda/ppa1a/cox11/ATP8/si:dkey-85n7.8/atp5mea/cox17/ndufa3/cox7b/atp6v1f/ndufc1/atp6v0b/ndufv1/cycsb/atp5mf/atp6ap1b/atpv0e2/sdhdb/atp6v1e1a/ndufb8/atp5mc3a/uqcrc2b/atp5meb/ndufb6/ndufa10/atp5po/cox7c/atp5f1e/ndufa8/cox5b2/cox6b2/atp5pd/cox7a1/ndufb7/atp5mc1/cox6c/ndufs3/ndufs4/cox7a2a/sdhc/cox4i1/ndufa7/uqcr10/sdha/ndufa6/uqcrc1/cox5ab/ndufa4b/atp5f1c/uqcrb/ndufa2/uqcrh/ndufs8a/ND3/ndufs6/ndufab1a/COX2/atp6v0e1/ndufs2/ndufa12/ndufb4/atp6v1c2/atp5pb/cox7a2l/cox8b/sdhb/uqcrq/cox6a1/ndufa5/ndufb5/ND5/si:ch1073-325m22.2/ndufb10/atp5mc3b/atp6v1c1b/ndufb9/ndufc2/ndufab1b/uqcrfs1/ndufa1/uqcrc2a/ndufa9a/atp5pf/atp6v1e1b/cox4i1l/ndufs5/CYTB/cox5aa/atp6v1h/atp6v0a1a/atp6v1g1/ndufa11/ATP6/cyc1/ND4L/ppa2 |
| dre04082 | Neuroactive ligand signaling | -2.39461733568973 | 6.16666666666667e-09 | penkb/si:dkey-206f10.1/chrna3/si:ch211-132f19.7/adcy6a/p2rx3b/slc17a9b/prkacab/chata/gnb1b/adcy1a/htr1fa/grm6b/gng13b/slc1a8a/slc18a3a/gabra1/slc6a5/grm4/gnb1a/gabrb4/gabra4/LOC101886206/gria3b/gnai2b/adrb2b/p2ry1/hrh3/gabrb3/gabbr1a/grin1a/glra3/htr1d/penka/grik2/chrna11/htr1aa/gabrb2a/grin2aa/grin2da/gnal/pdyn/adra2da/grm8b/adcy2b/grin3bb/gnb3a/gria2b/grm5a/drd4a/adcy8/gnb2/LOC100149482/htr7c/grin2ab/slc17a6b/grm2a/htr3a/gnao1b/gnao1a/glra1/pomca/gabrg2/slc32a1/grik5/chrna4b/grin1b/gria1a/slc6a1b/chrnb2/slc6a1a/gabbr1b/LOC101882451/pnocb/oprd1b/si:dkey-247m21.3/grm2b/oprl1/grik1a/grin2bb/gabrr2a/gabrb1/slc1a2b/chrm4a/tac3b/grm1a/grm8a/gria4b/p2ry13 |
| dre04080 | Neuroactive ligand-receptor interaction | -2.39295294436978 | 6.16666666666667e-09 | gabrb4/gabra4/sst1.1/gria3b/sstr1a/adrb2b/p2ry1/hrh3/gabrb3/crhb/gabbr1a/f2rl2/grin1a/f2rl1.1/glra3/pth2/htr1d/penka/uts2d/grik2/chrna11/htr1aa/chrna5/gabrb2a/grin2aa/grin2da/pdyn/rxfp1/adra2da/grm8b/grin3bb/grid1a/gria2b/chrnb3a/cckbra/grm5a/drd4a/LOC100149482/s1pr1/htr7c/grin2ab/grm2a/glra1/pomca/gpr156/vipr1b/gabrg2/grik5/chrna4b/grin1b/gria1a/chrng/ltb4r/sstr1b/mtnr1c/chrnb2/gabbr1b/LOC101882451/pnocb/oprd1b/uts1/si:dkey-247m21.3/npb/grm2b/oprl1/prlhr2a/grik1a/grin2bb/gabrr2a/ltb4r2a/gabrb1/trh/chrm4a/nmur3/adcyap1r1a/vgf/oxt/tac3b/npvf/grm1a/rln3b/mchr1b/si:dkey-148a17.6/rln3a/grm8a/c3a.2/ptger4c/gria4b/tshba/lepa/p2ry13 |
| dre04310 | Wnt signaling pathway | -1.79169862488582 | 0.000378862952474737 | senp2/cul1a/ctnnb1/wnt4b/prkcbb/nlk1/vangl2/fbxw11b/csnk2b/tbl1xr1a/csnk2a1/tle3b/dkk2/bambib/crebbpa/csnk2a2b/zgc:136929/ccdc88c/chd8/ankrd6b/ctbp1/daam1a/tbl1xr1b/camk2g1/wnt6b/gsk3ba/ep300b/prkacab/tle2a/camk2b1/celsr1b/tcf7l2/mapk10/wnt7bb/cul1b/rac3a/rspo1/ppp3cca/camk2a/ctnnd2b/ppp3r1b/csnk1e/mycb/camk2g2/tle3a/LOC101886206/fosl1a/apc/wnt16/ctbp2a/ctnnd2a/wnt3/fzd8a/lef1/fzd8b/fzd3a/ror1/prickle2b/camk2d2/psen1/ppp3ca/apc2/sox17/prickle2a/nfatc4/ppp3cb/rac1l |
| dre04514 | Cell adhesion molecule (CAM) interaction | -1.67001284519405 | 0.0101775110712772 | cldn5b/si:busm1-266f07.2/ptprfb/sdc3/si:ch73-22o12.1/ptprfa/pvrl2l/nfasca/nrxn3a/mpz/nrcama/slitrk2/cldnf/cldn3d/lrrc4ba/ptprc/lrrc4cb/nrxn1a/slitrk4/cdh2/negr1/itgb2/nlgn4xa/nrxn3b/nectin3b/lrrc4.1/cadm1b/cntnap2a/cntnap2b/esama/cldn30/si:dkey-1h24.6/lrrc4ca/cntn1b/nlgn3a/LOC559501/ncam1b/nlgn2a/ptprd/nlgn3b/ntng1a/itga6l |
| dre04519 | Cadherin signaling | -1.6653062034737 | 0.000319865137159646 | krt92/ppp1cab/dspa/mapk14b/ctnnb1/wnt4b/vav3b/map2k1/actb1/vangl2/arpc5la/fat4/krt98/ywhabl/pcdh20/krt15/pcdh9/lima1a/fat3a/krt91/vim/cdc42/pcdh1g29/crebbpa/igf1rb/lhfpl2a/tmc8/ankrd6b/daam1a/krt99/chuk/lats1/ppp1cb/wnt6b/fat1b/gsk3ba/ep300b/ctnnd1/pkp1b/actb2/scn1ba/ywhah/nckap1/celsr1b/tcf7l2/cdh10a/fat1a/si:dkey-92j12.5/mapk10/krt17/tjp1b/wnt7bb/myo1ca/pcdh17/ikbkg/hrasb/rac3a/gabra1/pcdh11/pkp2/lhfpl5a/taok3b/ctnnd2b/pcdh7b/abi2b/dchs1a/cyfip2/gabrb4/gabra4/gabrb3/scn2b/ctnna2/cdh2/apc/cdh8/pcdh10b/gabrb2a/wnt16/dsc2l/ctnnd2a/wnt3/pcdh7a/fzd8a/lef1/fzd8b/fzd3a/prickle2b/krt93/pcdh10a/psen1/pcdh19/cdh13/pcdh8/gabrg2/si:ch73-233f7.1/mapk12b/pcdh1g30/vcla/dchs1b/ush1ga/scn5lab/apc2/krt95/viml/prickle2a/LOC101886358/gabrr2a/gabrb1/rac1l/pcdh1gc5/ngs/LOC100006777/cdh7a/si:ch211-199f5.1 |
| dre04517 | IgSF CAM signaling | -1.65198049494317 | 0.000319865137159646 | unc5b/map2k1/prkcbb/actb1/rapgef2a/arpc5la/dynll2b/nrp2b/tubb6/ptprz1b/fyna/tubb2/grb2b/kirrel3a/rims1a/plxna4/cdc42/itga5/grb2a/kcna6a/ptprfb/ank3a/unc5db/rims1b/robo4/sptan1/si:ch73-22o12.1/tuba2/ptprfa/gsk3ba/pvrl2l/ank2b/si:ch73-264p11.1/nfasca/tuba1b/mpz/actb2/nrcama/akt3a/slitrk2/si:dkey-92j12.5/kcna1a/plcg1/tubb5/mapk10/tjp1b/pak1/rapgef2b/lrrc4ba/trioa/nrp2a/grip2a/caskin1/zgc:158689/lrrc4cb/zgc:65894/slitrk4/slit2/baiap2a/mpp3a/lrfn5b/appa/robo1/pik3ca/caskb/pik3r3b/cdh2/tuba1a/appb/slit3/slit1b/srgap3/si:dkey-178k16.1/itgb2/ntrk3a/nlgn4xa/ntn1b/nectin3b/cadm1b/cntnap2a/ppfia4/ppfia3/kirrel3l/kirrel1b/cadm4/robo3/lcp2a/cntnap2b/esama/si:dkeyp-9d4.3/si:dkey-1h24.6/tuba4l/il1rapl1a/lrrc4ca/magi1b/tuba1c/srgap1b/mapk12b/cntn1b/nlgn3a/vcla/ncam1b/nlgn2a/ptprd/grip2b/nlgn3b/epb41l3a/ntrk3b/plxna1a/tubb4bl/rac1l/triob/LOC100537363/pak6b |
| dre04916 | Melanogenesis | -1.58602647482429 | 0.0381596612181351 | map2k2a/creb3l3l/fzd2/tyrp1a/tcf7l1b/calm1a/adcy2a/zgc:55558/plcb3/adcy5/ctnnb1/wnt4b/map2k1/prkcbb/adcy7/calm1b/kitlga/crebbpa/si:dkey-206f10.1/si:ch211-132f19.7/creb3l3a/adcy6a/camk2g1/wnt6b/gsk3ba/ep300b/prkacab/camk2b1/adcy1a/tcf7l2/wnt7bb/hrasb/camk2a/camk2g2/LOC101886206/gnai2b/wnt16/wnt3/fzd8a/adcy2b/mitfa/lef1/fzd8b/fzd3a/adcy8/camk2d2/gnao1b/gnao1a/pomca |
| dre04081 | Hormone signaling | -1.44522264541335 | 0.0585001489998652 | smad1/hamp/cga/igf1rb/penkb/si:dkey-206f10.1/si:ch211-132f19.7/adcy6a/bmpr1ba/irs2b/insb/prkacab/sstr5/jak2a/gnb1b/adcy1a/gng13b/gnb1a/crhr1/thrb/LOC101886206/sst1.1/sstr1a/gnai2b/adrb2b/crhb/pik3ca/pik3r3b/penka/pdyn/adra2da/adcy2b/nppal/gnb3a/cckbra/drd4a/adcy8/gnb2/gnao1b/gnao1a/pomca/vipr1b/bmpr1ab/dio2/sstr1b/pnocb/oprd1b/uts1/oprl1/prlhr2a/gper1/trh/oxt/mchr1b/tshba/lepa |
| dre04020 | Calcium signaling pathway | -1.41820224968641 | 0.0294459304579805 | atp2a3/plcd4b/erbb4b/kdrl/camk1gb/plcd3a/pdgfd/orai1b/si:dkey-206f10.1/si:ch211-132f19.7/camk2g1/ret/camk1db/p2rx3b/prkacab/camk1b/camk2b1/adcy1a/si:dkey-28b4.8/plcg1/vegfc/ppp3cca/atp2b2/camk2a/ppp3r1b/cacna1da/camk2g2/LOC101886891/LOC101886206/adrb2b/grin1a/atp2b3b/cacna1e/chrna11/grin2aa/grin2da/gnal/ntrk3a/adcy2b/grin3bb/tpcn1/camk1ga/mcoln3a/mcoln1a/plcd4a/cckbra/grm5a/adcy8/LOC100149482/camk2d2/htr7c/grin2ab/ppp3ca/slc8a4b/tnnc2.1/grin1b/si:ch73-60h1.1/si:ch73-383l1.1/LOC101882451/hgfa/si:dkey-247m21.3/cacna1ab/ntrk3b/fgf17/nfatc4/ppp3cb/grin2bb/ltb4r2a/slc8a2b/plcd3b/grm1a |

**Table S5.** Enriched Kyoto Encyclopedia of Genes and Genomes (KEGG) pathways and their associated genes following M2 exposure, identified by gene set enrichment analysis (GSEA).

| ID | Description | NES | FDR (p adjusted value) | Genes implicated |
| --- | --- | --- | --- | --- |
| GO:0042254 | ribosome biogenesis | 2.68255980674827 | 1.14653846153846e-08 | imp3/rpl26/mrpl10/mrm3a/nle1/noc2l/rbis/urb2/exosc2/polr1c/rpp40/exosc6/abt1/mphosph6/snu13b/emg1/wdr18/znhit3/pwp2h/utp4/utp23/mrm3b/nop14/heatr1/mrto4/pin4/sbds/lsm6/ddx31/wdr12/nom1/nifk/utp11/tfb1m/ddx27/mrpl22/ngdn/pih1d1/wdr3/ddx51/heatr3/gnl3/mrpl20/rrp36/gar1/nvl/riok2/slx9/bxdc2/pdcd11/trmt112/nop58/dimt1l/bud23/rexo4/nip7/mettl16/mphosph10/wdr36/dkc1/npm1a/nsun5/rpusd2/ppan/mdn1/ddx49/rpl7l1/ddx52/bop1/abce1/nol6/sdad1/tbl3/nop10/ddx10/mrpl36/pak1ip1/rpl35/rbfa/zcchc4/rrp9/rps7/si:dkey-251i10.3/utp15/snu13a/rrp7a/nop56/nhp2/wdr75/ddx18/c1d/rcl1/wdr55/rrp8/tsr2/eif6/nat10/utp25/nop2/nop16/polr1d/mterf3/ebna1bp2/rps21/ftsj3/bysl/utp6/rrs1/gnl3l/rps8a/mpv17l2/rps14/nol10/mtrex/rpl5a/riok1/esf1/rpl35a/pop5/ran/rpf1/mrps11/gnl2/rpl14/rpf2/tfb2m/fbl/utp3/dhx37/rrp1/rps16/ddx54/gtpbp10/rpl7/tsr1/ddx28/znf622/malsu1/rps9/mak16/c1qbp/lsg1/naf1/kri1/rpl23a/urb1/exosc8/fcf1/pes/pop4/rps15/rpsa/mtg2/rrp15/rpl7a/mterf4/exosc7/dcaf13/rps5/rplp0/exosc9/tsr3 |
| GO:0006364 | rRNA processing | 2.64217411952833 | 1.14653846153846e-08 | imp3/mrm3a/exosc2/rpp40/exosc6/abt1/mphosph6/snu13b/emg1/wdr18/znhit3/pwp2h/utp4/utp23/mrm3b/heatr1/mrto4/pin4/lsm6/wdr12/nifk/utp11/tfb1m/ddx27/ngdn/pih1d1/wdr3/ddx51/gnl3/rrp36/gar1/riok2/slx9/pdcd11/trmt112/dimt1l/bud23/rexo4/mettl16/mphosph10/wdr36/dkc1/nsun5/rpusd2/ddx49/rpl7l1/ddx52/bop1/nol6/tbl3/nop10/ddx10/rpl35/rbfa/zcchc4/rrp9/rps7/si:dkey-251i10.3/utp15/snu13a/rrp7a/nhp2/wdr75/ddx18/c1d/rcl1/wdr55/rrp8/tsr2/eif6/nat10/utp25/nop2/ebna1bp2/rps21/ftsj3/bysl/utp6/rrs1/gnl3l/rps8a/rps14/nol10/mtrex/riok1/esf1/pop5/rpf1/mrps11/gnl2/rpf2/tfb2m/fbl/utp3/dhx37/rrp1/rps16/ddx54/rpl7/tsr1/mak16/naf1/kri1/urb1/exosc8/fcf1/pes/pop4 |
| GO:0034470 | ncRNA processing | 2.61312605149024 | 1.14653846153846e-08 | ints6/prkra/imp3/thumpd2/pus3/wdr4/qtrt2/trmu/mrm3a/mtfmt/trmt10c/exosc2/rpp21/rnf113a/rpp40/exosc6/abt1/mphosph6/snu13b/rpp30/emg1/henmt1/wdr18/tprkb/piwil2/znhit3/elp3/zgc:136564/pwp2h/elp5/utp4/trmt61a/spout1/zc3h7bb/tp53rk/utp23/mrm3b/trmt44/heatr1/mrto4/pin4/trmt6/exosc5/tyw3/rpusd4/adat1/tsen2/trmt12/lsm6/wdr12/nsun2/nifk/utp11/thumpd3/tfb1m/osgep/ddx27/urm1/ctu2/mettl3/ngdn/pih1d1/wdr3/srrt/ddx51/gnl3/rrp36/gar1/riok2/ints6l/slx9/pdcd11/trmt112/dimt1l/bud23/osgepl1/rexo4/ptcd1/mettl16/mphosph10/wdr36/dkc1/mettl2a/thada/pus1/nsun5/gtpbp3/rpusd2/ddx49/rpl7l1/ddx52/bop1/trmt10b/tdrd9/nol6/clp1/ctu1/tsen15/rtcb/tbl3/ints9/nop10/ddx10/rpl35/rbfa/zcchc4/rrp9/rpph1/rps7/si:dkey-251i10.3/utp15/snu13a/rrp7a/elp6/nhp2/wdr75/ddx18/c1d/rcl1/wdr55/rrp8/tsr2/trmt61b/elac1/trpt1/eif6/nat10/utp25/nop2/ebna1bp2/tut1/rps21/elp2/ftsj3/bysl/mettl1/pus7/utp6/rrs1/dtwd1/gnl3l/rps8a/rps14/nol10/rpp25l/thg1l/mtrex/riok1/sepsecs/esf1/eif1axb/pop5/cdk5rap1/zbtb8os/rpf1/mrps11/gnl2/rpf2/tfb2m/trmt2b/fbl/utp3/rpp38/dhx37/rtraf/rrp1/wdr6/ints11/rps16/ddx54/aars1/rpl7/tsr1/mak16/trmt5/trmt9b/naf1/kri1/urb1/snip1/exosc8/fcf1/pes/pop4 |
| GO:0016072 | rRNA metabolic process | 2.5944143069453 | 1.14653846153846e-08 | imp3/mrm3a/exosc2/rpp40/exosc6/abt1/mphosph6/snu13b/emg1/wdr18/znhit3/pwp2h/utp4/utp23/mrm3b/heatr1/mrto4/pin4/exosc5/lsm6/wdr12/nifk/utp11/tfb1m/ddx27/ngdn/pih1d1/wdr3/ddx51/gnl3/rrp36/gar1/riok2/slx9/pdcd11/trmt112/dimt1l/bud23/rexo4/mettl16/mphosph10/wdr36/dkc1/polr1e/nsun5/rpusd2/ddx49/rpl7l1/ddx52/bop1/nol6/tbl3/nop10/ddx10/rpl35/rbfa/zcchc4/rrp9/rps7/si:dkey-251i10.3/utp15/snu13a/rrp7a/nhp2/wdr75/ddx18/c1d/rcl1/wdr55/rrp8/tsr2/eif6/nat10/utp25/nop2/taf1b/ebna1bp2/rps21/ftsj3/bysl/dis3/utp6/rrs1/gnl3l/rps8a/rps14/nol10/mtrex/riok1/esf1/pop5/rpf1/mrps11/gnl2/rpf2/tfb2m/fbl/utp3/polr1a/dhx37/rrp1/rps16/ddx54/pelo/rpl7/tsr1/mak16/naf1/kri1/urb1/exosc8/fcf1/pes/pop4 |
| GO:0034660 | ncRNA metabolic process | 2.54552940229038 | 1.14653846153846e-08 | ints6/prkra/fars2/imp3/thumpd2/pus3/farsa/wdr4/qtrt2/trmu/mrm3a/dars2/wars2/mtfmt/trmt10c/exosc2/rpp21/rnf113a/rpp40/exosc6/dars1/abt1/mphosph6/snu13b/rpp30/emg1/henmt1/wdr18/tprkb/piwil2/znhit3/elp3/zgc:136564/pwp2h/elp5/utp4/trmt61a/spout1/zc3h7bb/tp53rk/utp23/mrm3b/trmt44/heatr1/mrto4/pin4/trmt6/exosc5/tyw3/lars1b/rpusd4/adat1/tsen2/trmt12/lsm6/wdr12/nsun2/nifk/utp11/thumpd3/si:ch211-256m1.8/tfb1m/mars2/osgep/ddx27/urm1/ctu2/mettl3/ngdn/pih1d1/wdr3/srrt/ddx51/snapc3/gnl3/rrp36/gar1/riok2/ints6l/slx9/pdcd11/trmt112/qrsl1/dimt1l/bud23/osgepl1/rexo4/ptcd1/mettl16/gatb/mphosph10/wdr36/dkc1/mettl2a/thada/polr1e/pus1/nsun5/gtpbp3/rpusd2/iars1/ddx49/hars/rpl7l1/ddx52/bop1/trmt10b/tdrd9/qars1/nol6/nars2/vars1/clp1/ctu1/tsen15/rtcb/tbl3/ints9/nop10/ddx10/rpl35/rbfa/zcchc4/rrp9/rpph1/rps7/si:dkey-251i10.3/utp15/snu13a/rrp7a/elp6/nhp2/wdr75/ddx18/yars1/c1d/rcl1/wdr55/rrp8/tsr2/trmt61b/elac1/rpap2/trpt1/rars1/eif6/nat10/utp25/nop2/taf1b/ebna1bp2/tut1/rps21/elp2/tars1/ftsj3/bysl/gars1/mettl1/pars2/dis3/pus7/iars2/utp6/rrs1/dtwd1/gnl3l/rps8a/rps14/nol10/rpp25l/thg1l/mtrex/riok1/sars1/sepsecs/esf1/eif1axb/pop5/vars2/mepceb/cdk5rap1/zbtb8os/rpf1/mrps11/gnl2/sars2/rpf2/tfb2m/trmt2b/fbl/utp3/polr1a/rpp38/dhx37/rtraf/rrp1/wdr6/ints11/rps16/ddx54/aars1/pelo/rpl7/tsr1/mak16/trmt5/trmt9b/naf1/kri1/dtd2/urb1/snip1/exosc8/fcf1/pes/pop4 |
| GO:0022613 | ribonucleoprotein complex biogenesis | 2.40988428678125 | 1.14653846153846e-08 | imp3/rpl26/mrpl10/mrm3a/nle1/noc2l/rbis/gemin5/urb2/exosc2/polr1c/rpp40/exosc6/abt1/mphosph6/snu13b/emg1/wdr18/znhit3/pwp2h/trnau1apa/utp4/utp23/mrm3b/nop14/heatr1/mrto4/pin4/sbds/snrpd2/lsm6/ddx31/wdr12/nom1/nifk/utp11/tfb1m/prmt7/ddx27/mrpl22/ngdn/pih1d1/gemin2/wdr3/ddx51/heatr3/gnl3/mrpl20/rrp36/gar1/nvl/snrpg/riok2/slx9/bxdc2/pdcd11/trmt112/nop58/dimt1l/bud23/prpf18/mcts1/rexo4/nip7/mettl16/mphosph10/wdr36/dkc1/npm1a/nsun5/rpusd2/ppan/mdn1/ddx49/rpl7l1/ddx52/bop1/abce1/nol6/clp1/sdad1/tbl3/nop10/ddx10/mrpl36/pak1ip1/rpl35/rbfa/zcchc4/rrp9/ptges3b/rps7/si:dkey-251i10.3/utp15/snu13a/rrp7a/nop56/eif3m/nhp2/wdr75/ddx18/c1d/nopchap1/snrpd3l/eif3ea/rcl1/wdr55/rrp8/tsr2/coil/eif6/snrpf/nat10/utp25/nop2/nop16/usp39/polr1d/mterf3/ebna1bp2/rps21/eif3jb/ftsj3/snrpe/bysl/utp6/rrs1/gnl3l/rps8a/mpv17l2/rpl38/rps14/nol10/shq1/mtrex/rpl5a/riok1/esf1/rpl35a/pop5/eif5/ran/eif3ha/rpf1/trnau1apb/mrps11/gnl2/scaf11/rpl14/rpf2/tfb2m/fbl/utp3/dhx37/rrp1/eif3s10/cdc73/rps16/ddx54/denr/gtpbp10/rpl7/tsr1/ddx28/znf622/malsu1/rps9/mak16/c1qbp/lsg1/naf1/kri1/rpl23a/urb1/exosc8/eif3s6ip/fcf1/pes/pop4/rps15/rpsa/eif2s2/mtg2/polr2d/rrp15/rpl7a/srpk1a/mterf4/exosc7/dcaf13/rps5/gemin8/rplp0/exosc9/tsr3/snrpd1/rps27.1/aar2/rpl10/rpl5b/eif3g/gtpbp4/eif3d/rpl6/yju2/noc4l |
| GO:0140053 | mitochondrial gene expression | 2.16306020123481 | 0.000153043817326382 | fastkd3/tefm/trmu/fastkd2/tbrg4/wars2/trmt10c/mrpl58/fastkd1/coa3a/mtg1/mrps18b/tfb1m/uqcc2/chchd1/tsfm/mtrf1l/qrsl1/gatb/gtpbp3/mtif2/hars/mrps24/rcc1l/tfam/trmt61b/mrpl47/mterf3/gars1/iars2/mrpl23/sars2/tfb2m/ptcd3/fastk/mrpl16 |
| GO:0043043 | peptide biosynthetic process | 2.02998011519709 | 1.14653846153846e-08 | dazl/fars2/farsa/zc3h12ab/dph7/rpl26/mrpl10/dars2/wars2/mtfmt/trmt10c/gemin5/exosc2/mrpl58/mrps5/dars1/rpp30/zar1/mrpl9/dcp1b/piwil2/coa3a/trnau1apa/mrpl35/mif4gdb/mrpl30/eif2ak1/dcps/mrpl17/dph5/lars1b/mtg1/mrps17/mrps18b/mrpl4/si:ch211-256m1.8/zgc:171480/mars2/uqcc2/chchd1/tsfm/gss/eif4ea/mrpl22/mettl3/ngdn/mtrf1l/faub/mrps18c/drg2/lsm1/mrpl27/mrpl20/dph6/magoh/eif1ad/eif2a/eefsec/qrsl1/mrpl51/mrps23/mcts1/ago3b/mettl16/gatb/mrpl18/mrps21/gtpbp3/mtif2/iars1/mrpl14/rpl12/hars/mrpl24/pdf/eif2b1/eif4e2rs1/abce1/eif2s1a/qars1/nars2/mrpl41/impact/vars1/mrpl34/fxr2/larp6a/mrps24/mrpl36/rpl35/cnot8/mrps12/mrpl13/rcc1l/eif5a2/rpl32/mrps2/rps7/mrpl12/brf1a/eif2b2/eif3m/eif4ebp3l/yars1/bdh2/eef1b2/eif3ea/mrpl11/rpl22l1/mrpl33/rpl34/rars1/eif6/dph2/ogfod1/gspt1/pde12/mrpl47/rps21/rps17/eef1a1l3/eif3jb/tars1/cnot6l/eef1da/eif4ebp1/gars1/mrps10/pars2/iars2/rc3h2/mrpl23/rpl29/zgc:171772/rpl37/rps8a/gatc/rpl38/rps3a/rps14/rps15a/srp9/larp4b/mrps30/rps20/rps29/tcea3/rpl5a/rpl15/sars1/sepsecs/mrpl3/eif1axb/rpl35a/vars2/eif5/rps18/eif3ha/zc3h15/mrpl32/mrpl37/trnau1apb/mrps11/rpl11/rpl14/rpl39/sars2/uba52/rpl36a/rps26/eif1b/ssb/igf2bp3/rpl24/eif4eb/ptcd3/rpl27/rpl23/eif3s10/rpl31/rps24/rps16/rwdd1/denr/aars1/rpl18/rpl13a/rpl22/mrps16/rbm24a/rplp2/mrpl16/mettl17/mtrfr/drg1/malsu1/rps9/rbm8a/rpl28/rack1/rplp1/rpl23a/eif3f/gspt1l/mrpl2/exosc8/larp1b/eif3s6ip/rps15/rpsa/eef1g/eif2s2/cnbpa/ftsj1/rps11/eif4e3/rplp2l/zgc:114188/polr2d/rpl17/cnot11/faua/pabpc1a/exosc7/rpl19/rps5/eif2s1b/rplp0/rpl18a/exosc9/tufm/rpl21/kars1/rpl4/eef2b/jmjd4/cnbpb/rps13/rpl8/eif5b/rps27.1/rpl10/rpl5b/eif3g/eif3d/mrps35/rpl6/gfm1/eif4e1c/rps3/eif1axa/cnot7/eif2b4/rps12/eef1a1l1/znf598/rpl9/cnot10/eef2a.1/rpl10a/cars1/rps6/rps27a/mars1/nck2a/rps27.2/rps28/mrps7/eif4bb/rpl36/rps23/eif2s3/gtpbp1l/nck2b/eif4g1a/cnot9/dcp2/rps19/dph1/ybx1/dhx29/eif3ja/mrpl15/eif4g3b/samd4a |
| GO:0006412 | translation | 2.02950881785419 | 1.14653846153846e-08 | dazl/fars2/farsa/zc3h12ab/dph7/rpl26/mrpl10/dars2/wars2/mtfmt/trmt10c/gemin5/exosc2/mrpl58/mrps5/dars1/rpp30/zar1/mrpl9/dcp1b/piwil2/coa3a/trnau1apa/mrpl35/mif4gdb/mrpl30/eif2ak1/dcps/mrpl17/dph5/lars1b/mtg1/mrps17/mrps18b/mrpl4/si:ch211-256m1.8/zgc:171480/mars2/uqcc2/chchd1/tsfm/eif4ea/mrpl22/mettl3/ngdn/mtrf1l/faub/mrps18c/drg2/lsm1/mrpl27/mrpl20/dph6/magoh/eif1ad/eif2a/eefsec/qrsl1/mrpl51/mrps23/mcts1/ago3b/mettl16/gatb/mrpl18/mrps21/gtpbp3/mtif2/iars1/mrpl14/rpl12/hars/mrpl24/pdf/eif2b1/eif4e2rs1/abce1/eif2s1a/qars1/nars2/mrpl41/impact/vars1/mrpl34/fxr2/larp6a/mrps24/mrpl36/rpl35/cnot8/mrps12/mrpl13/rcc1l/eif5a2/rpl32/mrps2/rps7/mrpl12/brf1a/eif2b2/eif3m/eif4ebp3l/yars1/eef1b2/eif3ea/mrpl11/rpl22l1/mrpl33/rpl34/rars1/eif6/dph2/ogfod1/gspt1/pde12/mrpl47/rps21/rps17/eef1a1l3/eif3jb/tars1/cnot6l/eef1da/eif4ebp1/gars1/mrps10/pars2/iars2/rc3h2/mrpl23/rpl29/zgc:171772/rpl37/rps8a/gatc/rpl38/rps3a/rps14/rps15a/srp9/larp4b/mrps30/rps20/rps29/tcea3/rpl5a/rpl15/sars1/sepsecs/mrpl3/eif1axb/rpl35a/vars2/eif5/rps18/eif3ha/zc3h15/mrpl32/mrpl37/trnau1apb/mrps11/rpl11/rpl14/rpl39/sars2/uba52/rpl36a/rps26/eif1b/ssb/igf2bp3/rpl24/eif4eb/ptcd3/rpl27/rpl23/eif3s10/rpl31/rps24/rps16/rwdd1/denr/aars1/rpl18/rpl13a/rpl22/mrps16/rbm24a/rplp2/mrpl16/mettl17/mtrfr/drg1/malsu1/rps9/rbm8a/rpl28/rack1/rplp1/rpl23a/eif3f/gspt1l/mrpl2/exosc8/larp1b/eif3s6ip/rps15/rpsa/eef1g/eif2s2/cnbpa/ftsj1/rps11/eif4e3/rplp2l/zgc:114188/polr2d/rpl17/cnot11/faua/pabpc1a/exosc7/rpl19/rps5/eif2s1b/rplp0/rpl18a/exosc9/tufm/rpl21/kars1/rpl4/eef2b/jmjd4/cnbpb/rps13/rpl8/eif5b/rps27.1/rpl10/rpl5b/eif3g/eif3d/mrps35/rpl6/gfm1/eif4e1c/rps3/eif1axa/cnot7/eif2b4/rps12/eef1a1l1/znf598/rpl9/cnot10/eef2a.1/rpl10a/cars1/rps6/rps27a/mars1/nck2a/rps27.2/rps28/mrps7/eif4bb/rpl36/rps23/eif2s3/gtpbp1l/nck2b/eif4g1a/cnot9/dcp2/rps19/dph1/ybx1/dhx29/eif3ja/mrpl15/eif4g3b/samd4a |
| GO:0045333 | cellular respiration | 1.99293254414966 | 4.41989406426173e-05 | ccn6/tefm/sfxn4/uqcc3/si:dkey-85n7.8/ugl/sdhda/ndufs8b/atpsckmt/mdh1b/mdh1ab/dhtkd1/wfs1b/uqcc2/idh3b/sdhaf2/atp5mf/atp5po/sdhdb/ndufb6/lyrm7/cox7b/uqcr10/ndufb8/mrpl24/etfdh/ndufa10/cox6c/cox4i1/ndufv1/ndufaf1/suclg2/si:dkey-190g11.3/antkmt/cox7c/ndufa7/atp5f1e/ndufa8/ndufc2/cox7a2a/sdhc/ogdhb/slc25a55b/slc25a12/coa6/sdha/ndufs6/pm20d1.1/cox6a1/uqcrq/lyrm5a/ndufs2/ndufb9/noa1/sucla2/idh1/uqcrfs1/ndufs8a/cox6a2/cox5ab/lyrm5b/uqcrh/ndufa5/ndufv2/idh3a/mdh2/idh2/cox7a2l/sdhb/dlst/uqcrb/ndufs7/cox5aa/cyc1/ND3/fh/cox5b2/ndufv3/ogdha/cox8a |
| GO:0006518 | peptide metabolic process | 1.98118588803671 | 1.14653846153846e-08 | dazl/fars2/farsa/zc3h12ab/dph7/parla/rpl26/mrpl10/dars2/wars2/mtfmt/trmt10c/gemin5/exosc2/mrpl58/mrps5/dars1/rpp30/zar1/erap2/mrpl9/dcp1b/piwil2/ggt1b/coa3a/trnau1apa/mrpl35/mif4gdb/mrpl30/eif2ak1/dcps/mrpl17/dph5/lars1b/mtg1/mrps17/mrps18b/cpz/mrpl4/si:ch211-256m1.8/zgc:171480/mars2/uqcc2/chchd1/tsfm/gss/eif4ea/mrpl22/mettl3/ngdn/mtrf1l/faub/mrps18c/drg2/lsm1/mrpl27/mrpl20/dph6/magoh/eif1ad/eif2a/eefsec/qrsl1/mrpl51/mrps23/mcts1/ago3b/mettl16/gatb/mrpl18/immp2l/mrps21/gtpbp3/mtif2/iars1/mrpl14/rpl12/hars/mrpl24/pdf/enpep/eif2b1/eif4e2rs1/abce1/eif2s1a/qars1/nars2/mrpl41/pcsk5b/impact/vars1/mrpl34/sec11a/nln/fxr2/larp6a/mrps24/mrpl36/rpl35/cnot8/mrps12/mrpl13/rcc1l/eif5a2/rpl32/mrps2/rps7/mrpl12/brf1a/eif2b2/eif3m/eif4ebp3l/yars1/bdh2/eef1b2/eif3ea/mrpl11/rpl22l1/mrpl33/rpl34/rars1/eif6/dph2/ogfod1/gspt1/pde12/sppl3/mrpl47/rps21/rps17/eef1a1l3/eif3jb/tars1/cnot6l/eef1da/eif4ebp1/gars1/mrps10/pars2/iars2/rc3h2/mrpl23/rpl29/zgc:171772/rpl37/rps8a/gatc/rpl38/spcs3/rps3a/rps14/rps15a/srp9/larp4b/mrps30/rps20/rps29/tcea3/rpl5a/rpl15/sars1/sepsecs/mrpl3/eif1axb/rpl35a/vars2/eif5/rps18/eif3ha/zc3h15/mrpl32/mrpl37/trnau1apb/mrps11/rpl11/rpl14/rpl39/sars2/uba52/rpl36a/rps26/eif1b/thop1/ssb/igf2bp3/rpl24/eif4eb/ptcd3/rpl27/rpl23/eif3s10/rpl31/rps24/rps16/rwdd1/denr/aars1/rpl18/rpl13a/rpl22/mrps16/mipepb/rbm24a/spcs1/rplp2/mrpl16/mettl17/mtrfr/drg1/malsu1/rps9/rbm8a/rpl28/rack1/rplp1/rpl23a/eif3f/gspt1l/mrpl2/exosc8/larp1b/eif3s6ip/rps15/rpsa/eef1g/eif2s2/cnbpa/ftsj1/rps11/eif4e3/rplp2l/zgc:114188/polr2d/rpl17/cnot11/faua/pabpc1a/exosc7/rpl19/rps5/eif2s1b/rplp0/rpl18a/exosc9/tufm/rpl21/kars1/rpl4/eef2b/jmjd4/cnbpb/rps13/rpl8/eif5b/rps27.1/rpl10/rpl5b/eif3g/eif3d/mrps35/rpl6/gfm1/ide/eif4e1c/rps3/eif1axa/cnot7/eif2b4/rps12/eef1a1l1/znf598/rpl9/cnot10/eef2a.1/rpl10a/spcs2/cars1/rps6/rps27a/mars1/nck2a/rps27.2/rps28/mrps7/eif4bb/rpl36/rps23/eif2s3/gtpbp1l/nck2b/lnpep/eif4g1a/cnot9/dcp2/rps19/dph1/ybx1/dhx29/eif3ja/mrpl15/eif4g3b/samd4a |
| GO:0007005 | mitochondrion organization | 1.97295174722294 | 2.83369625164005e-07 | si:dkey-32e23.4/bnip4/adck1/sqstm1/mtm1/mtx1b/mtx1a/rrm2b/sco2/si:dkey-82o10.4/vps13d/sdhaf1/uqcc3/sdhaf3/chchd4b/atp23/phb2b/timm21/ppargc1a/timm29/maip1/coa3a/sdhaf4/aifm1/ndufs8b/chchd4a/alkbh7/chchd6b/cox17/chchd7/coa4/mgme1/cox20/slc25a42/bcs1l/timm13/uqcc2/surf1/sdhaf2/timm50/spata18/lonp1/tmem126a/dnajc19/grpel1/bida/chchd3b/atp13a2/immp2l/wdr45b/lyrm7/uqcc6/cox19/etfa/cluha/dna2/coa5/snx7/apooa/hspd1/atpaf2/phb/ndufaf1/atg5/snx30/polg2/uqcc1/higd1a/chchd10/cox7a2a/primpol/atpaf1/higd2a/si:ch211-166a6.5/pet100/tomm40/apool/fam162a/agtpbp1/pmaip1/romo1/ndufa11/oxa1l/twnk/samm50/rpl37/ndufs8a/rhot1a/slc25a4/nubpl/ndufaf8/ttc19/dnajc11a/mfn1b/miga1/tmem70/dnlz/dnaja3a/mipepb/foxred1/cox7a2l/dlst/atg7/tafazzin/tomm40l/fis1 |
| GO:0002088 | lens development in camera-type eye | 1.96162227904054 | 0.000326438123176076 | fgf19/crygm2a/crybb2/crybb1l3/crygm5/crygm2d6/crygm2b/mipb/cryba1l2/crygm2d9/crygm2d15/dnajb1b/crygm2d12/crygm2d18/cryba2a/crygm2d17/crygm2d7/crygm2d1/lgsn/crygm2d5/crygm2f/cryba1l1/crybb1l2/dlx3b/crygm2d3/crygm2d20/crygm2d11/crygm2d10/crygm2d2/crygm2d21/cryba1a/dlx4b/mipa/lama1/crygm2d13/crygmxl2/crygn2/crygm2d16/cryba4/crygm2d8/tdrd7b/crygm2d14/cryba1b/crygn1/psmc3/foxe3/cryba2b/hsf4/unc45b/crygmx/crygmxl1/si:dkey-57a22.15/sox11b/crybb1l1/psmd6 |
| GO:0009145 | purine nucleoside triphosphate biosynthetic process | 1.93013788013767 | 0.00988355843878473 | nme9/uqcc3/atp5mc3a/atpsckmt/atp5mf/atp5pd/atp5po/atp5mea/atp5mc1/nme2b.2/nme6/atp5f1c/mrpl24/atp5meb/nme2a/si:dkey-190g11.3/antkmt/atp5f1e/nme3/atp5pf/atp5pb/atp5mc3b/nme7/atp5fa1/dlst/ATP6 |
| GO:0009206 | purine ribonucleoside triphosphate biosynthetic process | 1.93013788013767 | 0.00988355843878473 | nme9/uqcc3/atp5mc3a/atpsckmt/atp5mf/atp5pd/atp5po/atp5mea/atp5mc1/nme2b.2/nme6/atp5f1c/mrpl24/atp5meb/nme2a/si:dkey-190g11.3/antkmt/atp5f1e/nme3/atp5pf/atp5pb/atp5mc3b/nme7/atp5fa1/dlst/ATP6 |
| GO:0008202 | steroid metabolic process | 1.89342793339491 | 0.000798277428982877 | ch25h/sdr42e1/dhcr24/cyp46a1.1/ch25hl1.1/msmo1/amacr/nsdhl/cyp46a1.3/cyp39a1/cyp46a1.2/hmgcra/mvk/ebpl/hsd17b12b/bdh1/srd5a1/cyp51/gba/cyp21a2/cyp7a1/lipeb/slc27a2b/faxdc2/cyp46a1.4/apoea/cyp2r1/mvda/ch25hl2/fgfr4/lss/hmgcs1/ebp/cyp26c1/apoa1a/cyp27a1.4/hsd17b7/cyb5r2/cyp27a1.1/hsd17b14/rdh1/srd5a2b/erlin1/slc27a2a |
| GO:0006979 | response to oxidative stress | 1.83959019549846 | 0.00355248534040968 | bco2l/fancc/anxa1b/glrx2/keap1a/per2/prdx1/ptgs2a/sod2/park7/ptgs1/ptgs2b/lonp1/prdx3/si:ch211-120k19.1/trpm2/gpx4a/diabloa/impact/gpx1a/gpx9/hmox2a/gpx4b/gpx8/naprt/mpv17/brf2/sirt2/romo1/sod1/txndc12/ndufa6/mpv17l2/hmox1a/ryr1b/msraa |
| GO:0006260 | DNA replication | 1.7556677318185 | 0.00194590276805455 | dbf4/zranb3/polq/zgc:173742/wrn/lig4/pold3/rpa3/mcm8/wdr18/tonsl/terf1/rmi1/poln/ctc1/mgme1/rnaseh1/ssrp1a/gins3/rpa2/stn1/orc4/dna2/noc3l/mcm7/rfc5/gins2/orc5/tatdn1/pole2/polg2/rrm1/tipin/fen1/primpol/mcm3/rrm2/traip/orc1/lig1/igfbp2a/mcmbp/mcm2/wrnip1/twnk/recql/rad51/rfc2/mcm5/rpa1/esco2/cenps/mms22l/eme1/orc6/rbbp4/gins4/blm/rnaseh2a/cdc45/prim1/rfc3/helb/pcna/rbb4l/lrwd1/samhd1/orc3/dscc1/ssbp1/top1a/mcm4/pole |
| GO:0044743 | protein transmembrane import into intracellular organelle | 1.73136835035816 | 0.0470692918806812 | pex10/chchd4b/timm21/chchd4a/timm50/dnajc19/grpel1/hspd1/tomm40/romo1/pex1/dnlz/pex13/tomm40l/pex5/timm23a/tomm70a/tomm7/pex7/pam16/tomm20b/pex26 |
| GO:0050731 | positive regulation of peptidyl-tyrosine phosphorylation | 1.72535656967289 | 0.0678490288954661 | unc119a/epgn/arl2bp/btc/smtlb/hbegfa/zgc:153759 |
| GO:0034599 | cellular response to oxidative stress | 1.71630178467435 | 0.0639818065496334 | bco2l/fancc/anxa1b/glrx2/keap1a/prdx1/sod2/park7/lonp1/si:ch211-120k19.1/diabloa/impact/mpv17/brf2/sirt2/romo1/sod1/mpv17l2/ryr1b/msraa |
| GO:0042744 | hydrogen peroxide catabolic process | 1.71035395380329 | 0.0737625497040733 | prdx1/hbae5/zgc:163057/hbbe2/prdx3/hbae1.3/hbae1.1/hbae3/gpx1a |
| GO:0006974 | cellular response to DNA damage stimulus | 1.70302954476004 | 6.9422407694291e-06 | uba7/wdr76/hrob/mms19/ddias/cry5/slx1b/zswim7/fancc/zranb3/tdp2b/xpc/parpbp/rad51d/swsap1/chek2/cry-dash/si:ch211-127m7.2/msh3/palb2/polq/hic1/phkg1b/xndc1/neil1/rfwd3/rrm2b/ogg1/sfr1/wrn/fanci/rad54b/brip1/cpdp/lig4/pargl/mre11a/zgc:112496/rad51b/faap100/pold3/rpa3/atm/cdc7/mcm8/nthl1/mcm9/ddb2/ap5s1/ercc2/kmt5ab/aptx/nbn/tonsl/cep63/eya4/rmi1/alkbh7/poln/mnat1/si:dkey-74k8.3/mgme1/park7/dek/mpg/smyd2a/mettl3/spata18/polh/ssrp1a/tdp1/neil3/rpa2/stn1/poldip2/xrcc3/gtf2h3/trip13/zfyve26/fancg/msh2/dna2/mcm7/smarcad1a/rfc5/uba1/swi5/gins2/fxr2/xrcc6/rad51c/pole2/polg2/tipin/yars1/ercc6/ube2nb/prmt6/fen1/primpol/tigara/mcm3/unga/sem1/bcl2l16/rad23b/pmaip1/lig1/polr2i/bccip/nek4/vhl/mcm2/abraxas1/txndc12/rbbp8/ttf2/mlh1/rmi2/wrnip1/recql/poli/rad1/rad51/rfc2/mcm5/rpa1/ercc1/plrg1/smyd2b/cenps/psme4a/mms22l/eme1/cdca5/si:ch211-141o9.10/hltf |
| GO:0044283 | small molecule biosynthetic process | 1.67134004649573 | 0.000341676076873402 | gls2b/tecrl2b/pycr3/fads2/mcat/msmo1/sdsl/asns/gatm/chst15/elovl8a/gpib/anxa1b/prxl2b/mthfr/acacb/cyp39a1/elovl2/pycr1b/hmgcra/mvk/gch2/qdprb.1/otc/uros/ptgs2a/olah/hsd17b12b/glsl/coq5/aass/akr1a1b/srd5a3/moxd1l/pdss2/asnsd1/srr/cyp51/cyp7a1/adka/park7/lgsn/slc27a2b/apip/rfk/ptgs1/ptgs2b/mvda/pnpo/abhd2a/pcbd1/kmo/ptgesl/pah/elovl6l/fgfr4/tecra/asah2/lss/hmgcs1/ebp/apoa1a/cyp27a1.4/gch1/acmsd/hsd17b7/gnmt/coq4/pck2/elovl8b/sephs1/cdo1/ptges3b/gamt/cyp27a1.1/ndufab1b/fbp1b/impa1/bhmt/upb1/acot7/rgn/fa2h/slc27a2a/aprt/sptlc3/prtfdc1b/agxtb/g6pc3/hprt1/pdss1/ubiad1/coq6/cth/ndufa9a/qdpra |
| GO:0052646 | alditol phosphate metabolic process | 1.62136572553731 | 0.0983308153791125 | gpd1l/rfk/gpam/gpd1c/gpd1a/gpd2 |
| GO:0032259 | methylation | 1.61555246309617 | 0.00489199162953128 | hnmt/thumpd2/suv39h1a/zgc:113054/mettl21cb/wdr4/tpmt.2/etfbkmt/mrm3a/dnmt3bb.2/eef1akmt1/cskmt/trmt10c/dnmt3bb.3/emg1/ash2l/tyms/kmt5ab/henmt1/piwil2/coq5/trmt61a/vcpkmt/mrm3b/trmt44/atpsckmt/mettl4/trmt6/tyw3/trmt2a/dph5/trmt12/nsun2/eef1akmt2/thumpd3/tfb1m/prmt9/smyd2a/tpmt.1/hells/prmt7/mettl3/smyd1a/prmt3/trmt11/trmt112/dimt1l/bud23/ntmt1/mettl16/mettl21a/mettl2a/thada/nsun5/gtpbp3/amt/trmt10b/tdrd9/dnmt3bb.1/gnmt/zcchc4/gamt/cebpb/si:dkey-190g11.3/antkmt/cxxc1b/rrp8/mrm2/setd3/bhmt/prmt6/skic8/trmt61b/comta/lyn/unga/nop2/comtd1 |
| GO:0043436 | oxoacid metabolic process | 1.61491700112608 | 2.31796694375566e-05 | gls2b/tecrl2b/pycr3/fars2/fads2/ddo/mcat/farsa/amacr/sdsl/accs/etfbkmt/asns/si:dkey-238o13.4/gatm/elovl8a/gpib/anxa1b/acot20/prxl2b/dars2/acot17/wars2/mthfr/asah1a/acot11b/ugl/sdhaf3/acacb/cyp39a1/elovl2/hoga1/pycr1b/dars1/nit2/qdprb.1/gatd1/otc/abcd3b/uros/ptgs2a/olah/hsd17b12b/glsl/tdo2a/gcat/enosf1/uevld/aass/amdhd2/alkbh7/asnsd1/srr/mdh1b/mdh1ab/dhtkd1/marc1/lars1b/lpin1b/cyp7a1/tysnd1/park7/si:ch211-256m1.8/acaa1/tat/acads/lgsn/slc27a2b/mars2/apip/ptgs1/idh3b/ptgs2b/pnpo/tph1b/gpam/abhd2a/hacl1/hsd17b4/cpt2/adpgk2/kyat1/decr1/kmo/qrsl1/bpgm/ptgesl/gck/pah/elovl6l/si:ch211-212d10.2/gatb/zgc:162816/fgfr4/tecra/gtpbp3/asah2/iars1/amt/etfa/cpt1b/hars/acss1/gcshb/cmah/cyp26c1/ctps1a/acadm/qars1/nars2/vars1/hadhab/auh/bckdha/acmsd/gnmt/pck2/elovl8b/sephs1/cdo1/mrps36/ptges3b/eno1b/abcd3a/gamt/pgam2/gnpda1/ndufab1b/mccc2/pklr/yars1/aacs/hadh/gcdha/me1/bhmt/hpdb/upb1/acot7/rgn/rars1/tigara/fa2h/slc27a2a/kyat3.2/hibch/mpst/acss2/echdc2/eci1/hgd/agxtb/pm20d1.1/tars1/hadhaa/gars1/pars2/cth/qdpra/ptgr1.1/iars2/acsf2/aspg/gcsha/kynu/hadhb/acsl5/glud1b/got2b/hpdl/cpt1aa/hk2/idh1/acsl1b/sars1/ivd/acat2/vars2/acaa2/ehhadh/acot18/nags/amdhd1/sars2/phyh/ftcd/sgpl1/gip/slc25a44b/shmt1/pfas/acot15/mgst2/aldoab/elovl4b/idh3a/tecrb/aars1/ldha/elovl7a |
| GO:0044282 | small molecule catabolic process | 1.59271540808738 | 0.00441611574194559 | gls2b/cyp46a1.1/ddo/sdsl/etfbkmt/si:dkey-238o13.4/cyp46a1.3/bpnt1/cyp46a1.2/hoga1/qdprb.1/gatd1/abcd3b/glsl/tdo2a/gcat/enosf1/aass/srd5a3/amdhd2/srr/zgc:103586/rbks/miox/lpin1b/cyp7a1/tysnd1/acaa1/tat/acads/cyp46a1.4/apoea/uraha/sord/abhd2a/hacl1/hsd17b4/cpt2/decr1/kmo/pah/zgc:162816/amt/etfa/cpt1b/gcshb/cdaa/acadm/hadhab/auh/bckdha/pck2/cdo1/abcd3a/adh8b/gnpda1/mccc2/impa1/hadh/gcdha/hpdb/kyat3.2/hibch/echdc2/eci1/hgd/hadhaa/qdpra/gcsha/kynu/hadhb/glud1b/got2b/esd/cpt1aa/ivd/acat2/acaa2/ehhadh/amdhd1/phyh/ftcd/slc25a44b/shmt1 |
| GO:1901361 | organic cyclic compound catabolic process | 1.58649464452635 | 0.00292032014134612 | hnmt/nudt4b/cyp46a1.1/dazl/dnase1l1/blvra/fastkd3/fitm2/zc3h12ab/fastkd2/cyp46a1.3/tbrg4/nudt18/exosc2/cyp46a1.2/fastkd1/exosc6/qdprb.1/entpd3/dcp1b/pde4cb/tdo2a/nudt15/moxd1l/zgc:103586/dut/heatr1/mrto4/dcps/exosc5/cyp7a1/gda/acaa1/tat/dctpp1/cyp46a1.4/tent5bb/apoea/rnaseh1/mettl3/uraha/lsm1/magoh/kmo/lsm3/pah/rexo4/ago3b/mettl16/gtpbp3/xdh/cdaa/dna2/nudt4a/fxr2/cnot8/rnaset2l/entpd5b/dnase1l4.1/hsd17b14/hmox2a/fhit/hint1/dpys/hpdb/comta/fen1/kyat3.2/pde12/allc/hgd/cnot6l/edc3/nbas/dis3/qdpra/plex9.2/rc3h2/nudt12/kynu/tet2/syncripl/endog/hmox1a/mtrex/entpd5a |
| GO:0071826 | ribonucleoprotein complex subunit organization | 1.57792704614773 | 0.0130343312989884 | nle1/gemin5/zfand1/abt1/znhit3/pwp2h/trnau1apa/mrto4/snrpd2/prmt7/pih1d1/gemin2/mrpl20/snrpg/bxdc2/prpf18/mcts1/ppan/mdn1/clp1/ptges3b/rrp7a/eif3m/nopchap1/snrpd3l/eif3ea/coil/eif6/snrpf/usp39/eif3jb/snrpe/rpl38/rps14/shq1/rpl5a/eif5/eif3ha/trnau1apb/mrps11/scaf11/rpf2/eif3s10/cdc73/denr/ddx28/naf1/rpl23a/eif3s6ip/rps15/rpsa/eif2s2/polr2d/srpk1a/rps5/gemin8/rplp0/snrpd1/rps27.1/aar2/rpl10/rpl5b/eif3g/eif3d/rpl6/yju2/rps27.2/rps28/mrps7/gemin6/eif2s3/nudt21/dyrk3/lsm4/snrpc/rps19/eif3ja/eif3c |
| GO:0055088 | lipid homeostasis | 1.54238929377417 | 0.0961825238314319 | adck1/fitm2/zgc:162608/cyp39a1/spx/abcg4b/ormdl2/apobb.1/psmd10/apobb.2/cyp7a1/apoa4a/pnpla3/tlcd1/apoa4b.2/apoba/fabp2/angptl8 |
| GO:0140014 | mitotic nuclear division | 1.53553828880172 | 0.0229731377900925 | ncaph/epgn/btc/pdcd2/nde1/chtf8/si:ch211-244o22.2/cep192/sgo1/map10/mad1l1/incenp/gnl3/kif20bb/ndc80/anapc5/gcna/ncapg/bub3/nme6/mad2l1/reep3b/anapc15/birc5b/cdca8/esco1/ncapg2/abraxas2/ccnb1/ranbp1/aurkb/racgap1/anapc11/ube2s/zgc:86764/pmaip1/bccip/abraxas1/zw10/esco2/ncaph2/cenpk/spdl1/prc1b/ran/cdca5/gnl2/map9/mki67/kifc1/ube2ib/chmp7/kif11 |
| GO:0019439 | aromatic compound catabolic process | 1.4470736645875 | 0.0239204409208845 | hnmt/nudt4b/dazl/dnase1l1/blvra/fastkd3/fitm2/zc3h12ab/fastkd2/tbrg4/nudt18/exosc2/fastkd1/exosc6/qdprb.1/entpd3/dcp1b/pde4cb/tdo2a/nudt15/moxd1l/zgc:103586/dut/heatr1/mrto4/dcps/exosc5/gda/acaa1/tat/dctpp1/tent5bb/rnaseh1/mettl3/uraha/lsm1/magoh/kmo/lsm3/pah/rexo4/ago3b/mettl16/gtpbp3/xdh/cdaa/dna2/nudt4a/fxr2/cnot8/rnaset2l/entpd5b/dnase1l4.1/hmox2a/fhit/hint1/hpdb/comta/fen1/kyat3.2/pde12/hgd/cnot6l/edc3/nbas/dis3/qdpra/plex9.2/rc3h2/nudt12/kynu/tet2/syncripl/endog/hmox1a/mtrex/entpd5a/ephx1 |
| GO:0044270 | cellular nitrogen compound catabolic process | 1.42325048500041 | 0.0467129045300582 | hnmt/nudt4b/dazl/dnase1l1/blvra/fastkd3/fitm2/zc3h12ab/fastkd2/tbrg4/nudt18/exosc2/fastkd1/exosc6/entpd3/dcp1b/pde4cb/tdo2a/nudt15/zgc:103586/dut/heatr1/mrto4/dcps/exosc5/gda/dctpp1/tent5bb/rnaseh1/mettl3/uraha/lsm1/magoh/kmo/lsm3/rexo4/ago3b/mettl16/gtpbp3/xdh/cdaa/dna2/nudt4a/fxr2/cnot8/rnaset2l/entpd5b/dnase1l4.1/hmox2a/fhit/hint1/dpys/bhmt/fen1/pde12/cnot6l/edc3/nbas/dis3/plex9.2/rc3h2/nudt12/kynu/tet2/syncripl/endog/hmox1a/mtrex/entpd5a |
| GO:0007601 | visual perception | 1.4025303608205 | 0.0873969864431149 | unc119a/gprc5c/crygm2a/crybb2/crybb1l3/crygm5/grk7a/crygm2d6/crygm2b/cryba1l2/arr3b/crygm2d9/crygm2d15/crygm2d12/crygm2d18/cryba2a/opn1sw1/crygm2d17/crygm2d7/crygm2d1/arl6/crygm2d5/crygm2f/cryba1l1/pdcb/crybb1l2/crygm2d3/crygm2d20/crygm2d11/crygm2d10/crygm2d2/crygm2d21/cryba1a/opn1mw1/crygm2d13/crygmxl2/guca1b/crygn2/crygm2d16/arr3a/cryba4/crygm2d8/crygm2d14/cryba1b/crygn1/prph2a/lum/cryba2b/opn1sw2/crygmx/pdca/crygmxl1/si:dkey-57a22.15/crybb1l1/pde6a/gucy2f/grk7b/opn1lw2/nphp3/prph2lb/pde6d/crybgx/tmtops2a |
| GO:0099537 | trans-synaptic signaling | -2.69921105199218 | 1.14653846153846e-08 | dlg1b/dtnbp1b/ncdn/gabrr2a/gabrb1/cplx2b/drd4-rs/cacng8b/otofa/sv2bb/dlg3/cdh2/penka/cep89/chrnb5a/synj1/cart2/cspg5b/gria3a/dlgap1a/chrm2a/snap25a/napbb/penkb/glrbb/slc5a7a/napba/chata/spg11/apba1b/erc1a/grid1a/stxbp1b/cplx2a/clstn2b/cacnb1/gabrb3/chrng/hdc/sncb/glra1/drd3/cacng5a/si:ch211-81a5.8/syn1/erc2/gria1a/grm4/grm6b/stx3a/caskb/chrnb3a/ppp1r9ba/slc18a3a/snap25b/jph3b/mctp2b/grm8b/cart3/slc6a5/calb1/chrna3/htr1aa/calb2b/snap47/htr1d/pdyn/jph3a/cacnb2b/ptchd1/calb2a/syt7b/syn2a/cacng7b/chrna5/grin2aa/rims1b/shisa9a/clstn1/syt4/rab3ab/dlgap4b/lrrc4ca/chrna2a/gabra4/glra4b/nlgn4xa/gria2b/nsmfb/gria4a/slc17a6b/clstn2a/sncgb/htr3a/gabrb2a/lrrc4.1/cadpsb/grin1a/grm2a/ppp1r9bb/syt11a/unc13a/cacng2b/stxbp1a/syt1a/chrnb2/slc12a5b/chrna6/cacng3b/shisa7b/nlgn3a/gria3b/stx1b/dlg2/ppfia3/grin2ab/cacng2a/grm8a/lrrc4cb/atcaya/slc12a5a/chrna4b/grm2b/nlgn3b/cacnb4b/grid2/clstn3/syt2a/apba2b/grik1a/cspg5a/sv2a/pnocb/grin1b/baiap3/cacna1ab/grin2da/nlgn2a/nrxn2a/si:dkey-174m14.3/drd2a/rph3ab/dagla/grik5/htr1fa/shisa9b/gabrp/chrm4a/grin3bb/grin2bb/insyn1/grik4/gabrg2/grm1a/cacng8a/stx11a/gabrb4/dlgap3/htr7c/neto1l/nalcn/gria4b |
| GO:0007268 | chemical synaptic transmission | -2.68729704840182 | 1.14653846153846e-08 | dlg1b/dtnbp1b/ncdn/gabrr2a/gabrb1/cplx2b/drd4-rs/cacng8b/otofa/sv2bb/dlg3/cdh2/penka/cep89/chrnb5a/synj1/cart2/cspg5b/gria3a/dlgap1a/chrm2a/snap25a/napbb/penkb/glrbb/slc5a7a/napba/chata/spg11/apba1b/erc1a/grid1a/stxbp1b/cplx2a/clstn2b/cacnb1/gabrb3/chrng/hdc/sncb/glra1/drd3/cacng5a/si:ch211-81a5.8/syn1/erc2/gria1a/grm4/grm6b/stx3a/caskb/chrnb3a/ppp1r9ba/slc18a3a/snap25b/jph3b/mctp2b/grm8b/cart3/slc6a5/calb1/chrna3/htr1aa/calb2b/snap47/htr1d/pdyn/jph3a/cacnb2b/ptchd1/calb2a/syt7b/syn2a/cacng7b/chrna5/grin2aa/rims1b/shisa9a/clstn1/syt4/rab3ab/dlgap4b/lrrc4ca/chrna2a/gabra4/glra4b/nlgn4xa/gria2b/nsmfb/gria4a/slc17a6b/clstn2a/sncgb/htr3a/gabrb2a/lrrc4.1/cadpsb/grin1a/grm2a/ppp1r9bb/syt11a/unc13a/cacng2b/stxbp1a/syt1a/chrnb2/slc12a5b/chrna6/cacng3b/shisa7b/nlgn3a/gria3b/stx1b/dlg2/ppfia3/grin2ab/cacng2a/grm8a/lrrc4cb/atcaya/slc12a5a/chrna4b/grm2b/nlgn3b/cacnb4b/grid2/clstn3/syt2a/apba2b/grik1a/cspg5a/sv2a/pnocb/grin1b/baiap3/cacna1ab/grin2da/nlgn2a/nrxn2a/si:dkey-174m14.3/drd2a/rph3ab/grik5/htr1fa/shisa9b/gabrp/chrm4a/grin3bb/grin2bb/insyn1/grik4/gabrg2/grm1a/cacng8a/stx11a/gabrb4/dlgap3/htr7c/neto1l/nalcn/gria4b |
| GO:0098916 | anterograde trans-synaptic signaling | -2.68729704840182 | 1.14653846153846e-08 | dlg1b/dtnbp1b/ncdn/gabrr2a/gabrb1/cplx2b/drd4-rs/cacng8b/otofa/sv2bb/dlg3/cdh2/penka/cep89/chrnb5a/synj1/cart2/cspg5b/gria3a/dlgap1a/chrm2a/snap25a/napbb/penkb/glrbb/slc5a7a/napba/chata/spg11/apba1b/erc1a/grid1a/stxbp1b/cplx2a/clstn2b/cacnb1/gabrb3/chrng/hdc/sncb/glra1/drd3/cacng5a/si:ch211-81a5.8/syn1/erc2/gria1a/grm4/grm6b/stx3a/caskb/chrnb3a/ppp1r9ba/slc18a3a/snap25b/jph3b/mctp2b/grm8b/cart3/slc6a5/calb1/chrna3/htr1aa/calb2b/snap47/htr1d/pdyn/jph3a/cacnb2b/ptchd1/calb2a/syt7b/syn2a/cacng7b/chrna5/grin2aa/rims1b/shisa9a/clstn1/syt4/rab3ab/dlgap4b/lrrc4ca/chrna2a/gabra4/glra4b/nlgn4xa/gria2b/nsmfb/gria4a/slc17a6b/clstn2a/sncgb/htr3a/gabrb2a/lrrc4.1/cadpsb/grin1a/grm2a/ppp1r9bb/syt11a/unc13a/cacng2b/stxbp1a/syt1a/chrnb2/slc12a5b/chrna6/cacng3b/shisa7b/nlgn3a/gria3b/stx1b/dlg2/ppfia3/grin2ab/cacng2a/grm8a/lrrc4cb/atcaya/slc12a5a/chrna4b/grm2b/nlgn3b/cacnb4b/grid2/clstn3/syt2a/apba2b/grik1a/cspg5a/sv2a/pnocb/grin1b/baiap3/cacna1ab/grin2da/nlgn2a/nrxn2a/si:dkey-174m14.3/drd2a/rph3ab/grik5/htr1fa/shisa9b/gabrp/chrm4a/grin3bb/grin2bb/insyn1/grik4/gabrg2/grm1a/cacng8a/stx11a/gabrb4/dlgap3/htr7c/neto1l/nalcn/gria4b |
| GO:0050804 | modulation of chemical synaptic transmission | -2.68416739586281 | 1.14653846153846e-08 | erc1a/grid1a/cplx2a/clstn2b/drd3/cacng5a/si:ch211-81a5.8/erc2/gria1a/grm4/grm6b/caskb/ppp1r9ba/jph3b/mctp2b/grm8b/calb1/calb2b/jph3a/calb2a/cacng7b/grin2aa/rims1b/shisa9a/clstn1/dlgap4b/lrrc4ca/nlgn4xa/gria2b/nsmfb/gria4a/clstn2a/lrrc4.1/grm2a/ppp1r9bb/cacng2b/cacng3b/shisa7b/nlgn3a/gria3b/stx1b/grin2ab/cacng2a/grm8a/lrrc4cb/atcaya/grm2b/nlgn3b/grid2/clstn3/grik1a/baiap3/grin2da/nlgn2a/si:dkey-174m14.3/drd2a/grik5/shisa9b/grin3bb/grin2bb/grik4/grm1a/cacng8a/dlgap3/neto1l/nalcn/gria4b |
| GO:0099177 | regulation of trans-synaptic signaling | -2.68416739586281 | 1.14653846153846e-08 | erc1a/grid1a/cplx2a/clstn2b/drd3/cacng5a/si:ch211-81a5.8/erc2/gria1a/grm4/grm6b/caskb/ppp1r9ba/jph3b/mctp2b/grm8b/calb1/calb2b/jph3a/calb2a/cacng7b/grin2aa/rims1b/shisa9a/clstn1/dlgap4b/lrrc4ca/nlgn4xa/gria2b/nsmfb/gria4a/clstn2a/lrrc4.1/grm2a/ppp1r9bb/cacng2b/cacng3b/shisa7b/nlgn3a/gria3b/stx1b/grin2ab/cacng2a/grm8a/lrrc4cb/atcaya/grm2b/nlgn3b/grid2/clstn3/grik1a/baiap3/grin2da/nlgn2a/si:dkey-174m14.3/drd2a/grik5/shisa9b/grin3bb/grin2bb/grik4/grm1a/cacng8a/dlgap3/neto1l/nalcn/gria4b |
| GO:0099536 | synaptic signaling | -2.6833837570563 | 1.14653846153846e-08 | dlg1b/dtnbp1b/ncdn/gabrr2a/gabrb1/cplx2b/drd4-rs/cacng8b/otofa/sv2bb/dlg3/cdh2/penka/cep89/chrnb5a/synj1/cart2/cspg5b/gria3a/dlgap1a/chrm2a/snap25a/napbb/penkb/glrbb/slc5a7a/napba/chata/spg11/apba1b/erc1a/grid1a/stxbp1b/cplx2a/clstn2b/cacnb1/gabrb3/chrng/hdc/sncb/glra1/drd3/cacng5a/si:ch211-81a5.8/syn1/erc2/gria1a/grm4/grm6b/stx3a/caskb/chrnb3a/ppp1r9ba/slc18a3a/snap25b/jph3b/mctp2b/grm8b/cart3/slc6a5/calb1/chrna3/htr1aa/calb2b/snap47/htr1d/pdyn/jph3a/cacnb2b/ptchd1/calb2a/syt7b/syn2a/cacng7b/chrna5/grin2aa/rims1b/shisa9a/clstn1/syt4/rab3ab/dlgap4b/lrrc4ca/chrna2a/gabra4/glra4b/nlgn4xa/gria2b/nsmfb/gria4a/slc17a6b/clstn2a/sncgb/htr3a/gabrb2a/lrrc4.1/cadpsb/grin1a/grm2a/ppp1r9bb/syt11a/unc13a/cacng2b/stxbp1a/syt1a/chrnb2/slc12a5b/chrna6/cacng3b/shisa7b/nlgn3a/gria3b/stx1b/dlg2/ppfia3/grin2ab/cacng2a/grm8a/lrrc4cb/atcaya/slc12a5a/chrna4b/grm2b/nlgn3b/cacnb4b/grid2/clstn3/syt2a/apba2b/grik1a/cspg5a/sv2a/pnocb/grin1b/baiap3/cacna1ab/grin2da/nlgn2a/nrxn2a/si:dkey-174m14.3/drd2a/rph3ab/dagla/grik5/htr1fa/shisa9b/gabrp/chrm4a/grin3bb/grin2bb/insyn1/grik4/gabrg2/grm1a/cacng8a/stx11a/gabrb4/scn1lab/dlgap3/htr7c/neto1l/nalcn/gria4b |
| GO:0098742 | cell-cell adhesion via plasma-membrane adhesion molecules | -2.34722281326859 | 1.14653846153846e-08 | arvcfb/zeb1a/cdh2/prnprs3/ptk7a/pcdh2ab10/zgc:165604/si:ch211-186j3.6/tenm3/dchs1a/ret/pcdh8/ptprsa/clstn2b/dscama/pcdh17/pcdh2ab9/pecam1a/pcdh10b/pcdh7a/tenm2b/robo3/si:ch211-199f5.1/dsc2l/pcdh1g22/jag2b/nectin3b/tenm4/pcdh1g31/pcdh7b/pcdh1g29/pcdh2g13/pcdh1a/clstn1/pcdh19/cadm1b/lrrc4ca/pcdh1a4/pcdh2ab3/cadm3/dscamb/ptprdb/pcdh10a/pcdh11/clstn2a/lrrc4.1/igsf21b/robo4/fat3a/cadm4/si:ch211-214p13.3/cdh18a/si:dkey-237h12.3/lrrc4cb/igsf21a/tenm2a/esama/cdh13/clstn3/cdh15/cntn3a.1/si:ch73-233f7.1/pcdh1g32/cdh4/pcdh1g11/si:ch73-233f7.3/cdh7a/pcdh1g18/adgrl1a/pcdh1gc5 |
| GO:0050808 | synapse organization | -2.21669824394954 | 9.94928637272983e-08 | dbn1/ctnna2/cdh2/synj1/ptk7a/dbnlb/cbln2b/ephb3a/sez6a/tjp1b/ctbp2a/erc1a/fbxo45/gphnb/neurod2/ptprsa/clstn2b/cacnb1/sncb/erc2/tanc2a/ppfia4/gfpt1/slitrk4/gphna/cacnb2b/myo9ab/inpp5ka/caprin2/pcloa/clstn1/rab3ab/lrrc4ca/ptprdb/nlgn4xa/clstn2a/sncgb/sybu/lrrc4.1/arhgef9b/unc13a/dock10/ctnnd2b/nbeaa/nlgn3a/bsnb/cacna1da/ppfia3/lrrc4cb/epb41l3a/grna/bdnf/nlgn3b/ctnnd2a/cacnb4b/pclob/clstn3/slitrk2/sez6l2/nlgn2a/nrxn2a |
| GO:0048667 | cell morphogenesis involved in neuron differentiation | -2.08310048291656 | 1.14653846153846e-08 | plxnb1b/pik3cd/mycbp2/rb1/ptprfb/zgc:100846/ptprfa/isl2b/nrp2b/arhgap35a/rapgef1a/whrna/emb/pik3ca/rbpms2a/aplp2/mapk8ip3/npr1a/sema3fa/emx2/olfm1a/unc5b/nptna/vegfc/dbn1/kctd12.1/si:ch211-129c21.1/phox2bb/unk/ubap1/ctnna2/cntn4/cdh2/trarg1a/kalrnb/ulk2/ptk7a/dbnlb/efna1b/ephb3a/slc35b2/rbpms2b/fus/map1ab/dock7/foxb1a/nsfa/rab33ba/sipa1l1/nrp2a/ush1ga/pnpla6/kctd13/isl1a/fezf2/slit2/tenm3/rufy3/nfasca/hecw1b/sim1a/adcy8/pak1/spg11/ret/kif1b/arnt2/sema3bl/zdhhc17/plxna4/appa/nedd4l/arhgap35b/efna2a/dscama/apc/robo2/pcdh17/casp9/epha4a/hdc/anos1a/trioa/lhx9/epha7/rtn4r/cyfip2/dpysl3/draxinb/slit1b/robo3/usp33/map1sb/appb/tanc2a/dzip1/sema5a/kifbp/chad/robo1/tnr/whrnb/tenm4/hecw2a/epha4l/kif5c/slitrk4/islr2/nptnb/lhx2b/myo9ab/caprin2/kalrna/cdkl5/ephb2b/epha4b/clstn1/rac3a/brsk1b/aplp1/rab3ab/gap43/lrrc4ca/srrm4/lhx1a/map1aa/prmt8b/atl1/nkx2.2a/arhgef25b/dscamb/triob/tmem59l/myo10l3/nsmfb/slit3/dpysl5a/chst11/fzd3a/kif5aa/robo4/efnb3b/vax1/stat3/dock10/ctnnd2b/zgc:77784/sema4c/si:ch211-113g11.6/rac1l/elavl4/myo10l1/ntn1b/scn8aa/lrrc4cb/plppr4a/grna/bdnf/ctnnd2a/slitrk2/nkx6.1/ncam1b/cntn3a.1/nrxn2a/dagla/grxcr1a/efna3a/dpysl5b/thsd7aa/si:ch211-241e1.3/plxna3/l1camb/plxna1a/nrcama/sema6ba/sema3b |
| GO:0048858 | cell projection morphogenesis | -2.05831448361924 | 1.14653846153846e-08 | plxnb1b/tmem108/pik3cd/mycbp2/rb1/ptprfb/zgc:100846/ptprfa/spag6/isl2b/nrp2b/arhgap35a/rapgef1a/emb/pik3ca/rbpms2a/aplp2/mapk8ip3/npr1a/sema3fa/olfm1a/unc5b/nptna/vegfc/dbn1/kctd12.1/si:ch211-129c21.1/phox2bb/ubap1/ctnna2/cntn4/cdh2/trarg1a/kalrnb/ulk2/ptk7a/dbnlb/cspg5b/efna1b/ephb3a/slc35b2/rbpms2b/fus/map1ab/dock7/foxb1a/nsfa/rab33ba/sipa1l1/tnika/nrp2a/pnpla6/kctd13/isl1a/fezf2/mink1/slit2/tenm3/rufy3/nfasca/hecw1b/sim1a/adcy8/pak1/spg11/ret/kif1b/arnt2/sema3bl/zdhhc17/plxna4/appa/nedd4l/arhgap35b/efna2a/dscama/apc/robo2/pcdh17/casp9/epha4a/hdc/anos1a/trioa/lhx9/epha7/rtn4r/cyfip2/dpysl3/draxinb/slit1b/robo3/usp33/map1sb/appb/tanc2a/dzip1/nyap2b/sema5a/kifbp/chad/pacsin1a/robo1/tnr/tenm4/hecw2a/epha4l/kif5c/slitrk4/islr2/nptnb/lhx2b/myo9ab/caprin2/kalrna/cdkl5/ephb2b/epha4b/clstn1/rac3a/brsk1b/aplp1/rab3ab/gap43/lrrc4ca/srrm4/lhx1a/map1aa/prmt8b/atl1/nkx2.2a/arhgef25b/dscamb/triob/tmem59l/myo10l3/gas7a/nsmfb/slit3/dpysl5a/chst11/fzd3a/kif5aa/robo4/efnb3b/vax1/stat3/dock10/ctnnd2b/zgc:77784/sema4c/si:ch211-113g11.6/rac1l/elavl4/myo10l1/ntn1b/scn8aa/lrrc4cb/plppr4a/grna/bdnf/ctnnd2a/nrn1la/slitrk2/nkx6.1/cspg5a/ncam1b/cntn3a.1/nrxn2a/dagla/efna3a/dpysl5b/inab/thsd7aa/si:ch211-241e1.3/plxna3/l1camb/plxna1a/nrcama/sema6ba/sema3b |
| GO:0048812 | neuron projection morphogenesis | -2.04924995520432 | 1.14653846153846e-08 | plxnb1b/tmem108/pik3cd/mycbp2/rb1/ptprfb/zgc:100846/ptprfa/spag6/isl2b/nrp2b/arhgap35a/rapgef1a/emb/pik3ca/rbpms2a/aplp2/mapk8ip3/npr1a/sema3fa/olfm1a/unc5b/nptna/vegfc/dbn1/kctd12.1/si:ch211-129c21.1/phox2bb/ubap1/ctnna2/cntn4/cdh2/trarg1a/kalrnb/ulk2/ptk7a/dbnlb/efna1b/ephb3a/slc35b2/rbpms2b/fus/map1ab/dock7/foxb1a/nsfa/rab33ba/sipa1l1/tnika/nrp2a/pnpla6/kctd13/isl1a/fezf2/mink1/slit2/tenm3/rufy3/nfasca/hecw1b/sim1a/adcy8/pak1/spg11/ret/kif1b/arnt2/sema3bl/zdhhc17/plxna4/appa/nedd4l/arhgap35b/efna2a/dscama/apc/robo2/pcdh17/casp9/epha4a/hdc/anos1a/trioa/lhx9/epha7/rtn4r/cyfip2/dpysl3/draxinb/slit1b/robo3/usp33/map1sb/appb/tanc2a/dzip1/nyap2b/sema5a/kifbp/chad/pacsin1a/robo1/tnr/tenm4/hecw2a/epha4l/kif5c/slitrk4/islr2/nptnb/lhx2b/myo9ab/caprin2/kalrna/cdkl5/ephb2b/epha4b/clstn1/rac3a/brsk1b/aplp1/rab3ab/gap43/lrrc4ca/srrm4/lhx1a/map1aa/prmt8b/atl1/nkx2.2a/arhgef25b/dscamb/triob/tmem59l/myo10l3/gas7a/nsmfb/slit3/dpysl5a/chst11/fzd3a/kif5aa/robo4/efnb3b/vax1/stat3/dock10/ctnnd2b/zgc:77784/sema4c/si:ch211-113g11.6/rac1l/elavl4/myo10l1/ntn1b/scn8aa/lrrc4cb/plppr4a/grna/bdnf/ctnnd2a/nrn1la/slitrk2/nkx6.1/ncam1b/cntn3a.1/nrxn2a/dagla/efna3a/dpysl5b/inab/thsd7aa/si:ch211-241e1.3/plxna3/l1camb/plxna1a/nrcama/sema6ba/sema3b |
| GO:0120039 | plasma membrane bounded cell projection morphogenesis | -2.0490209096944 | 1.14653846153846e-08 | plxnb1b/tmem108/pik3cd/mycbp2/rb1/ptprfb/zgc:100846/ptprfa/spag6/isl2b/nrp2b/arhgap35a/rapgef1a/emb/pik3ca/rbpms2a/aplp2/mapk8ip3/npr1a/sema3fa/olfm1a/unc5b/nptna/vegfc/dbn1/kctd12.1/si:ch211-129c21.1/phox2bb/ubap1/ctnna2/cntn4/cdh2/trarg1a/kalrnb/ulk2/ptk7a/dbnlb/efna1b/ephb3a/slc35b2/rbpms2b/fus/map1ab/dock7/foxb1a/nsfa/rab33ba/sipa1l1/tnika/nrp2a/pnpla6/kctd13/isl1a/fezf2/mink1/slit2/tenm3/rufy3/nfasca/hecw1b/sim1a/adcy8/pak1/spg11/ret/kif1b/arnt2/sema3bl/zdhhc17/plxna4/appa/nedd4l/arhgap35b/efna2a/dscama/apc/robo2/pcdh17/casp9/epha4a/hdc/anos1a/trioa/lhx9/epha7/rtn4r/cyfip2/dpysl3/draxinb/slit1b/robo3/usp33/map1sb/appb/tanc2a/dzip1/nyap2b/sema5a/kifbp/chad/pacsin1a/robo1/tnr/tenm4/hecw2a/epha4l/kif5c/slitrk4/islr2/nptnb/lhx2b/myo9ab/caprin2/kalrna/cdkl5/ephb2b/epha4b/clstn1/rac3a/brsk1b/aplp1/rab3ab/gap43/lrrc4ca/srrm4/lhx1a/map1aa/prmt8b/atl1/nkx2.2a/arhgef25b/dscamb/triob/tmem59l/myo10l3/gas7a/nsmfb/slit3/dpysl5a/chst11/fzd3a/kif5aa/robo4/efnb3b/vax1/stat3/dock10/ctnnd2b/zgc:77784/sema4c/si:ch211-113g11.6/rac1l/elavl4/myo10l1/ntn1b/scn8aa/lrrc4cb/plppr4a/grna/bdnf/ctnnd2a/nrn1la/slitrk2/nkx6.1/ncam1b/cntn3a.1/nrxn2a/dagla/efna3a/dpysl5b/inab/thsd7aa/si:ch211-241e1.3/plxna3/l1camb/plxna1a/nrcama/sema6ba/sema3b |
| GO:0032990 | cell part morphogenesis | -2.03560178882976 | 1.14653846153846e-08 | plxnb1b/tmem108/pik3cd/mycbp2/rb1/ptprfb/zgc:100846/ptprfa/spag6/isl2b/nrp2b/arhgap35a/rapgef1a/emb/pik3ca/rbpms2a/aplp2/mapk8ip3/npr1a/sema3fa/olfm1a/unc5b/nptna/vegfc/dbn1/kctd12.1/si:ch211-129c21.1/phox2bb/ubap1/ctnna2/cntn4/cdh2/trarg1a/kalrnb/ulk2/ptk7a/dbnlb/supv3l1/cspg5b/efna1b/ephb3a/slc35b2/rbpms2b/fus/map1ab/dock7/foxb1a/nsfa/rab33ba/sipa1l1/tnika/nrp2a/pnpla6/kctd13/isl1a/fezf2/mink1/slit2/tenm3/rufy3/nfasca/hecw1b/sim1a/adcy8/pak1/spg11/ret/kif1b/arnt2/sema3bl/zdhhc17/plxna4/appa/nedd4l/arhgap35b/efna2a/dscama/apc/robo2/pcdh17/casp9/epha4a/hdc/anos1a/trioa/lhx9/epha7/rtn4r/cyfip2/dpysl3/draxinb/slit1b/robo3/usp33/map1sb/appb/tanc2a/dzip1/nyap2b/sema5a/kifbp/chad/pacsin1a/robo1/tnr/tenm4/hecw2a/epha4l/kif5c/slitrk4/islr2/nptnb/lhx2b/myo9ab/caprin2/kalrna/cdkl5/ephb2b/epha4b/clstn1/rac3a/brsk1b/aplp1/rab3ab/gap43/lrrc4ca/srrm4/lhx1a/map1aa/prmt8b/atl1/nkx2.2a/arhgef25b/dscamb/triob/tmem59l/myo10l3/gas7a/nsmfb/slit3/dpysl5a/chst11/fzd3a/kif5aa/robo4/efnb3b/vax1/stat3/dock10/ctnnd2b/zgc:77784/sema4c/si:ch211-113g11.6/rac1l/elavl4/myo10l1/ntn1b/scn8aa/lrrc4cb/plppr4a/grna/bdnf/ctnnd2a/nrn1la/slitrk2/nkx6.1/cspg5a/ncam1b/cntn3a.1/nrxn2a/dagla/efna3a/dpysl5b/inab/thsd7aa/si:ch211-241e1.3/plxna3/l1camb/plxna1a/nrcama/sema6ba/sema3b |
| GO:0000904 | cell morphogenesis involved in differentiation | -2.02983448887318 | 1.14653846153846e-08 | pik3cd/mycbp2/rb1/ptprfb/adgrg1/zgc:100846/ptprfa/isl2b/nrp2b/arhgap35a/rapgef1a/whrna/emb/pik3ca/erbb2/rbpms2a/aplp2/mapk8ip3/npr1a/sema3fa/emx2/olfm1a/unc5b/nptna/vegfc/dbn1/kctd12.1/si:ch211-129c21.1/phox2bb/unk/ubap1/ctnna2/cntn4/cdh2/trarg1a/kalrnb/ulk2/ptk7a/dbnlb/efna1b/rilpl2/ephb3a/slc35b2/rbpms2b/fus/map1ab/dock7/foxb1a/nsfa/rab33ba/sipa1l1/nrp2a/ush1ga/pnpla6/kctd13/isl1a/fezf2/slit2/tenm3/rufy3/nfasca/hecw1b/sim1a/adcy8/pak1/spg11/ret/kif1b/arnt2/sema3bl/zdhhc17/plxna4/appa/nedd4l/arhgap35b/efna2a/dscama/apc/robo2/pcdh17/casp9/epha4a/hdc/anos1a/trioa/lhx9/epha7/rtn4r/cyfip2/dpysl3/draxinb/slit1b/robo3/usp33/map1sb/appb/tanc2a/dzip1/sema5a/kifbp/chad/robo1/tnr/whrnb/tenm4/hecw2a/epha4l/kif5c/slitrk4/islr2/nptnb/lhx2b/tet3/myo9ab/caprin2/kalrna/cdkl5/ephb2b/epha4b/clstn1/rac3a/brsk1b/aplp1/rab3ab/gap43/lrrc4ca/srrm4/lhx1a/map1aa/prmt8b/atl1/nkx2.2a/arhgef25b/dscamb/triob/tmem59l/myo10l3/nsmfb/slit3/cxcl8a/dpysl5a/chst11/fzd3a/kif5aa/robo4/efnb3b/vax1/stat3/dock10/ctnnd2b/zgc:77784/sema4c/si:ch211-113g11.6/rac1l/elavl4/myo10l1/ntn1b/scn8aa/lrrc4cb/plppr4a/grna/bdnf/ctnnd2a/slitrk2/nkx6.1/ncam1b/cntn3a.1/nrxn2a/dagla/grxcr1a/efna3a/dpysl5b/thsd7aa/si:ch211-241e1.3/plxna3/l1camb/plxna1a/nrcama/sema6ba/sema3b |
| GO:0007409 | axonogenesis | -2.02083525635161 | 1.14653846153846e-08 | plxnb1b/pik3cd/mycbp2/rb1/ptprfb/zgc:100846/ptprfa/isl2b/nrp2b/arhgap35a/rapgef1a/emb/pik3ca/rbpms2a/aplp2/mapk8ip3/npr1a/sema3fa/olfm1a/unc5b/nptna/vegfc/dbn1/si:ch211-129c21.1/phox2bb/ubap1/ctnna2/cntn4/cdh2/trarg1a/kalrnb/ulk2/ptk7a/dbnlb/efna1b/ephb3a/slc35b2/rbpms2b/fus/map1ab/dock7/foxb1a/nsfa/rab33ba/nrp2a/pnpla6/kctd13/isl1a/fezf2/slit2/tenm3/rufy3/nfasca/sim1a/adcy8/pak1/spg11/ret/kif1b/arnt2/sema3bl/zdhhc17/plxna4/appa/arhgap35b/efna2a/dscama/apc/robo2/pcdh17/casp9/epha4a/hdc/anos1a/trioa/lhx9/epha7/rtn4r/cyfip2/dpysl3/draxinb/slit1b/robo3/usp33/map1sb/appb/dzip1/sema5a/kifbp/chad/robo1/tnr/tenm4/epha4l/kif5c/slitrk4/islr2/nptnb/lhx2b/myo9ab/kalrna/cdkl5/ephb2b/epha4b/clstn1/rac3a/brsk1b/aplp1/rab3ab/gap43/lrrc4ca/srrm4/lhx1a/map1aa/atl1/nkx2.2a/arhgef25b/dscamb/triob/tmem59l/myo10l3/nsmfb/slit3/dpysl5a/chst11/fzd3a/kif5aa/robo4/efnb3b/vax1/stat3/zgc:77784/sema4c/si:ch211-113g11.6/rac1l/elavl4/myo10l1/ntn1b/scn8aa/lrrc4cb/plppr4a/grna/bdnf/slitrk2/nkx6.1/ncam1b/cntn3a.1/nrxn2a/dagla/efna3a/dpysl5b/thsd7aa/si:ch211-241e1.3/plxna3/l1camb/plxna1a/nrcama/sema6ba/sema3b |
| GO:0001505 | regulation of neurotransmitter levels | -2.01367130950093 | 0.000141537068751352 | snap25a/napbb/slc5a7a/napba/chata/stxbp1b/cplx2a/comtb/si:ch211-81a5.8/dbh/syn1/stx3a/caskb/snap25b/mctp2b/snap47/syt7b/syn2a/rims1b/gad1b/syt4/rab3ab/moxd1/cadpsb/syt11a/unc13a/stxbp1a/syt1a/ppfia3/gad2/syt2a/baiap3/hrh3/rph3ab/th2/stx11a |
| GO:0061564 | axon development | -1.97936421112892 | 1.14653846153846e-08 | plxnb1b/pik3cd/mycbp2/rb1/ptprfb/zgc:100846/plp1b/ptprfa/isl2b/nrp2b/arhgap35a/rapgef1a/emb/pik3ca/erbb2/rbpms2a/aplp2/mapk8ip3/npr1a/sema3fa/olfm1a/unc5b/nptna/vegfc/dbn1/gpm6bb/si:ch211-129c21.1/phox2bb/ubap1/ctnna2/cntn4/cdh2/trarg1a/kalrnb/ulk2/ptk7a/dbnlb/nfia/efna1b/ephb3a/slc35b2/rbpms2b/fus/map1ab/dock7/foxb1a/nsfa/rab33ba/nrp2a/pnpla6/kctd13/isl1a/fezf2/gpm6ba/slit2/tenm3/rufy3/nfasca/sim1a/adcy8/pak1/spg11/ret/kif1b/arnt2/sema3bl/zdhhc17/plxna4/appa/arhgap35b/efna2a/dscama/apc/robo2/pcdh17/casp9/epha4a/hdc/anos1a/trioa/lhx9/epha7/rtn4r/cyfip2/dpysl3/draxinb/slit1b/robo3/usp33/map1sb/appb/dzip1/sema5a/kifbp/chad/il6st/robo1/tnr/tenm4/epha4l/kif5c/slitrk4/islr2/nptnb/lhx2b/myo9ab/kalrna/cdkl5/ephb2b/epha4b/ek1/clstn1/rac3a/brsk1b/aplp1/rab3ab/gap43/lrrc4ca/srrm4/lhx1a/map1aa/atl1/nkx2.2a/arhgef25b/dscamb/triob/tmem59l/myo10l3/nsmfb/slit3/dpysl5a/sarm1/chst11/fzd3a/kif5aa/robo4/efnb3b/vax1/stat3/zgc:77784/sema4c/si:ch211-113g11.6/rac1l/elavl4/myo10l1/ntn1b/scn8aa/lrrc4cb/plppr4a/grna/bdnf/slitrk2/nkx6.1/ncam1b/cntn3a.1/nrxn2a/dagla/efna3a/dpysl5b/thsd7aa/si:ch211-241e1.3/plxna3/l1camb/plxna1a/nrcama/sema6ba/sema3b |
| GO:0051049 | regulation of transport | -1.97152747166447 | 1.14653846153846e-08 | prnprs3/gem/kcne4/kcna1a/unc45a/dmxl2/mffa/scn5lab/mcoln1a/pxk/abcg1/kcnj13/rab33ba/fgf13b/cep290/c2cd5/cldnb/abca1a/hecw1b/cacna1ia/kctd7/nedd4l/cplx2a/chp2/kcnab1b/si:cabz01076231.1/cacnb1/bmp2k/cacna1fb/mcu/si:ch211-204c21.1/fgf13a/msnb/kcng1/cacng5a/cacna1ha/si:ch211-81a5.8/stim2b/cnstb/syt6b/stxbp5b/c2cd2l/caskb/fxyd6/crhbp/scn1ba/kcnab2a/pacsin1a/mctp2b/fgf12a/plcb3/rab3c/htr1aa/ank3a/dnajc5aa/hecw2a/ptprn2/rab3da/cacnb2b/syt7b/cacng7b/rab3b/rims1b/shisa9a/syt9b/zgc:91999/syt4/rab3ab/kcnj3b/kcnc2/kcnj19a/sgip1b/cadpsb/scn3b/rrad/syt13/stxbp5a/syt11a/cacng2b/kcnj1a.1/syt1a/cacng3b/shisa7b/syt9a/syt10/ptprna/kcnc1b/stx1b/cacna1da/stxbp4/anxa1c/scn8aa/cacng2a/rab44/stc1l/cacna1c/cacnb4b/fgf11b/cacna2d3/kcnj14/kcnj19b/syt2a/nkx6.1/kcnd3/il1rapl1a/syt3/cabp2a/baiap3/cacna1ab/kcnc1a/rph3ab/shisa9b/kcnd2/cnih2/npvf/kcnab2b/trh/cacng8a/cacna2d2b/scn1lab/kcnj5/syt6a/cacna2d4b/kcnh4a/kcnb1 |
| GO:0032879 | regulation of localization | -1.91598630922618 | 1.14653846153846e-08 | dtnbp1b/scn4ba/cplx2b/plcg1/gpr137ba/cacng8b/ptprnb/hip1rb/kcnh7/efcab7/scn8ab/aplnrb/adcyap1b/prnprs3/gem/kcne4/kcna1a/unc45a/letm1/dmxl2/mffa/scn5lab/mcoln1a/pxk/abcg1/kcnj13/rab33ba/fgf13b/cep290/c2cd5/cldnb/abca1a/hecw1b/cacna1ia/kctd7/nedd4l/cplx2a/chp2/plin2/kcnab1b/si:cabz01076231.1/cacnb1/bmp2k/cacna1fb/mcu/si:ch211-204c21.1/fgf13a/msnb/kcng1/plch1/cacng5a/cacna1ha/si:ch211-81a5.8/stim2b/cnstb/syt6b/stxbp5b/c2cd2l/caskb/fxyd6/crhbp/scn1ba/kcnab2a/pacsin1a/mctp2b/fgf12a/plcb3/rab3c/htr1aa/ank3a/dnajc5aa/hecw2a/wnt5b/ptprn2/rab3da/cacnb2b/plce1/plch2a/syt7b/cacng7b/rab3b/rims1b/shisa9a/syt9b/zgc:91999/syt4/gpc6a/rab3ab/kcnj3b/kcnc2/kcnj19a/sgip1b/zgc:77486/cadpsb/scn3b/rrad/syt13/stxbp5a/syt11a/cacng2b/kcnj1a.1/syt1a/gpc5b/cacng3b/shisa7b/syt9a/syt10/ptprna/kcnc1b/stx1b/cacna1da/stxbp4/anxa1c/scn8aa/cacng2a/rab44/stc1l/cacna1c/cacnb4b/fgf11b/cacna2d3/kcnj14/kcnj19b/syt2a/nkx6.1/kcnd3/il1rapl1a/syt3/cabp2a/baiap3/cacna1ab/kcnc1a/rph3ab/shisa9b/kcnd2/si:ch211-13f8.1/cnih2/npvf/kcnab2b/trh/cacng8a/cacna2d2b/scn1lab/kcnj5/syt6a/cacna2d4b/kcnh4a/kcnb1 |
| GO:1901701 | cellular response to oxygen-containing compound | -1.89692649413327 | 1.56824905636127e-05 | zgc:172120/pik3r3b/myd88/carf/stat6/nr4a2b/cpeb4a/chrng/glra1/syt6b/il13/c2cd2l/phip/htr1aa/ptger4c/ticam1/htr1d/ptprn2/tet3/gnao1b/inpp5ka/syt7b/syt9b/syt4/insig1/esr2b/ptger4a/cpne4a/vtcn1/cxcl8a/cxcl8b.3/nr4a2a/syt13/syt11a/stat3/nsg2/syt1a/sstr1a/syt9a/syt10/gnao1a/ptprna/cpne2/stxbp4/adgrv1/anxa1c/nr4a3/inpp5kb/syt2a/syt3/cpne7/irs2b/drd2a/htr1fa/chrm4a/gnb5a/cpne8/syt6a/il1b/gper1/cxcl18b |
| GO:0048640 | negative regulation of developmental growth | -1.88113815885922 | 0.0105693423015173 | sema6d/sema6e/dusp6/sema3fa/si:ch211-129c21.1/ulk2/slit2/sema3bl/robo2/epha4a/wwc1/sema5a/epha4b/slit3/sema4c/si:ch211-113g11.6/wwc3/sema6ba/sema3b |
| GO:0007019 | microtubule depolymerization | -1.8750187848337 | 0.021391144294225 | map1ab/stmn4/apc/map1b/stmn2a/map1sb/stmn2b/stmn3/map1aa/camsap1a/stmn4l/apc2 |
| GO:0032409 | regulation of transporter activity | -1.86261008963063 | 0.00378678429698509 | cabp1a/cabp5a/scn4ba/cacng8b/prnprs3/gem/kcne4/fgf13b/hecw1b/kctd7/nedd4l/chp2/kcnab1b/si:cabz01076231.1/cacnb1/fgf13a/kcng1/cacng5a/stim2b/fxyd6/fgf12a/ank3a/hecw2a/cacnb2b/cacng7b/shisa9a/rrad/cacng2b/cacng3b/shisa7b/cacng2a/cacnb4b/fgf11b/cabp2a/shisa9b/cnih2/cacng8a |
| GO:0003310 | pancreatic A cell differentiation | -1.86173412990953 | 0.00748525766259099 | isl1a/pax6b/insm1b/nkx6.2/myt1b/nkx6.1/arxa |
| GO:0036269 | swimming behavior | -1.85563104344825 | 0.0118323095206783 | dmrt3a/myo9aa/sin3b/panx1b/kif2a/gdpd5a/zgc:100846/ca10a/klhl41a/spg11/scdb/micall2b/ca10b/myo9ab/cdkl5/grna/tcap/cacna1ab/dagla/syngap1b/gnb5a/scn1lab/dnmt3aa/oxt |
| GO:0019226 | transmission of nerve impulse | -1.84484111835001 | 0.0190096470355517 | cacng8b/scn8ab/cacna1ia/cacng5a/cacna1ha/ank3a/cacng7b/cacng2b/cacng3b/scn8aa/cacng2a/kcnab2b/cacng8a/scn1lab |
| GO:0021545 | cranial nerve development | -1.80828563846919 | 0.00979204145463202 | rb1/adgrg1/klf6a/hgfa/mib1/erbb2/nes/tuba1a/nsfa/six3a/slit2/kif1b/chn1/robo2/klf7b/slit1b/six3b/kifbp/diras1a/lhx2b/vax1/mdga2a/cdh4/foxi1 |
| GO:0007186 | G protein-coupled receptor signaling pathway | -1.79358601966978 | 1.60681800279223e-06 | penka/gpr27/rgs18/trpm1b/gpr78a/cart2/opn1mw2/gnb1a/gpr182/npy/chrm2a/rgrb/penkb/gng3/glrbb/adcy2a/gal/adcy6b/adcy8/adgre15/ptgir/f2rl2/rgra/grm5a/gpr22a/gnal/lpar1/or95a1/gpr35.1/s1pr1/glra1/ccl38.6/gnai2b/tmem116/drd3/oprd1b/aplnr2/gpr22b/usp33/grm4/grm6b/npffl/lmbrd2b/rgs7a/ccl34b.4/grm8b/cart3/gpr174/htr1aa/ptger4c/tafa5a/gpr61l/adgrf3b/pde6ga/htr1d/pdyn/gnb2/si:ch211-117l17.6/gnao1b/adgra2/xcr1a.1/gnb3a/ccl44/gnb4b/agtr2/rgs9b/mrap2a/gpr63/ptger4a/adra2da/opn6b/prlhr2a/adcy6a/nmur3/adcy2b/adgrl3.1/adcy1a/si:dkey-6n21.13/grin1a/grm2a/tas1r2.1/sstr1a/gabbr1b/pth2/adra2b/trpm1a/git2b/tmtops2b/gnao1a/gpr85/gpr158b/adgrv1/ccl35.1/si:dkey-172j4.3/adcy5/grm8a/tshba/adgra1a/tafa5b/grm2b/prlhr2b/gpr75/vipr1b/adgrb3/pnocb/adgre5b.2/hrh3/adrb1/rgs11/drd2a/dagla/adgrb1a/htr1fa/npb/si:ch211-132p1.2/kctd8/chrm4a/ltb4r2a/crhr1/gpr158a/npvf/ltb4r/grm1a/adcyap1r1a/pomca/adgrg3/gnb5a/rxfp1/tac3b/htr7c/adgrl1a/adgrb1b/gper1/or92a7/p2ry13 |
| GO:0010721 | negative regulation of cell development | -1.7837780436552 | 0.00979204145463202 | sema3fa/gpr137ba/si:ch211-129c21.1/irf2bp2b/spi1b/ulk2/slit2/sema3bl/tmem178a/robo2/epha4a/bmp15/sema5a/epha4b/slit3/sema4c/si:ch211-113g11.6/anxa1c/mmp9/sema6ba/gper1/sema3b |
| GO:0006935 | chemotaxis | -1.78340911781747 | 1.1812068993636e-05 | unc5b/nptna/vegfc/tnfrsf1a/si:ch211-129c21.1/cntn4/cdh2/kalrnb/maco1a/tdgf1/ptk7a/efna1b/ephb3a/slc35b2/rbpms2b/rab33ba/nrp2a/fezf2/slit2/tenm3/maco1b/nfasca/sim1a/adcy8/kif1b/arnt2/sema3bl/myd88/plxna4/efna2a/dscama/apc/robo2/pcdh17/hdc/anos1a/trioa/s1pr1/lhx9/epha7/ccl38.6/rtn4r/cyfip2/dpysl3/draxinb/slit1b/robo3/usp33/aanat2/dock4b/dzip1/trim33/ccl34b.4/sema5a/chad/vegfd/dock4/robo1/tnr/tenm4/epha4l/kif5c/islr2/nptnb/lhx2b/kalrna/xcr1a.1/ccl44/ephb2b/rac3a/gap43/lhx1a/atl1/nkx2.2a/arhgef25b/dscamb/triob/slit3/cxcl8a/dpysl5a/cxcl8b.3/chst11/fzd3a/kif5aa/si:dkey-12h9.6/robo4/efnb3b/vax1/stat3/zgc:77784/sema4c/si:ch211-113g11.6/rac1l/anxa1c/ntn1b/scn8aa/ccl35.1/cxcl8b.1/ncam1b/cntn3a.1/nrxn2a/efna3a/dpysl5b/si:ch211-241e1.3/plxna3/l1camb/plxna1a/nrcama/sema6ba/il1b/sema3b/cxcl18b |
| GO:0042330 | taxis | -1.78340911781747 | 1.1812068993636e-05 | unc5b/nptna/vegfc/tnfrsf1a/si:ch211-129c21.1/cntn4/cdh2/kalrnb/maco1a/tdgf1/ptk7a/efna1b/ephb3a/slc35b2/rbpms2b/rab33ba/nrp2a/fezf2/slit2/tenm3/maco1b/nfasca/sim1a/adcy8/kif1b/arnt2/sema3bl/myd88/plxna4/efna2a/dscama/apc/robo2/pcdh17/hdc/anos1a/trioa/s1pr1/lhx9/epha7/ccl38.6/rtn4r/cyfip2/dpysl3/draxinb/slit1b/robo3/usp33/aanat2/dock4b/dzip1/trim33/ccl34b.4/sema5a/chad/vegfd/dock4/robo1/tnr/tenm4/epha4l/kif5c/islr2/nptnb/lhx2b/kalrna/xcr1a.1/ccl44/ephb2b/rac3a/gap43/lhx1a/atl1/nkx2.2a/arhgef25b/dscamb/triob/slit3/cxcl8a/dpysl5a/cxcl8b.3/chst11/fzd3a/kif5aa/si:dkey-12h9.6/robo4/efnb3b/vax1/stat3/zgc:77784/sema4c/si:ch211-113g11.6/rac1l/anxa1c/ntn1b/scn8aa/ccl35.1/cxcl8b.1/ncam1b/cntn3a.1/nrxn2a/efna3a/dpysl5b/si:ch211-241e1.3/plxna3/l1camb/plxna1a/nrcama/sema6ba/il1b/sema3b/cxcl18b |
| GO:0098953 | receptor diffusion trapping | -1.77120547755989 | 0.0221781634063151 | cacng8b/gphnb/cacng5a/gphna/cacng7b/cacng2b/cacng3b/cacng2a/cacng8a |
| GO:0098970 | postsynaptic neurotransmitter receptor diffusion trapping | -1.77120547755989 | 0.0221781634063151 | cacng8b/gphnb/cacng5a/gphna/cacng7b/cacng2b/cacng3b/cacng2a/cacng8a |
| GO:0099628 | neurotransmitter receptor diffusion trapping | -1.77120547755989 | 0.0221781634063151 | cacng8b/gphnb/cacng5a/gphna/cacng7b/cacng2b/cacng3b/cacng2a/cacng8a |
| GO:0071867 | response to monoamine | -1.76002500271274 | 0.0460556112254136 | htr1aa/htr1d/gnao1b/nsg2/gnao1a/drd2a/htr1fa/gnb5a |
| GO:0071869 | response to catecholamine | -1.76002500271274 | 0.0460556112254136 | htr1aa/htr1d/gnao1b/nsg2/gnao1a/drd2a/htr1fa/gnb5a |
| GO:0051960 | regulation of nervous system development | -1.75557639124564 | 0.00248666027456893 | nlk1/si:zfos-741a10.3/plxnb1b/hes6/neurod6a/adnpb/sema3fa/btbd6a/her12/dbn1/si:ch211-129c21.1/lef1/ulk2/dbnlb/ephb3a/dhrs12la/slit2/ctbp2a/sema3bl/plxna4/clstn2b/bhlhe40/robo2/epha4a/neurod4/gata3/her13/gbx2/tnfrsf21/sema5a/foxo3a/tnr/tenm4/slitrk4/caprin2/zbtb16a/cdkl5/epha4b/clstn1/nsmfb/slit3/clstn2a/elavl3/stat3/sema4c/si:ch211-113g11.6/gbx1/grna/bdnf/clstn3/slitrk2/dmbx1a/plxna3/l1camb/mmp9/plxna1a/sema6ba/sema3b |
| GO:0097120 | receptor localization to synapse | -1.75544036908873 | 0.0351674145187552 | dlg1b/cacng8b/grip2b/gphnb/cacng5a/stx3a/anks1b/kif5c/gphna/cacng7b/kif5aa/cacng2b/cacng3b/stx1b/cacng2a/rab44/cacng8a |
| GO:0031532 | actin cytoskeleton reorganization | -1.74088656656218 | 0.0402190977319739 | klf6a/anxa1a/baiap2a/sipa1l1/mink1/nedd9/thsd7bb/carmil2/parvg/carmil3/plek/thsd7ba/anxa1c/plek2/phactr1/thsd7aa |
| GO:0040013 | negative regulation of locomotion | -1.73929262815316 | 0.018759055936431 | sema3fa/lrch1/si:ch211-129c21.1/lbx2/prox1a/miip/lef1/phf8/srgap3/sema3bl/robo2/anos1a/rap2b/sema5a/srgap1b/sinhcaf/arhgap4b/sema4c/si:ch211-113g11.6/rtn4rl2a/sema6ba/sema3b |
| GO:0031110 | regulation of microtubule polymerization or depolymerization | -1.72739295421889 | 0.0437181181328161 | ska3/cdk5rap2/stmn1b/map1sa/camsap2a/camsap1b/slain2/map1ab/traf3ip1/zgc:66447/clasp2/mapre3b/stmn4/apc/map1b/stmn2a/map1sb/stmn2b/stmn3/clip1a/map1aa/camsap1a/stmn4l/apc2 |
| GO:0040011 | locomotion | -1.71881331498307 | 4.74228301214316e-06 | plcg1/unc5b/nptna/lrch1/vegfc/tnfrsf1a/si:ch211-129c21.1/lbx2/cntn4/cdh2/prox1a/kalrnb/miip/maco1a/lef1/tdgf1/foxj1a/ptk7a/efna1b/ephb3a/slc35b2/gnb1a/rbpms2b/sst7/slkb/rab33ba/nrp2a/fezf2/phf8/slit2/tenm3/maco1b/srgap3/nfasca/sim1a/adcy8/kif1b/arnt2/sema3bl/myd88/plxna4/efna2a/dscama/apc/robo2/pcdh17/hdc/asap1a/arhgef39/anos1a/trioa/s1pr1/lhx9/epha7/hace1/peak1/ccl38.6/rtn4r/cyfip2/dpysl3/draxinb/gata3/slit1b/robo3/usp33/tek/aanat2/dock4b/dzip1/rap2b/trim33/ccl34b.4/sema5a/chad/vegfd/pdgfbb/dock4/robo1/tnr/tenm4/epha4l/kif5c/islr2/nptnb/htr1d/lhx2b/kalrna/xcr1a.1/ccl44/ephb2b/sst1.1/srgap1b/rac3a/gap43/lhx1a/atl1/nkx2.2a/arhgef25b/dscamb/amot/triob/nsmfb/slit3/cxcl8a/dpysl5a/cxcl8b.3/chst11/fzd3a/kif5aa/si:dkey-12h9.6/sinhcaf/robo4/efnb3b/vax1/arhgap4b/stat3/dock10/zgc:77784/sema4c/adam8a/si:ch211-113g11.6/rac1l/anxa1c/ntn1b/scn8aa/ccl35.1/cxcl8b.1/mdga2a/zgc:153631/asap1b/ncam1b/cntn3a.1/nrxn2a/klhl40b/efna3a/chrm4a/dpysl5b/si:ch211-241e1.3/plxna3/l1camb/plxna1a/nrcama/rtn4rl2a/sema6ba/il1b/sema3b/cxcl18b |
| GO:0006959 | humoral immune response | -1.69323952530372 | 0.0594675488145613 | c4/tnfrsf21/c1qa/tfe3a/cfb/cxcl8b.3/c6/susd4/c7b/c4b/c3a.2/itln2/cxcl18b |
| GO:0007420 | brain development | -1.67227058783588 | 2.31796694375566e-05 | her12/wnt3/macroh2a2/marcksa/kctd12.1/daam1a/pknox1.1/tgif1/ctnna2/cntn4/cdh2/adcyap1b/prox1a/zfhx3b/prnprs3/sbno2a/lef1/tdgf1/ulk2/tsc22d3/bag6l/pou3f3a/atg4da/creb1a/pafah1b1a/birc6-as2/sox14/foxb1a/nsfa/zfhx3a/kdm6al/slc2a3a/cep290/kctd13/fezf2/phf8/six3a/nfasca/sim1a/pax6b/arnt2/raraa/grhl3/tbr1b/kirrel3l/cars2/ptprsa/kdm7ab/nr4a2b/ralgapa1/apc/epha4a/hdc/anos1a/lhx9/neurod4/pcdh10b/pbx1a/mvp/six3b/mdkb/si:ch73-119p20.1/gbx2/dzip1/lgi1a/pbx1b/epb41l5/auts2a/pbx3b/pcare1/hecw2a/znf503/meis2a/wnt5b/lhx6a/irx7/lhx2b/brf1b/rbfox1/tcf7l2/nkx2.1/pcdh19/srrm4/lhx1a/asphd1/pou4f2/prmt8b/tbxta/aatkb/rtn1b/sp5l/nr2e3/tbcd/has2/lhx5/fzd3a/pou3f2b/nr4a2a/efnb3b/ndrg4/vax1/pax2a/git2b/gbx1/ntn1b/pou3f2a/bdnf/atp1a3a/sez6l2/mdga2a/rx3/dmbx1a/otpb/six6a/cdk5r2a/arxa/nkx2.4a/syngap1b/gnrh2/l1camb/cnnm2a/otpa/kcnb1/nrcama/gper1/bicdl1 |
| GO:0060322 | head development | -1.66225293057786 | 4.42238345198842e-05 | her12/wnt3/macroh2a2/marcksa/kctd12.1/daam1a/pknox1.1/tgif1/ctnna2/cldn7a/cntn4/cdh2/adcyap1b/prox1a/zfhx3b/prnprs3/sbno2a/lef1/tdgf1/ulk2/tsc22d3/bag6l/pou3f3a/atg4da/creb1a/pafah1b1a/birc6-as2/sox14/foxb1a/nsfa/zfhx3a/kdm6al/slc2a3a/cep290/kctd13/fezf2/phf8/six3a/nfasca/sim1a/pax6b/arnt2/raraa/grhl3/tbr1b/kirrel3l/cars2/ptprsa/kdm7ab/nr4a2b/ralgapa1/apc/epha4a/hdc/wdr11/anos1a/lhx9/neurod4/pcdh10b/pbx1a/mvp/six3b/mdkb/si:ch73-119p20.1/gbx2/dzip1/lgi1a/pbx1b/epb41l5/auts2a/pbx3b/pcare1/hecw2a/znf503/meis2a/wnt5b/lhx6a/irx7/lhx2b/brf1b/rbfox1/tcf7l2/nkx2.1/pcdh19/srrm4/lhx1a/asphd1/pou4f2/prmt8b/tbxta/aatkb/rtn1b/sp5l/nr2e3/tbcd/has2/lhx5/fzd3a/pou3f2b/nr4a2a/efnb3b/ndrg4/vax1/pax2a/git2b/gbx1/ntn1b/chd5/pou3f2a/bdnf/atp1a3a/sez6l2/mdga2a/specc1/rx3/dmbx1a/otpb/six6a/cdk5r2a/arxa/nkx2.4a/syngap1b/gnrh2/l1camb/cnnm2a/otpa/kcnb1/nrcama/gper1/bicdl1 |
| GO:0045732 | positive regulation of protein catabolic process | -1.66175262841073 | 0.0607582432020156 | oaz1a/csnk1db/tmem259/oaz2b/socs5b/prkn/apc/rnf217/si:ch211-278j3.3/trib1/fzr1b/rnf144ab/csnk1e/adam8a/klhl40b/apc2/rnf144aa |
| GO:1901797 | negative regulation of signal transduction by p53 class mediator | -1.65413798029984 | 0.0829144295829561 | cd74b/cd74a/bcor/vax1/bcl6aa |
| GO:0006688 | glycosphingolipid biosynthetic process | -1.64952029857802 | 0.097836707744085 | b4galt6/b4galt3/st6galnac3 |
| GO:0031644 | regulation of nervous system process | -1.64521543139582 | 0.0906720571933235 | dlgap1a/tnfrsf21/tenm4/dlgap4b/si:dkey-174m14.3/kcnab2b/dlgap3/neto1l |
| GO:0060560 | developmental growth involved in morphogenesis | -1.64279725335724 | 0.0128583805736124 | plxnb1b/tmem108/pik3cd/mycbp2/spag6/cbsb/smad2/pik3ca/zeb2a/mapk8ip3/sema3fa/dbn1/si:ch211-129c21.1/ubap1/cdh2/trarg1a/tmem67/ulk2/ptk7a/dbnlb/creb1a/fus/isl1a/slit2/ppp1cb/spg11/ret/kif1b/sema3bl/bbs1/robo2/pcdh17/casp9/epha4a/robo3/appb/sema5a/wnt5b/myo9ab/cdkl5/epha4b/clstn1/tbxta/zeb2b/tmem59l/myo10l3/nsmfb/slit3/stat3/sema4c/si:ch211-113g11.6/myo10l1/ntn1b/grna/bdnf/nrn1la/nkx6.1/nrxn2a/thsd7aa/plxna3/l1camb/sema6ba/sema3b |
| GO:0016049 | cell growth | -1.63666909504427 | 0.0141269580867656 | plxnb1b/tmem108/pik3cd/mycbp2/spag6/ewsr1a/pik3ca/mapk8ip3/sema3fa/dbn1/si:ch211-129c21.1/ubap1/trarg1a/ulk2/dbnlb/supv3l1/dnph1/fus/isl1a/slit2/spg11/ret/kif1b/sema3bl/robo2/pcdh17/casp9/epha4a/robo3/appb/sema5a/wnt5b/myo9ab/cdkl5/epha4b/clstn1/tmem59l/myo10l3/nsmfb/slit3/sema4c/si:ch211-113g11.6/myo10l1/ntn1b/epb41l3a/grna/bdnf/nrn1la/nkx6.1/nrxn2a/thsd7aa/plxna3/l1camb/sema6ba/sema3b |
| GO:0034643 | establishment of mitochondrion localization. microtubule-mediated | -1.63631487834659 | 0.0931198545493737 | rhot1b/trak1a/sybu/kif5aa/rhot2 |
| GO:0047497 | mitochondrion transport along microtubule | -1.63631487834659 | 0.0931198545493737 | rhot1b/trak1a/sybu/kif5aa/rhot2 |
| GO:0051654 | establishment of mitochondrion localization | -1.63631487834659 | 0.0931198545493737 | rhot1b/trak1a/sybu/kif5aa/rhot2 |
| GO:0032102 | negative regulation of response to external stimulus | -1.63570938930065 | 0.0540332674423811 | sema3bl/gramd4b/robo2/epha4a/il13/nr1h5/sema5a/cart3/ptger4c/irak3/epha4b/gramd4a/ptger4a/sema4c/si:ch211-113g11.6/susd4/grna/serpine1/sc:d189/sema6ba/sema3b |
| GO:0048885 | neuromast deposition | -1.63446228315307 | 0.0960340406478379 | lbx2/prox1a/lef1/phf8/anos1a/rtn4rl2a |
| GO:0071407 | cellular response to organic cyclic compound | -1.62675574144756 | 0.0543657683428525 | abhd2b/htr1aa/htr1d/gnao1b/insig1/esr2b/cxcl8a/nsg2/sstr1a/gnao1a/anxa1c/drd2a/htr1fa/gnb5a/gper1 |
| GO:0022603 | regulation of anatomical structure morphogenesis | -1.62242060545465 | 0.00196692398869123 | plcd4a/vegfab/stab1/hif1ab/amotl2b/sema6d/ptenb/alcama/sema6e/ptk2ab/thbs2b/brwd1/rspo3/tspan18a/gata2a/zeb1b/plxnb1b/pik3cd/disc1/mia3/anxa1a/fn1b/arhgap35a/pik3ca/flot2a/sema3fa/camsap1b/plcg1/unc5b/vegfc/dbn1/aplnrb/kctd12.1/si:ch211-129c21.1/rhogd/zeb1a/lbx2/cdh2/fmnl2b/ulk2/elmo1/marchf5/ptk7a/dbnlb/dab2ipb/lurap1/ephb3a/mffa/sipa1l1/tnika/vat1/palm1b/slit2/mief2/rufy3/hecw1b/pak1/sema3bl/plxna4/nedd4l/arhgap35b/robo2/casp9/epha4a/s1pr1/msnb/tanc2a/phip/sema5a/vegfd/tenm4/hecw2a/wnt5b/caprin2/cdkl5/epha4b/myo10/clstn1/rac3a/lrrc4ca/lhx1a/nsmfb/slit3/basp1/stat3/prkd1/sema4c/si:ch211-113g11.6/rac1l/gbx1/anxa1c/lrrc4cb/hif1aa/bdnf/serpine1/adgrb3/scg2b/gdf3/adgrb1a/thbs1a/dab2ipa/plxna3/l1camb/eve1/plxna1a/adgrb1b/sema6ba/sema3b |
| GO:0000381 | regulation of alternative mRNA splicing. via spliceosome | -1.62121505428978 | 0.0619371464051369 | hnrnpa0a/rbm38/rbm25a/celf2/rbfox3a/rbfox3b/celf6/rbfox1l/rbfox1/srrm4/nova2/celf3a/khdrbs2/celf5a/celf4 |
| GO:0001933 | negative regulation of protein phosphorylation | -1.61410605566215 | 0.0735990755145654 | spry4/aida/dtnbp1b/ptprc/ldlrad4a/camk2n2/cbl/camk2n1a/cblc/prkn/sh3bp5la/sh3bp5b/si:dkey-175m17.7/ibtk/camk2n1/irak3/inpp5ka/cadm4/inpp5kb/smcr8b |
| GO:0008361 | regulation of cell size | -1.59323427771673 | 0.0658674351521172 | arhgap35a/pik3ca/sema3fa/arhgap5/dbn1/si:ch211-129c21.1/ulk2/dbnlb/zgc:153039/slit2/sema3bl/arhgap35b/robo2/casp9/epha4a/sema5a/piezo1/cdkl5/epha4b/clstn1/nsmfb/slit3/si:ch211-178n15.1/stat3/slc12a5b/sema4c/si:ch211-113g11.6/ntn1b/slc12a5a/bdnf/nkx6.1/plxna3/l1camb/sema6ba/sema3b |
| GO:0043087 | regulation of GTPase activity | -1.57086875382947 | 0.0297635976408028 | tbc1d9/rapgef1a/flcn/sipa1l2/rasgef1ba/iqsec1b/ralgapa2/rasa4/gpr137ba/rgl1/nf1a/usp6nl/adcyap1b/vav2/agap1/rapgef5b/rapgef2a/si:dkey-91m11.5/dab2ipb/rgl3a/ephb3a/rapgef2b/dock7/rapgef4b/sipa1l1/gapvd1/rasal2/plxna4/chn1/ralgapa1/ccpg1/asap1a/rasgef1bb/ccl38.6/bcr/rgs7a/ccl34b.4/si:ch211-288d18.1/evi5l/tbc1d30/elmod1/vav3b/myo9ab/ccl44/ek1/rgs9b/sgsm3/syngap1a/arap3/rap1gapa/unm_sa1614/ralgps1/dock10/agap2/ccl35.1/asap1b/rasgrf2b/rgs11/syngap1b/dab2ipa/plxna3/rasgrf1/plxna1a/arhgap25 |
| GO:0002252 | immune effector process | -1.52782636998837 | 0.0607582432020156 | cd74b/c4/c1qa/ticam1/cd74a/cfb/cadm1b/cxcl8a/c6/si:ch211-196f2.6/susd4/anxa1c/zgc:153631/c7b/c4b/zgc:123107/c3a.2/csf2rb/mmp9/mhc2d4b |
| GO:0001755 | neural crest cell migration | -1.51280994298295 | 0.0960340406478379 | hif1ab/sema6d/pax6a/med13b/sema6e/lrp5/disc1/nrp2b/erbb2/pdgfrb/sema3fa/radil/si:ch211-129c21.1/cdh2/lef1/dmxl2/pax3a/nrp2a/fscn1a/pak1/ret/pax6b/sema3bl/mmp17b/wdr11/sema5a/zeb2b/sh3pxd2aa/sema4c/si:ch211-113g11.6/rx3/foxi1/sema6ba/sema3b |
| GO:0051240 | positive regulation of multicellular organismal process | -1.44230693231483 | 0.0657742225763037 | gata2a/slc27a1b/nlk1/si:zfos-741a10.3/plxnb1b/anxa1a/avpr1ab/neurod6a/tnfsf13b/nr2c2/cactin/apln/tlr5b/olfm1a/vegfc/dbn1/aplnrb/lef1/elmo1/dbnlb/lurap1/ephb3a/npy/tapt1b/dhrs12la/loxl2a/zgc:172120/plxna4/clstn2b/cd74b/neurod4/gata3/aanat2/vegfd/enpp5/tenm4/ticam1/slitrk4/caprin2/zbtb16a/cdkl5/clstn1/cd74a/lhx1a/vtcn1/nsmfb/clstn2a/basp1/stat3/si:ch211-196f2.6/anxa1c/grna/bdnf/zbtb16b/clstn3/slitrk2/zgc:153631/adrb1/npvf/plxna3/l1camb/zgc:123107/plxna1a/panx2/gper1/mhc2d4b |
| GO:0030334 | regulation of cell migration | -1.43156853115565 | 0.0696903189146766 | plcg1/lrch1/vegfc/si:ch211-129c21.1/lbx2/prox1a/miip/lef1/gnb1a/sst7/slkb/phf8/srgap3/sema3bl/plxna4/asap1a/arhgef39/anos1a/s1pr1/hace1/peak1/dpysl3/gata3/tek/rap2b/sema5a/vegfd/pdgfbb/sst1.1/srgap1b/amot/nsmfb/sinhcaf/robo4/arhgap4b/dock10/sema4c/adam8a/si:ch211-113g11.6/anxa1c/mdga2a/zgc:153631/asap1b/si:ch211-241e1.3/plxna3/plxna1a/rtn4rl2a/sema6ba/sema3b |
| GO:0009719 | response to endogenous stimulus | -1.32546647522578 | 0.0666420813824336 | tbx16/irs1/skia/igfbp3/pik3r3a/gsk3ba/rbm14b/gna11b/slc39a14/slc27a1b/esr2a/zyx/smad7/tll1/ltbp3/tacr1a/spry4/dusp6/trim71/anxa1a/avpr1ab/nrp2b/nr1d1/smad2/grb10a/flcn/smurf1/zeb2a/asic4b/fcho1/skib/rgmb/fgfbp1a/irs2a/drd4-rs/nptna/vegfc/fstl4/si:rp71-46j2.7/ldb2a/timp2a/tgif1/ldlrad4a/caly/fstl3/castor2/ppp4cb/zdhhc13/tsc22d3/il13ra1/dab2ipb/lifrb/golm1/rerea/chrm2a/slc2a3a/nrp2a/pnpla6/asip1/tcf21/grem2b/pik3r3b/raraa/thrb/zdhhc17/bmpr1ab/smad3b/stat6/gpr22a/nr4a2b/cpeb4a/chrng/glra1/sall1a/gata3/bmp15/dzip1/fabp4b/phip/crhbp/sp9/snx25/inhbab/vegfd/fgf12a/abhd2b/htr1aa/bmpr2a/scube1/ptger4c/ccbe1/stat1b/nrros/htr1d/lemd3/gnao1b/inpp5ka/smad6b/insig1/esr2b/ptger4a/atl1/tbxta/inhbaa/asic2/timp4.3/has2/adgrl3.1/kcp/grem1a/nr4a2a/bmpr2b/stat3/nsg2/wbp2/gnao1a/stxbp4/anxa1c/nr4a3/bdnf/elapor2b/inpp5kb/ntrk3a/frs3/irs2b/gdf3/drd2a/ntrk3b/htr1fa/chrm4a/trh/dio2/dab2ipa/gnb5a/rxfp1/c3a.2/sostdc1b/mmp9/timp2b/hpxa |
